# Range-wide phylogeography, population genomics, and demography of three widespread *Ara* macaws (Psittacidae)

**DOI:** 10.64898/2026.05.22.726827

**Authors:** Jaime G. Morin-Lagos, Vanessa C. Bieker, Ole K. Tørresen, Taylor Hains, Maanasa Raghavan, Letty Salinas, Cristina Y. Miyaki, Shannon J. Hackett, John Bates, Michael D. Martin

## Abstract

Macaws of the genus *Ara* comprise eight extant species distributed throughout the Neotropics. Among them, four have broad geographic ranges, yet little is known about the evolutionary history and demographic processes that shaped their genomic variation and present-day distributions. This is particularly relevant because, although these wide-ranging macaws are classified as Least Concern by the IUCN, many of their populations are declining due to habitat fragmentation, illegal trade, and climate change. Here, we used nuclear and mitochondrial genomic data to characterize the evolutionary relationships, population structure, genetic diversity, and demographic histories of three widely distributed species (*A. ararauna*, *A. chloropterus*, and *A. severus*) across their geographic distributions. We identified two main populations within *Ara severus*, and this species showed the highest heterozygosity levels among the three species. In *A. ararauna* and *A. chloropterus*, we observed four main genetic clusters corresponding to two populations in the Amazon rainforest biome and and two populations in the Cerrado savanna biome. Cerrado populations in both species exhibited markedly reduced heterozygosity and elevated inbreeding relative to Amazonian populations, consistent with smaller effective population sizes and increased isolation. Genome-wide scans suggested that genetic drift and divergent demographic histories played a predominant role in driving the strong differentiation between Amazon and Cerrado in these two species. Nevertheless, we detected two candidate genes, *NALCN* and *RBBP6*, with convergent selection signals across *A. ararauna* and *A. chloropterus*, suggesting possible local adaptation to the Cerrado biome.

## Introduction

The genus *Ara* (Psittacidae) comprises eight extant species of macaw parrots distributed throughout the Neotropics, with half of them widely distributed across the region (*Ara ararauna*, *A. chloropterus*, *A. severus*, and *A. macao*), while the others (*A. militaris*, *A. glaucogularis*, *A. rubrogenys*, and *A. ambiguus*) occupy smaller ranges. These birds are characterized by their large body size, long tails, strikingly colored plumage, and extensive bare facial skin around the eyes. In addition, they are known for their high cognitive abilities (Torres Ortiz et al., 2022), exhibiting complex social behaviors, the ability to imitate human sounds, and the capacity to make facial expressions associated with social communication (Bertin et al., 2018, 2023). Thus, *Ara* species are among the most charismatic bird species globally, highly valued in the pet trade, and frequently depicted in movies, advertisements, and broader popular culture. Moreover, the archaeological record suggests that a close relationship between humans and macaws dates back to pre-Hispanic times (Capriles et al., 2021; George et al., 2018).

Humans have significantly contributed to the loss and fragmentation of macaw habitats, resulting in dramatic population declines, particularly over the last 50 years (Herzog et al., 2021, 2023), which is also exacerbated by climate change and illegal capture and trafficking (Tella & Hiraldo, 2014). As a result, the four *Ara* species with restricted ranges are currently considered by the IUCN as threatened to different degrees, while the widely distributed *Ara* macaws are regarded as not threatened. Despite this, some studies indicate that many populations of widespread *Ara* species are experiencing rapid declines in population size (Ragusa-Netto, 2024; Vaughan et al., 2005). Particularly, *A. chloropterus* has undergone significant population reductions within the Cerrado biome (Ragusa-Netto, 2024)).

Comprehensive knowledge of genetic diversity within species has the potential to provide valuable information for conservation purposes (Cassin-Sackett et al., 2019; Gonçalves et al., 2015; Olah et al., 2021; Vilaça et al., 2024). Classical single-locus genetic markers, particularly mitochondrial ones such as cytochrome-b (CYTB), and microsatellites have been used for this kind of studies in *Ara* macaws, and have been mostly applied to narrowly-distributed species such as *A. rubrogenys* (Blanco et al., 2021), *A. glaucogularis* (C. I. Campos et al., 2021), and *A. militaris* (F. A. Rivera-Ortíz et al., 2017, 2023) to explore the intra-specific genetic diversity and population structure within these species. This type of studies can also support or challenge putative subspecies as in *A. militaris* (Eberhard et al., 2015; F. A. Rivera-Ortíz et al., 2023), and *A. macao* (Schmidt et al., 2020). Furthermore, the genetic information obtained can be directly transferred to conservation initiatives. For example they have been used to propose the creation of conservation units (Blanco et al., 2021; F. A. Rivera-Ortíz et al., 2017); to identify the species and population identity of individuals from the illegal trade market (Fernandes & Caparroz, 2013; Gonçalves et al., 2015; F. Rivera-Ortíz et al., 2021); and to select captive individuals suitable for reintroduction efforts (Estrada, 2014).

Among the four widespread *Ara* macaw species, only *A. macao* has been studied across its entire range (Schmidt et al., 2020). However, while this study based on the single mitochondrial marker CYTB detected genetic structure in Central and North America, it did not detect significant genetic differentiation among South American populations. In contrast, other studies using mitochondrial markers have detected some degree of differentiation between South American populations in smaller ranged parrot species (Blanco et al., 2021; Masello et al., 2011). Similarly, a regional study of *A. ararauna* in Brazil using the mitochondrial control region and microsatellites revealed genetic differentiation among populations (Caparroz et al., 2009). However, no comprehensive population-level genetic analyses covering the full range have yet been conducted for *A. ararauna* or the two other widespread *Ara* species (*A. chloropterus*, and *A. severus*), limiting our understanding of genetic structure in these macaws across the South American subcontinent.

Defining population boundaries in birds is particularly challenging, especially for large, long-distance flying species such as macaws. According to the climatic variability hypothesis, species in the Neotropics may be adapted to a narrow range of temperature because they experience relatively stable climatic conditions throughout the year (Janzen, 1967). Genetic evidence has revealed population structure in every *Ara* macaw species that has been studied to date, which could be associated with dispersal barriers, and habitat heterogeneity, particularly along mountain ranges (Remsen et al., 2025; F. A. Rivera-Ortíz et al., 2017), and even habitat preference (Caparroz et al., 2009). Accordingly, *A. macao*, *A. militaris*, and *A. ambiguus* currently comprise two or more recognized subspecies (Remsen et al., 2025). In contrast, the lack of genomic studies in *A. ararauna*, *A. chloropterus*, and *A. severus* is probably a major reason that they are still considered monotypic despite their comparatively broader ranges (Collar et al., 2020a, 2020b, 2020c).

The advent of high-throughput sequencing technologies has enabled the incorporation of whole-genome sequence data into modern studies of avian population genetics, allowing the investigation of population structuring and dynamics at finer spatial and temporal scales (Elgvin et al., 2017; Kersten et al., 2021; Vianna et al., 2020; Vilaça et al., 2024). Such genomic information has been already studied for its implementation to enhance further conservation initiatives. For instance, it has been shown that genomic diversity indices can be directly related to IUCN conservation categories (Jeon et al., 2024). However, despite some efforts to generate genomic data in parrots (Hains et al., 2022; Olah et al., 2021), it remains underemployed in parrot population studies. To date, SNP-based population studies have been conducted in the kakapo (*Strigops habroptilus*) (Dussex et al., 2021; Guhlin et al., 2023), and the Norfolk parakeet (Gautschi et al., 2024), both endangered endemic species of New Zealand and the Norfolk island respectively. In the Neotropics some notable efforts have been conducted on the Hyacinth macaw (*Anodorhynchus hyacinthinus*) (Vilaça et al., 2024), the Sun Parakeet (*Aratinga solstitialis*) (Spitzer et al., 2020), and the scarlet macaw (*Ara macao*) (Aardema et al., 2023), although the latter was restricted in sampling size and scope, focusing only on Central American populations.

In the past, subtle differences in body size have led to the proposal of a subspecies within *A. severus* (*A. s. castaneifrons;* Lafresnaye, 1847) which has never been tackled using genetic data. Moreover, *A. ararauna* and *A. chloropterus* are not only widely ranged in South America, but notably they are the only *Ara* species with populations in the Cerrado biome, the world’s most diverse savanna. This area is characterized by seasonal dry periods, in contrast to the Amazon, which is characterized by tropical rainforests (Antonelli et al., 2018). Heterogeneity of habitats can significantly impact population dynamics and population structure across bird species (Bocalini et al., 2021). For instance, a recent genomic study in the gray-breasted sabrewing (*Campylopterus largipennis*) species complex revealed deep genetic structure separating species from both biomes (Oliveira et al., 2024), evidencing the role of these biomes in population divergence. However, no further studies focusing on bird species with ranges across both biomes have been carried out using genetic data. In addition, the Cerrado biome is currently experiencing significant habitat loss due to high rates of land conversion, threatening local biodiversity (Vieira-Alencar et al., 2023), such as the documented recent population declines in *A. chloropterus* (Ragusa-Netto, 2024). This warrants further research to assess whether habitat heterogeneity and anthropogenic pressures are shaping genetic structure within these species.

In this study, we sought to perform phylogenomic and population genomic analyses to benefit macaw conservation, generating whole-genome sequence data of the three widespread *Ara* species that are endemic to South America (*A. chloropterus, A. ararauna, A. severus*), encompassing their entire range. We assessed population structure and genetic diversity, and identified potential conservation units. In addition to assessing present-day population structure, we also inferred demographic history to identify past population expansions and contractions, including potential bottlenecks, that may have shaped observed genetic patterns, particularly in response to climatic fluctuations during the Last Glacial period. Moreover, this is the first comprehensive intraspecific study done in these three highly charismatic bird species, providing valuable information that has implications for the systematics of these species and should be considered in conservation efforts. Finally, we compared the genomic results with mitogenome-based phylogenies finding population structure that cannot be detected using mitochondrial data alone. Complete mitochondrial genome data captures partially the broad-scale geographic patterns observed in the nuclear data, especially in *A. chloropterus* and *A. severus*, suggesting that they may offer a cost-effective alternative for preliminary assessments in conservation contexts. We therefore encourage broader application of integrative approaches in *Ara* macaw research.

## Material & Methods

### Sample acquisition

We collected feathers from wild individuals of the three *Ara* species in the Peruvian Amazon under the research and sampling permit issued by the local authorities (SERFOR N° AUT-IFS-2022-042). We deposited all feathers in the Ornithology Collection of the Natural History Museum of Universidad Nacional Mayor de San Marcos (MUSM, Peru) from which we took a subsample for DNA extraction. We also obtained toe pad and feather samples from the MUSM historical feather and skin collections. Additionally, we obtained blood samples from the Laboratório de Genética e Evolução Molecular de Aves (LGEMA, Brazil), skin samples from ornithological collections in the United States (Field Museum, Louisiana State University, Museum of Vertebrate Zoology, and Yale Peabody Museum), and Europe (NTNU University Museum, Norway; National Museum of Natural History, France; and Naturalis Biodiversity Center, Netherlands). A subset of *A. severus* samples was processed from DNA extraction through library preparation at the University of Chicago (ST3). The rest of the samples were sent to the NTNU University Museum in Trondheim, Norway for molecular analysis under CITES permits PE005039/SP, PE005040/SP, and PE005045/SP for Peruvian samples, CITES permit 24BR048160/DF for Brazilian samples, and CITES permit 22US701766/9 for US samples as part of exchanges between CITES COSE-registered institutions. We processed samples from a total of 92 *Ara ararauna*, 90 *A. chloropterus*, and 57 *A. severus* individuals from across their natural ranges (Tables S1-3).

### DNA extraction, library preparation, and sequencing

Samples from individuals collected before 1960 were considered historical and as such were processed in dedicated, positively pressurized ancient DNA laboratory facilities either at the Department of Human Genetics at the University of Chicago or the NTNU University Museum. Before DNA extraction, we pre-treated toe pads and feathers as described in Zhang et al. (2021). For historical samples from U.S. institutions, we first lysed each sample in 600 µL of extraction solution containing 532 µl HPLC-grade water, 160 µl of extraction buffer (5X TE Buffer, 95 mM EDTA, and 50 mM NaCL), 80 µl SDS (10%), 35 µl proteinase K (10 mg/mL), and 20 µl DTT (400 mg/mL). For other historical samples, we utilized the extraction solution used in Dabney et al. (2013). All samples were incubated at 55 °C for 3–5 days until fully digested, adding more proteinase K and DTT as needed. DNA was purified using the MinElute PCR Purification kit (Qiagen) following the manufacturer’s instructions. For contemporary samples and feathers, we performed genomic DNA extraction using the DNeasy Blood and Tissue kit (QIAGEN, California) according to the manufacturer’s instructions, except that for feather samples we added 20 µL of 1 M DTT during the lysis step. We quantified the DNA concentration using a Qubit 2.0 fluorometer (Thermo Fisher Scientific, Indiana, USA) with the dsDNA HS (High Sensitivity) Assay kit and then evaluated the DNA integrity of historical samples with 2% agarose gel electrophoresis or TapeStation Genomic DNA ScreenTape Analysis (Agilent).

Contemporary and selected historical samples exhibiting high molecular weight DNA processed at NTNU were sheared to a mean length of 500 bp using a Covaris M220 focused-ultrasonicator. We constructed blunt-end Illumina libraries using the single-tube BEST protocol (Carøe et al., 2018). Sample-specific, dual-indexing barcodes were incorporated into each library during the indexing PCR using custom index primers. For modern samples, indexing PCR was carried out in 100-µl reactions with 10 µl library template, 0.25 mM each dNTP, 0.25 µM each index primer, 1 µl Herculase II Fusion DNA polymerase, 20 µl 5 × Herculase II Reaction Buffer, and 63 µl molecular biology-grade water, under the following conditions: 95°C for 3 min, an appropriate number of cycles of 95°C for 20 s, 60°C for 20 s, 72°C for 40 s and a final extension of 72°C for 5 min. For historical samples, indexing PCR was carried out in 100-µl reactions with 10 µl library template, 0.2 mM each dNTP, 0.2 µM each index primer, 0.05 U/µl AmpliTaq Gold polymerase, 1X AmpliTaq Gold Buffer, 2.5 mM MgCl_2_, and 0.4 mg/ml bovine serum albumin under the following conditions: 95°C for 10 min, appropriate number of cycles of 95°C for 30 s, 60°C for 1 min, 72°C for 45 s and a final extension of 72°C for 5 min. The optimal number of index PCR cycles was previously estimated using quantitative PCR (qPCR). We included multiple negative controls (blanks) throughout molecular procedures to monitor potential environmental contamination during DNA extraction, library building, qPCR, and index PCR. The amplified libraries were purified using AMPure XP beads (Beckman Coulter), with a bead-to-sample ratio of 1:1 for modern samples and 1.2:1 for historical samples. DNA then was eluted in 33 µl EB buffer (Qiagen). We estimated the molarity and fragment distribution of the amplified libraries using the High Sensitivity D1000 ScreenTape assay (Agilent) to create equimolar sequencing pools.

Historical DNA extracts of *A. severus* processed at the University of Chicago were built into libraries using the KAPA HyperPrep Library Kit, following the manufacturer’s protocol using the iTru indexing strategy (Illumina). Libraries were then quantified using a High Sensitivity D1000 ScreenTape assay on a 4200 TapeStation System instrument (Agilent). All samples were sequenced by Novogene Europe on the Illumina NovaSeq platform in 150-bp paired-end format.

### Genomic data processing

We used the PALEOMIX pipeline v1.2.13 (Schubert et al., 2014) to process raw fastq files. Adapter trimming and paired-end read collapsing were performed using AdapterRemoval v2.3.1 (Schubert et al., 2016)), as implemented in PALEOMIX (min overlap of 11 bp), and only reads ≥ 25 bp were retained. Trimmed reads were aligned to the *Ara ararauna* nuclear genome assembly bAraAra1.hap1 (GCA_028858755.1, unpublished) using the Burrows-Wheeler Aligner implemented in BWA v0.7.15 (Li & Durbin, 2009), with the ‘backtrack’ algorithm and keeping only those aligned reads with a Phred-scaled mapping quality (mapQ) ≥25. The endogenous DNA content, sequencing depth, and clonality were estimated using PALEOMIX, based on the raw sequences retained after trimming and quality filtering. Samples with a mean coverage ≥0.6X after mapQ filtering were included in the whole-genome analysis. Using R v4.4.3, we performed a Welch two-sample t-test to evaluate the difference in endogenous DNA content between contemporary and historical specimens (collected before 1960) of the same species and modern feather vs. modern blood samples, and one-way ANOVA to assess the pairwise differences in endogenous content between sample types (i.e. blood, tissue, toe pad, and feather).

### Population genomic analysis

The estimated individual nuclear-genome sequencing depths in the final data set ranged from 0.7 to 12× in *A. ararauna*, 0.6 to 20.7× in *A. chloropterus*, and 0.5 to 8.9× in *A. severus*. Given the uneven sequencing depth, we used ANGSD (Korneliussen et al., 2014) to generate genotype likelihoods in BEAGLE format for each species with command-line arguments *-GL 2 –doGlf 2 –doMajorMinor 1 –SNP_pval 1e-6 –doGeno –1 –doPost 1 –minMapQ 30 –minQ 20 –minMaf 0.05 –doCounts 1 –doMaf 3 –geno_minDepth 2 –setMinDepthInd 2 –postCutoff 0.95 –remove_bads 1 –uniqueOnly 1 –baq 1 –C 50*, additionally we set *-minInd* to only retain sites covered in at least 50% of the individuals (38 in *A. ararauna*, 28 in *A. chloropterus*, and 19 in *A. severus*). We then performed linkage disequilibrium (LD) pruning using PLINK v1.9 (Chang et al., 2015) (window size: 50 SNPs; step size: 5 SNPs; *r²* threshold: 0.3), allowing unrecognized chromosome codes (*-allow-extra-chr*). Finally, we removed the Z chromosome (scaffold CM054410.1), retaining 350,541 SNPs in *A. ararauna*, 169,796 SNPs in *A. chloropterus*, and 507,423 SNPs in *A. severus*. To evaluate the presence of any additional sex-linked scaffolds, we performed coverage comparison analysis through the sex assignment through coverage (SATC) framework (Nursyifa et al., 2022) implemented in R (https://github.com/popgenDK/SATC). However, no additional sex-linked scaffolds were identified. In addition, using individuals with known sex as positive controls, we used SATC to determine the sex of the sampled individuals otherwise lacking this information to evaluate if our data sets were sex biased.

We conducted principal component analysis (PCA) to explore population structure based on genotype likelihoods. Covariance matrices were generated using PCAngsd v1.21 (Meisner & Albrechtsen, 2018). These matrices were then imported into R v4.4.3 (R Core Team, 2023), where we compute principal components using the *prcomp* function from base R. We calculated the proportion of variance explained by each principal component and visualized the results using ggplot2 (Wickham, 2016). To further investigate genetic structure within each species, we used NGSadmix (Nursyifa et al., 2022; Skotte et al., 2013) to estimate individual ancestry proportions. We tested values of *K* (the number of ancestral populations) from 1 to 15, performing 50 independent runs for each *K*. We used an in-house bash script to extract likelihood values from each run and identify the best replicate (the run with the highest likelihood) for each *K* value. The best runs (.qopt files) for each *K* value and its likelihood values were loaded into R v4.4.3 (R Core Team, 2023) for downstream analysis and visualization using *ggplot2 (Wickham, 2016)*. We calculated Δ*K* following the method of Evanno et al. (2005) to objectively determine the optimal *K* value for each species.

### Mitogenome assembly and phylogeny

Adapter sequences and low-quality bases were trimmed from the raw fastq data using AdapterRemoval v2.3.1 (Schubert et al., 2016). Mitochondrial genomes were *de-novo* assembled using NOVOPlasty v4.3 (Dierckxsens et al., 2017). One mitochondrial genome for each species was annotated using the MITOS2 tool within the Galaxy platform (https://usegalaxy.eu/). Annotations were refined by comparison with previously annotated *Ara* mitogenomes deposited in NCBI (KF946546.1, NC_029319.1, and NC_047199.1). We used the Transfer Annotations tool implemented in Geneious Prime to annotate the remaining mitogenomes by species. We recovered 80 complete mitochondrial genomes for *A. ararauna*, 73 complete mitogenomes for *A. chloropterus*, and 45 complete mitogenomes and six partial mitogenomes for *A. severus*. The sequences were deposited in GenBank (NCBI) under the accession codes PV918056-PV918259. In addition, we obtained *A. macao* (GenBank accession NC_045076.1) and *A. glaucogularis* (GenBank accession JQ782215) mitogenome sequences to serve as outgroups for *A. chloropterus* and *A. ararauna*, respectively. We also used one of our *A. ararauna* sequences (LGEMA 4437) as an outgroup for *A. severus.* Finally, we included the mitogenome sequences of captive specimens of *A. chloropterus* (GenBank accession NC_047199) and *A. severus* (GenBank accession KF946546).

Mitogenome sequences were aligned using MUSCLE v3.8 (Edgar, 2004) with default parameters. We created a partitioned nexus file with separate partitions for genes, codons in protein coding genes (PCGs), and the mitochondrial control region. A maximum-likelihood phylogeny was inferred using IQ-TREE v2.0.3 (Minh et al., 2020) with 1000 rapid bootstraps and using the option *-m MFP+MERGE*, which performs nucleotide substitution model selection and chooses the best-fit partitioning scheme under the Bayesian information criterion (BIC) via an implementation of PartitionFinder (Lanfear et al., 2012) (See ST 10-12). Genetic distances between highly supported haplogroups were calculated using the *Species Delimitation* tool (Masters et al., 2011) implemented in Geneious Prime.

### *F_ST_* and genetic diversity, and inbreeding coefficients

Within each species, we estimated genome-wide genetic differentiation (*F_ST_*) between population pairs. First, we used ANGSD v0.940 (Korneliussen et al., 2014) without specifying an ancestral state to compute per-site allele frequency (SAF) likelihoods (*-dosaF 1, –GL 2*) for each population, excluding sexual chromosomes. We applied quality filters (*-minMapQ 30, –minQ 20*) and removed poorly aligned or duplicated reads (*-remove_bads 1, –uniqueOnly 1, –baq 1, –C 50*) to account for low-coverage data. Then, we estimated a folded 2D site frequency spectrum (2D-SFS) for each population pair, followed by the generation of per-site *F_ST_* indices and the calculation of global *F_ST_* values. Additionally, we inferred a folded one-dimensional SFS for each population using realSFS. This was followed by estimation of population genetic summary statistics using realSFS saf2theta and thetaStat do_stat. Per-site Watterson’s theta (θW) and nucleotide diversity (π) was calculated by summing these statistics estimates across autosomal scaffolds and dividing by the total number of callable autosomal sites. Tajima’s D and Fay & Wu’s H were summarized as site-weighted scaffold-level estimates, with non-finite values excluded. The latter two describe deviations in the site frequency spectrum that may reflect demographic change and/or selection. To estimate individual inbreeding coefficients, GLs were first computed for each population within *Ara ararauna* and *Ara chloropterus* and for *Ara severus* as a whole, set using ANGSD with the csame filtering parameters as in population genomic analysis section, but specifying –doGlf 3, and filtering out sex-linked chromosomes. Inbreeding coefficients were then estimated for each individual using ngsF (Vieira et al., 2013).

### Heterozygosity

We estimated the genome-wide heterozygosity of each individual genome within each species based on the number of polymorphic and monomorphic sites. We first used ANGSD v0.940 (Korneliussen et al., 2014) to estimate per-site allele frequency (SAF) likelihoods (*-dosaF 1, –GL 2*) from BAM files, excluding sexual chromosomes. We applied quality filters (*-minMapQ 30, –minQ 20*) and removed poorly aligned or duplicated reads (*-remove_bads 1, –uniqueOnly 1, –baq 1, –C 50*) to account for low-coverage data. Then, we used realSFS, a tool within ANGSD v0.940 (Korneliussen et al., 2014), to obtain the folded site frequency spectrum (SFS). Autosomal heterozygosity was then estimated for each individual as the number of polymorphic sites divided by the total number of callable sites. To test whether heterozygosity values differed across biomes within *A. ararauna* and *A. chloropterus*, we used the Kruskal-Wallis rank-sum test (Kruskal & Wallis, 1952). Additionally, we performed pairwise Wilcoxon tests (Wilcoxon, 1945) for specific biome comparisons. These tests and visualizations of the heterozygosity results were performed in R v.4.5.0 (2025-04-11) (R Core Team, 2023), using ggplot2 (Wickham, 2016) and ggpubr (Kassambara, 2025) packages.

### Demographic analysis

We used *Stairway Plot* v2.1.1 (X. Liu & Fu, 2020) to model effective population size (*N_e_*) dynamics based on folded SFS for each species (all individuals, except the captive specimens, see ST1-3) following the package’s instructions (https://github.com/xiaoming-liu/stairway-plot-v2). In addition, we modeled the *N_e_* dynamics of the three main populations within *A. ararauna* and *A. chloropterus*: West Amazon, East Amazon, and Cerrado. Plotting the results requires scaling the data to real-time using an appropriate substitution rate (sites/year/generation) and generation time. However, knowledge about the generation time is very limited in these three species. Therefore, we estimated the generation time (*g*) by multiplying the age of sexual maturity by a factor of two, as recommended by Nadachowska-Brzyska et al. (2015). For this, we assumed a maturity age of four years for all the species, consistent with findings from other macaws (Bueno, 2000; Collar et al., 2020d; Voss, 2005). The substitution rate (*u*) was set to 2.21 × 10^-9^ substitutions/site/year (Nam et al., 2010), multiplied by the estimated generation time. Therefore, we used a mutation rate of 1.768 × 10^-8^ mutations/site/generation. Plotting files were rescaled and visualized in R v.4.5.0 (2025-04-11).

### Genome annotation

Genome assemblies of *Ara ararauna* (GCA_028858755.1), *Ara chloropterus* (GCA_010014725.2) and *Ara severus* (GCA_024733695.1) were obtained from the NCBI public repository, and BUSCO (Tegenfeldt et al., 2025) and gfastats (Formenti et al., 2022) were applied to assess the assemblies’ quality based on conserved genes and their contiguity. The assemblies were annotated using a pre-release version of the EBP-Nor genome annotation pipeline (https://github.com/ebp-nor/GenomeAnnotation). Through this pipeline, first AGAT (https://zenodo.org/record/7255559) agat_sp_keep_longest_isoform.pl and agat_sp_extract_sequences.pl were applied to the GRCg7b (GCA_016699485.1) chicken genome assembly and annotation to generate one protein sequence (the longest isoform) per gene. Miniprot (Li, 2023) was used to align the protein sequences to the curated assemblies. UniProtKB/Swiss-Prot (UniProt Consortium, 2023) release 2023_02 and the Vertebrata portion of OrthoDB v11 (Kuznetsov et al., 2023) were also aligned separately to the assemblies. RED (Girgis, 2015) was applied via redmask (https://github.com/nextgenusfs/redmask) to the assemblies to mask repetitive regions. In addition to gene models from protein alignments, GALBA (Brůna et al., 2023; Hoff & Stanke, 2019; Stanke et al., 2006) was run with the chicken proteins using the miniprot mode on the masked assemblies to generate *ab initio* predicted gene models. The funannotate-runEVM.py script from Funannotate was used to run EvidenceModeler (Haas et al., 2008) on the alignments of chicken proteins to the parrot genome assemblies, UniProtKB/Swiss-Prot proteins, Vertebrata proteins from OrthoDB, and the predicted genes from GALBA. The resulting predicted proteins were compared to the protein repeats that Funannotate distributes, using DIAMOND blastp, and the predicted genes were filtered based on this comparison using AGAT. The filtered proteins were compared to the UniProtKB/Swiss-Prot release 2023_02 using DIAMOND (Buchfink et al., 2015) blastp to find gene names, and InterProScan (Jones et al., 2014) was used to discover functional domains. The agat_sp_manage_functional_annotation.pl tool of AGAT was used to attach the gene names and functional annotations to the predicted genes. BUSCO (Tegenfeldt et al., 2025) analysis was conducted on the predicted proteins. A full list of relevant software tools and versions is presented in Table S16.

### Ancestral state estimation

We generated the ancestral state for *A. ararauna* and *A. chloropterus* using shotgun sequencing reads of three closely related taxa each based on the genome-wide phylogeny (Smith et al., 2023). For *A. ararauna* we selected an *A. glaucogularis* (ENA: SRR8064809), *A. ambiguus* (ENA: SRR12045419), and *A. chloropterus* individual LGEMA11461; and for *A. chloropterus* we selected an *A. glaucogularis*, *A. ambiguus*, and *A. ararauna* individual LGEMA15314. The raw sequencing data for these individuals was mapped against the *A. ararauna* reference genome assembly bAraAra1.hap1 (GCA_028858755.1, unpublished) using the same methods as described above in the ‘Genomic data processing’ section. To avoid bias in generating the ancestral state, we downsampled the bam files using SAMtools to obtain an equal sequencing depth of 9.55x for all taxa. We then generated the ancestral state FASTA files using angsd v0.931 –doFasta 2 with a minimum base quality of 20, minimum mapping quality of 25, and the options –remove_bads 1, and –uniqueOnly 1.

### Selection scanning

To identify genomic regions potentially associated with adaptation to the Cerrado biome, we performed genome-wide selection scans in *Ara ararauna* and *A. chloropterus*. Because pairwise population structure analyses revealed substantial differentiation within Amazonia, we did not pool Amazonian individuals into a single comparison group. Instead, for both species, we contrasted Cerrado populations independently against each Amazonian subpopulation (Cerrado vs. West Amazon and Cerrado vs. East Amazon).

For each species and population, we estimated site allele frequency likelihoods (SAFs) using ANGSD v0.940 with the following settings: –GL 1 –doSaf 1 –baq 1 –C 50 –minMapQ 30 –minQ 20 –setMinDepthInd 2 –remove_bads 1 –uniqueOnly 1. The –minInd threshold was set to 50% of the sample size for each population. The ancestral state was specified using the species-specific ancestral reference described above. Pairwise unfolded two-dimensional site frequency spectra (2D-SFS) were then estimated for each population comparison using realSFS. These 2D-SFS were used as priors to estimate *F_ST_* with realSFS fst index, and windowed *F_ST_* values were extracted using realSFS fst stats2 in 50-kbp sliding windows with 10-kbp steps.

We estimated site-frequency-spectrum-based diversity and neutrality statistics from the SAF files. Per-site theta likelihoods were first computed using realSFS saf2theta, and subsequently summarized into window-based estimates of nucleotide diversity and neutrality statistics (e.g., Tajima’s *D* and Fay and Wu’s *H*) using thetaStat do_stat, applying the same window size and step size as for the *F_ST_*scans. We focused particularly on Fay and Wu’s *H*, as strongly negative values can indicate an excess of high-frequency derived variants consistent with recent positive selection.

Outputs were imported into R for filtering, outlier detection, and gene annotation. Windows with fewer than 100 informative sites were excluded, and the Z chromosome was excluded from downstream analyses. *F_ST_*outlier windows were defined separately for each species and population comparison as windows in the upper 1% of the empirical *F_ST_* distribution. Low-*H* windows were defined separately for each species and biome as windows in the lower 1% of the empirical Fay and Wu’s *H* distribution. Candidate genes were identified by intersecting outlier windows with gene annotations from the *A. ararauna* reference genome. We generated separate gene lists for: (i) genes overlapping *F_ST_* outlier windows, (ii) genes overlapping low-*H* windows, and (iii) genes with both signals, meaning overlapping *F_ST_* outlier windows per comparison and low Fay and Wu’s *H* within the same a species and biome.

To evaluate whether genomic regions with elevated population differentiation also showed significantly lower Fay and Wu’s H values. We tested whether Fay and Wu’s H was lower in *F_ST_* outlier windows than in non-outlier windows for each species and pairwise comparison using a one-sided Wilcoxon rank-sum test. P-values were adjusted for multiple testing using the Benjamini–Hochberg correction (Benjamini & Hochberg, 1995).

### GO enrichment

Gene ontology (GO) enrichment analyses were performed using the clusterProfiler package (Yu et al., 2012), restricting the analysis to biological process (BP) terms. GO annotations were parsed directly from the *Ara ararauna* GFF file. Enrichment was assessed using scan-aware background sets to account for gene detectability biases. Thus, for *F_ST_*analyses the background consisted of all genes overlapping windows that were testable in the corresponding comparison scan after filtering. For the combined *H*-*F_ST_* analyses, the background consisted of genes testable in both. Enrichment significance was evaluated using a hypergeometric overrepresentation test with Benjamini–Hochberg correction (Benjamini & Hochberg, 1995). A summary of the number of genes included in each analysis can be found in Table S19.

## Results

### Genome annotations

The downloaded genome assemblies of *Ara ararauna*, *Ara chloropterus* and *Ara severus* have vastly different contiguity. *Ara chloropterus* had the most fragmented assembly, with nearly 460,000 contigs and a contig N_50_ of 4.2 kb, followed by *A. severus* with approximately 183,000 contigs and a contig N_50_ of 16.9 kb. In contrast, *A. ararauna* had a highly contiguous assembly, with 353 contigs and a contig N_50_ of 38.8 Mb (Table S17). These differences were also reflected in the annotation statistics where the most fragmented assembly, *A. chloropterus*, exhibited only 39.3 % complete BUSCO genes, whereas the highly contiguous assembly, *A. ararauna*, exhibited the higher proportion of complete BUSCO genes (Table S17). Although *A. chloropterus* and *A. severus* had more annotated genes, these gene models were generally shorter and more fragmented than those in *A. ararauna* (Table S17). To minimize biases associated with comparing analyses across assemblies of uneven quality, we therefore mapped all sequencing data to the *A. ararauna* reference genome and used this genome as the common coordinate system for downstream population genomic analyses.

### Population structure

This final dataset for downstream nuclear DNA analysis included 65 *A. ararauna*, 57 *A. chloropterus*, and 38 *A. severus* samples (Table S1). The endogenous content in the remaining samples was too low to attempt further sequencing to reach a desirable depth. However, several of these were used for mitochondrial genome assembly (Tables S1-3). Significant sex bias in sampling was tested using the sex identifications from our SATC analysis for *A. ararauna* and *A. chloropterus* (Figures S5–7). The observed male-to-female ratios were 1.03 in *A. ararauna*, 1.29 in *A. chloropterus*, and 1.70 in *A. severus* (Table S1-3). The principal component analysis (PCA) based on genotype likelihoods revealed clear genetic structure within the three *Ara* species: *A. ararauna*, *A. chloropterus*, and *A. severus* (Figure 1).

**Figure 1.**
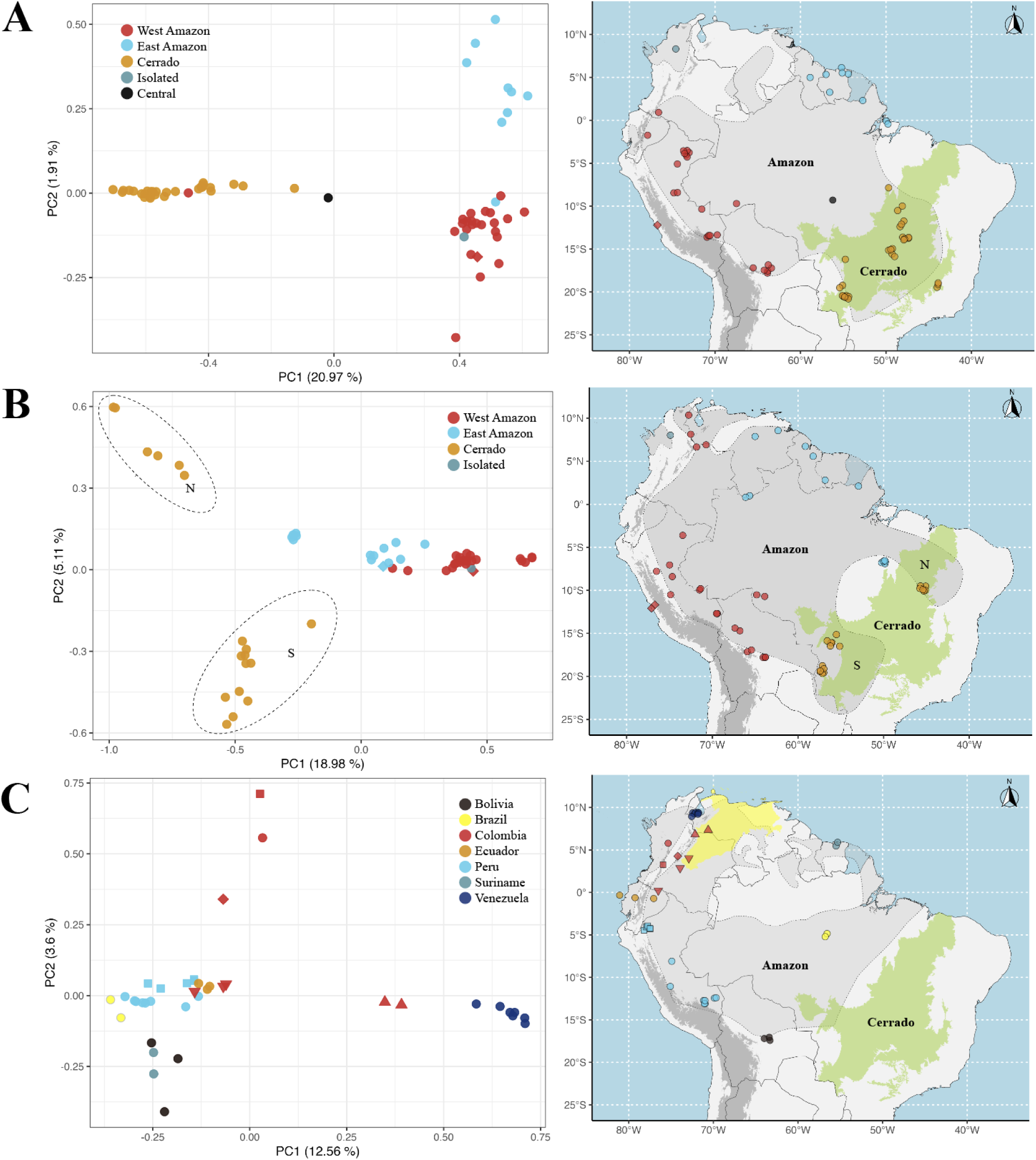
PCA and geographic location of the samples. Panels correspond to (A) *Ara ararauna*, (B) *Ara chloropterus*, and (C) *Ara severus*. In *A. severus*, points are color-coded by country, and distinctive shapes are used to differentiate localities of Peru (Square = Amazonas, Circles = All other localities) and Colombia (Squares = Cauca, Diamond = Cundimarca, Up-triangle = Arauca and Boyaca, and down-triangle = Meta and Putumayo). In *A. ararauna* and *A. chloropterus*, points are color-coded to correspond to the Cerrado and Amazon (West and East) biomes; captive specimens are represented by diamond-shaped symbols, including the single Norwegian captive individual of *A. chloropterus* (light-blue diamond); the isolated populations are shown with a teal dot, and the central population in *A. ararauna* is shown with a black dot. Background layers include the South American continent (light gray), the Cerrado biome (green), and the IUCN species distribution range (gray polygon). Locations were jittered slightly to minimize overlap. Elevations over 3000 meters a.s.l. are highlighted in darker grey and roughly correspond to the Andes mountain ranges.

#### Ara ararauna

In *Ara ararauna*, individuals clustered into three major groups corresponding to the West and East regions of the Amazon biome, and the Cerrado biome, with only a few individuals grouping with individuals from different biomes (Figure 1A). In *Ara ararauna*, individuals from the Cerrado biome formed a distinct cluster in the PCA space, clearly separated from Amazonian individuals along PC1, which explains 21% of the genomic variation, indicating strong genetic differentiation. In addition, individuals from Cerrado showed greater dispersion along PC1, suggesting higher within-population genetic variation. In contrast, individuals from the two different Amazonian populations (West and East) showed some separation between each other along PC2, which accounts for only 1.9% of the variation, indicating much lower levels of population structure. Moreover, individuals within both populations exhibited lower dispersion along PC1, consistent with weaker differentiation relative to Cerrado individuals. In addition, global *F_ST_* among these three populations was in general low or moderate (Table 1). *F_ST_* was highest between Cerrado and East Amazon (*F_ST_* = 0.07), and was the lowest between the two Amazonian populations (*F_ST_* = 0.047). This is consistent with our admixture analysis (Figure 2). Based on the Δ*K* method, the best-supported admixture result was *K* = 2 (Figure S22), which separated individuals from the Amazon and Cerrado biomes, consistent with the PCA results (Figure 2A). However, higher *K* values revealed additional population structure that was suggested by our PCA analysis (Figure 1A). For example, at *K* = 4 and higher, the admixture analysis discriminated the two Amazonian populations as in our PCA analysis. Moreover, it uncovered genetic structure within the Cerrado that was consistent with the differentiation we found between Cerrado individuals from the North and South (Figure S11).

**Figure 2.**
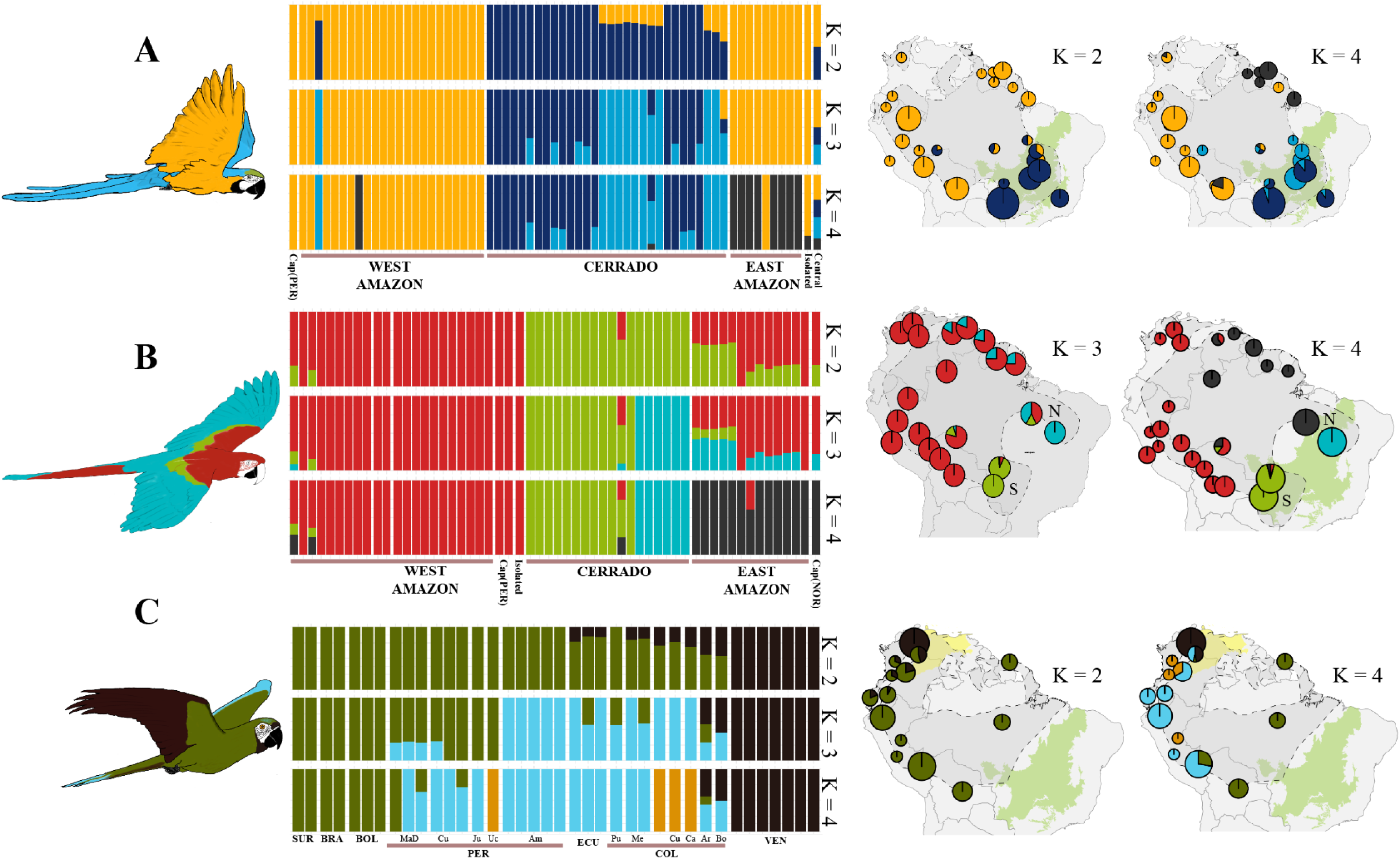
Admixture analysis of. (A) *Ara ararauna,* (B) *Ara chloropterus,* and © *Ara severus*. Bar plots for *Ara ararauna* and *Ara chloropterus* are grouped according to the biome of origin (West Amazon, East Amazon, and Cerrado), while in *Ara severus,* individuals are grouped by country of origin. The bar plots also differentiate captive individuals and the isolated populations (if any) and samples that don’t belong to either biome (Central). Spatial admixture results are shown for each species, with samples clustered geographically within 200 km. For each species, the best-supported *K* (based on the Δ*K* method) value (left) and *K* = 4 (right) are plotted. The Cerrado biome is highlighted in light green in all maps, while the Dry Northern South America biome is additionally shown in yellow in the *A. severus* maps. Dashed outlines represent species distribution range

**Table 1.** Global weighted pairwise *F_ST_* values estimated among populations in *A. chloropterus* and *A. ararauna*.

| Species | Population 1 | Population 2 | $F_{ST}$ | Interpretation |
| --- | --- | --- | --- | --- |
| *A. chloropterus* | Amazon | Cerrado | 0.0486 | Low |
|  | West Amazon | East Amazon | 0.044 | Low |
|  | Cerrado North | Cerrado South | 0.103 | Moderate |
|  | Cerrado North | East Amazon | 0.060 | Moderate |
|  | Cerrado South | West Amazon | 0.077 | Moderate |
| *A. ararauna* | Amazon | Cerrado | 0.047 | Low |
|  | West Amazon | East Amazon | 0.047 | Low |
|  | West Amazon | Cerrado | 0.051 | Moderate |
|  | East Amazon | Cerrado | 0.070 | Moderate |

Patterns of genome-wide heterozygosity and inbreeding partially mirrored the population structure inferred from PCA and admixture analyses, whereas nucleotide diversity remained broadly comparable among populations (Table 2). In *A. ararauna*, nucleotide diversity (π) was comparable across populations, but individuals from Cerrado exhibited lower mean genome-wide heterozygosity (*H_G_* = 8e-4) and higher inbreeding coefficients (*F* = 0.015) relative to Amazonian populations (West Amazon: *H_G_* = 1.2e-3 and *F* = 0.006; East Amazon: *H_G_* = 1.2e-3 and *F* = 0.011). In contrast, the two Amazonian populations showed similar heterozygosity and inbreeding values, in agreement with their weak differentiation in PCA and relatively lower pairwise differentiation based on *F_ST_* values. Tajima’s *D* and Fay & Wu’s *H* were positive across all populations, with no strong deviations among biomes. (Table 2, Figures S40-S41)

**Table 2.** Population-level summary of global autosomal *θ_W_* and neutrality statistics. For each population, sample size (*n*), and total sites (sum of callable sites) are shown. Per-site Watterson’s theta (θ_W_) and nucleotide diversity (π) are reported, calculated as the summed theta estimate divided by total sites. Neutrality statistics (Tajima’s *D*, and Fay & Wu’s *H*) were summarized as site-weighted averages of scaffold-level estimates, excluding non-finite values. We also show mean individual inbreeding coefficients (*F*), mean heterozygosity (*H_G_*), and their respective standard deviations (SD).

| Species | Population | $n$ | Total sites | $\theta_w$ | $\pi$ | Weighted Tajima's $D$ | Weighted Fay & Wu's $H$ | inbreeding coefficient ($F$) ± SD | Genome- wide heterozygosity ($H_G$) ± SD |
| --- | --- | --- | --- | --- | --- | --- | --- | --- | --- |
| *Ara ararauna* | Amazon West | 23 | 4.738e+08 | 9.029e-04 | 1.031e-03 | 4.901e-01 | 4.483e-01 | 6.550e-03 ± 2.2e-02 | 1.22e-03 ± 3.8e-04 |
| *Ara ararauna* | Amazon East | 9 | 8.688e+08 | 8.840e-04 | 1.067e-03 | 8.537e-01 | 5.414e-01 | 1.052e-02 ± 2.2e-02 | 1.24e-03 ± 4.8e-04 |
| *Ara ararauna* | Cerrado | 30 | 1.000e+09 | 8.792e-04 | 9.749e-04 | 3.633e-01 | 4.398e-01 | 1.523e-02 ± 1.1e-02 | 7.86e-04 ± 8.9e-05 |
| *Ara chloropterus* | Amazon West | 24 | 6.147e+08 | 4.812e-04 | 5.693e-04 | 6.687e-01 | 3.901e-01 | 6.360e-03 ± 1.8e-02 | 9.13e-04 ± 3.9e-04 |
| *Ara chloropterus* | Amazon East | 13 | 9.241e+08 | 7.940e-04 | 1.143e-03 | 1.765e+00 | 2.971e-01 | 2.755e-02 ± 7.5e-02 | 7.97e-04 ± 4.3e-04 |
| *Ara chloropterus* | Cerrado | 18 | 9.596e+08 | 5.083e-04 | 5.796e-04 | 5.274e-01 | 4.408e-01 | 3.257e-02 ± 3.4e-02 | 4.11e-04 ± 5.3e-05 |
| *Ara chloropterus* | Cerrado North | 6 | 9.16e+08 | 1.17e-03 | 1.76e-03 | 2.35e+00 | 2.21e-01 | 5.012e-02 ± 5.3e-02 | 4.11e-04 ± 5.3e-05 |
| *Ara chloropterus* | Cerrado South | 12 | 9.59e+08 | 5.23e-04 | 6.83e-04 | 1.24e+00 | 4.38e-01 | 2.280e-02 ± 1.6e-02 | 4.11e-04 ± 5.5e-05 |
| *Ara severus* | All | 38 | 9.514e+06 | 1.795e-03 | 1.534e-03 | -5.387e-01 | 3.631e-01 | 6.250e-03 ± 1.1e-02 | 1.71e-03 ± 3.5e-04 |

#### Ara chloropterus

In *A. chloropterus*, the PCA revealed high genetic differentiation between individuals from the Cerrado and Amazon biomes, primarily along PC1, which explains 19% of the genomic variation, although some overlap between groups was observed (Figure 1B). Within the Amazon biome, individuals from the West and East populations showed moderate separation along PC1 suggesting lower but significant genomic differentiation. In contrast, two clearly distinct clusters were found within the Cerrado biome, corresponding to the two disjunct distribution areas in the north and in the south (IUCN, 2025). These clusters were mainly separated along PC2, which explains 5.11% of the variation, indicating population structure within the Cerrado itself (see Table 1). Based on the Δ*K* method (Evanno et al., 2005), the best-supported number of ancestral groups was *K* = 3 (Figure S25), which separated individuals from the Amazon and Cerrado biomes and differentiated individuals from the North and South Cerrado, consistent with the PCA results (Figure 2B). Under this scenario, the East Amazon population is genetically similar to the West Amazon population but with some admixture with the North population of Cerrado. F_ST_ results showed more differentiation between Cerrado and Amazon (*F_ST_*= 0.049) than between the two Amazonian populations (*F_ST_* = 0.044). However, those values were low (*F_ST_* < 0.05, Table 1). Moderate *F_ST_*values (*F_ST_* > 0.5) were found between the two Cerrado subpopulations, and between those subpopulations and the closer Amazonian subpopulation (Table 1).

In *A. chloropterus*, patterns of genetic diversity further supported the population structure identified in PCA and admixture analyses (Table 2). Individuals from Cerrado exhibited substantially lower mean heterozygosity (*H_G_* = 4e-3, Figure 3) and higher inbreeding coefficients (*F* = 0.033, Table S13) compared to Amazonian populations (*H_G_*= 9e-4 and *F* = 0.006 in West Amazon, and *H_G_* = 8e-4 and *F* = 0.028 in East Amazon), consistent with the differentiation observed along PC1. Within the Cerrado, the northern subpopulation showed elevated nucleotide diversity (π = 1.8e−3), whereas the southern subpopulation exhibited lower diversity (π = 6.8e−4). Both Cerrado populations exhibited Tajima’s D values >1 (Table 2), suggesting structuring and potential differences in demographic history between these regions. Amazonian populations displayed moderate heterozygosity (Figure 3), and similar to Cerrado, these populations exhibited clearly positive Tajima’s *D* values (Table 2), particularly in the eastern population.

**Figure 3.**
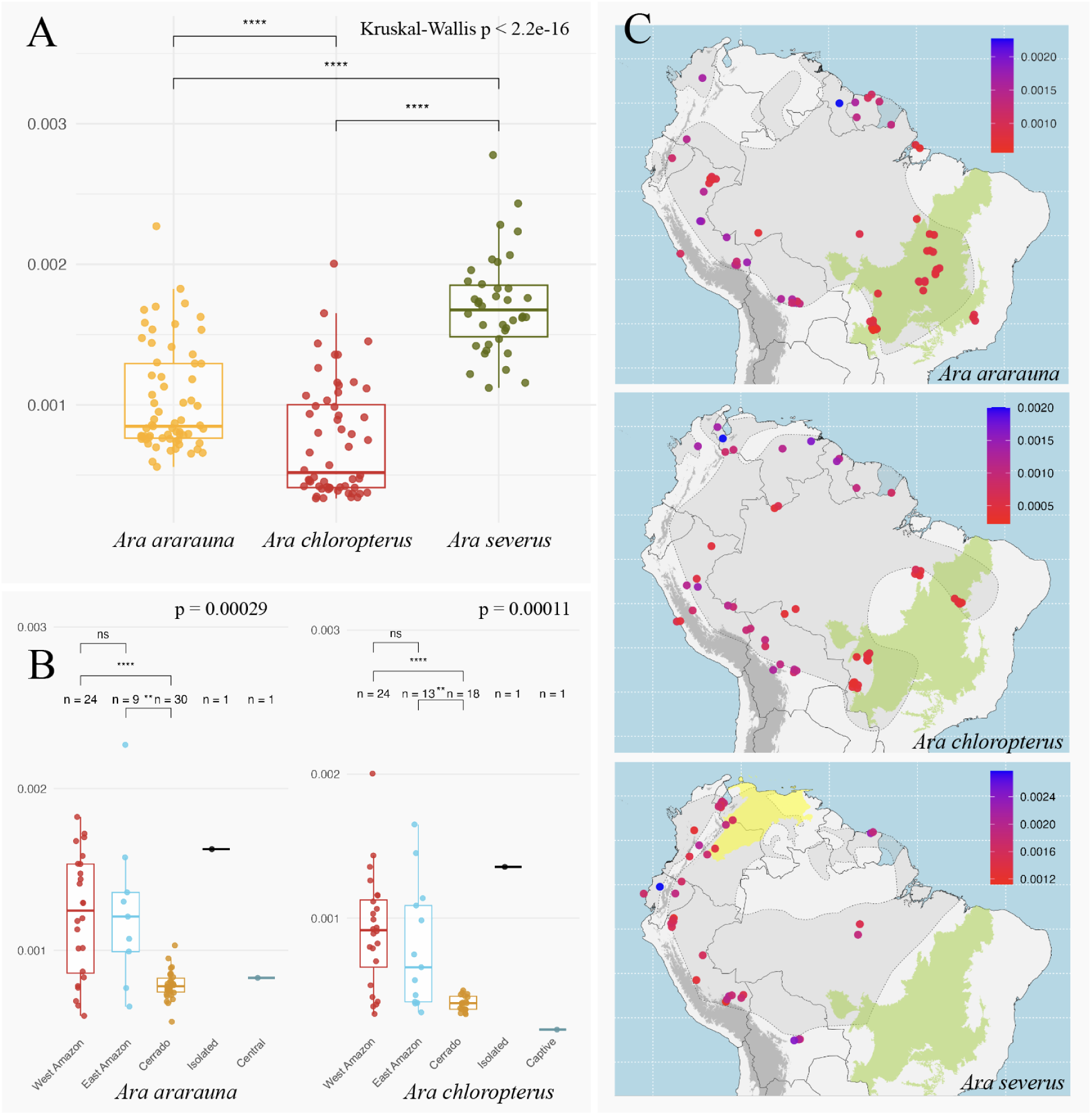
Genome-wide heterozygosity in *A. ararauna*, *A. chloropterus*, and *A. severus*. (A) Individuals’ heterozygosity values across the three species are shown as box plots. (B) Calculated heterozygosity compared across biomes for *A. ararauna* and *A. chloropterus*. We also show the sample found in the disconnected range of their respective species. Additionally, we present specimens that do not fit into the other categories. (C) Spatial distribution of heterozygosity values across their species’ ranges (shaded and delimited with dotted lines).

For both *A. ararauna* and *A. chloropterus*, we included one individual from an isolated population inhabiting the northwestern part of the Amazon isolated by the Andes mountain ranges (Figure 1A and 1B). Both PCA (Figure 1; grey dot) and admixture (Figure 2) analysis placed those samples within the West Amazon population for both species. In addition, we also included captive specimens in the analysis. Samples from captive individuals from Lima (Peru) clustered with the West Amazon population along with the rest of Peruvian samples for both *A. ararauna* and *A. chloropterus*. However, the captive Norwegian individual of *A. chloropterus* grouped with the East Amazon population. Finally, we note that the undersampled “central” Amazon population (n = 1; LGEMA 12339), represented only in *Ara ararauna*, showed a complex admixture pattern involving all distinct genetic clusters in the admixture results at any *K*. Moreover, it is in the middle of all other samples in the PCA plot (Figure 1; black dot).

#### Ara severus

In *A. severus*, individuals from Venezuela (n = 7) and two specimens from nearby localities in Colombia formed a distinct genetic cluster along PC1, which accounted for 12.6% of the genomic variation (Figure 1C), clearly separating them from the rest of the individuals sampled. Within this cluster, subtle differentiation between Venezuelan and Colombian individuals was also observed along PC1. Among the remaining individuals, minor structure was observed along PC2 (3.6% of variation), generally reflecting geographic proximity. Admixture analysis supported these patterns, with the best-supported model based on Δ*K (Evanno et al., 2005)* indicating *K* = 2 ancestral clusters: one corresponding to Venezuelan individuals, and the other to individuals from Peru, Bolivia, Brazil, and Suriname. Individuals sampled from regions between these two populations exhibited mixed ancestry, consistent with geographic intermediacy and potential gene flow.

### Heterozygosity

Genome-wide heterozygosity differed significantly among species and populations (Figure 3A–C). *Ara severus* exhibited the highest heterozygosity (*H_G_*= 1.7e-3), which was approximately 2.4 times higher than that of *A. chloropterus* (*H_G_* = 7e-4), the species with the lowest heterozygosity. Differences among species were significant (*p* < 0.05) under a Kruskal–Wallis test (Kruskal & Wallis, 1952), with all pairwise comparisons also significant under Wilcoxon tests (Wilcoxon, 1945). Within *A. ararauna* and *A. chloropterus*, heterozygosity varied significantly among populations (*p*-value: 3e-4 and 1e-4 respectively). Further pairwise tests indicated no significant differences between Amazonian populations, but individuals from Cerrado exhibited significantly reduced heterozygosity levels compared to both Amazonian populations individually. In addition, we observed that individuals from the isolated population in both species exhibited high heterozygosity (Table S7). In contrast, when heterozygosity was mapped by geographic location (Figure 3C), individuals from the Amazon Biome with low heterozygosity were consistently situated deeper within central Amazonia. In *A. ararauna*, this included the individual from the central population. Finally, the captive *A. chloropterus* individual from Norway exhibited the lowest heterozygosity across all samples with mean autosomal heterozygosity of 2e-4 (Table S7).

### Mitochondrial phylogeny

Our mitogenome data matrix contained assembled sequences from 80 *A. ararauna*, 74 *A. chloropterus*, and 52 *A. severus* individuals as well as one outgroup sample from closely related species in each data set. Within each species, pairwise identity was around 99%, but identical sites were higher in *A. chloropterus* (94.5%) and lower in *A. severus* (85.7%) (Table 3). The smallest median mitogenome length was 16,980 bp in *A. ararauna*, while the median in *A. chloropteru*s and *A. severus* was 16,994 and 16,996 bp respectively. Phylogenetic analysis of mitogenome sequences in each species revealed several major monophyletic clades with nested subclades (Figure 4). The phylogeographic patterns partially capture the broad-scale geographic patterns observed in the nuclear DNA analysis, particularly in *A. chloropterus* and *A. severus*. In addition, they reflect fine-scale genetic structuring among geographically proximate samples. In *A. ararauna* (Figure 4A) we identified a major clade (“A”) present across the whole species range, but more prevalent in the Andes population of the Amazon biome and in the Cerrado. In addition, individuals from the Cerrado carried only mitogenomes from subclade A1 (particularly A1a and A1c) and B3, whereas the Andes population was dominated by subclade A2, with the exception of the Bolivian individuals which exclusively exhibited A1b (see Figure S29). Moreover, the Andes population was also the only one to exhibit all three subclades of clade A, including A3, which was found exclusively there. Interestingly, the East Amazon population showed the greatest mitogenomic diversity, including representatives of all three main clades (A, B, C), and was the only population where haplogroup “C” was detected. The isolated population represented by individual FMNH190744 exhibited mitogenome form clade B3.

**Figure 4.**
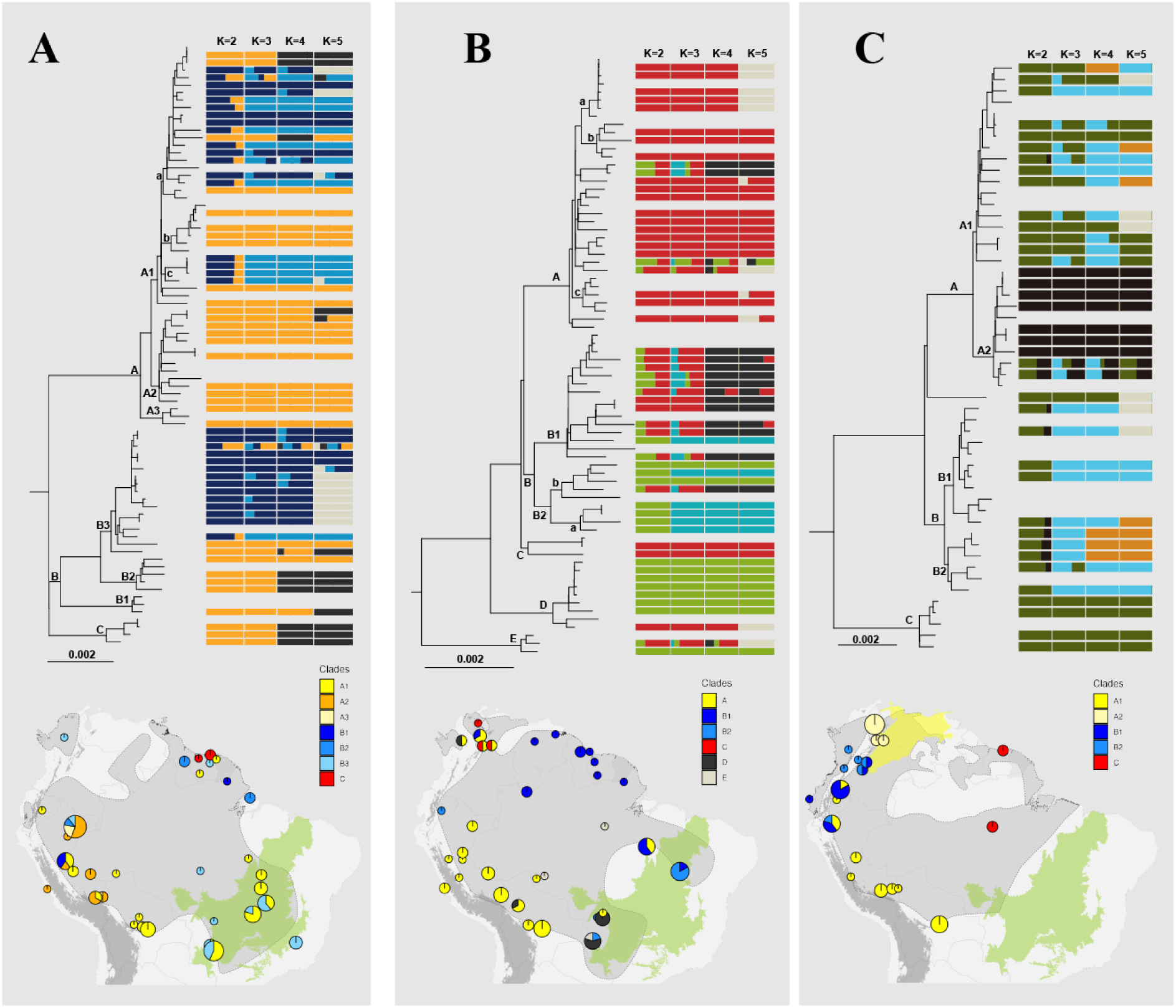
Mitochondrial haplogroups identified in. (A) *Ara ararauna*, (B) *A. chloropterus*, and (C) *A. severus*. For each species, we present a maximum-likelihood phylogeny based on complete mitogenome sequences with haplogroup labels in the nodes. Next to the mitochondrial phylogeny we show the barplots obtained using admixture. Corresponding maps display the geographic distribution of samples colored by their mitochondrial haplogroup identity and clustered within a 100-km radius; the sizes of the pie charts correspond to the number of samples represented by them. Species ranges (IUCN) are shown with blue lines. A summary table is also included in each panel to summarize average pairwise intra– and inter-genetic distances.

**Table 3.** Summary statistics of mitogenome alignments (ingroup) for the three *Ara* species.

|  | **n** | **pairwise identity** | **identical sites** | **Median** | **min length** | **max length** | **%CG** |
| --- | --- | --- | --- | --- | --- | --- | --- |
| *A. ararauna* | 80 | 99.5% | 92.6% | 16,980 | 16,979 | 16,982 | 46.5% |
| *A. chloropterus* | 74 | 99.6% | 94.5% | 16,994 | 16,991 | 16,995 | 46.7% |
| *A. severus* | 52 | 99.2% | 85.7% | 16,996 | 16,993 | 16,998 | 46.7% |

In *A. chloropterus* we identified five major mitochondrial clades (A, B, C, D, E; Figure 4B). We find mitogenome phylogeographic patterns concordant with the geographic patterns found in the nuclear analysis, particularly in the admixture results at *K* = 4 (Figure 2B). Thus, individuals from the West and East Amazon populations carried mitogenomes corresponding almost exclusively to the “A” and “B1” respectively, in agreement with the genomic structure observed in the PCA (Figure 1) and admixture analyses (Figure 2). Similarly, individuals from the North and South populations of the Cerrado biome carried predominantly mitogenomes from clades B2 and D respectively. Interestingly, we find great mitogenome diversity in the northern part of the West Amazon population (between Colombia and Venezuela), involving clades A, B1, D, and C. The latter being exclusive to this area. The isolated population represented by individual FMNH190745 exhibited a clade A1b mitogenome, in concordance with our genomic analysis, further supporting the affinity of this population with the West Amazon population.

In *A. severus* we find three main mitochondrial clades (A, B, C), which showed a spatial structure consistent with the genomic patterns observed in our analysis of the nuclear genome (Figure 1, Figure 2). In the northern part of the range (Venezuela), we found only clade “A2”, which corresponds to one of the genetically distinct groups revealed by our admixture analysis for *K* = 2 (Figure 2). Clades “A1” and “C” match the other distinct genetic cluster found in our admixture results (*K* = 2). Moreover, the mitogenome analysis can distinguish between individuals closer to the Andes (A1) and individuals that are deeper in the Amazon (Brazil and Suriname, C). This distinction corresponds roughly to the admixture results at *K* = 4. Finally, the mitogenomes of the admixed individuals found in our admixture analysis (at *K* = 2) grouped almost exclusively within clade “B”. This corresponds better to the admixture analysis at *K* =3 (Figure 4C).

### Demographic history

Stairway Plot analyses for the three species, including all populations, revealed broadly similar effective population sizes (*N_e_*) prior to the Last Glacial period (LG, ca. 116 Kya), but distinct demographic trajectories during it (Figure 5A). In *A. severus*, *N_e_*experienced a sudden expansion shortly after the Last Interglacial (LIG, ca. 116-129 Kya), peaking at over 100,000 individuals, the highest among the three species. This high *N_e_*was maintained throughout most of the LG before experiencing a steady decline to present times. In *A. ararauna*, a gradual *N_e_* increase began at the beginning of the LG but was briefly interrupted around 40 Kya, when *N_e_*returned to pre-glacial levels. A second phase of growth followed, reaching a peak near the end of the LG, followed again by a decline until recent times. *A. chloropterus*, in contrast, maintained relatively low *N_e_* throughout most of the LG until the Last Glacial Maximum (LGM, ca. 19-23 Kya), after which it underwent a rapid expansion, reaching *N_e_* sizes comparable to those of *A. ararauna* at the same point. This demographic growth peaked near the end of the LG (ca. 11.5 Kya) and persisted relatively stable until ca. 5 Kya before it started declining to its current level. Moreover, our results suggest the *N_e_* in *A. chloropterus* remained relatively similar to *A. ararauna* during that period, following a similar trajectory (Figure 5A).

**Figure 5.**
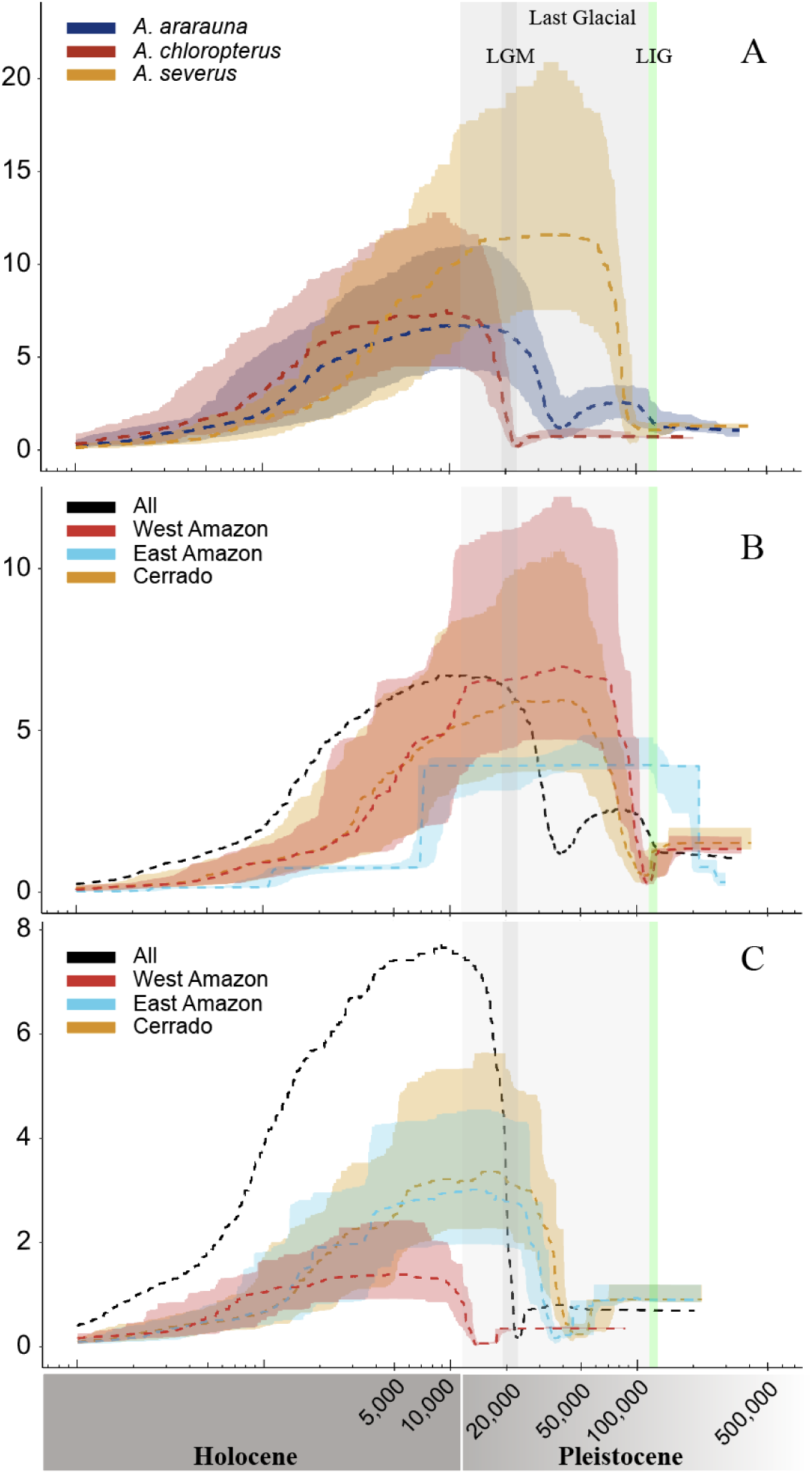
(A) Demographic histories of *A. ararauna*, *A. chloropterus*, and *A. severus*. Dotted lines represent the estimated median effective population size (*N_e_*x10^4^), and shaded areas indicate the 75% confidence intervals (CIs). Demographic histories of different populations within *A. ararauna* (B) and *A. chloropterus* (C), each shown with their respective 75% CIs; As reference, demographic history of the respective entire species is shown in black dots (with no CIs for clarity). The grey-shaded area represents the last glacial period (∼11.7-116 Kya), the darker grey band corresponds to the last glacial maximum (LGM, ∼19-23 Kya), and the green-shaded band marks the last interglacial period (LIG).

Within *A. ararauna*, Stairway Plot analyses for both the East Amazon and Cerrado populations revealed an initial increase in effective population size (*N_e_*) at the beginning of the LG period, followed by a decline, a pattern consistent with the trajectory inferred when all populations were analyzed jointly. However, this growth appeared uninterrupted, suggesting a continuous demographic expansion during the early LG (Figure 5B) in these populations. The East Amazon population began with the lowest initial *N_e_* estimate among *A. ararauna* populations. It experienced a pronounced increase in *N_e_,* reaching its maximum before the start of the LG. However, this maximum was comparatively lower than that observed in other populations and remained stable at this lower level throughout the LG and some time after it.

Within *A. chloropterus*, Stairway Plot analyses revealed distinct demographic trajectories among populations (Figure 5C). The East Amazon and Cerrado populations exhibited increases in *N_e_* prior to the Last Glacial Maximum (LGM), contrasting with the pattern observed in the combined analysis, where *N_e_* growth started around the LGM period. In contrast, the West Amazon population showed no substantial increase in *N_e_* until the late LG period. Notably, this population had the lowest maximum *N_e_* estimate among *A. chloropterus* populations.

### Selection scanning

Given the strong genetic differentiation between Cerrado and Amazonian populations in both *A. ararauna* and *A. chloropterus*, together with reduced heterozygosity and elevated inbreeding in Cerrado populations (see above), we aimed to test whether this divergence reflects local adaptation or demographic processes such as bottlenecks and reduced effective population size. Across all pairwise comparisons, we identified between 1,032 and 1,046 top-1% *F_ST_* outlier windows (Table S14, Figure S36, and Figure S37), corresponding to 168-269 annotated candidate genes (Table S19). To evaluate whether these highly differentiated regions also show signatures consistent with selection, we compared Fay & Wu’s *H* between *F_ST_* outlier and non-outlier windows (Table S14, Figure S42). In Amazonian populations of both species, *H* values were significantly lower in *F_ST_* outlier regions. In contrast, this pattern was not consistently observed in Cerrado populations.

We next identified genes located within regions showing both elevated Cerrado-Amazon pairwise *F_ST_* and reduced Fay & Wu’s *H* (Table S15). In both species, these genes were more frequently detected in Cerrado populations, with 37 in *Ara ararauna* and 33 in *Ara chloropteru*s, although we also found nine to 17 candidates in Amazonian populations (Table S15, S19). Most of the identified candidate genes were specific to particular species and populations, indicating that there is little convergence in the genomic regions potentially under selection across species. We identified only two genes (*NALCN* and *RBBP6*) within regions that were significantly differentiated between Cerrado and Amazonian populations in both *A. ararauna* and A*. chloropterus* and likely under selection in Cerrado (Figure 6). Specifically, these signals were detected in pairwise comparisons of Cerrado against Amazon East and/or Amazon West. No equivalent double-signal genes were identified for the Amazonian populations. Overall, GO enrichment provided limited evidence for functional enrichment. A few BP terms reached nominal Benjamini–Hochberg-adjusted significance for either *F_ST_* outlier genes or *F_ST_* and reduced Fay & Wu’s *H* candidate genes, but these results were weak and based on small numbers of GO-annotated genes with GO annotations.

**Figure 6.**
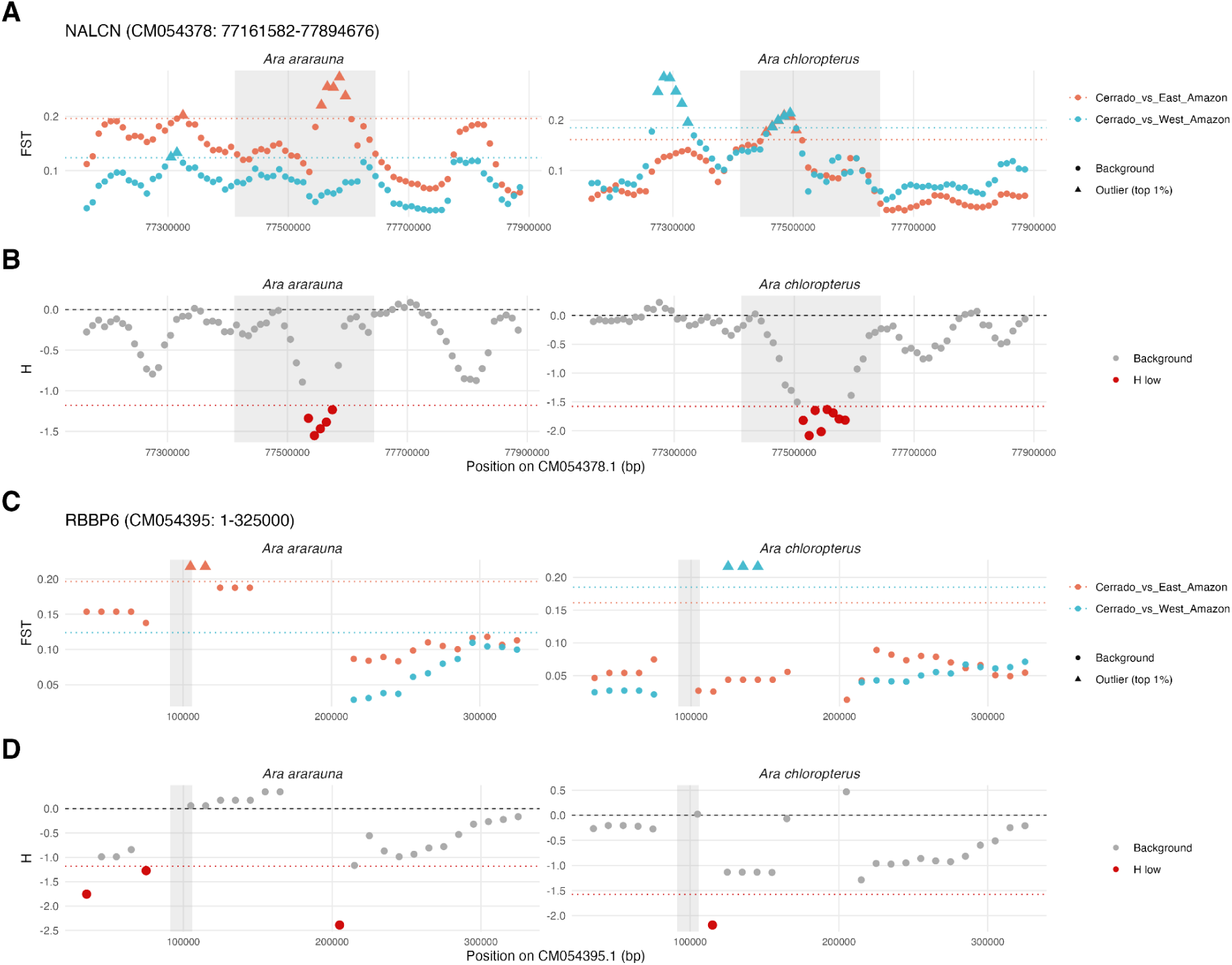
Two shared cross-species candidate loci showing concordant signatures of selection in Cerrado populations of *Ara ararauna* and *A. chloropterus*: (A-B) *NALCN* on chromosome CM054378 and (C-D) *RBBP6* on chromosome CM054395. For each locus, upper panels show windowed *F_ST_* values for comparisons between Cerrado and Amazon East (red) and between Cerrado and Amazon West (blue), with triangles indicating top 1% *F_ST_*outlier windows and dotted horizontal lines indicating the corresponding empirical *F_ST_* outlier thresholds (A and C). Lower panels show Fay & Wu’s *H* for the Cerrado population only, with red points indicating windows in the lowest 1% and the dotted horizontal line indicating the empirical H-low threshold. Shaded regions denote the annotated coordinates of the candidate gene.

## Discussion

### Population structure

The three *Ara* species examined here (*A. ararauna*, *A. chloropterus*, and *A. severus*) exhibit remarkably extensive geographic ranges across northern South America, each spanning over 8 million km² according to IUCN estimates (IUCN, 2025), which is comparable to the area of continental Europe. Such wide ranges can reflect the recognized high dispersal capacity of parrot species, especially of those with larger wing sizes (Sheard et al., 2020). Thus, our study offers the opportunity to better understand how landscape heterogeneity shapes genetic structure and connectivity in Neotropical birds, especially when no apparent barriers are present.

#### Ara ararauna

Our PCA and admixture analysis showed clear genetic structure within *A. ararauna*. Interestingly, the pattern is similar to *A. chloropterus*, with a clear differentiation between the individuals inhabiting the Amazon and Cerrado biomes. In *Ara ararauna*, at *K* = 2, which is the most likely number of ancestral clusters following Δ*K*, we can discriminate the Amazon from the Cerrado biomes (Figure 2). However, it has been described that the Δ*K* method can favor *K* = 2 and may underestimate the finer-scale population structure (Janes et al., 2017; D. J. Lawson et al., 2018). Our PCA analysis supports this, showing high variation along PC1 within this Cerrado (Figure 1), and some differentiation between West and East Amazon populations. Our mitogenome phylogeny identified distinct mitochondrial clades within Cerrado, agreeing with Caparroz et al. (2009), supporting at least two groups within Cerrado. In the Amazon biome, this analysis identified distinct clades associated with the West and East populations, suggesting phylogeographic structure (Figure 4A). These evidences suggested a preferred *K* = 4 for this species (Figure 2A). Similarly, within the same geographical area, admixture analysis using whole genome data of Hyacinth macaw (*Anodorhynchus hyacinthinus*) showed no signal of genetic structuring despite finding four clusters in their PCA analysis that were consistent with geographical isolation (Vilaça et al., 2024). This highlights the importance of integrating molecular evidence with ecological and biogeographic context when assessing population structure and differentiation in parrots.

It is worth noting that increasing the number of clusters beyond *K* = 4 (Figure S20 and S21) did not significantly improve resolution within the Amazon populations. Instead, additional clusters primarily reflected further subdivision within the Cerrado. Given that Cerrado individuals exhibit reduced heterozygosity and slightly elevated inbreeding coefficients relative to Amazonian populations, this additional clustering is consistent with stronger genetic drift, smaller effective population sizes, increased isolation, founder events, or some combination of these processes. In addition, the distributions of window-averaged values of Fay & Wu’s *H* and Tajima’s *D* broadly overlap among *A. ararauna* populations, with no marked genome-wide shift separating Cerrado from the Amazonian populations (Figures S43 and S44). This interpretation is further supported by the well-documented habitat fragmentation in this region (Vieira-Alencar et al., 2023). Moreover, neither the PCA analysis (PC1 vs PC2, and PC1 vs PC3) nor the mitogenome phylogeny supports further substructure within the Cerrado biome, though finer scale differentiation may still exist.

It is also important to note that mitochondrial DNA often reveals different patterns of population structure compared to nuclear DNA due to their different modes of inheritance (maternal and biparental respectively). Thus, mtDNA analysis can be useful to detect sex-biased dispersal and historical phylogeographic patterns in females, as documented for this species (Caparroz et al., 2009). Moreover, because mtDNA is a single, non-recombinant locus, it is more susceptible to genetic drift (Ferreira & Rodriguez, 2024). Our mitogenome analysis suggested a possible isolation of Bolivian individuals as nearly all of them exhibited mitogenomes from clade A1b, which is not found elsewhere. This pattern may represent cytonuclear discordance due to incomplete lineage sorting as seen in *A. macao* (Aardema et al., 2023) and other birds (DeRaad et al., 2023), or reflect female philopatry, which has been suggested for the species (Caparroz et al., 2009). Importantly, this differentiation is also supported by admixture results at *K* = 7 (Figure S20), pointing to an emerging signal of genetic divergence. The Bolivian population has also been shown to be genetically distinct from other populations in *A. militaris* based on the mitochondrial CYTB marker sequences (F. A. Rivera-Ortíz et al., 2023).

#### Ara chloropterus

In *A. chloropterus*, *K* = 3 was the best estimated number of ancestral clusters following Δ*K*, which agrees with the main clusters observed on our PCA analysis (Figure 1B). Under this scenario, there was a clear differentiation between individuals from the Amazon and Cerrado biomes, similar to *A. ararauna*. In contrast, it identified great differentiation between individuals from the Southern and Northern populations of Cerrado. In contrast to *A. ararauna*, these two populations are discontinuous in the Cerrado biome (Collar et al., 2020c). Although these populations are connected through the Amazon, genomic evidence does not support recent gene flow between them. Moreover, this is also reflected in the differentiation found in the mitochondrial phylogeny, low heterozygosity levels, and moderate F_ST_ values, which may indicate a history of reduced effective population size and potentially restricted gene flow between regional groups. This interpretation is consistent with the markedly lower heterozygosity and higher inbreeding coefficients observed in Cerrado populations, together with differences in nucleotide diversity and Tajima’s *D* between northern and southern subpopulations, suggesting heterogeneous demographic histories within the biome. However, it is unclear whether this geographical disconnection is recent or reflects a long term historical barrier. Future work could estimate divergence times between these two populations by applying SNP-based methods such as SNAPP (Bryant et al., 2012) and fastsimcoal2 (Excoffier et al., 2021), which would require increasing sample sizes and sequencing depths. High and moderate levels of structure within Cerrado have been observed in four passerine species based on reduced-representation genomic data, suggesting complex ancestral evolutionary histories (Lima-Rezende et al., 2022). Similar patterns of low genetic diversity and moderate population structure attributed to habitat loss have also been reported in Cerrado-endemic species, such as the Brazilian merganser (Ragusa-Netto, 2024). In other birds inhabiting Cerrado, habitat loss and population decline is also thought to be the reason for the observed low genetic diversity and moderate population structure (D. P. Campos et al., 2023).

Within the Amazon, individuals also showed some differentiation across PC1 on our PCA analysis which reflected better our admixture results at *K = 4*, therefore this is the preferred number of ancestral clusters. Under this scenario, individuals from the West Amazon were discriminated against those from the East Amazon. This structure was also observed in the mtDNA phylogeny, and even in a more pronounced manner than in *A. ararauna*, as the West Amazon population exhibited almost exclusively mitogenomes of the clade A, while the East Amazon population exhibited almost exclusively mitogenomes of the clade B1. Similar east–west genomic structuring within the Amazon has been documented in other bird species, such as the Striped Woodcreeper *Xiphorhynchus obsoletus* (Luna et al., 2022) and the Atlantic Plain-Xenops *Xenops minutus* (Harvey & Brumfield, 2015). However, in these studies, genetic differentiation has often been attributed to the isolating effects of major Amazonian rivers. Unfortunately the lack of sampling in the deep Amazon (Central population) limits our capacity of studying the role of major river basins in shaping population structure in *Ara* species. Future efforts should aim to increase sampling for this region in order to characterize the patterns of intraspecific genetic diversity within its macaw populations and identify possible dispersal barriers.

Mitogenome evidence also helped us to detect finer scale population structure. For instance, as in *A. ararauna*, Bolivian individuals exhibited almost exclusively mitogenomes of the Clade Aa, which is not found elsewhere. The combined evidence of these two species, and past works in *A. militaris* (F. A. Rivera-Ortíz et al., 2017) supports the treatment of the *Ara* populations of this region as a separate conservation unit. It is also worth mentioning that we found great mitochondrial diversity in the northern part of the species’ range, with the presence of the rare clade C. Interestingly, this region also harbors the only known divergent *A. macao* individuals in South America. Based on CYTB, Schmidt et al. (2020) found that while most individuals in this region cluster into Haplo4, individuals from the northernmost part of the range cluster into Haplo7. In contrast, *A. militaris* showed no evidence of population structure in this area (F. A. Rivera-Ortíz et al., 2017).

#### Ara severus

We did not detect significant genomic divergence between individuals from the eastern and western Amazon in *A. severus*. However, this could be because of the limited sampling size in the eastern Amazon (n = 4). Additionally, admixture analysis supported a model with only two genetic clusters. Under this scenario, individuals from the northernmost part of the range (Venezuela) and individuals from the southwest and east were entirely assigned to different ancestral populations, despite apparent geographical proximity, while individuals between them exhibited different and progressive levels of admixture along the west. This pattern resembles ring species such as *Ensatina eschscholtzii* (Pereira et al., 2011) and *Alophoixus bulbuls* (Pereira & Wake, 2015), in which gene flow occurs around a geographic barrier, but terminal populations are reproductively isolated. It is possible that the biome known as the Dry North South America acts as a dispersal barrier between peripheral populations of *A. severus*. However, further research is needed to evaluate whether reproductive isolation, particularly pre-mating barriers, has evolved between terminal populations. This scenario is possible considering that despite geographical proximity, eastern populations have been historically attributed to the nominal subspecies, while the other population (western) has been attributed to the *castaneifrons* morph, which is supposed to be slightly bigger than *A. severus severus* (Collar et al., 2020b). Pre-mating barriers due to morphological differences have been very well documented in birds (Uy et al., 2018). Further studies should examine morphological differences and its relationship with the genomic evidence The mitogenome phylogeny revealed four main mitochondrial clades, indicating greater genetic structure. Notably, individuals from the northern edge of the western population exclusively carried mitogenomes from clade A2, which was absent elsewhere, agreeing with his genomic distinction on our PCA and admixture analysis (*K* = 2). Individuals assigned entirely to the alternative genomic cluster carried mitogenomes from two other clades, which further differentiated individuals from the eastern and western portions of the Southern Amazon. Meanwhile, admixed individuals (at *K* = 2) possessed mitogenomes from clade B, which was not found elsewhere, with some exhibiting admixture with Clade A1 around the lower end of this population. This pattern more closely matched the structure inferred at *K* = 3, assigning the admixed individuals found at *K* = 2 to their own genetic cluster. However, we did not find nuclear genomic support for the east-west division within the Southern Amazon suggested by the mitochondrial phylogeny. This discrepancy may rather reflect cytonuclear discordances due to incomplete lineage sorting (Aardema et al., 2023), or female phylopatry (Caparroz et al., 2009).

### Demographic history

To investigate historical demographic events that may have shaped the current population structure in *A. ararauna, A. chloropterus,* and *A. severus*, we performed Stairway Plot analyses based on the site frequency spectrum (SFS) derived from genome-wide data. This analysis revealed broadly similar effective population sizes (*N_e_*) prior to the Last Glacial period (LG, ca. 116 Kya), but distinct demographic trajectories during this period.

In *A. severus*, *N_e_* experienced a sudden and pronounced expansion shortly after the Last Interglacial (LIG, ca. 116-129 Kya), that was sustained along this era, reaching the highest historical *N_e_*among the three species (Figure 5A). Moreover, this species has the highest genome-wide heterozygosity values across the species included here. In addition, it has been less affected by the illegal trade of *Ara* in modern times than the other species and no substantial population decline has been reported (Collar et al., 2020b). These results suggest that *A. severus* represents a historically stable and demographically robust species. Future works could explore the demographic trajectories within each of the described *A. severus* populations to determine whether the observed population structure corresponds to recent divergence or to long-term historical processes, using methods such as PSMC (https://github.com/lh3/psmc) and Stairway Plot 2 (X. Liu & Fu, 2020). The pronounced *N_e_* growth in *A. severus* could have caused extensive incomplete lineage sorting after explosive diversification or population growth (DeRaad et al., 2023), which would explain the cytonuclear discordances found in the species.

In contrast to *A. severus*, the eastern Amazon was better represented in *A. ararauna* and *A. chloropterus*, with larger numbers of sampled individuals. Furthermore, unlike *A. severus*, both species have populations occurring in the Cerrado biome. PCA and Admixture analyses (Figures 1 and 2), and mitochondrial-based phylogenies, (Figure 4) supported the genetic differentiation between individuals from the Amazon and Cerrado. Therefore we explored whether the demographic histories of these biomes followed distinct trajectories over time. When accounting for population structure, differences in *N_e_* trajectories became more pronounced within species, following different demographic histories. In *A. araruna, N_e_* expansions were observed around the beginning of the LG in the West Amazon and Cerrado populations while it started earlier, before the LIG (116-129 Kya), in the East Amazon population. However, the *N_e_* remained stable until the LGM (23 Kya) for all the populations. On the other hand, *A. chloropterus* populations began their *N_e_* expansions significantly later than those of *A. ararauna*, with the Cerrado and West Amazon populations showing expansions between approximately 50 Kya and the LGM (ca. 23 Kya). This discrepancy likely reflects species-specific responses to climatic conditions, particularly resource availability, as observed in other macaws (de la Parra-Martínez et al., 2019; Ragusa-Netto, 2024). It is also thought that climate change was more pronounced in the eastern than the western Amazon during the LGM (Dalapicolla et al., 2024). However, we did not find evidence to support differential responses between the Amazon (eastern) and Andes (western) populations in either species.

Notably, the West Amazon population of *A. chloropterus* followed a different post-LG trajectory. It remained low along the LG and started an expansion of population size after this period. This population also exhibited both the lowest estimated historical *N_e_*. Altogether, our results suggest a more recent colonization of this area (ca 12 Kya) by *A. chloropterus* than by *A. ararauna*, which exhibited *N_e_* since 100 Kya. Furthermore, it is possible that anthropogenic impacts, including habitat loss and fragmentation, have exacerbated post-glacial isolation, reinforcing genetic structure among populations (Ragusa-Netto, 2024). This could explain that despite a late colonization of the Andes area, we found considerable mitochondrial divergence among northern and southern populations, suggesting limited dispersal along the Andean foothills as in *A. militaris* (F. A. Rivera-Ortíz et al., 2023). This could be attributed to the suggested role of human settlements in landscape modification around this region during the Holocene (Schiferl et al., 2023). To this we add the acknowledged historical relationship between macaws and humans since Pre-Columbian times (Capriles et al., 2021; George et al., 2018), sourcing individuals from the wild. For instance, there are records of mummified *Ara* specimens outside their native range in archaeological context along the Pacific coast in Chile (Capriles et al., 2021), Peru (Celeste Asurza, personal communication), and Bolivia (Jose Capriles, personal communication). Moreover, an extensive record of featherwork using macaw feathers, particularly *A. chloropterus*, *A. macao* and *A. ararauna (King, 2012)*. Integrating genomic data in the study of archaeological artifacts could help us to better understand the role of ancient human populations in the observed isolation of the northern and southern individuals of the Andean population of *A. ararauna* and *A. chloropterus*.

### Selection scanning

The genomic selection scan provides limited support for a primary role of positive selection in driving the strong genome-wide differentiation observed between Cerrado and Amazonian populations. Although *F_ST_*outlier regions were identified in all pairwise comparisons, Fay & Wu’s *H* values were not consistently reduced in Cerrado outlier regions relative to non-outlier regions (Figure S42), suggesting that highly differentiated windows in Cerrado are not generally associated with an excess of high-frequency derived alleles. In contrast, Amazonian populations showed significantly lower *H* values within *F_ST_* outlier regions, indicating that some of the Amazonian populations’ differentiated regions may have been subject to localized selective sweeps or linked selection (Fay & Wu, 2000).

In Cerrado populations, the combination of elevated differentiation, reduced heterozygosity, and increased inbreeding coefficients, together with the absence of a consistent genome-wide shift in Fay & Wu’s *H*, is more consistent with demographic processes than with widespread positive selection. This is supported by the population-specific *N_e_*trajectories observed in *A. ararauna* and *A. chloropterus* (Figure 5). Thus, the genomic differentiation between Amazonian population and Cerradon may largely reflect genetic drift associated with smaller effective population size and increased isolation, rather than adaptive divergence alone. This is consistent with known history of habitat fragmentation and population decline in the Cerrado biome (Ragusa-Netto, 2024; Vieira-Alencar et al., 2023), and similar patterns have been observed in birds with known history of demographic erosion (Lu et al., 2024), and bottlenecks (Martin et al., 2021). The limited GO enrichment among candidate genes further supports this interpretation, suggesting that candidate loci are functionally heterogeneous and not concentrated within specific biological pathways.

Nevertheless, demographic processes do not exclude local adaptation. Interestingly, two genes (*NALCN* and *RBBP6*) exhibited overlapping signals of high differentiation and reduced Fay & Wu’s *H* in Cerrado populations of both *Ara ararauna* and *Ara chloropterus*, suggesting potential convergent evolutionary responses to the Cerrado biome. *NALCN* (sodium leak channel, non-selective) gene encodes a voltage-independent, nonselective cation channel that is present in zebra finch and chicken genomes (Friedrich et al., 2019). This gene has been proposed as a candidate locus associated with morphological divergence island birds, including beak morphology divergence in Galapagos finches (L. P. Lawson & Petren, 2017) and body and bill size increases in silvereyes (Sendell-Price et al., 2021). It has also been linked to adaptation to hot arid and harsh environments in chickens (Assiri et al., 2025; Gu et al., 2020), which could mirror the Amazonian-Cerrado transition. Fewer studies have directly linked *RBBP6* (retinoblastoma binding protein 6) to adaptive processes, although it has been identified in the relict gull (*Larus relictus*) among candidate genes associated with reproductive traits, including sex-ratio regulation and embryonic development, under varying environmental conditions (Yang et al., 2025). These gene-level signals should therefore be viewed as candidate hypotheses for exceptional cases of convergent local adaptation, rather than as evidence that selection broadly explains Cerrado–Amazon differentiation.

Finally, the absence of strong GO enrichment should also be interpreted with caution. Rather than indicating that candidate genes are biologically irrelevant, it suggests that the detected loci are functionally heterogeneous and not concentrated within a small number of annotated biological pathways. This limitation is particularly important in non-model organisms, where incomplete annotation can reduce the power of enrichment analyses. In this study, more than 50% of candidate and background genes lacked annotated GO terms (Table S19), meaning that additional convergent or functionally related signals may have gone undetected.

### Conservation implications

It has been presumed that humans contributed significantly to the extinction of at least two macaw species, endemic to the Caribbean Islands, due to hunting or trade (Olson & López, 2008; Wiley & Kirwan, 2013), leaving few records of those species. In addition, on the mainland of South America, human activity is probably the cause of the dramatic decline of the critically endangered Blue-throated macaw (*A. glaucogularis*) in the last 50 years (Herzog et al., 2023). While the widespread *Ara* species are not considered as threatened, genomic studies are important to evaluate potentially overlooked and vulnerable populations and establish conservation units. This kind of studies have been barely applied to neotropical parrot research so far.

Genetic studies of Neotropic species in general are very limited due to limited access to funds, genetic lab facilities, and trained labor (Napolitano et al., 2024). Moreover, research on parrots is heavily regulated under CITES, and the process to obtain research permits can be long, and different from country to country. This makes it hard to attempt sampling across the full species’ range. Although the advent of high-throughput sequencing has allowed us to obtain genomic data from birds coming from natural history collections, modern genomic research is more cost-effective when non-degraded fresh tissue is available to ensure high endogenic content. In general the endogenic content did not differ significantly between toe pad and feather samples (Figure S1). However, we found best results when the DNA extraction was carried out in freshly molted feathers rather than older museum toe pads. This highlights the value of freshly collected feathers as a practical and non-invasive alternative to blood samples.

Based on the presented evidence we suggest that more conservation initiatives are necessary in the Cerrado populations of *A. ararauna* and *A. chloropterus* as they exhibit high genetic differentiation and extremely low heterozygosity, which can correlate with threatened levels following the IUCN (Jeon et al., 2024). Outside the Cerrado, individuals exhibiting low heterozygosity were consistently located deeper within central Amazonia. Sampling in the Amazon, especially in its most remote areas, is notoriously challenging, as reflected by the geographic bias in natural history collections. It is possible that the low heterozygosity observed in these individuals reflects human activity where sampling is more accessible. Particularly, across *A. ararauna* and *A. chloropterus*, we find consistent low heterozygosity in Iquitos, the largest city of the Peruvian Amazon. This suggests that conservation initiatives are necessary in the region. To date, no substantial population decline has been reported in *A. severus*, although local population declines have been reported in its Venezuelan distribution (Collar et al., 2020b). Coincidentally, we find a distinct and structured population, based on both mitochondrial and nuclear data, in this threatened region.

We also acknowledge the role of captive individuals in reintroduction efforts to help critically endangered parrot populations (C. I. Campos et al., 2021; Estrada, 2014). However, to minimize risks such as outbreeding depression, or inbreeding, it is important to consider the genetic background of potential founders (Guhlin et al., 2023). For instance, our analysis shows the nuclear DNA of the captive species of *A. chloropterus* from Europe can be assigned to the East Amazon population, underscoring the power of population genomics to determine the ancestry of global captive macaws when a genomic reference database is available. Moreover, we did not find a signal of admixture even at higher *K* values in the admixture analysis (Figure S23), suggesting no mixing in captivity with individuals from other populations. However, we also detected that its heterozygosity levels were the lowest across all individuals analysed, which can also reflect inbreeding in captivity. Despite this, it is possible that captive individuals retain genetic variants no longer present in wild populations. We hope that this and future genomic works will provide guidance for potential genetic rescue initiatives, ensuring that reintroduction strategies are genetically informed.

Finally, this study also represents the most comprehensive study of Neotropical parrot populations based on complete mitochondrial sequences to date, and significantly expands the available mitochondrial data for *Ara* species, for which only five complete mitogenomes have been previously assembled and annotated for the studied species (D.-W. Liu et al., 2020; Urantowka et al., 2017; Urantówka et al., 2017). The phylogeographic patterns based on complete mitochondrial sequences partially captures the broad-scale geographic patterns observed in the SNP data (Figure 2 and 3). Thus, as the geographic pattern in nuclear genetic structure resembles that of the mitochondrial genome, we suggest that mitochondrial data alone can be used as a budget friendly proxy for genetic structure on these species. This information is also relevant when the sample material is not optimal, like in natural history skin collections or even biological samples found in archaeological sites.

## Acknowledgments

This study was supported by funds granted by NTNU to MDM as part of the Onsager Fellowship Programme, as well as by the Norwegian Directorate for Higher Education and Skills (HKdir) through NORPART award 2021/10475 (“BiGTREE: Biodiversity Genomics Teaching, Research, Exchanges and Education”). Funding was also provided in part by the Conselho Nacional de Desenvolvimento Científico e Tecnológico (CNPq, 306989/2023-9) and Fundación de Apoyo a la Investigación del Estado de São Paulo (FAPESP, 2013/50297-0) from Brazil and the Research Council of Norway project 326819 (Earth BioGenome Project Norway). We thank the Peruvian government for the research and sampling permit issued by the local authorities (SERFOR N° AUT-IFS-2022-042). Samples from Peru were exported to Norway under CITES permits PE005039/SP, PE005040/SP, and PE005045/SP, and samples from Brazil were exported to Norway under CITES 24BR048160/DF. We thank Constanza de la Fuente Castro for her valuable support in conducting lab work at the University of Chicago. The authors acknowledge support from the Norwegian National Infrastructure for High Performance Computing and resources provided by Sigma2 as well as Data Storage in Norway (project NN8013K).

## Data availability

Gene annotations and predicted proteins for *Ara ararauna, Ara chloropterus* and *Ara severus* are available at Zenodo at https://doi.org/10.5281/zenodo.19853893. Raw sequencing data generated for this study has been deposited on the European Nucleotide Archive under project accession code PRJEB91554.

## Contributions

JML conceived the original idea for the study. JML, MDM, and VCB designed the study. JML wrote the paper with input from all authors. JML carried out fieldwork and obtained feather samples from the Peruvian Amazon. LS, CYM, JB, TH, assisted in sample acquisition and preparation. JML and TH performed molecular preparations on the samples. JML and OKT analysed the data with input from MDM, and VCB. JML, MDM, VCB, contributed to interpreting the results.

## Notes

https://doi.org/10.5281/zenodo.19853893

### Competing Interest Statement

The authors have declared no competing interest.

