## Supplementary Material for "Range-wide phylogeography, population genomics, and demography of three widespread *Ara* macaws (Psittacidae)"

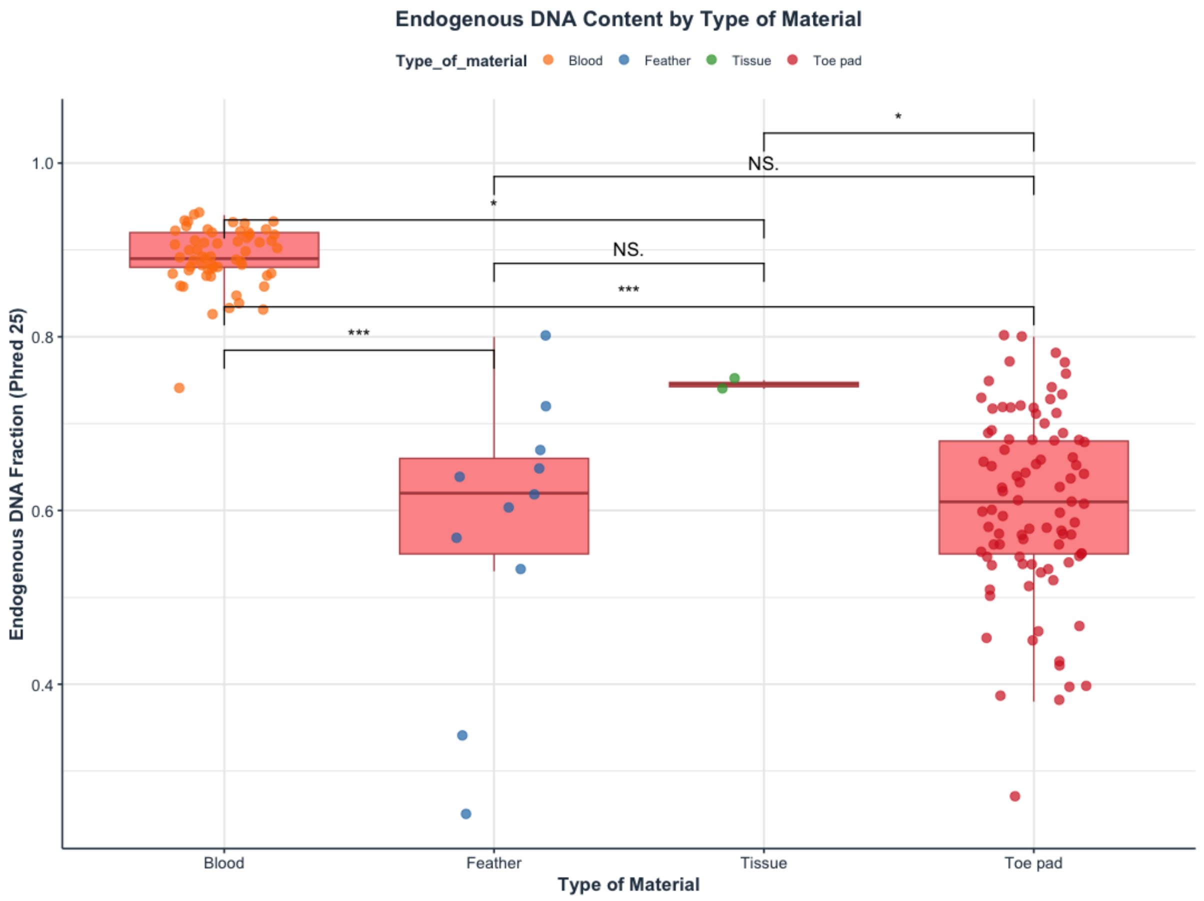

**Figure S1**. Endogenous DNA content (MAPQ > 25) across different sample types. Boxplots represent the distribution of endogenous content for each material type: blood, feather, toe pad, and tissue. Statistical significance was assessed using one-way ANOVA, followed by Tukey's HSD post-hoc test, with comparisons indicated by asterisks (*p < 0.05, **p < 0.01, ***p < 0.001). Asterisks above the boxplots indicate significant pairwise differences between sample types.

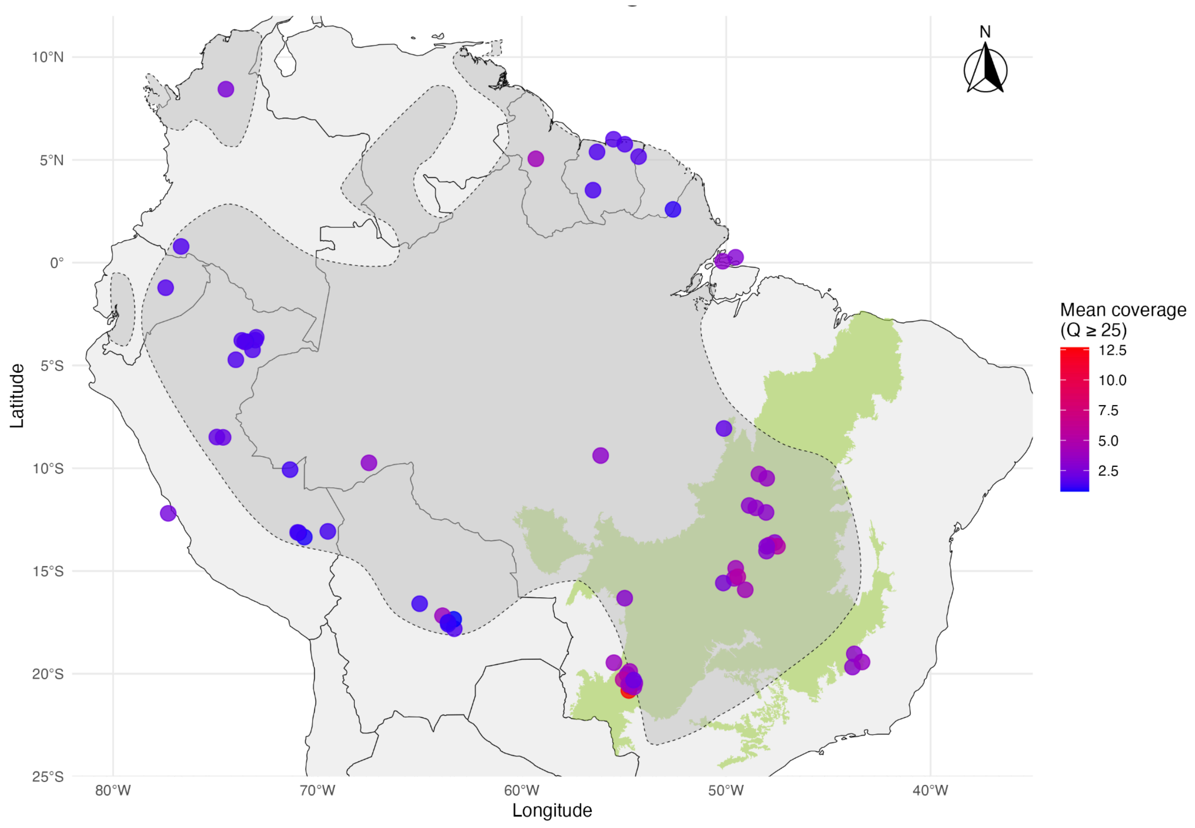

**Figure S2.** Geographic distribution of mean sequencing depth (Q ≥ 25) for *Ara ararauna* sampling localities. Background layers include the South American continent (light gray), the Cerrado biome (green), and the IUCN species distribution range (gray polygon). Locations were jittered slightly to minimize overlap.

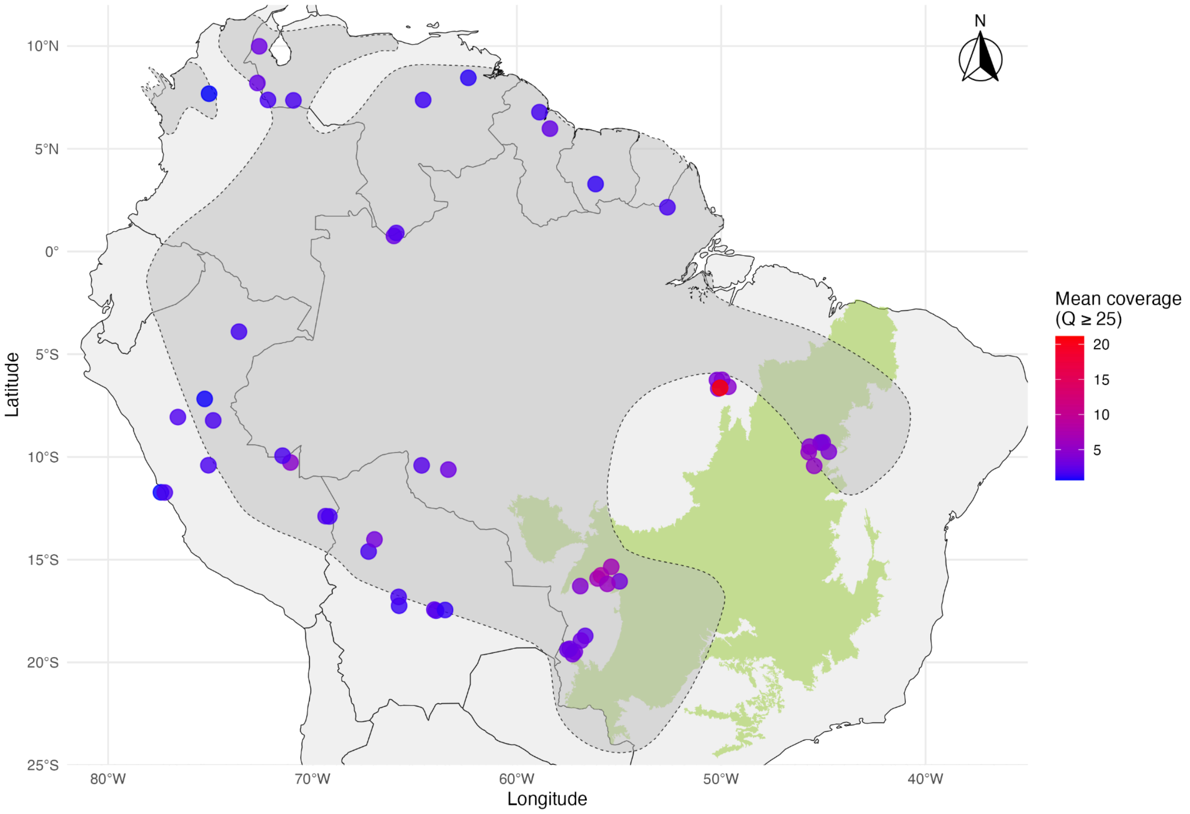

**Figure S3.** Geographic distribution of mean sequencing depth (Q ≥ 25) for *Ara chloropterus* sampling localities. Background layers include the South American continent (light gray), the Cerrado biome (green), and the IUCN species distribution range (gray polygon). Locations were jittered slightly to minimize overlap.

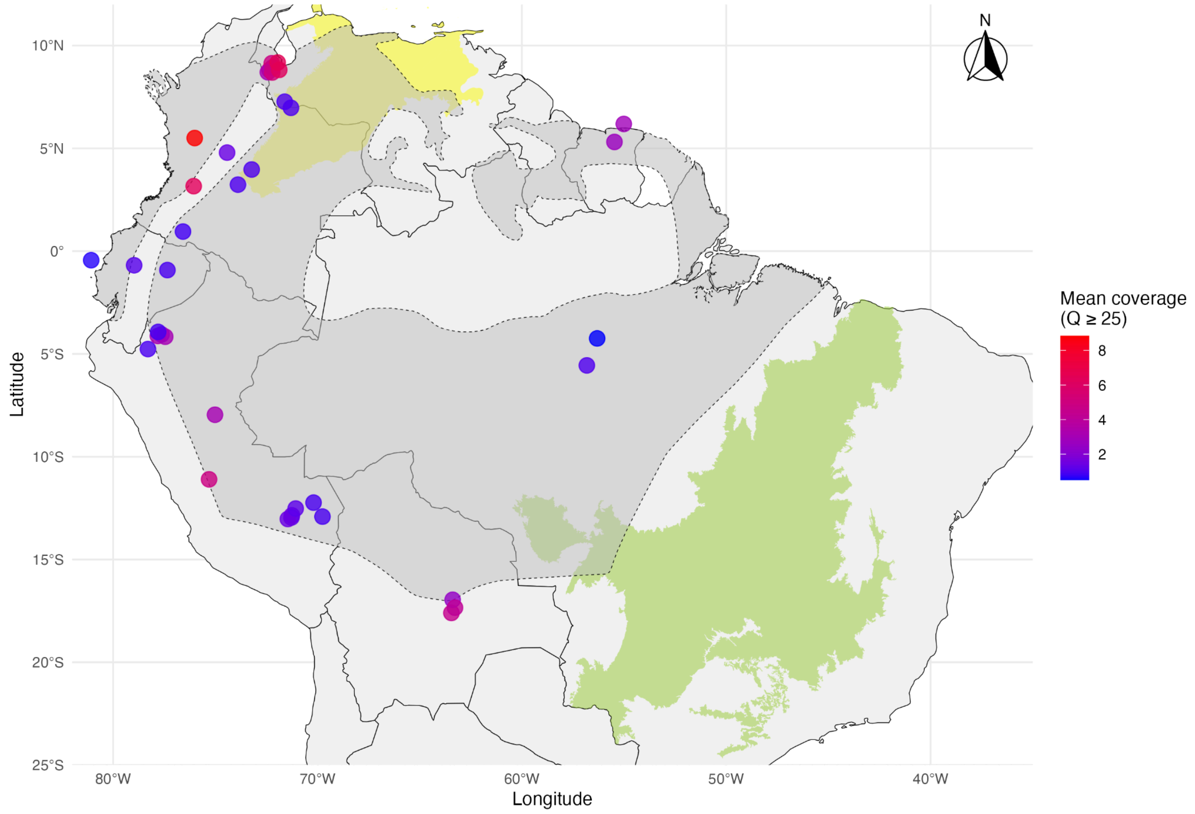

**Figure S4.** Geographic distribution of mean sequencing depth (Q ≥ 25) for *Ara severus* sampling localities. Background layers include the South American continent (light gray), the Cerrado biome (green), the Dry North of South America biome (yellow), and the IUCN species distribution range (gray polygon). Locations were jittered slightly to minimize overlap.

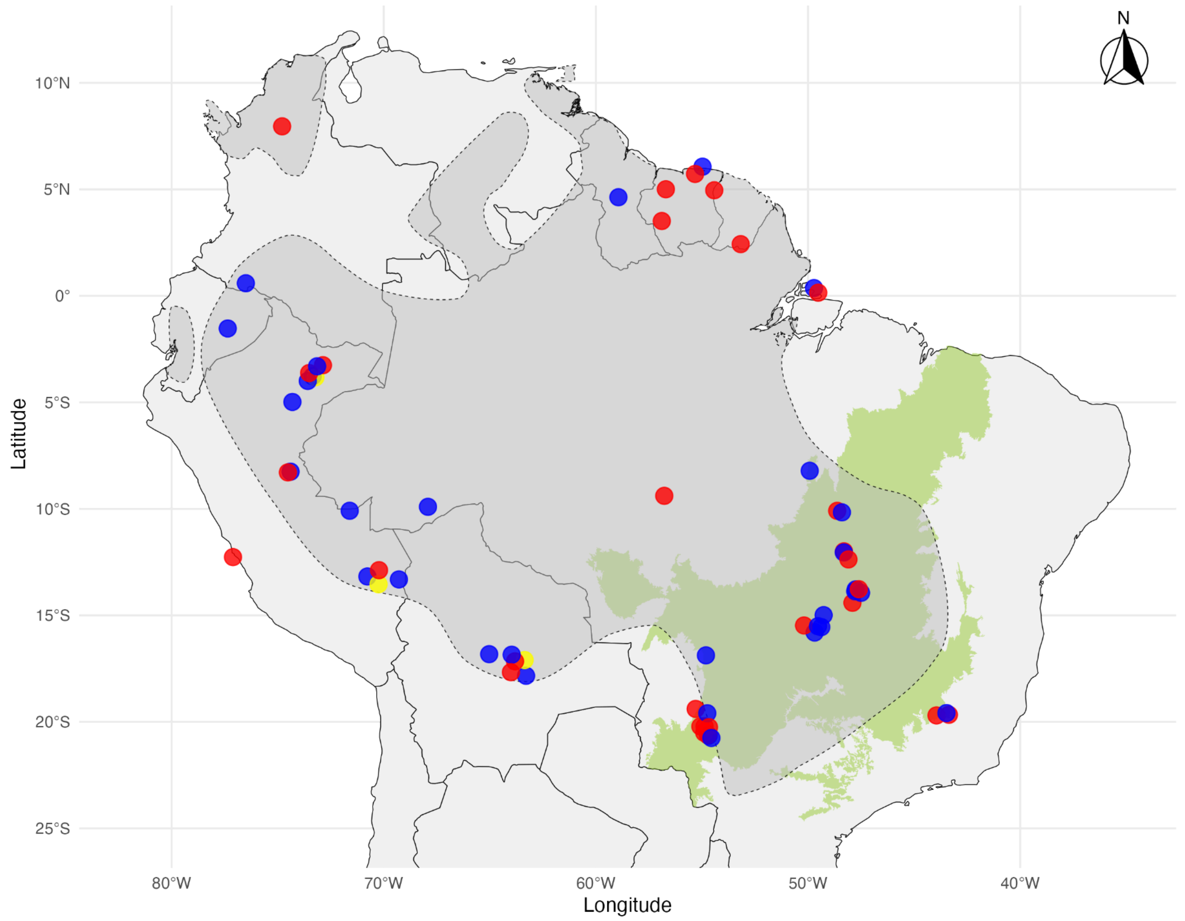

**Figure S5.** Geographic distribution of *Ara ararauna* individuals categorized by sex, based on Sex Assignment Through Coverage (SATC) analysis. Each point represents a sampled individual, colored by sex: blue for males, red for females, and yellow for individuals of undetermined sex. Background layers include the South American continent (light gray), the Cerrado biome (green), and the IUCN species distribution range (gray polygon). Locations were jittered slightly to minimize overlap.

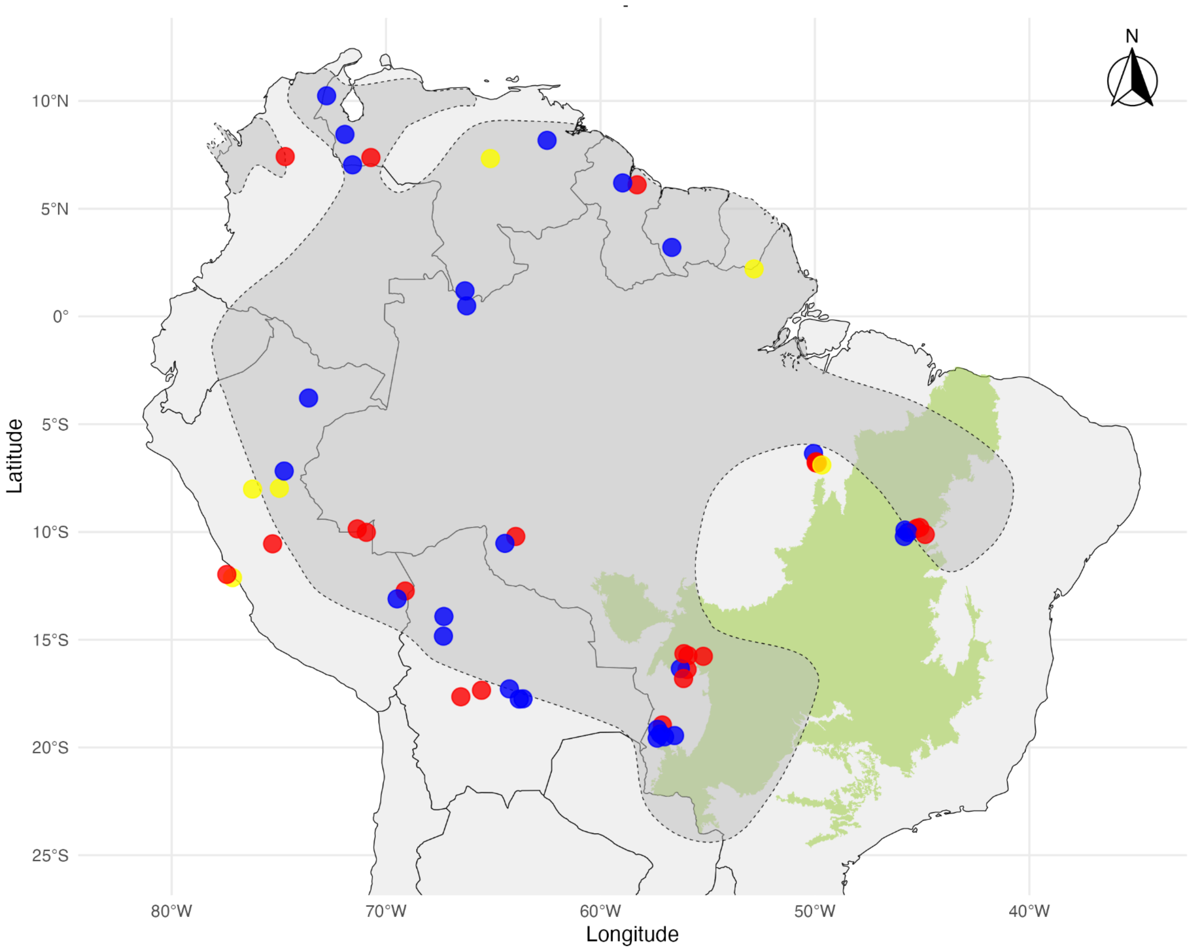

**Figure S6.** Geographic distribution of *Ara chloropterus* individuals categorized by sex, based on Sex Assignment Through Coverage (SATC) analysis. Each point represents a sampled individual, colored by sex: blue for males, red for females, and yellow for individuals of undetermined sex. Background layers include the South American continent (light gray), the Cerrado biome (green), and the IUCN species distribution range (gray polygon). Locations were jittered slightly to minimize overlap.

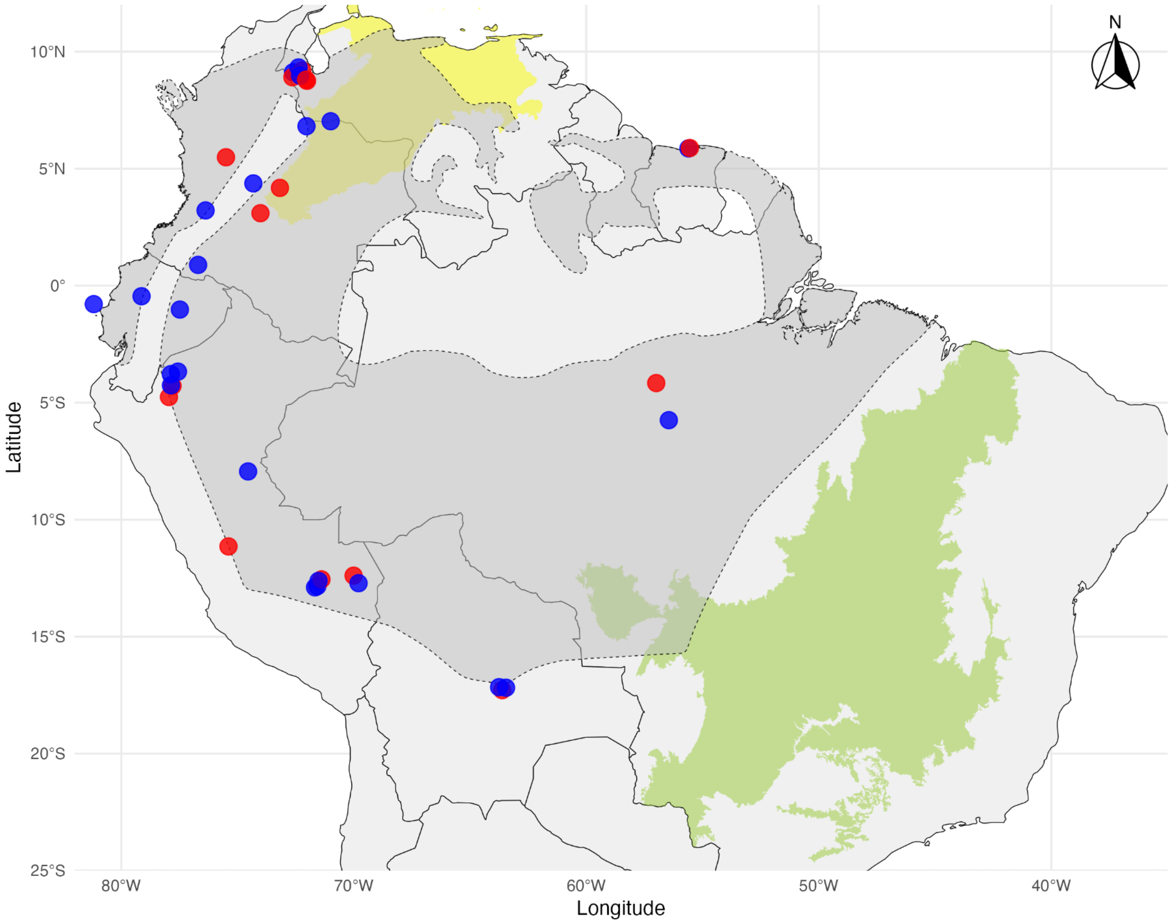

**Figure S7.** Geographic distribution of *Ara severus* individuals categorized by sex, based on Sex Assignment Through Coverage (SATC) analysis. Each point represents a sampled individual, colored by sex: blue for males, red for females, and yellow for individuals of undetermined sex. Background layers include the South American continent (light gray), the Cerrado biome (green), the Dry North of South America biome (yellow), and the IUCN species distribution range (gray polygon). Locations were jittered slightly to minimize overlap.

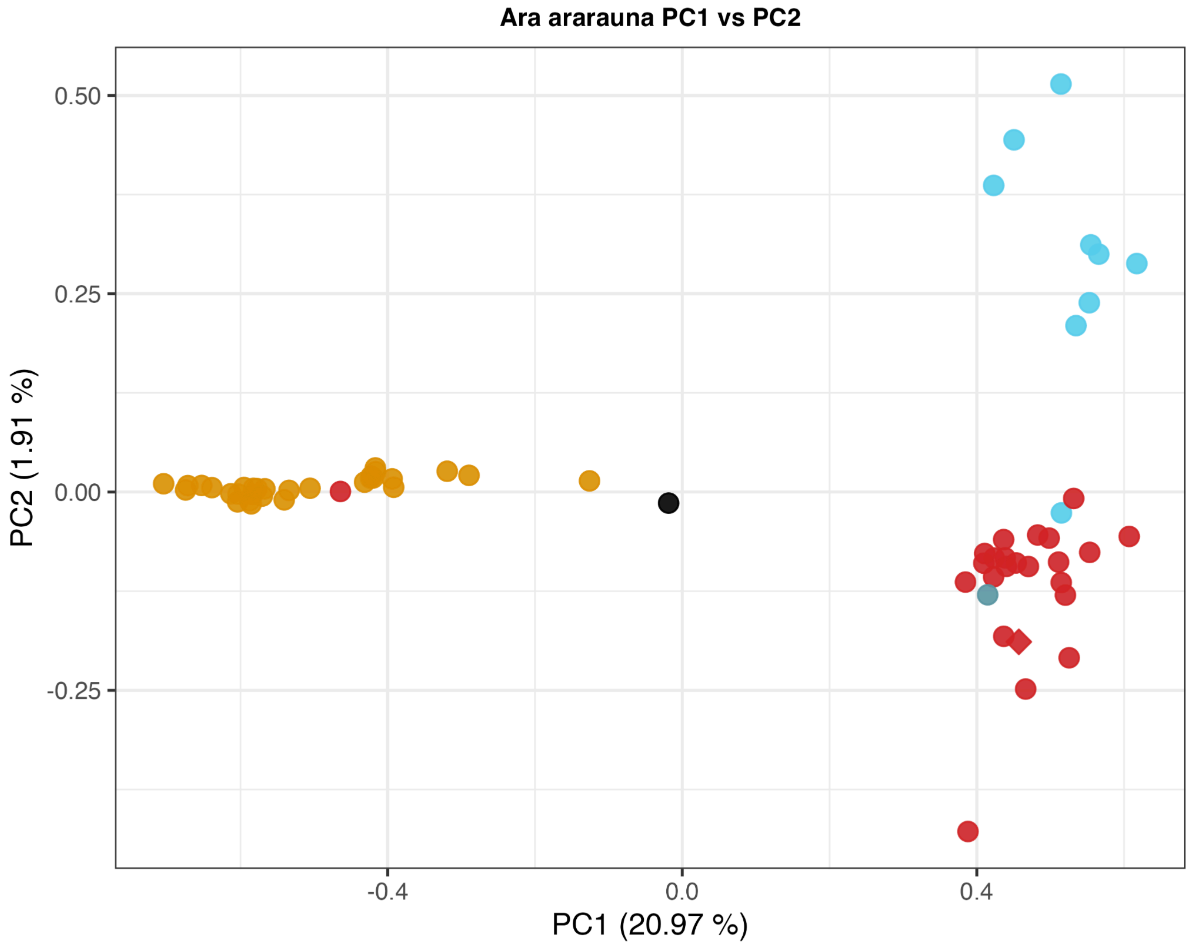

**Figure S8.** Principal component analysis (PC1 vs PC2) plot for *Ara ararauna*. Points are color-coded by the main populations analyzed: Cerrado (amber), Amazon-West (crimson), and Amazon-East (cyan). Diamond-shaped symbols represent captive specimens, while the isolated and "central" populations are differentiated in teal and black, respectively.

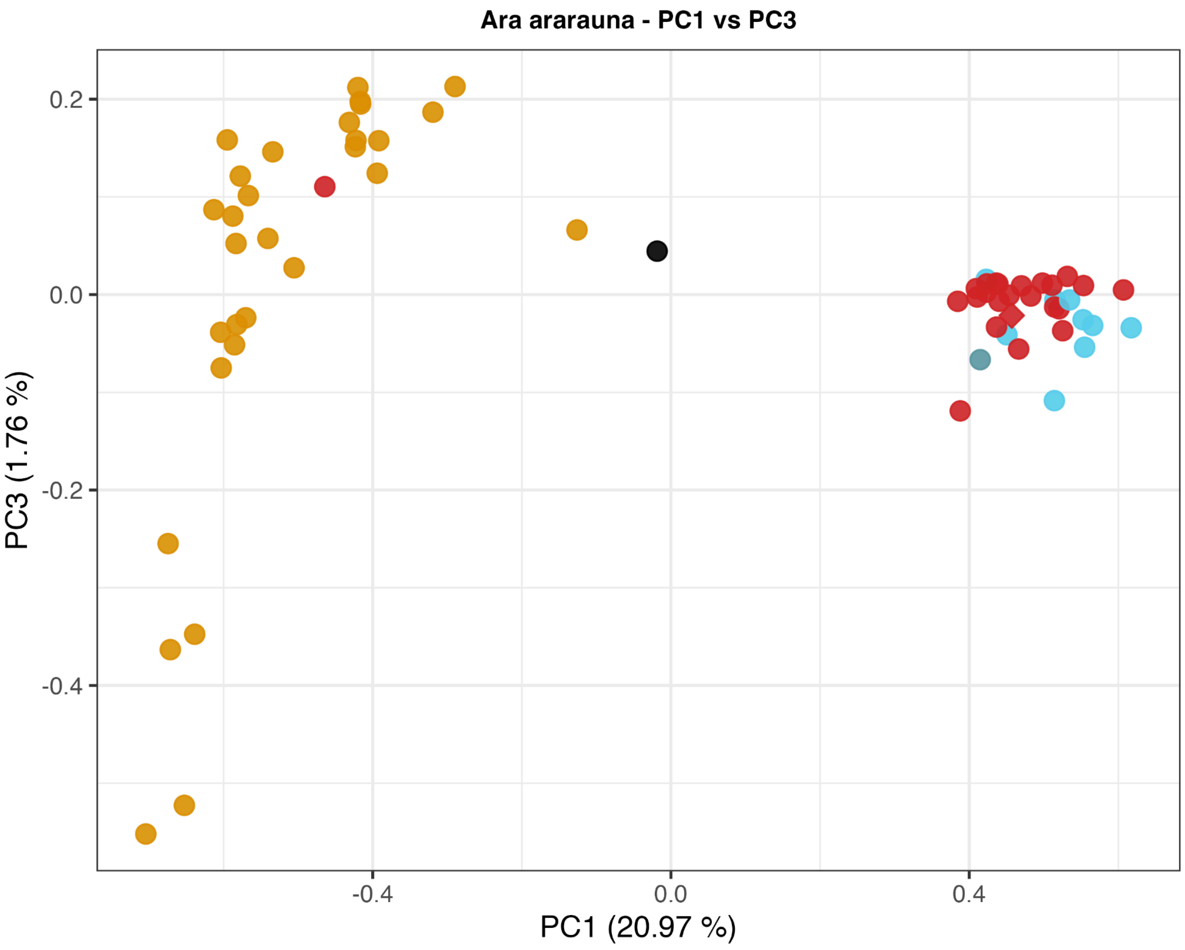

**Figure S9.** Principal Component Analysis (PC1 vs PC3) plot for *Ara ararauna*. Points are color-coded by the main populations analyzed: Cerrado (amber), Amazon-West (crimson), and Amazon-East (cyan). Diamond-shaped symbols represent captive specimens, while the isolated and "central" populations are differentiated in teal and black, respectively.

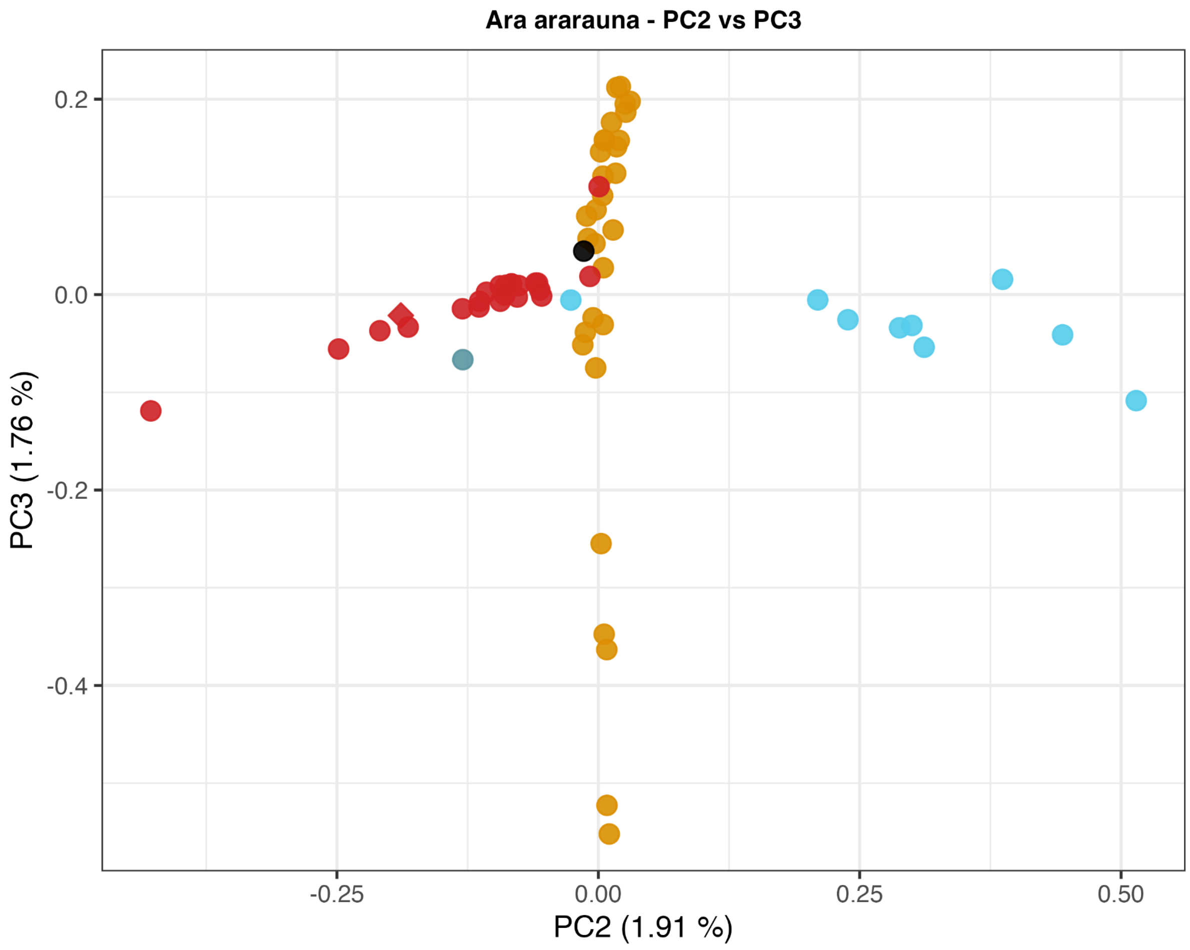

**Figure S10.** Principal Component Analysis (PC2 vs PC3) plot for *Ara ararauna*. Points are color-coded by the main populations analyzed: Cerrado (amber), Amazon-West (crimson), and Amazon-East (cyan). Diamond-shaped symbols represent captive specimens, while the isolated and “central” populations are differentiated in teal and black, respectively.

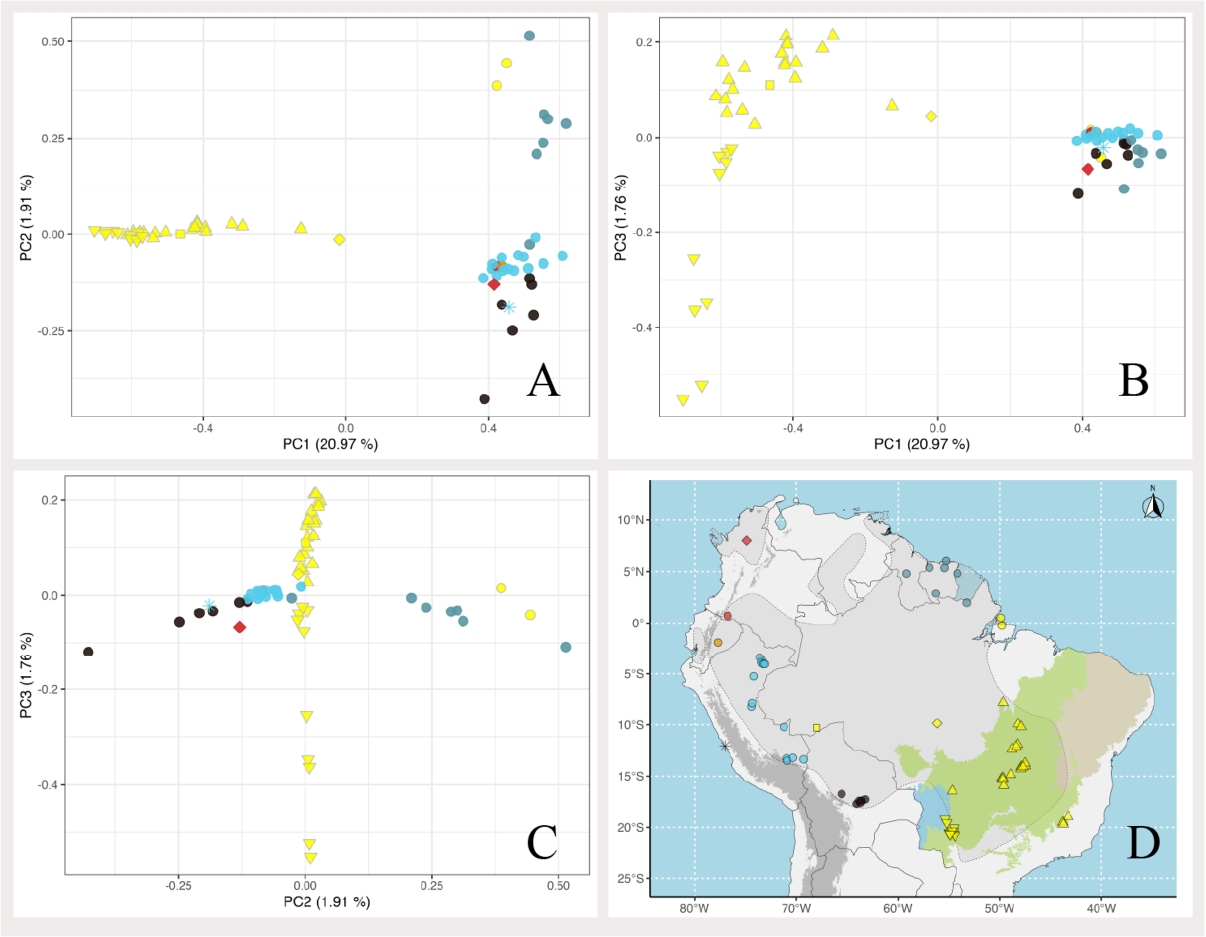

**Figure S11.** Exploratory PCA plots in *Ara ararauna*: PC1 vs PC2 (A), PC1 vs PC3 (B), and PC2 vs PC3 (C). Points are color-coded by country, and distinctive shapes are used to differentiate localities as depicted in the map (D). A captive individual from Lima (Peru) is shown by an asterisk, and the isolated population is diamond-shaped. The distribution range of the species is delimited with dotted lines, and the areas are shaded in gray. Elevations over 3000 meters a.s.l. are highlighted in dark grey and roughly correspond to the Andes. The Cerrado biome is highlighted in Brazil.

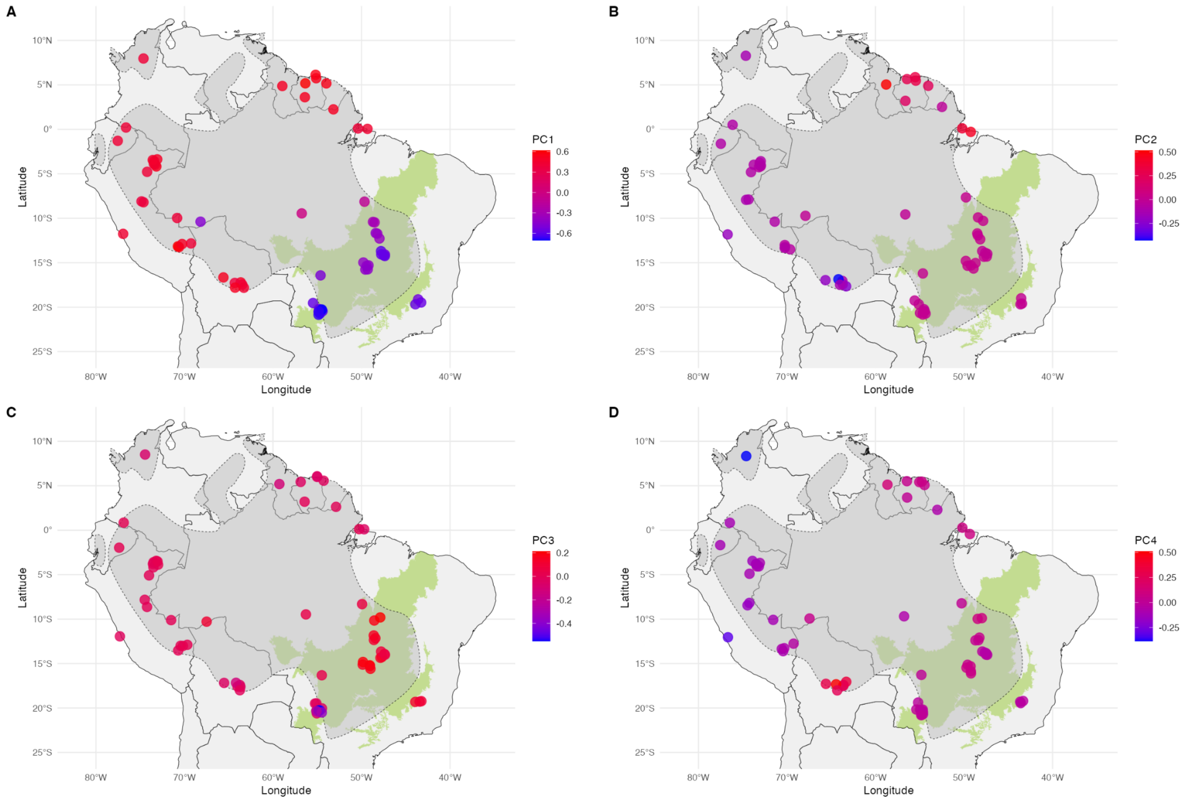

**Figure S12.** Spatial plots of principal components PC1 (A), PC2 (B), PC3 (C), and PC4 (D) across the range of *Ara ararauna.* Background layers include the South American continent (light gray), the Cerrado biome (green), and the IUCN species distribution range (gray polygon). Locations were jittered slightly to minimize overlap.

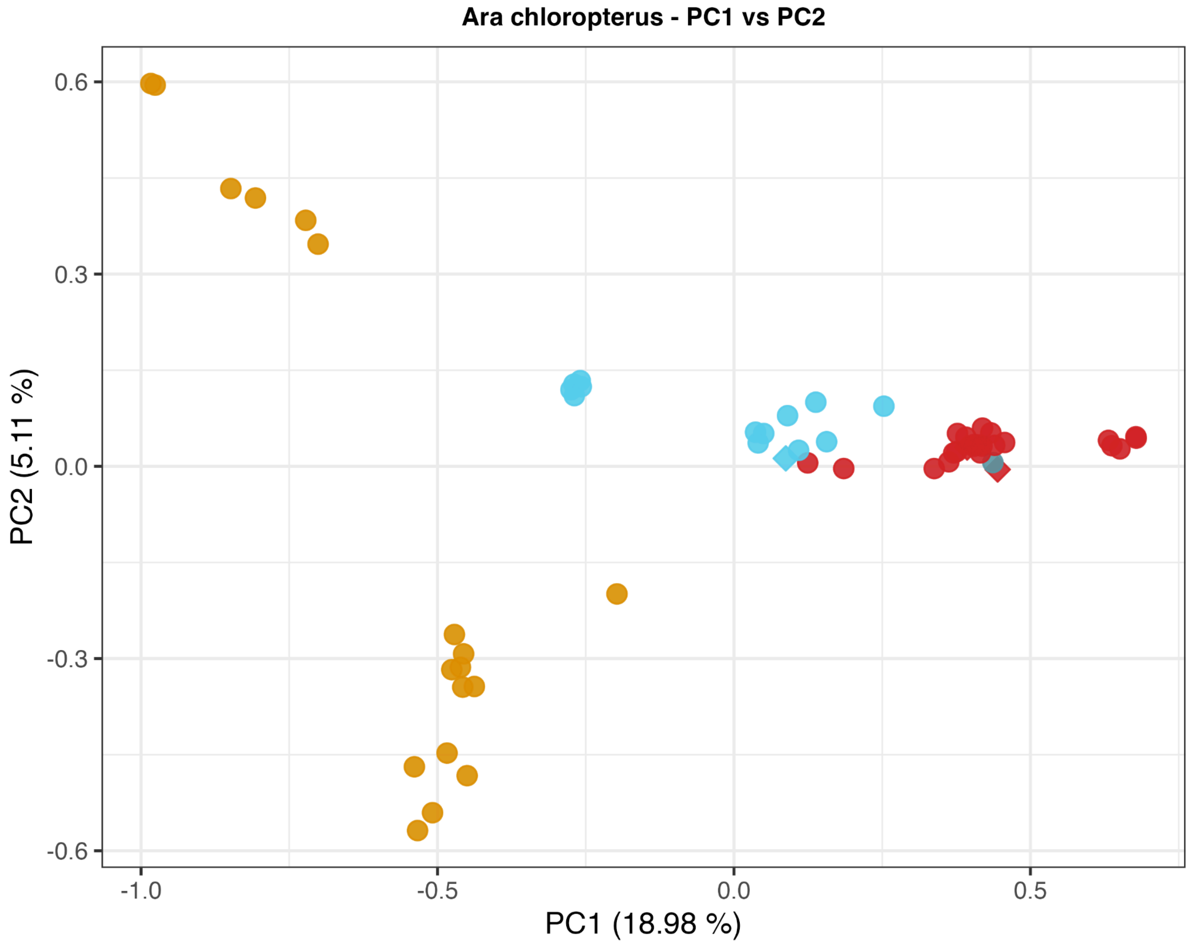

**Figure S13.** Principal Component Analysis (PC1 vs PC2) plot for *Ara chloropterus*. Points are color-coded by main populations analyzed: Cerrado (amber), Amazon-West (crimson), Amazon-East (cyan); the captive individuals from Norway and Lima (Peru) are diamond-shaped in cyan and crimson, respectively. The isolated population is shown with a teal dot.

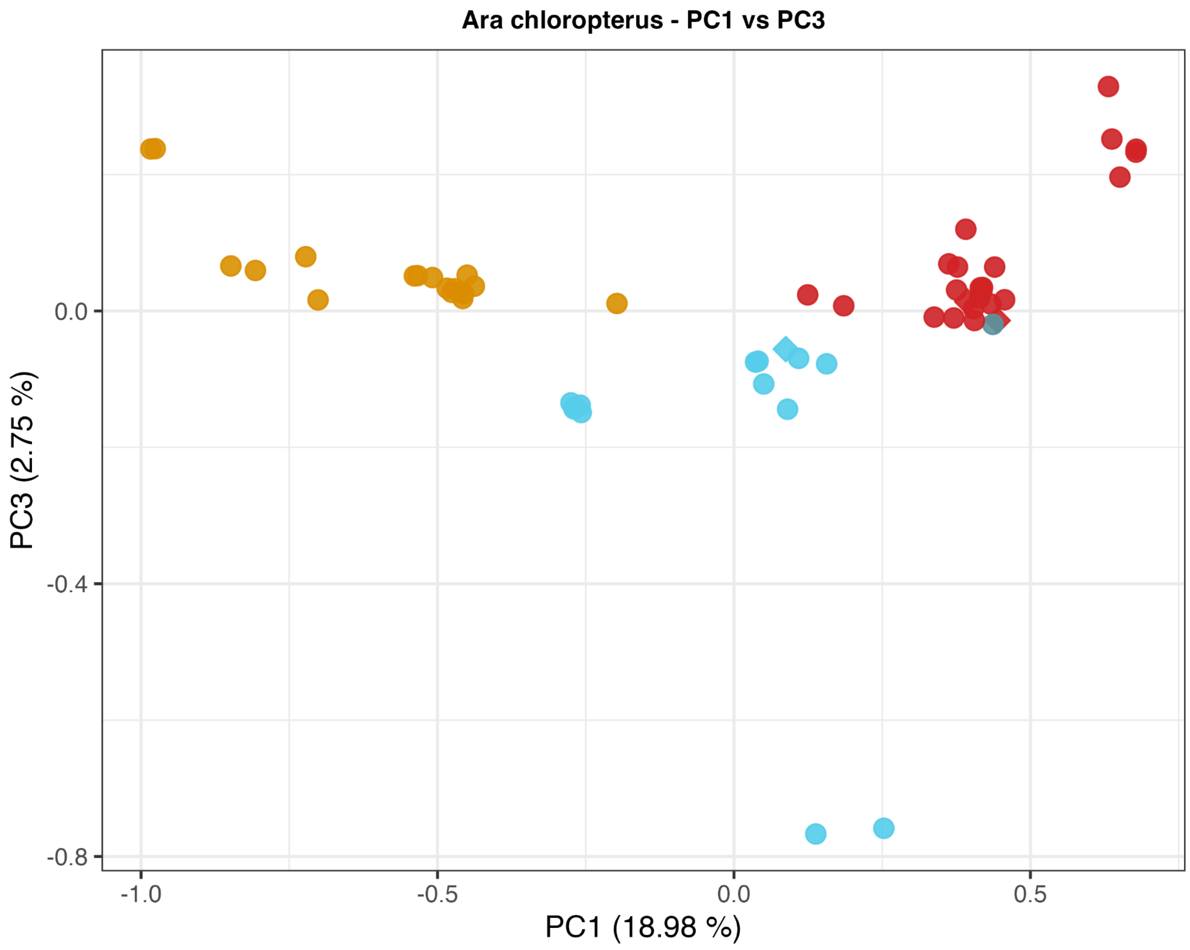

**Figure S14.** Principal component analysis (PC1 vs PC3) plot for *Ara chloropterus*. Points are color-coded by main populations analyzed: Cerrado (amber), Amazon-West (crimson), Amazon-East (cyan); the captive individuals from Norway and Lima (Peru) are diamond-shaped in cyan and crimson, respectively. The isolated population is shown with a teal dot.

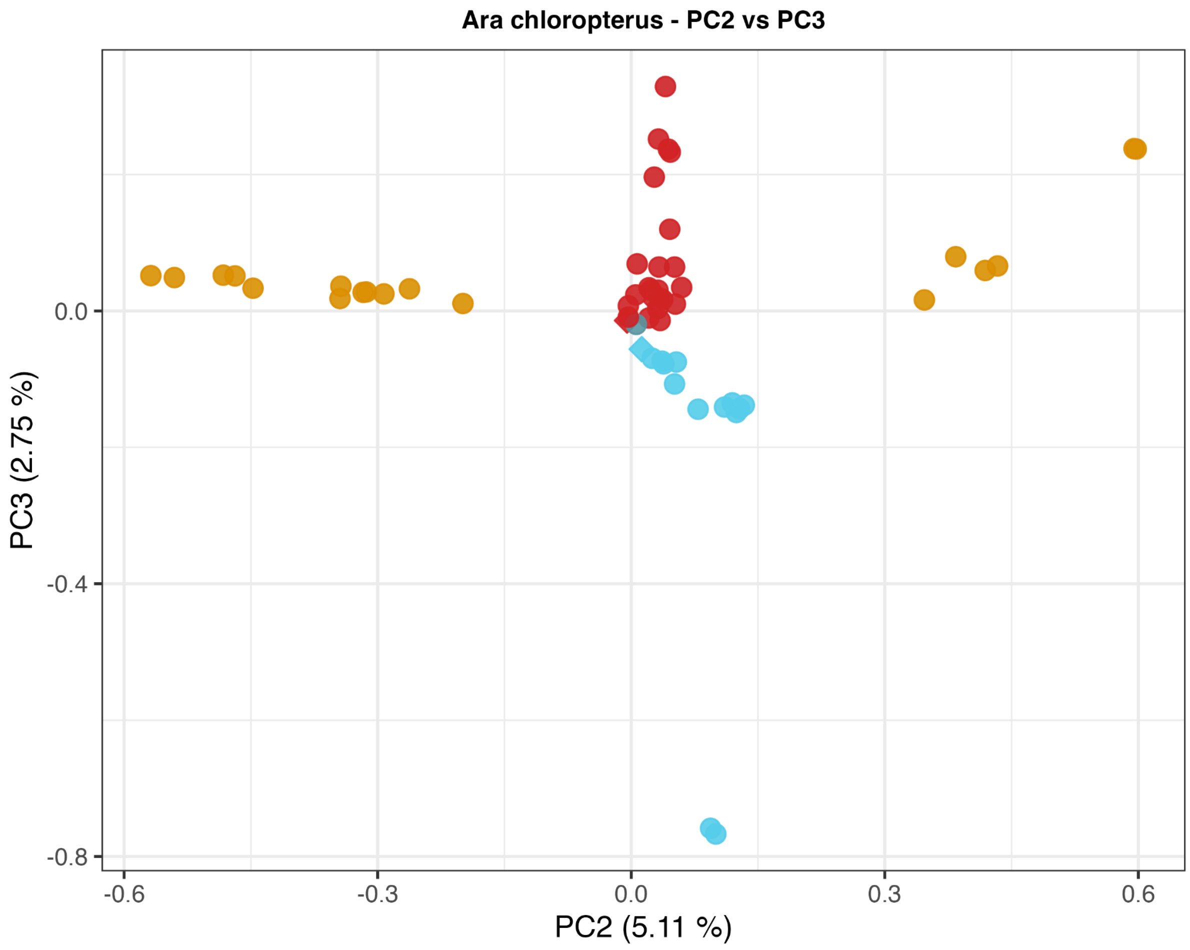

**Figure S15.** Principal Component Analysis (PC2 vs PC3) plot for *Ara chloropterus*. Points are color coded by main populations analyzed: Cerrado (amber), Amazon-West (crimson), Amazon-East (cyan); the captive individuals from Norway and Lima (Peru) are diamond-shaped in cyan and crimson, respectively. The isolated population is shown with a teal dot.

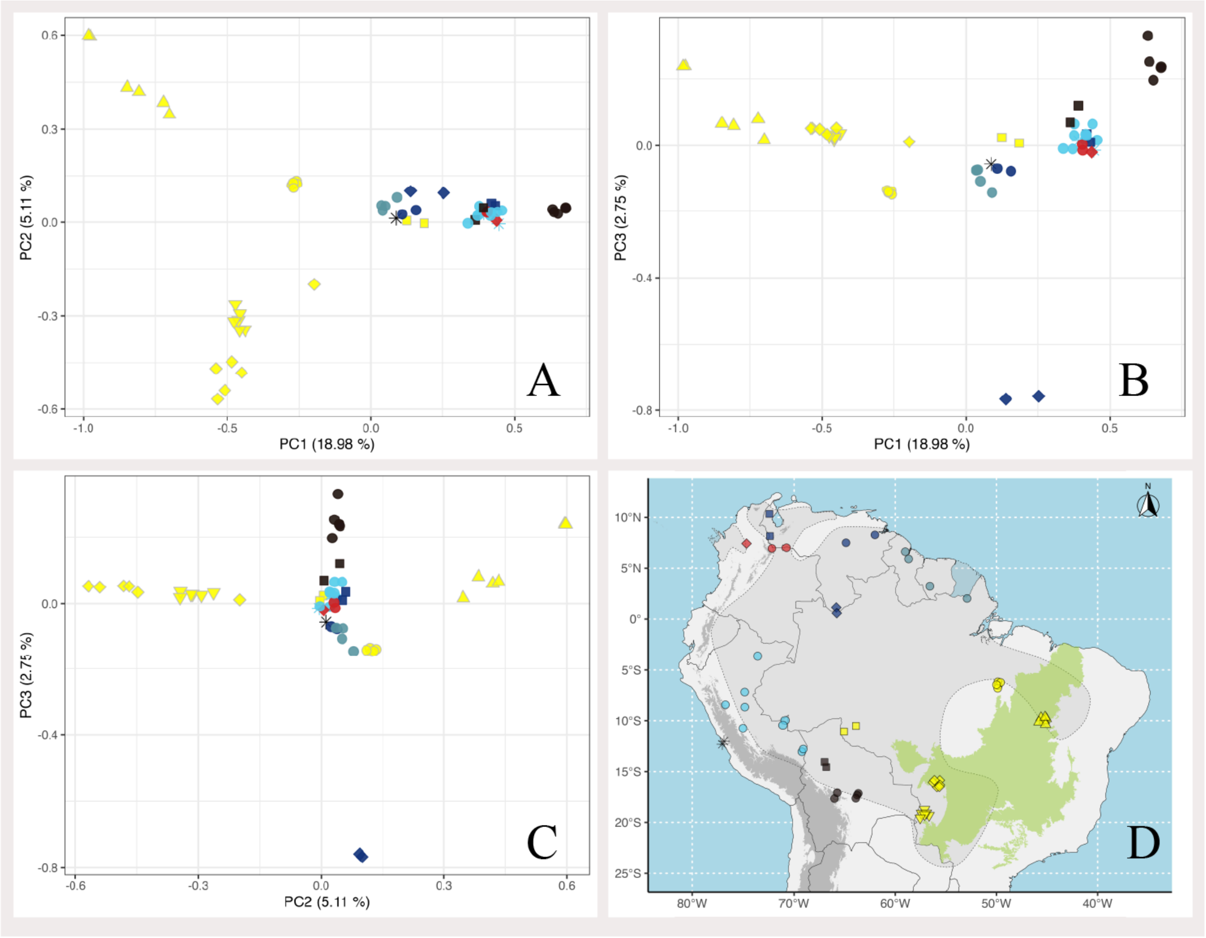

**Figure S16.** Exploratory PCA plots in *Ara chloropterus*: PC1 vs PC2 (A), PC1 vs PC3 (B), and PC2 vs PC3 (C). Points are color-coded by country, and distinctive shapes are used to differentiate localities as depicted in the map (D). On the PCA plot, captive individuals from Lima (Peru) are shown by an asterisk of the color of the country, and the captive individual from Norway is shown by a black asterisk. The distribution range of the species is delimited with dotted lines, and the areas are shaded in gray. Elevations over 3000 meters a.s.l. are highlighted in dark grey and roughly correspond to the Andes. The Cerrado biome is highlighted in Brazil.

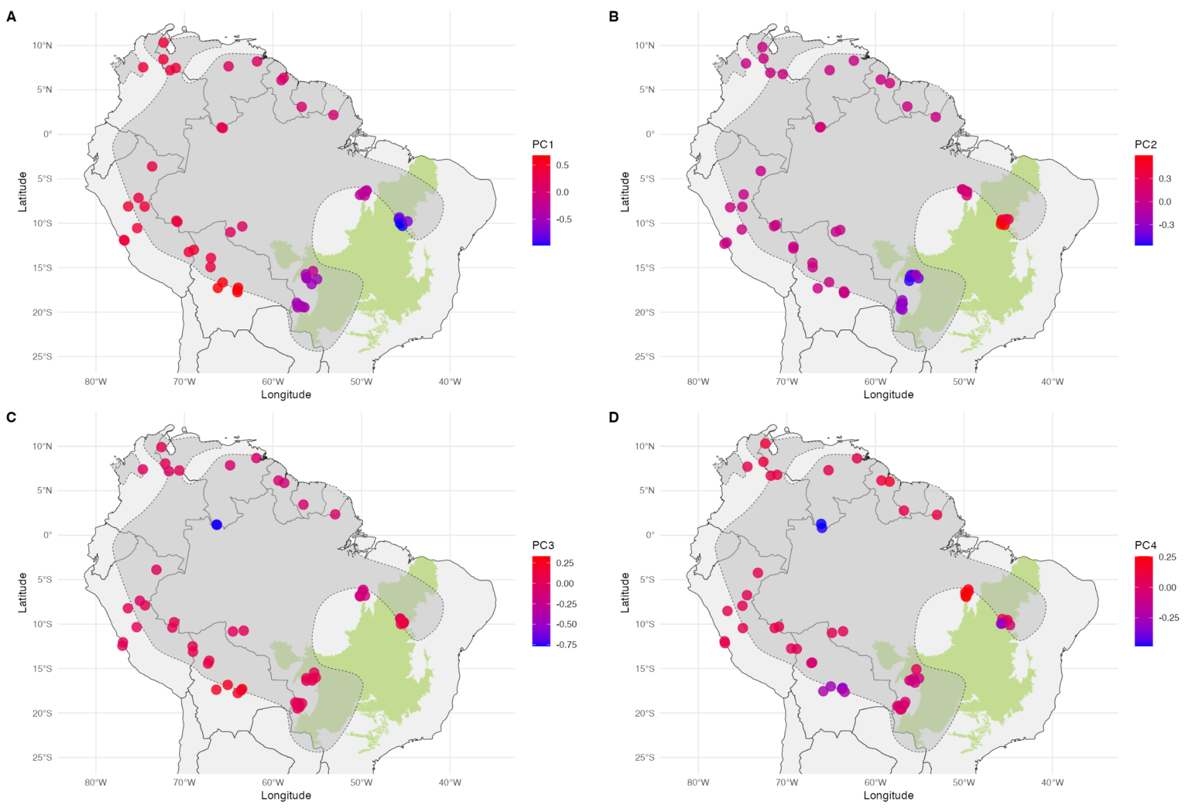

**Figure S17.** Spatial plots of principal components PC1 (A), PC2 (B), PC3 (C), and PC4 (D) across the range of *Ara chloropterus*. Background layers include the South American continent (light gray), the Cerrado biome (green), and the IUCN species distribution range (gray polygon). Locations were jittered slightly to minimize overlap.

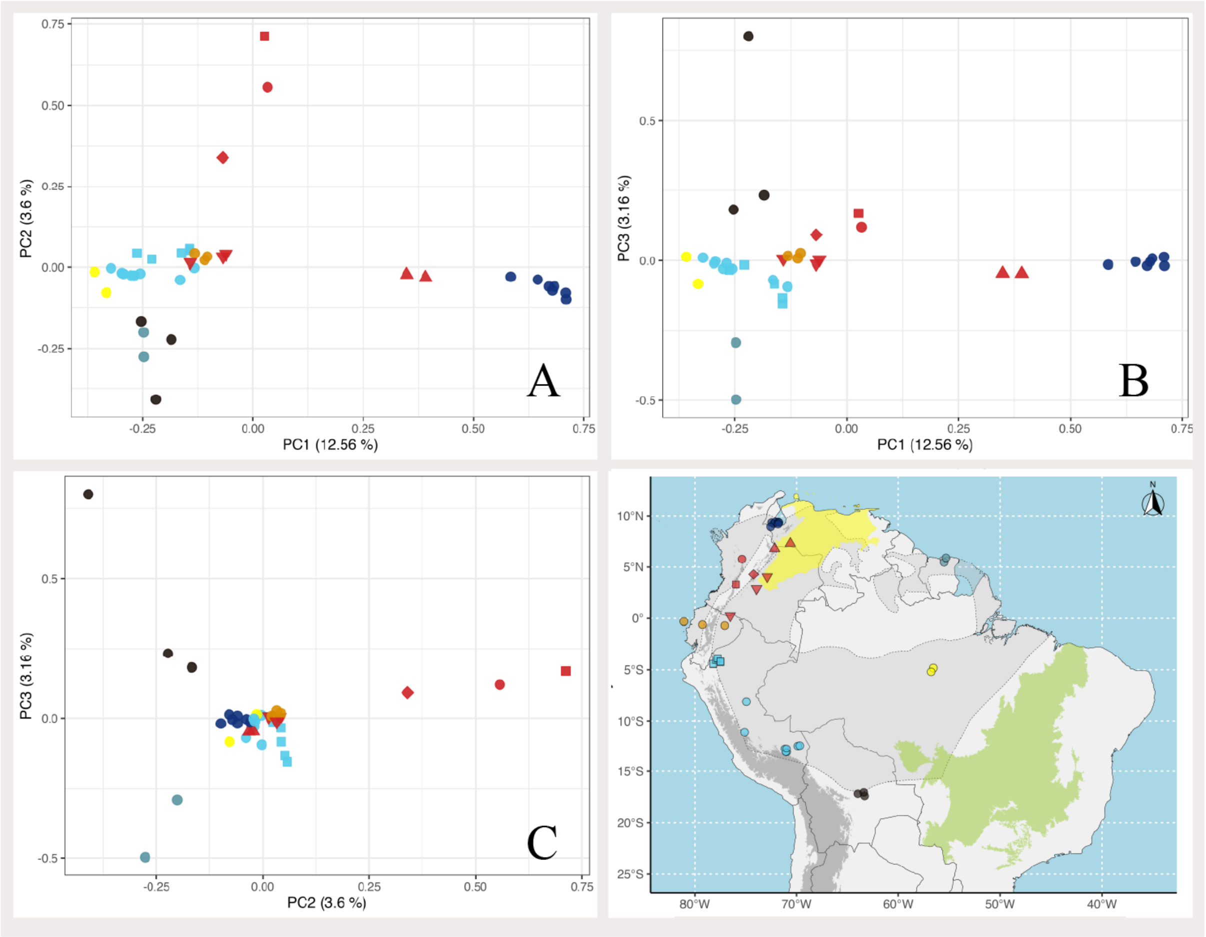

**Figure S18.** PCA plots in *Ara severus*: PC1 vs PC2 (A), PC1 vs PC3 (B), and PC2 vs PC3 (C). Points are color-coded by country, and distinctive shapes are used to differentiate localities as depicted in the map (D). The distribution range of the species is delimited with dotted lines, and the areas are shaded in gray. Elevations over 3000 meters a.s.l. are highlighted in dark grey and roughly correspond to the Andes. The Cerrado biome is highlighted in green in Brazil, the Dry North South America biome is highlighted in yellow.

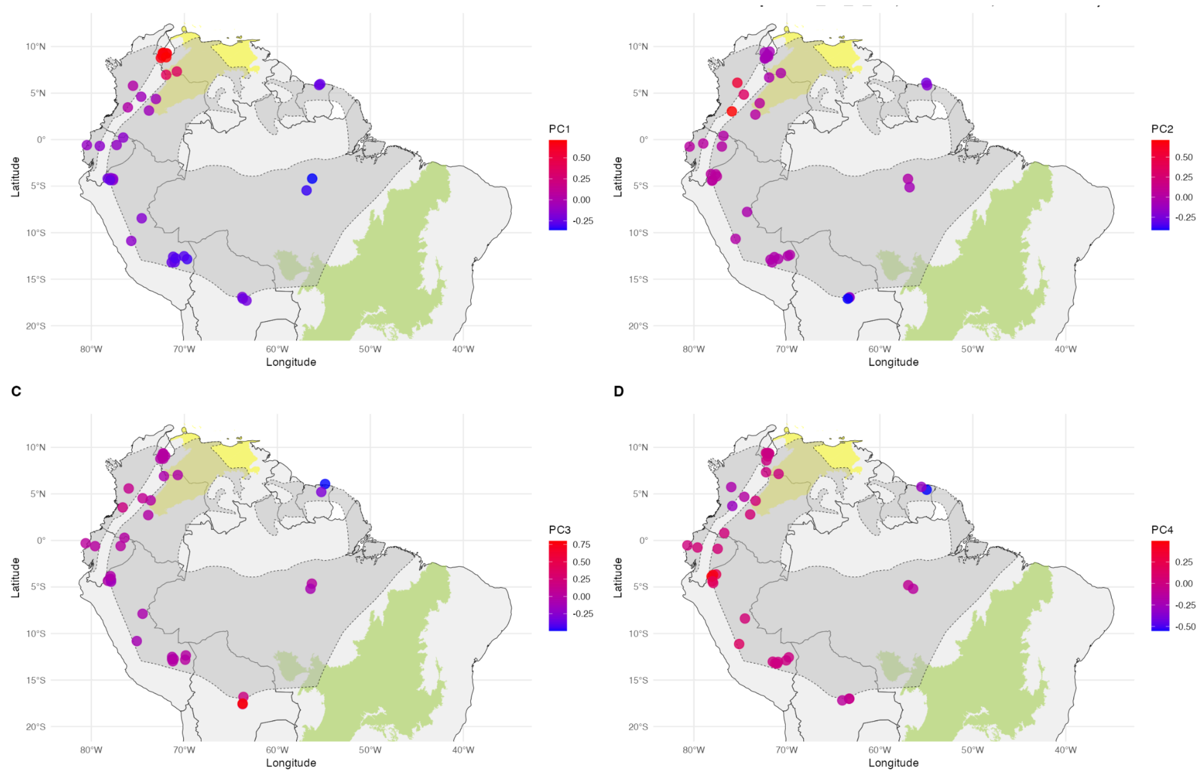

**Figure S19.** Spatial plots of principal components PC1 (A), PC2 (B), PC3 (C), and PC4 (D) across the range of *Ara severus*. Background layers include the South American continent (light gray), the Cerrado biome (green), and the IUCN species distribution range (gray polygon). Locations were jittered slightly to minimize overlap.

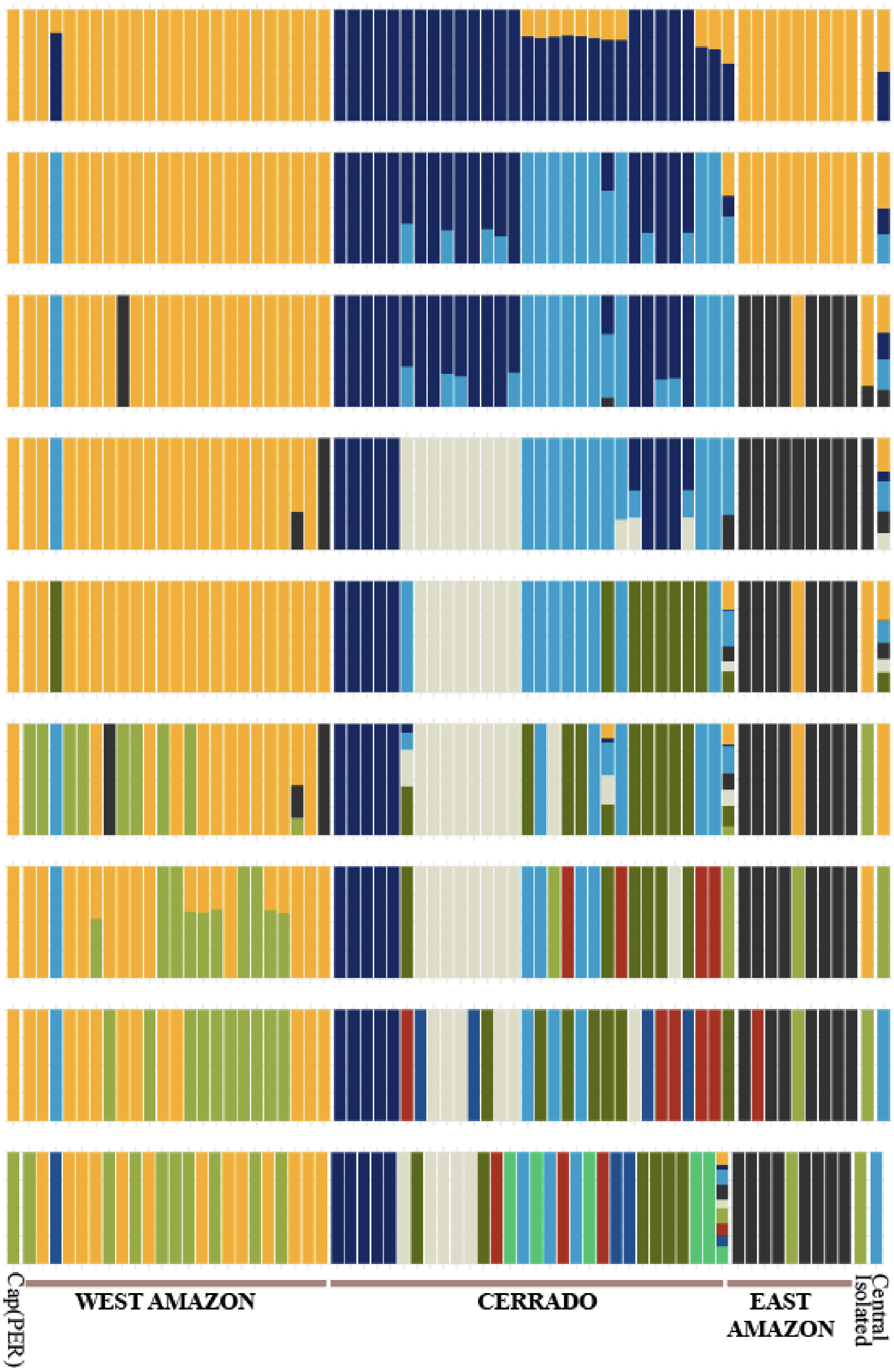

**Figure S20.** Admixture analysis of *Ara ararauna* (K2 to K10). Bar plots are grouped according to the biome of origin (Amazon-West, Amazon-East, and Cerrado). The bar plots differentiate captive individuals, the individual representing the isolated populations, and the individuals that don't belong to either biome, here referred to as the "central" population.

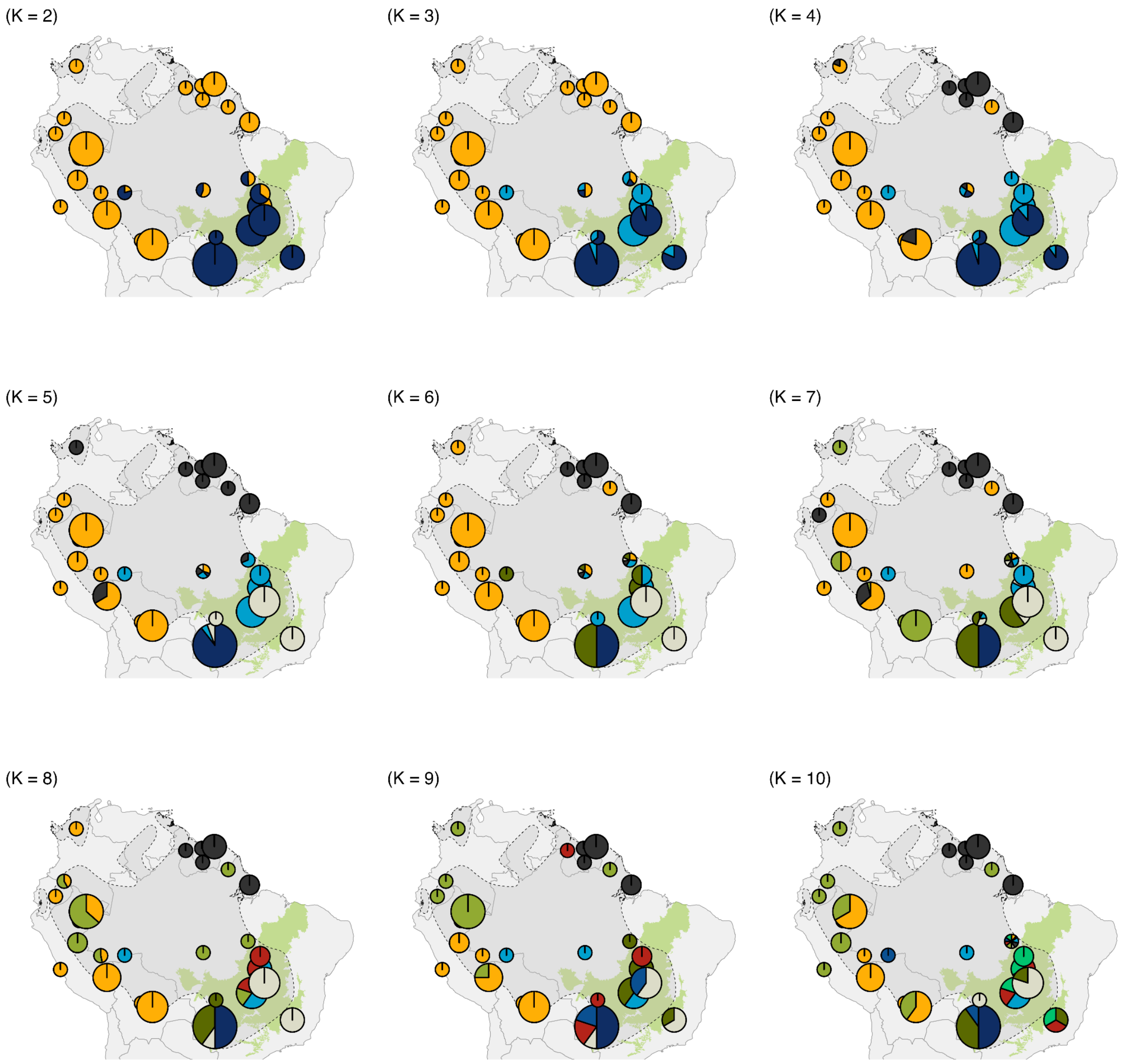

**Figure S21.** Admixture results for *Ara ararauna* plotted by sampling location, with individuals clustered within 200 km of spatial proximity. Each pie chart represents ancestry proportions for a given value of K (2 to 10). The Cerrado biome is shown in light green, and the species' IUCN distribution is shaded in dark grey with dashed outlines. The basemap includes country boundaries from South America for reference.

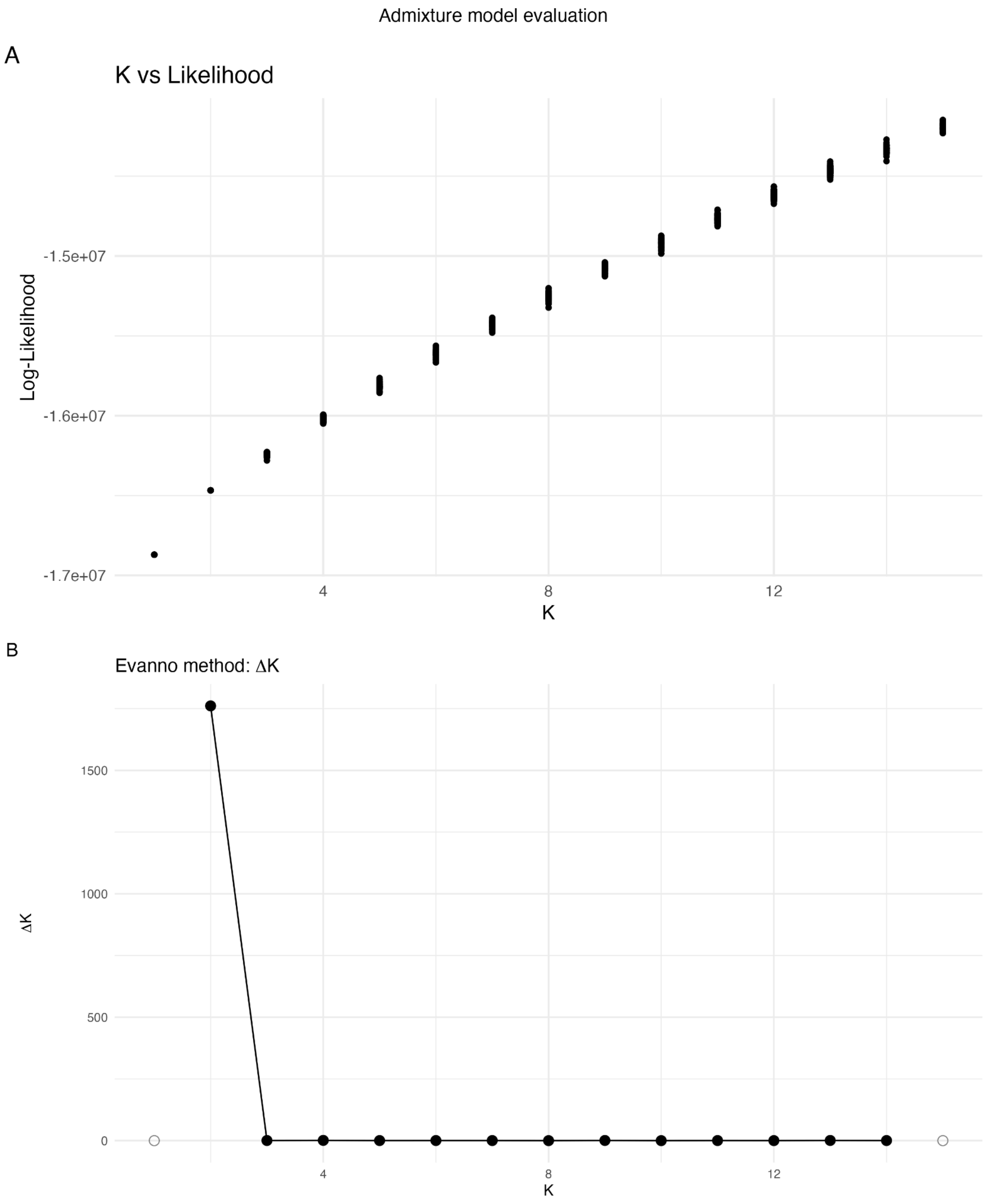

**Figure S22.** Evaluation of the ADMIXTURE results for *Ara ararauna* across values of K. (A) Mean log-likelihoods (± SD) from 50 replicate runs per K. (number of clusters evaluated). (B) ΔK values calculated using the Evanno method, which quantifies the second-order rate of change in log-likelihood. The peak at $K$ = 2 suggests this as the most likely number of genetic clusters. However, it is consistent with the known limitations of ΔK, which favors K = 2 and may underestimate the finer-scale population structure (Janes et al. 2017; Lawson, van Dorp, and Falush 2018). ΔK is undefined for $K$ = 1 and $K$ = 15

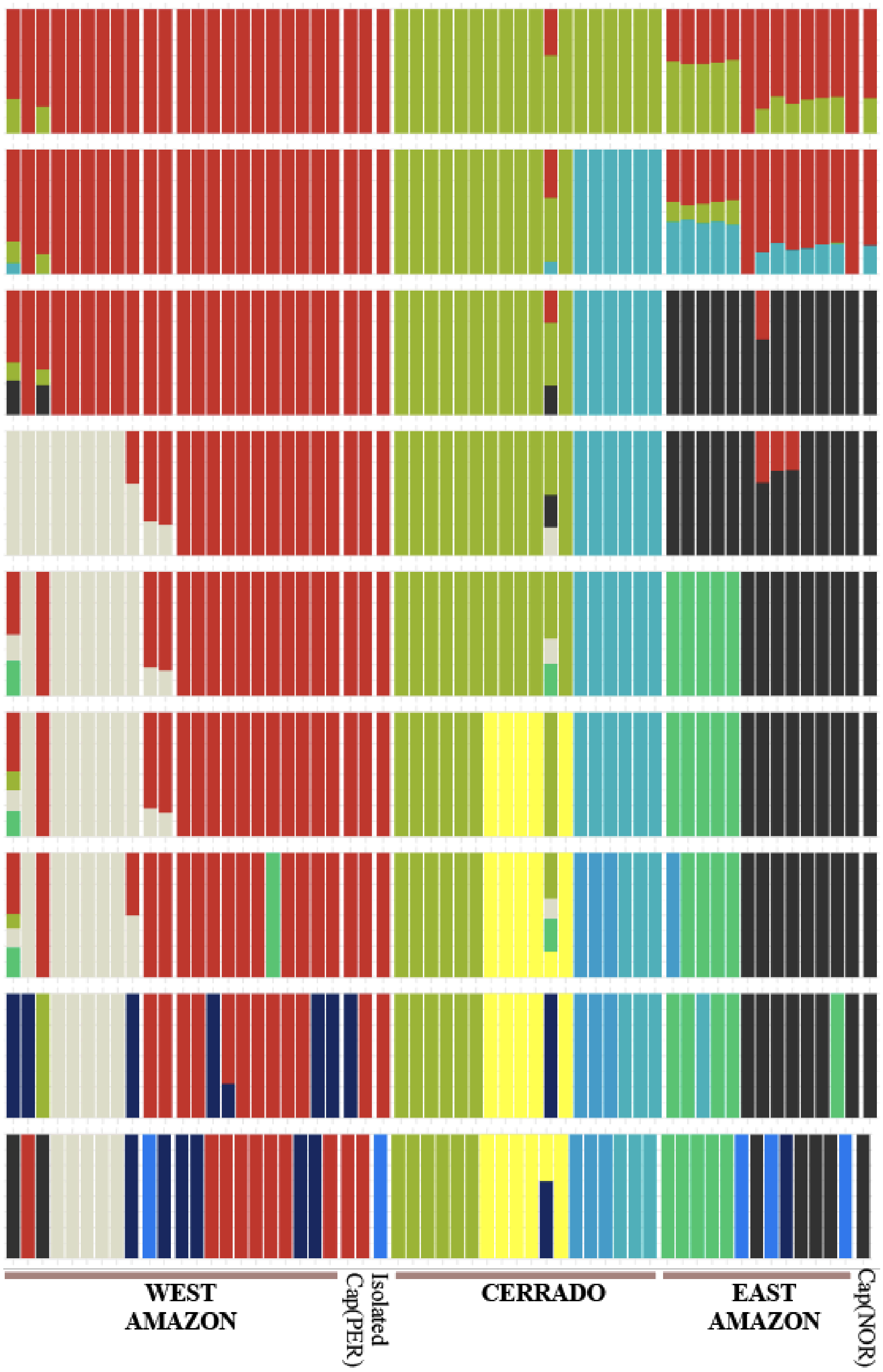

**Figure S23.** Admixture analysis of *Ara chloropterus* (K2 to K10). Bar plots are grouped according to the biome of origin (Amazon-West, Amazon-East, and Cerrado). The bar plots differentiate captive individuals (Peru = PER; Norway = NOR), and the individual representing the isolated populations.

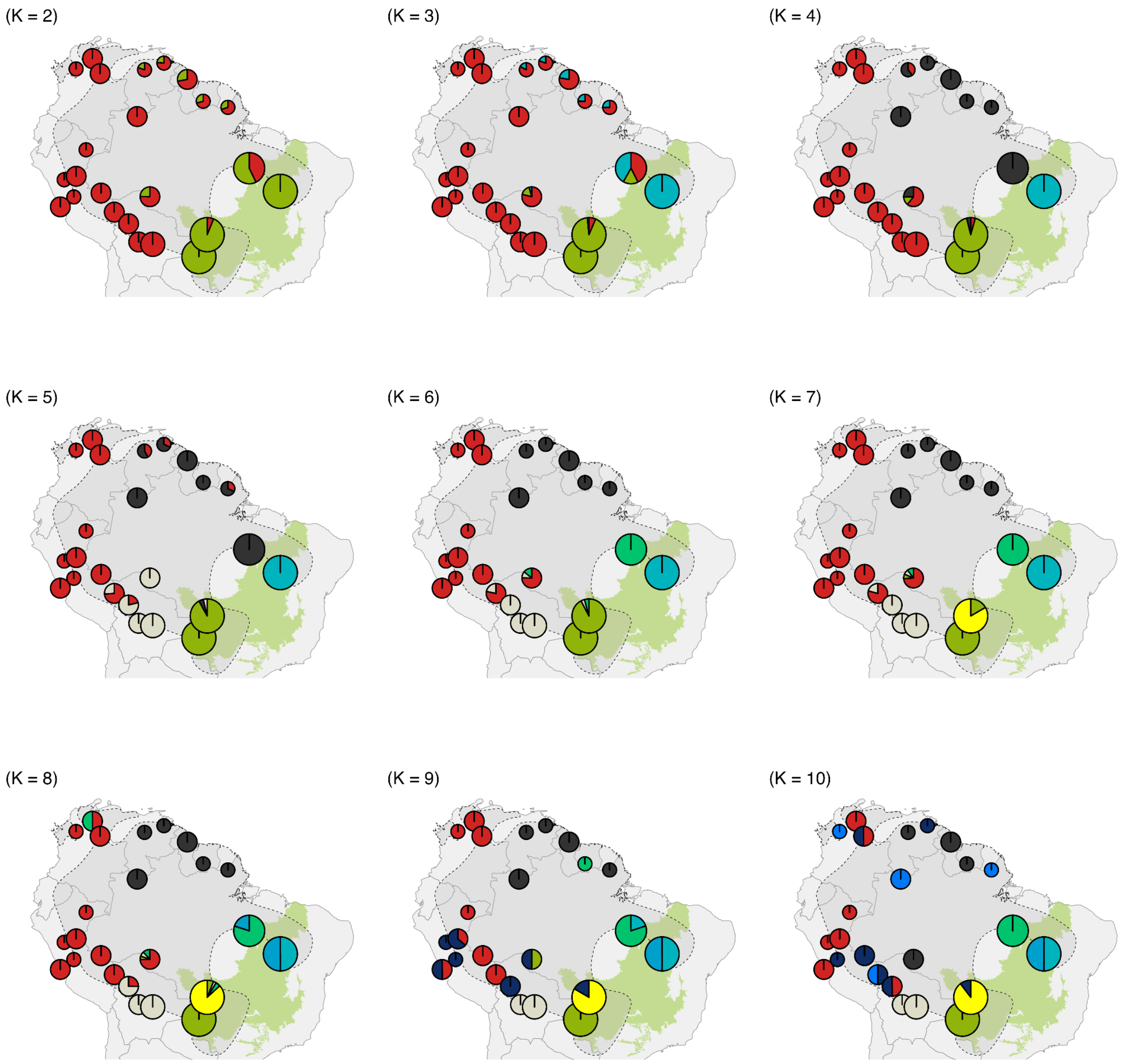

**Figure S24.** Admixture results for *Ara chloropterus* plotted by sampling location, with individuals clustered within 200 km of spatial proximity. Each pie chart represents ancestry proportions for a given value of K (2 to 10). The Cerrado biome is shown in light green, and the species' IUCN distribution is shaded in dark grey with dashed outlines. The basemap includes country boundaries from South America for reference.

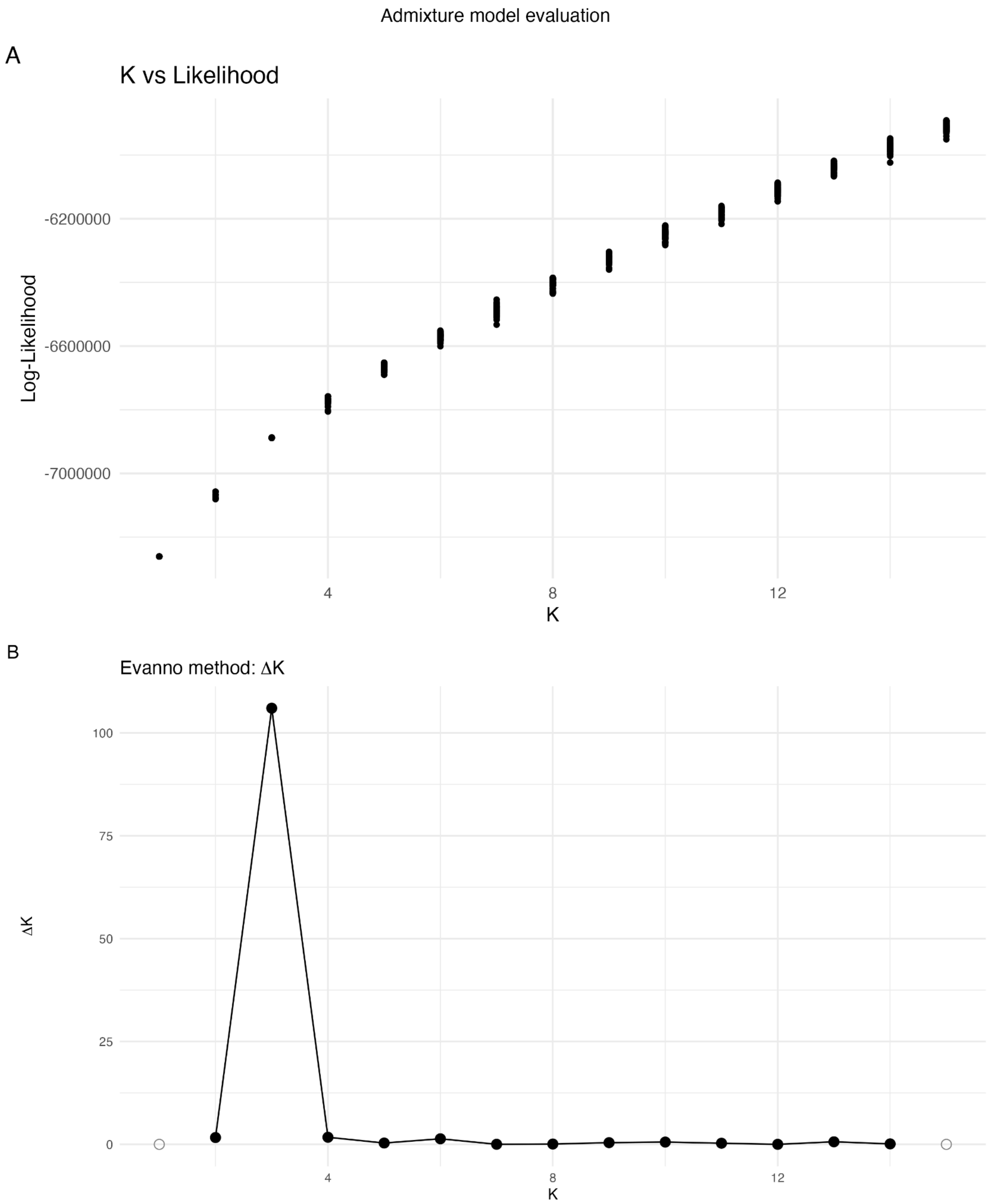

**Figure S25.** Evaluation of the ADMIXTURE results for *Ara chloropterus* across values of K. (A) Mean log-likelihoods (± SD) from 50 replicate runs per K. (number of clusters evaluated). (B) ΔK values calculated using the Evanno method, which quantifies the second-order rate of change in log-likelihood. The peak at $K$ = 3 suggests this as the most likely number of genetic clusters.

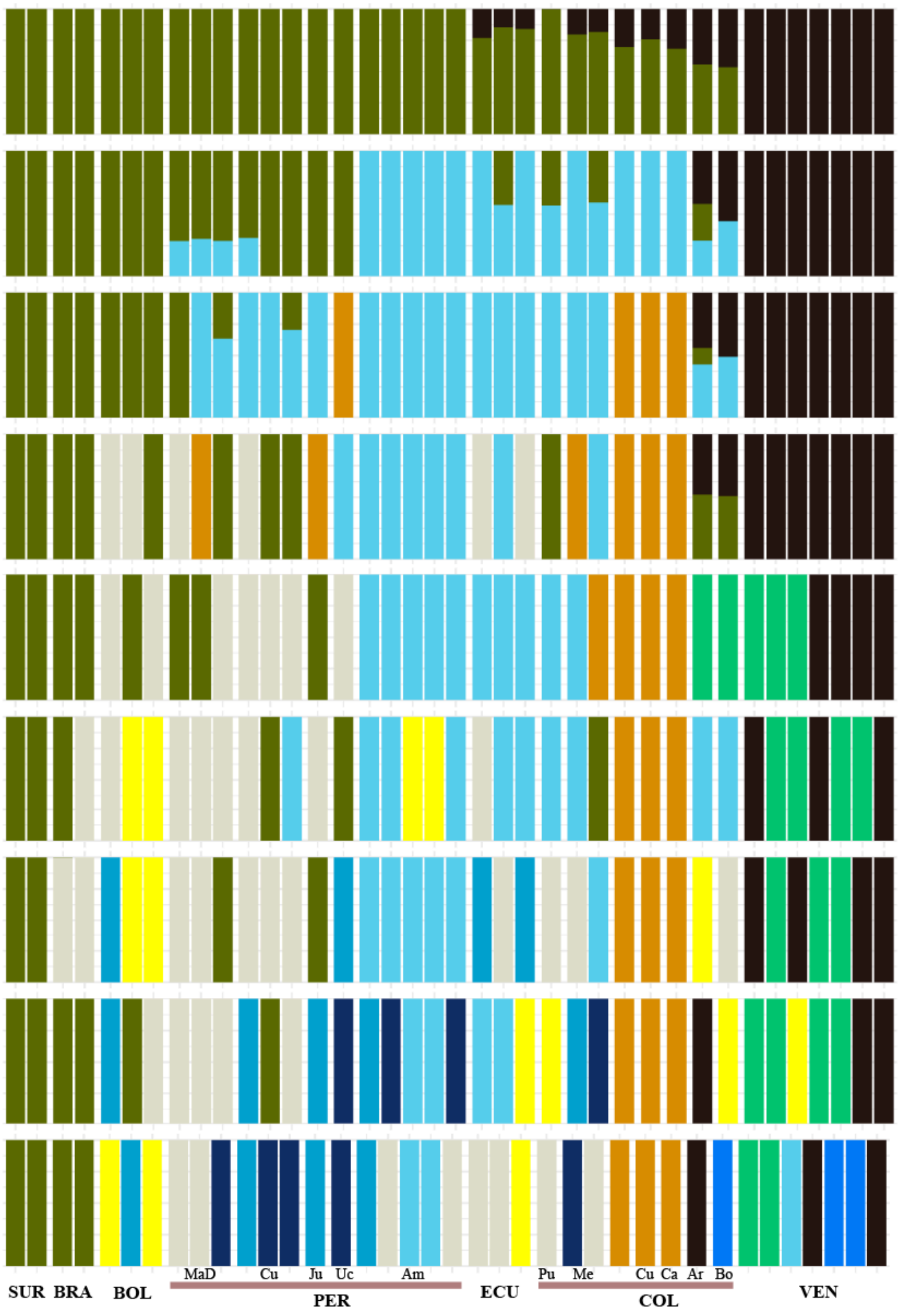

**Figure S26.** Admixture analysis of *Ara severus* (K2 to K10). Bar plots are grouped by country of origin. Certain provinces are denoted for Peru (MaD = Madre de Dios, Cu = Cuzco, Ju = Junín, Uc = Ucayali, Am = Amazonas) and Colombia (Pu = Putumayo, Me = Meta, Cu = Cundinamarca, Ca = Cauca, Ar = Arauca, Bo = Boyacá).

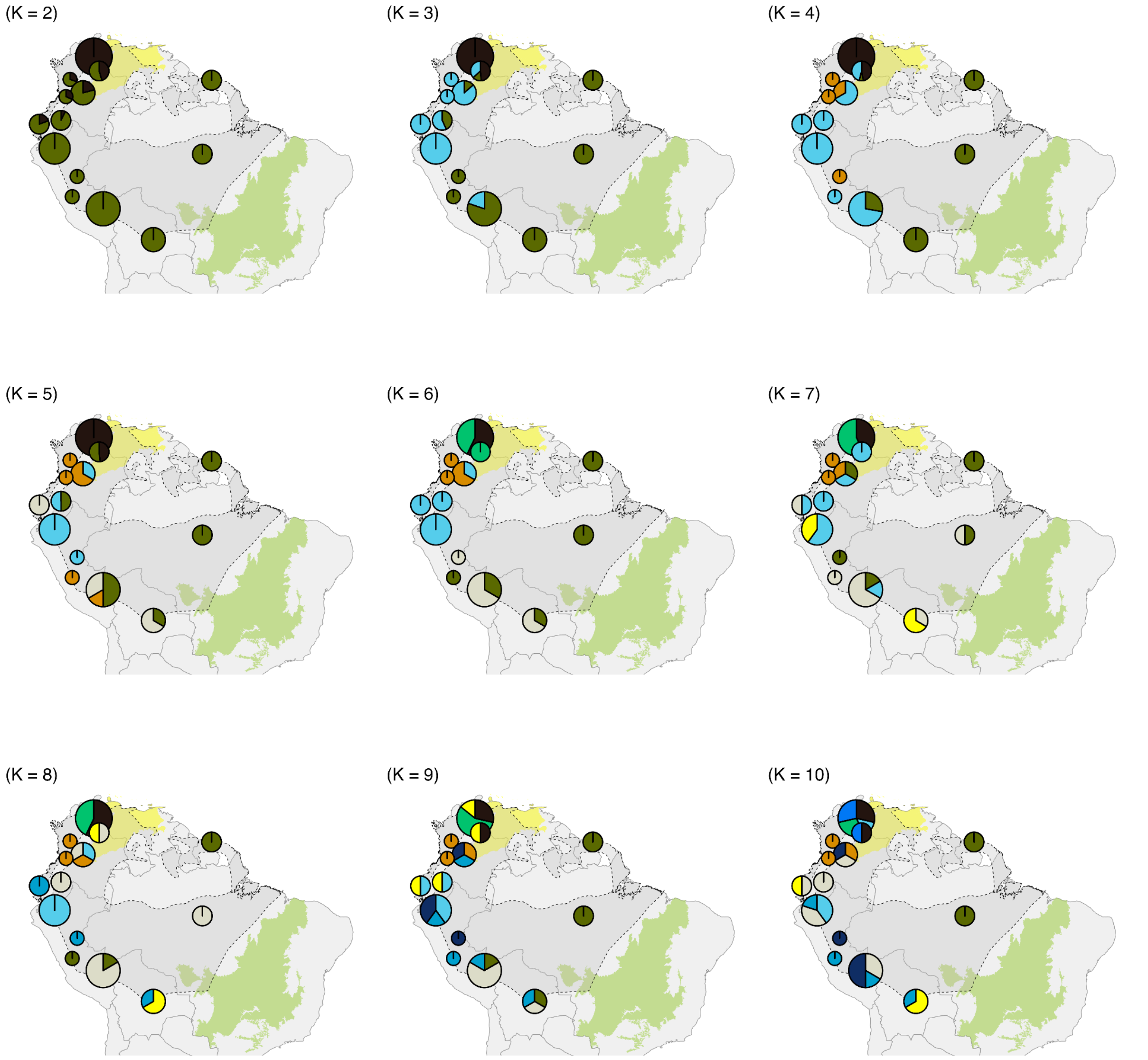

**Figure S27.** Admixture results for *Ara severus* plotted by sampling location, with individuals clustered within 200 km of spatial proximity. Each pie chart represents ancestry proportions for a given value of K (2 to 10). The Cerrado biome is shown in light green, and the Dry North of South America is in yellow. and the species' IUCN distribution is shaded in dark grey with dashed outlines. The basemap includes country boundaries from South America for reference.

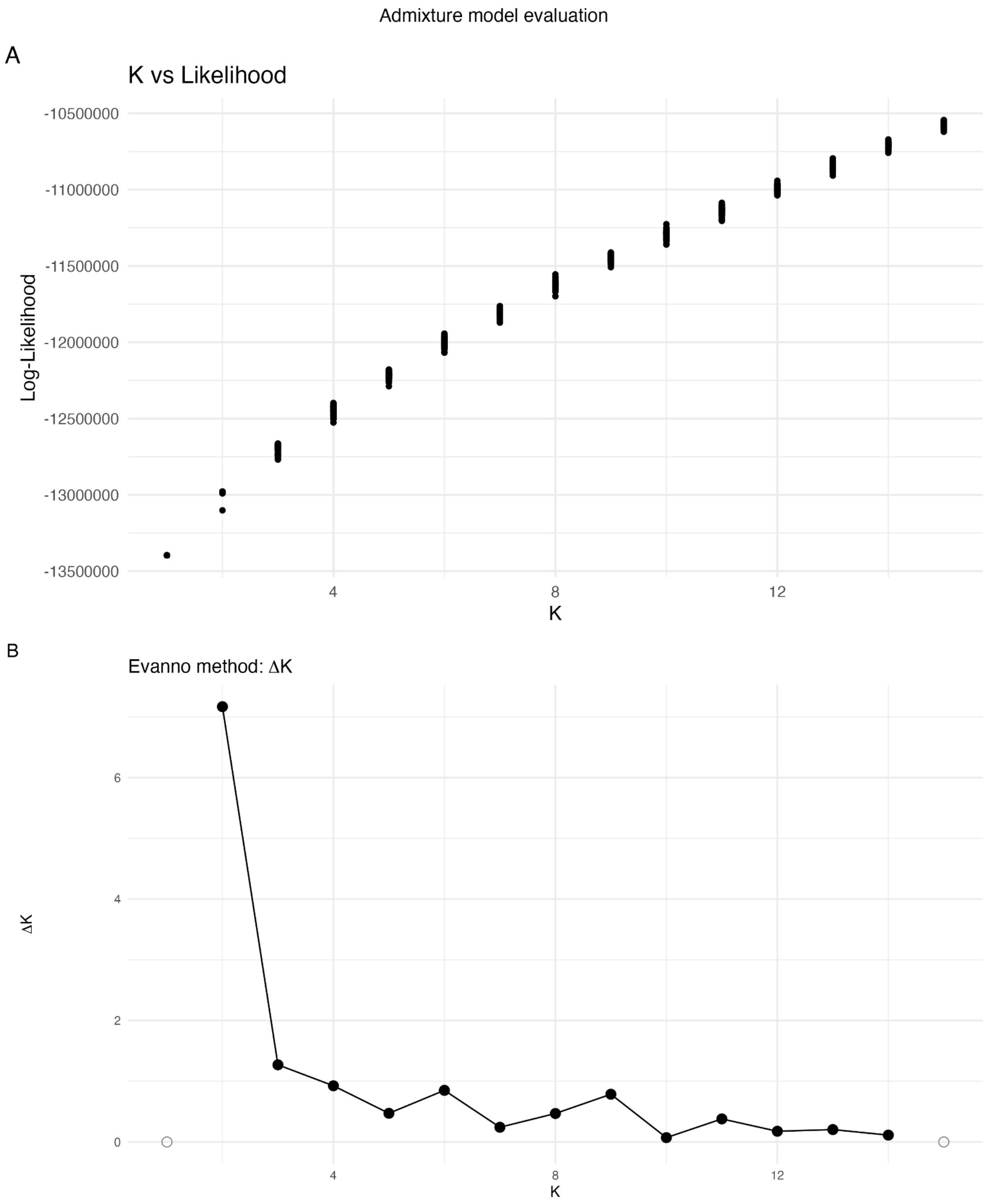

**Figure S28.** Evaluation of the ADMIXTURE results for *Ara severus* across values of K. (A) Mean log-likelihoods (± SD) from 50 replicate runs per K. (number of clusters evaluated). (B) ΔK values calculated using the Evanno method, which quantifies the second-order rate of change in log-likelihood. The peak at $K = 2$ suggests this as the most likely number of genetic clusters. However, it is consistent with the known limitations of ΔK, which favors K = 2 and may underestimate the finer-scale population structure (Janes et al. 2017; Lawson, van Dorp, and Falush 2018). ΔK is undefined for $K = 1$ and $K = 15$

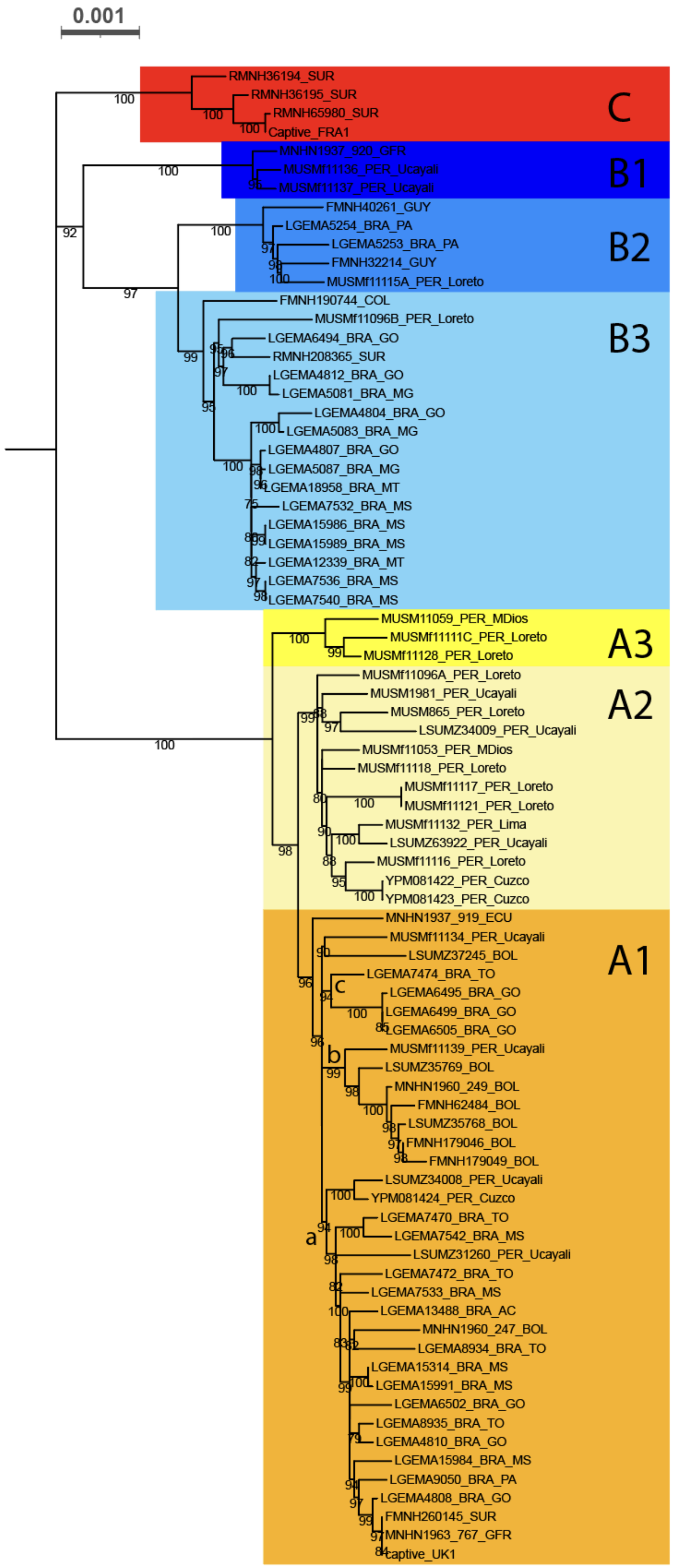
0.001
C
B1
B2
B3
A3
A2
A1
RMNH36194_SUR
RMNH36195_SUR
RMNH65980_SUR
Captive_FRA1
MNHN1937_920_GFR
MUSMf11136_PER_Ucayali
MUSMf11137_PER_Ucayali
FMNH40261_GUY
LGEMA5254_BRA_PA
LGEMA5253_BRA_PA
FMNH32214_GUY
MUSMf11115A_PER_Loreto
FMNH190744_COL
MUSMf11096B_PER_Loreto
LGEMA6494_BRA_GO
RMNH208365_SUR
LGEMA4812_BRA_GO
LGEMA5081_BRA_MG
LGEMA4804_BRA_GO
LGEMA5083_BRA_MG
LGEMA4807_BRA_GO
LGEMA5087_BRA_MG
LGEMA18958_BRA_MT
LGEMA7532_BRA_MS
LGEMA15986_BRA_MS
LGEMA15989_BRA_MS
LGEMA12339_BRA_MT
LGEMA7536_BRA_MS
LGEMA7540_BRA_MS
MUSM11059_PER_MDios
MUSMf11111C_PER_Loreto
MUSMf11128_PER_Loreto
MUSMf11096A_PER_Loreto
MUSM1981_PER_Ucayali
MUSM865_PER_Loreto
LSUMZ34009_PER_Ucayali
MUSMf11053_PER_MDios
MUSMf11118_PER_Loreto
MUSMf11117_PER_Loreto
MUSMf11121_PER_Loreto
MUSMf11132_PER_Lima
LSUMZ63922_PER_Ucayali
MUSMf11116_PER_Loreto
YPM081422_PER_Cuzco
YPM081423_PER_Cuzco
MNHN1937_919_ECU
MUSMf11134_PER_Ucayali
LSUMZ37245_BOL
LGEMA7474_BRA_TO
LGEMA6495_BRA_GO
LGEMA6499_BRA_GO
LGEMA6505_BRA_GO
MUSMf11139_PER_Ucayali
LSUMZ35769_BOL
MNHN1960_249_BOL
FMNH62484_BOL
LSUMZ35768_BOL
FMNH179046_BOL
FMNH179049_BOL
LSUMZ34008_PER_Ucayali
YPM081424_PER_Cuzco
LGEMA7470_BRA_TO
LGEMA7542_BRA_MS
LSUMZ31260_PER_Ucayali
LGEMA7472_BRA_TO
LGEMA7533_BRA_MS
LGEMA13488_BRA_AC
MNHN1960_247_BOL
LGEMA8934_BRA_TO
LGEMA15314_BRA_MS
LGEMA15991_BRA_MS
LGEMA6502_BRA_GO
LGEMA8935_BRA_TO
LGEMA4810_BRA_GO
LGEMA15984_BRA_MS
LGEMA9050_BRA_PA
LGEMA4808_BRA_GO
FMNH260145_SUR
MNHN1963_767_GFR
captive_UK1
a
b
c

**Figure S29.** Maximum-likelihood phylogeny of *Ara ararauna* mitogenomes, inferred using IQ-TREE 2 (Minh et al. 2020). Bootstrap support values greater than 75 are displayed at the corresponding nodes. The outgroup is not shown. Major monophyletic clades (A, B, and C), and subclades (where applicable) have been distinctly highlighted. The isolated population is represented by individual FMNH190744 (Clade B3)

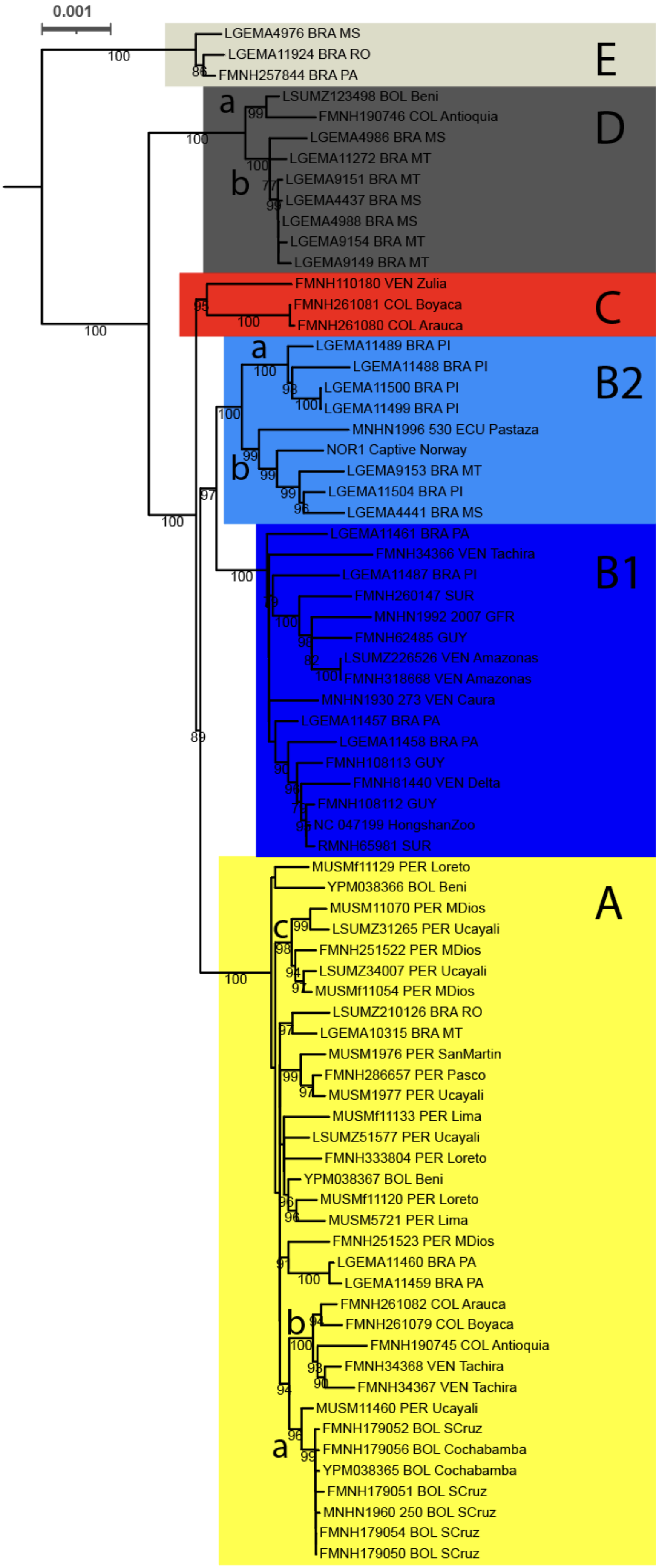
0.001
LGEMA4976 BRA MS
LGEMA11924 BRA RO
FMNH257844 BRA PA
E
LSUMZ123498 BOL Beni
FMNH190746 COL Antioquia
LGEMA4986 BRA MS
LGEMA11272 BRA MT
LGEMA9151 BRA MT
LGEMA4437 BRA MS
LGEMA4988 BRA MS
LGEMA9154 BRA MT
LGEMA9149 BRA MT
D
FMNH110180 VEN Zulia
FMNH261081 COL Boyaca
FMNH261080 COL Arauca
C
LGEMA11489 BRA PI
LGEMA11488 BRA PI
LGEMA11500 BRA PI
LGEMA11499 BRA PI
MNHN1996 530 ECU Pastaza
NOR1 Captive Norway
LGEMA9153 BRA MT
LGEMA11504 BRA PI
LGEMA4441 BRA MS
B2
LGEMA11461 BRA PA
FMNH34366 VEN Tachira
LGEMA11487 BRA PI
FMNH260147 SUR
MNHN1992 2007 GFR
FMNH62485 GUY
LSUMZ226526 VEN Amazonas
FMNH318668 VEN Amazonas
MNHN1930 273 VEN Caura
LGEMA11457 BRA PA
LGEMA11458 BRA PA
FMNH108113 GUY
FMNH81440 VEN Delta
FMNH108112 GUY
NC 047199 HongshanZoo
RMNH65981 SUR
B1
MUSMf11129 PER Loreto
YPM038366 BOL Beni
MUSM11070 PER MDios
LSUMZ31265 PER Ucayali
FMNH251522 PER MDios
LSUMZ34007 PER Ucayali
MUSMf11054 PER MDios
LSUMZ210126 BRA RO
LGEMA10315 BRA MT
MUSM1976 PER SanMartin
FMNH286657 PER Pasco
MUSM1977 PER Ucayali
MUSMf11133 PER Lima
LSUMZ51577 PER Ucayali
FMNH333804 PER Loreto
YPM038367 BOL Beni
MUSMf11120 PER Loreto
MUSM5721 PER Lima
FMNH251523 PER MDios
LGEMA11460 BRA PA
LGEMA11459 BRA PA
FMNH261082 COL Arauca
FMNH261079 COL Boyaca
FMNH190745 COL Antioquia
FMNH34368 VEN Tachira
FMNH34367 VEN Tachira
MUSM11460 PER Ucayali
FMNH179052 BOL SCruz
FMNH179056 BOL Cochabamba
YPM038365 BOL Cochabamba
FMNH179051 BOL SCruz
MNHN1960 250 BOL SCruz
FMNH179054 BOL SCruz
FMNH179050 BOL SCruz
A

**Figure S30.** Maximum-likelihood phylogeny of *Ara chloropterus* mitogenomes, inferred using IQ-TREE 2 (Minh et al. 2020). Bootstrap support values greater than 75 are displayed at the corresponding nodes. The outgroup is not shown. Major monophyletic clades (A, B, C, D, and E), and subclades (where applicable) have been distinctly highlighted. The isolated population is represented by individual FMNH190745 (Clade A1b)

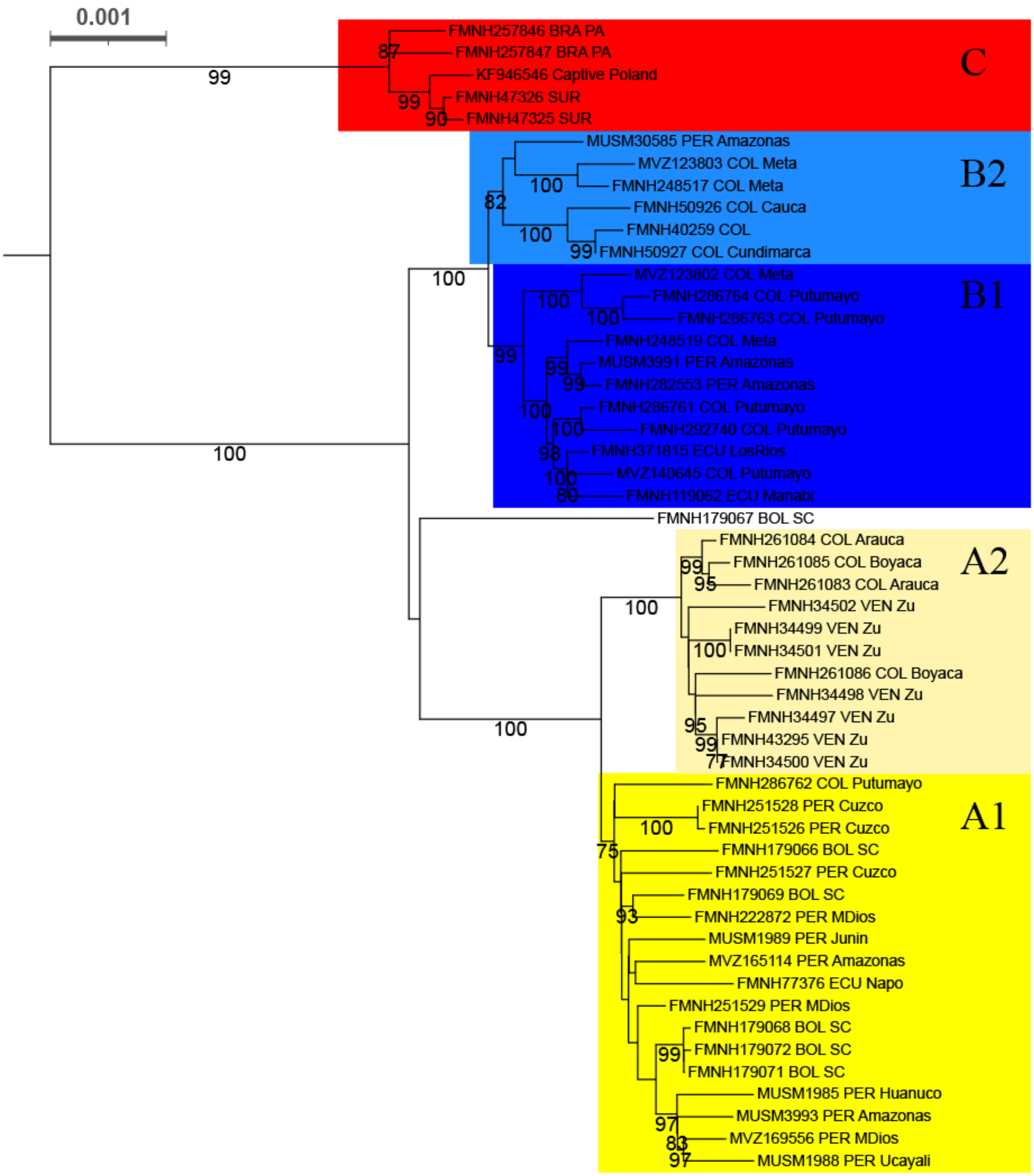

**Figure S31.** Maximum-likelihood phylogeny of *Ara severus* mitogenomes, inferred using IQ-TREE 2 (Minh et al. 2020). Bootstrap support values greater than 75 are displayed at the corresponding nodes. The outgroup is not shown. Major monophyletic clades (A, B, and C), and subclades (where applicable) have been distinctly highlighted.

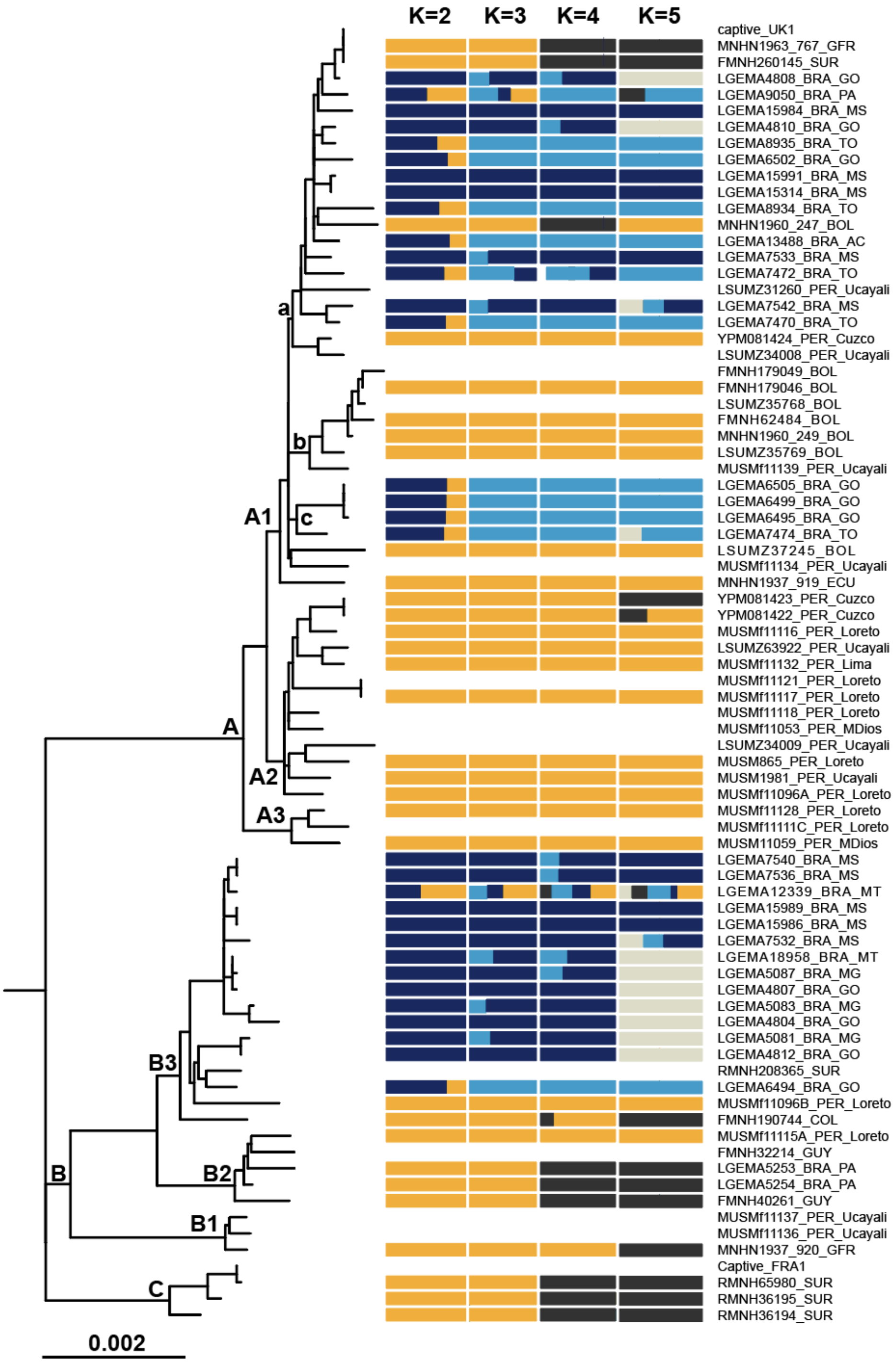

**Figure S32.** Maximum-likelihood phylogeny of *Ara ararauna* mitogenomes, inferred using IQ-TREE 2 (Minh et al. 2020). Outgroup is not shown. Major monophyletic clades, and subclades (where applicable), are labeled at the beginning of the clade. Individual admixture proportions (K = 2 to 5), calculated using NGSadmix (Skotte, Korneliussen, and Albrechtsen 2013) based on whole-genome data, are shown as bar plots adjacent to each corresponding sample.

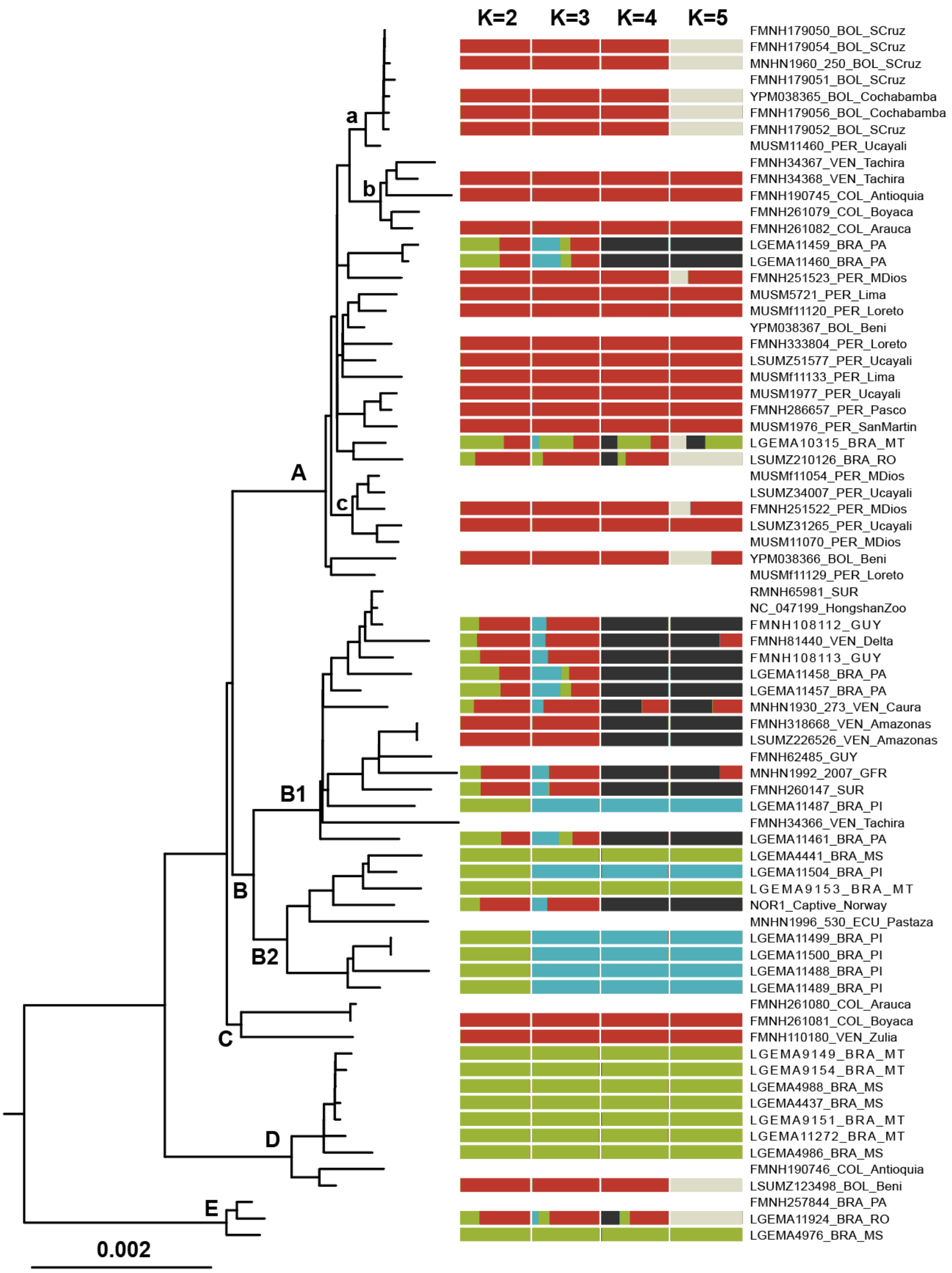

**Figure S33.** Maximum-likelihood phylogeny of *Ara chloropterus* mitogenomes, inferred using IQ-TREE 2 (Minh et al. 2020). Outgroup is not shown. Major monophyletic clades, and subclades (where applicable), are labeled at the beginning of the clade. Individual admixture proportions (K = 2 to 5), calculated using NGSadmix (Skotte, Korneliussen, and Albrechtsen 2013) based on whole-genome data, are shown as bar plots adjacent to each corresponding sample.

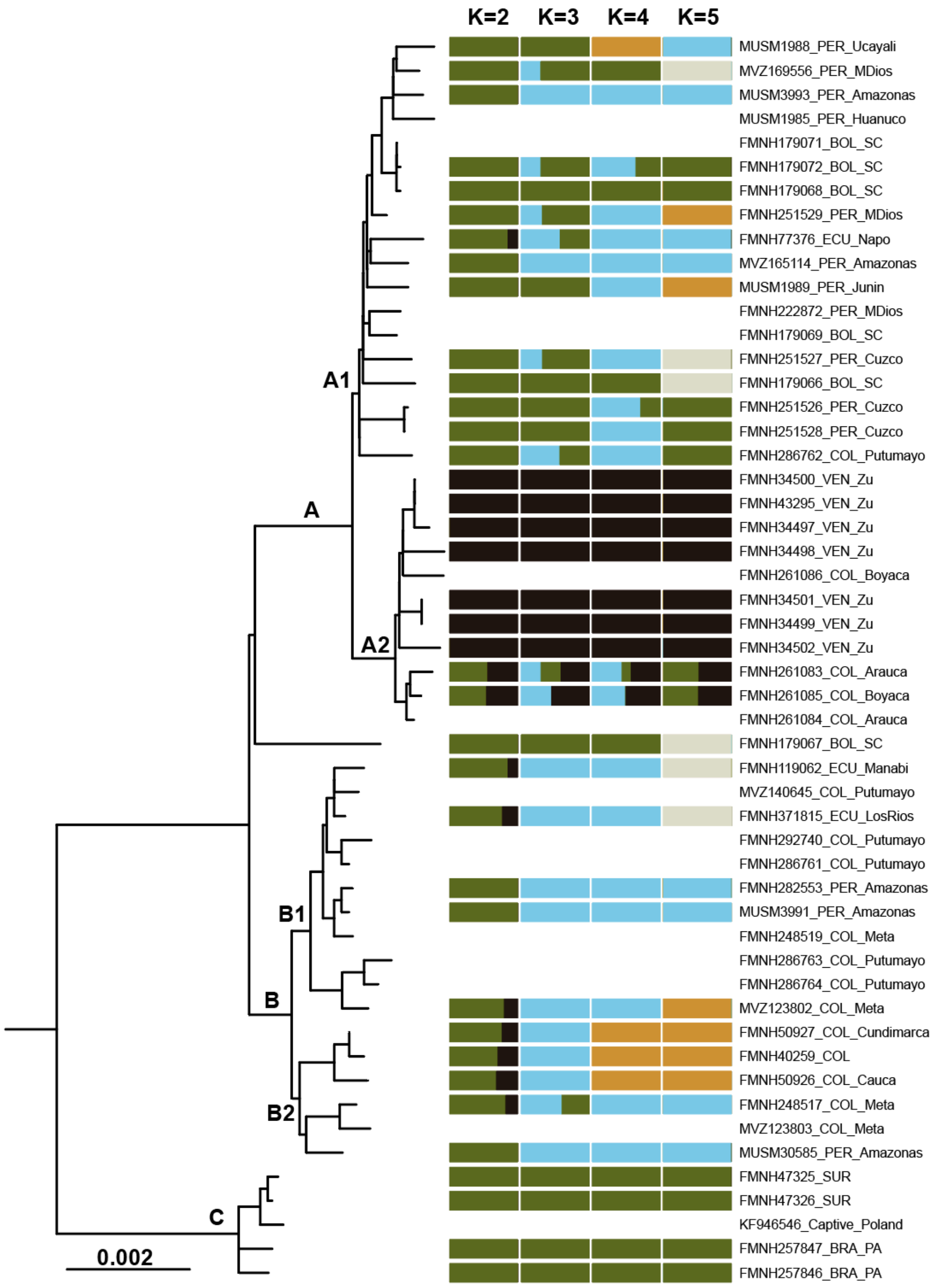

**Figure S34.** Maximum-likelihood phylogeny of *Ara severus* mitogenomes, inferred using IQ-TREE 2 (Minh et al. 2020). Outgroup is not shown. Major monophyletic clades, and subclades (where applicable), are labeled at the beginning of the clade. Individual admixture proportions (K = 2 to 5), calculated using NGSadmix (Skotte, Korneliussen, and Albrechtsen 2013) based on whole-genome data, are shown as bar plots adjacent to each corresponding sample.

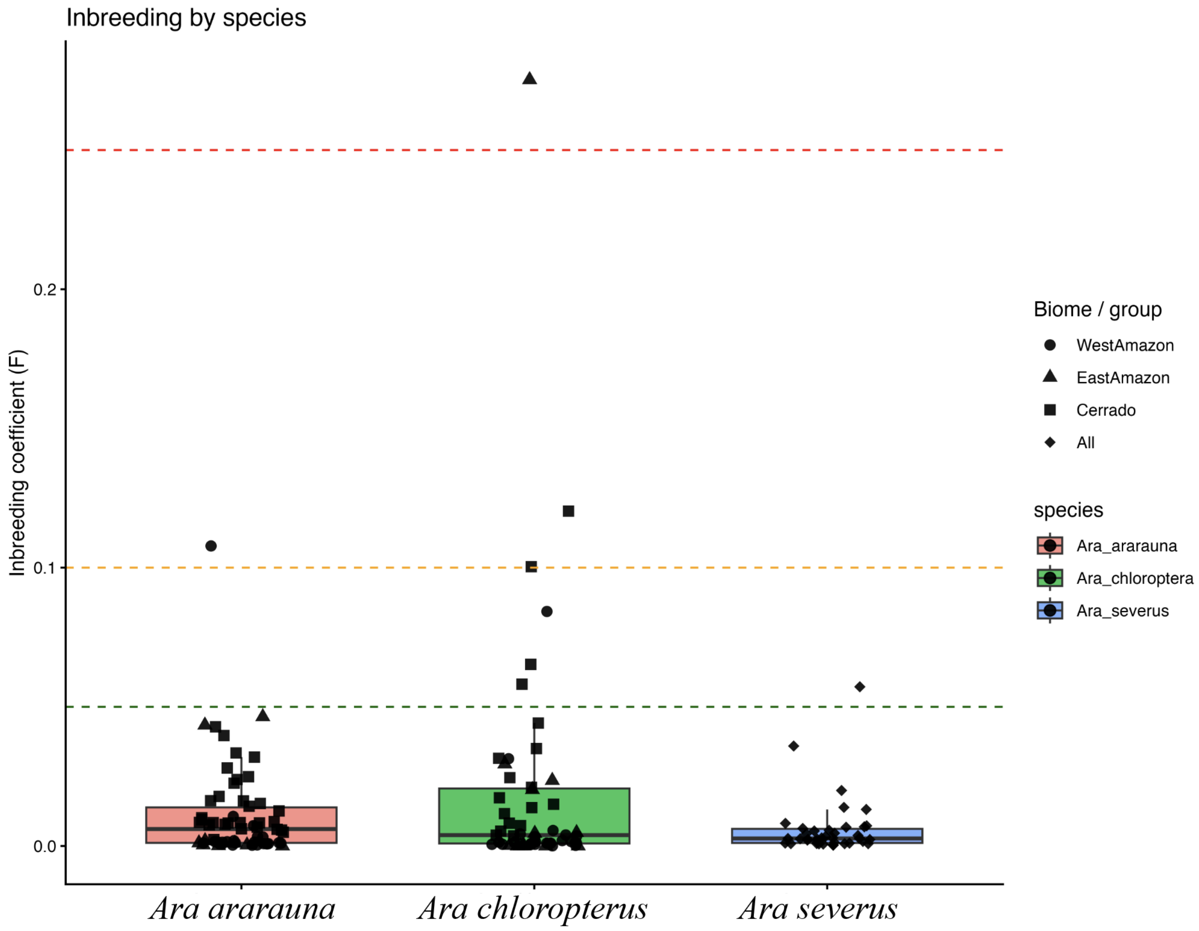

**Figure S35**. Distribution of individual inbreeding coefficients (F) across three *Ara* species. Points represent individual estimates, with symbol shapes denoting populations (West Amazon, East Amazon, Cerrado, and all individuals for *A. severus*).

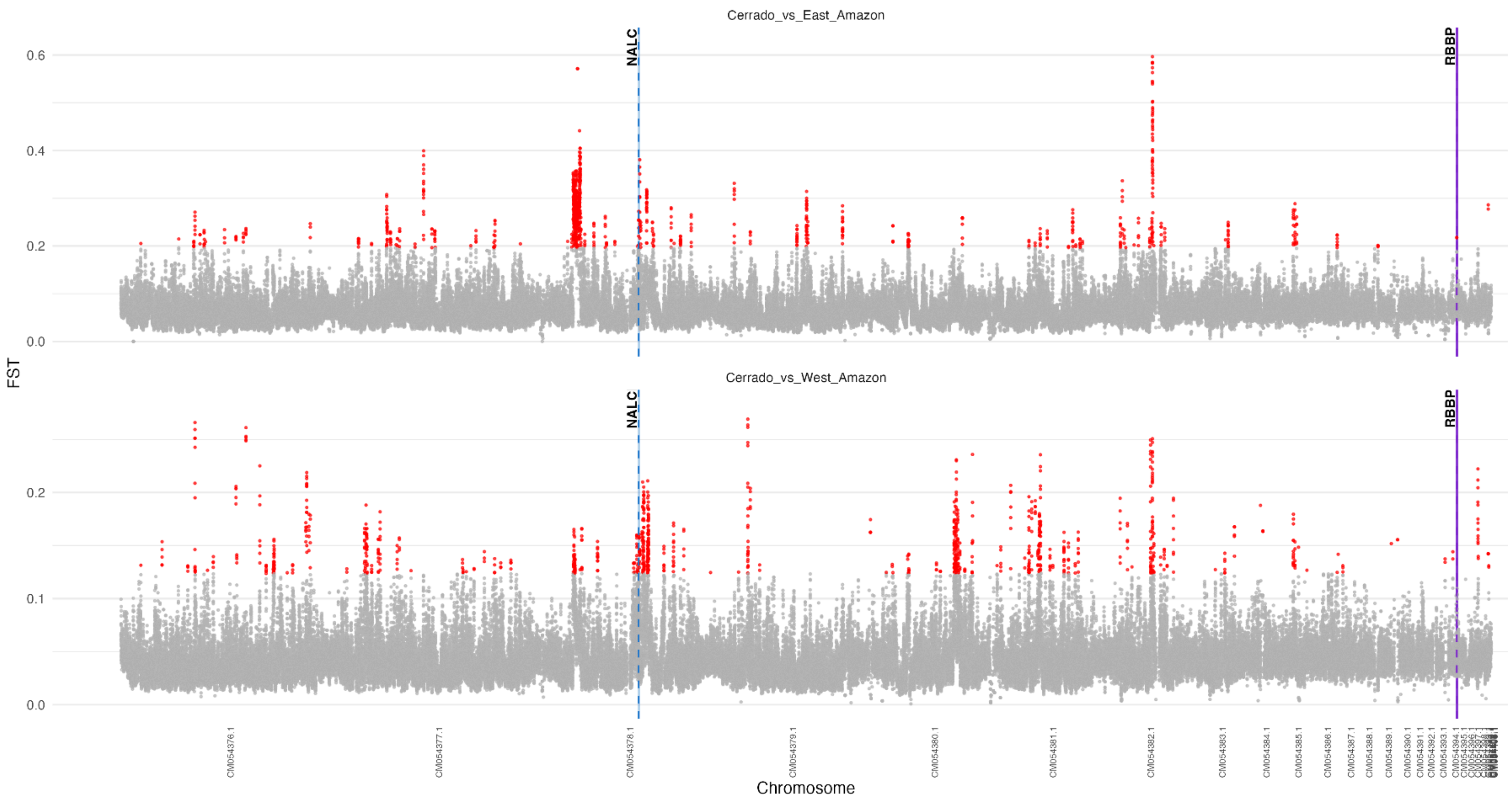

**Figure S36**. Genome-wide distribution of window-based (windows = 50 kb, step = 10 kb) $F_{ST}$ values in *Ara ararauna* for two pairwise comparisons: East Amazon vs. Cerrado (top), and West Amazon vs. Cerrado (down). Each point represents the midpoint of the window, the x axis represents the chromosomes. Windows with fewer than 100 sites and those located on the Z chromosome were excluded. For each comparison, the top 1% of $F_{ST}$ values (empirical 0.99 quantile) are highlighted in red and defined as outlier windows, while remaining windows are shown in grey. The two candidate genes (in $F_{ST}$ outlier windows and with low Fay & Wu's H) cross species are highlighted and labeled.

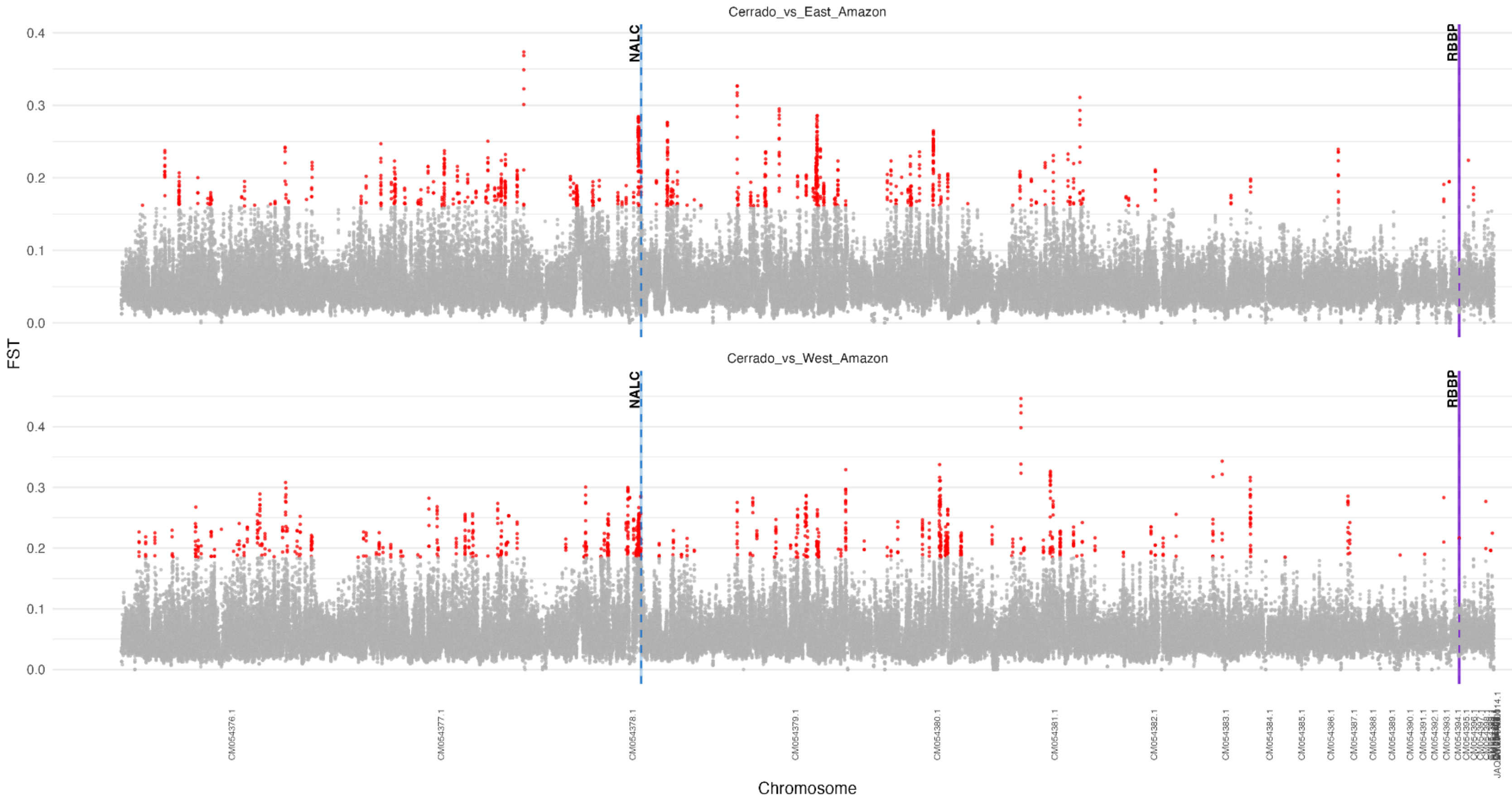

**Figure S37**. Genome-wide distribution of window-based (windows = 50 kb, step = 10 kb) $F_{ST}$ values in *Ara chloropterus* for two pairwise comparisons: East Amazon vs. Cerrado (top), and West Amazon vs. Cerrado (down). Each point represents the midpoint of the window, the x axis represents the chromosomes. Windows with fewer than 100 sites and those located on the Z chromosome were excluded. For each comparison, the top 1% of $F_{ST}$ values (empirical 0.99 quantile) are highlighted in red and defined as outlier windows, while remaining windows are shown in grey. The two candidate genes (in $F_{ST}$ outlier windows and with low Fay & Wu's H) cross species are highlighted and labeled.

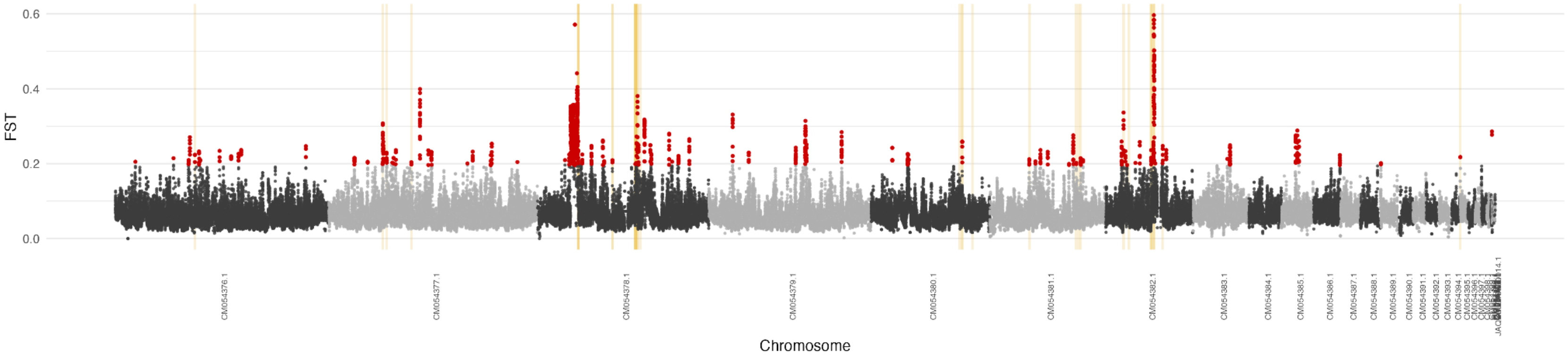

ararauna | Cerrado | Cerrado_vs_West_Amazon

FST Manhattan plot; red = FST outliers; gold bands = DOUBLE genes

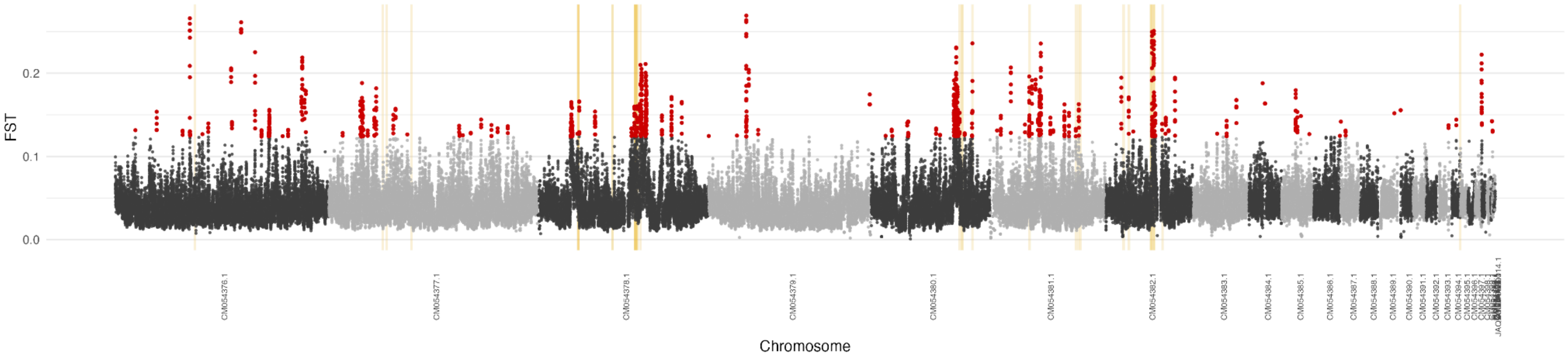

ararauna | Cerrado | Fay & Wu's H

Red = H-low windows; gold bands = DOUBLE genes

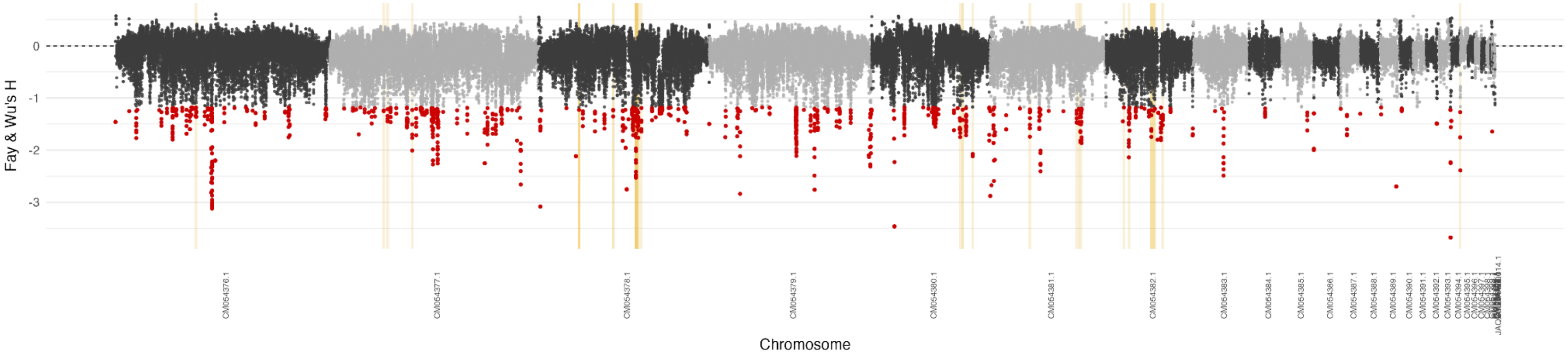

**Figure S38**. Genome-wide distribution of window-based (windows = 50 kb, step = 10 kb) $F_{ST}$ values in *Ara ararauna* for two pairwise comparisons, East Amazon vs. Cerrado (top), and West Amazon vs. Cerrado (middle), and Fay & Wu's Manhattan plot (down). Each point represents the midpoint of the window, the x axis represents the chromosomes. Windows with fewer than 100 sites and those located on the Z chromosome were excluded. For each comparison, the top 1% of $F_{ST}$ values (empirical 0.99 quantile) are highlighted in red and defined as outlier windows, while remaining windows are shown in grey. All candidate genes (in $F_{ST}$ outlier windows and with low Fay & Wu's H) are highlighted; a full list of genes can be found in Table S15.

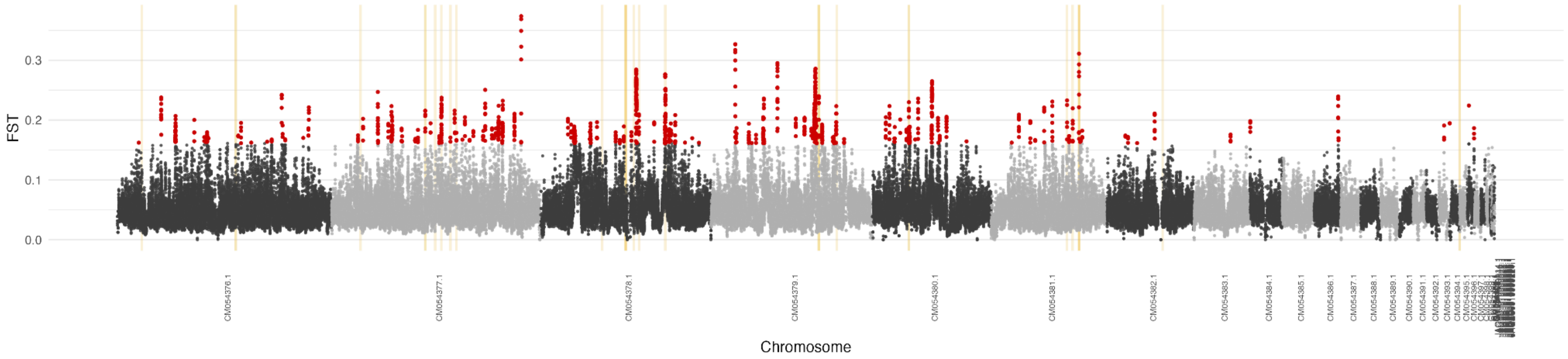

chloropterus | Cerrado | Cerrado_vs_West_Amazon

FST Manhattan plot; red = FST outliers; gold bands = DOUBLE genes

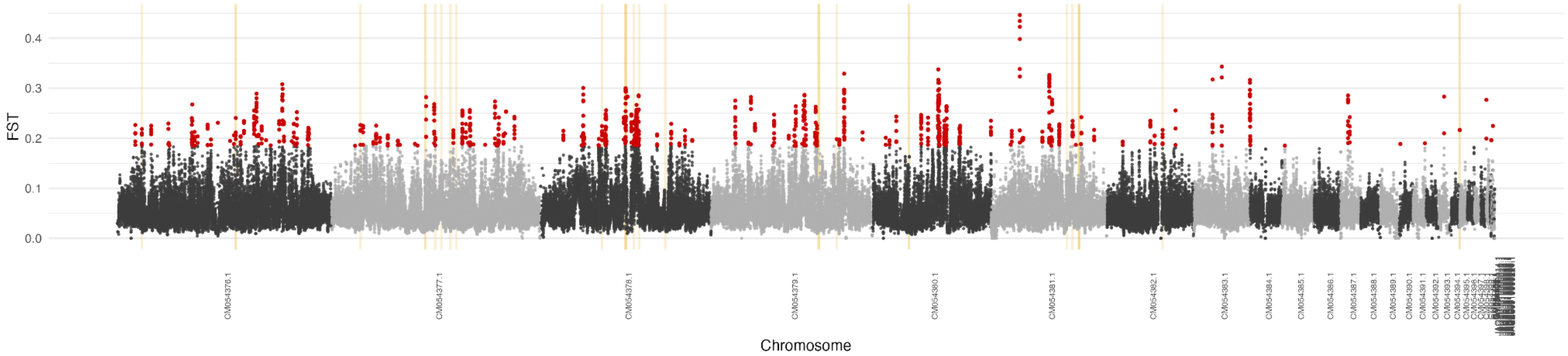

chloropterus | Cerrado | Fay & Wu's H

Red = H-low windows; gold bands = DOUBLE genes

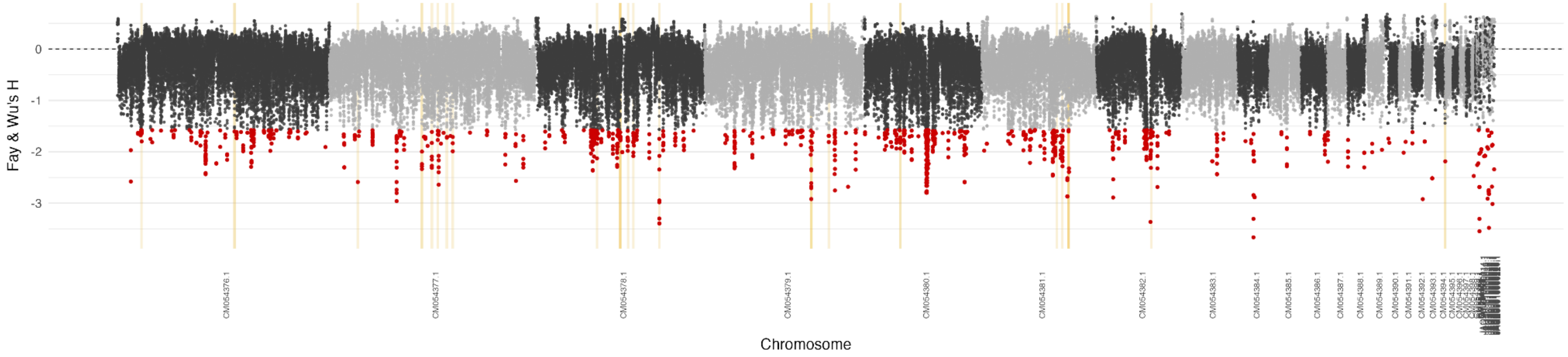

**Figure S39**. Genome-wide distribution of window-based (windows = 50 kb, step = 10 kb) $F_{ST}$ values in *Ara chloropterus* for two pairwise comparisons, East Amazon vs. Cerrado (top), and West Amazon vs. Cerrado (middle), and Fay & Wu's Manhattan plot (down). Each point represents the midpoint of the window, the x axis represents the chromosomes. Windows with fewer than 100 sites and those located on the Z chromosome were excluded. For each comparison, the top 1% of $F_{ST}$ values (empirical 0.99 quantile) are highlighted in red and defined as outlier windows, while remaining windows are shown in grey. All candidate genes (in $F_{ST}$ outlier windows and with low Fay & Wu's H) are highlighted; a full list of genes can be found in Table S15.

ararauna | Genome-wide Manhattan plots for Fay & Wu's H and Tajima's D

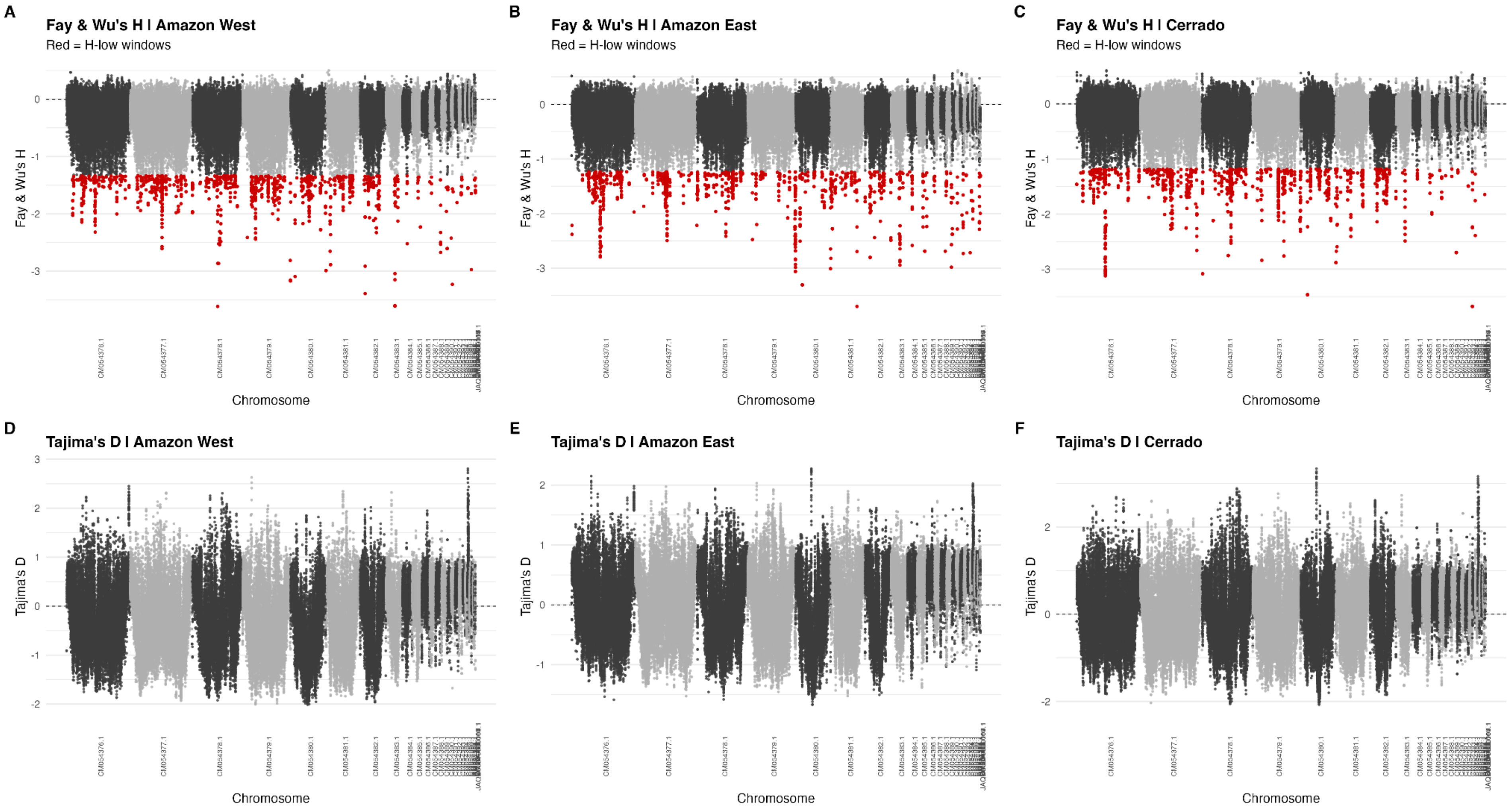

**Figure S40**. Genome-wide Manhattan plots for Fay & Wu's H and Tajima's D in *Ara ararauna*. Data was visualized independently for Amazon West (A and D), Amazon East (B and E), and Cerrado (C and F) populations. Low-H windows, defined separately for each species and biome as windows in the lower 1% of the empirical Fay and Wu's H distribution, are shown in red.

**chloropterus | Genome-wide Manhattan plots for Fay & Wu's H and Tajima's D**

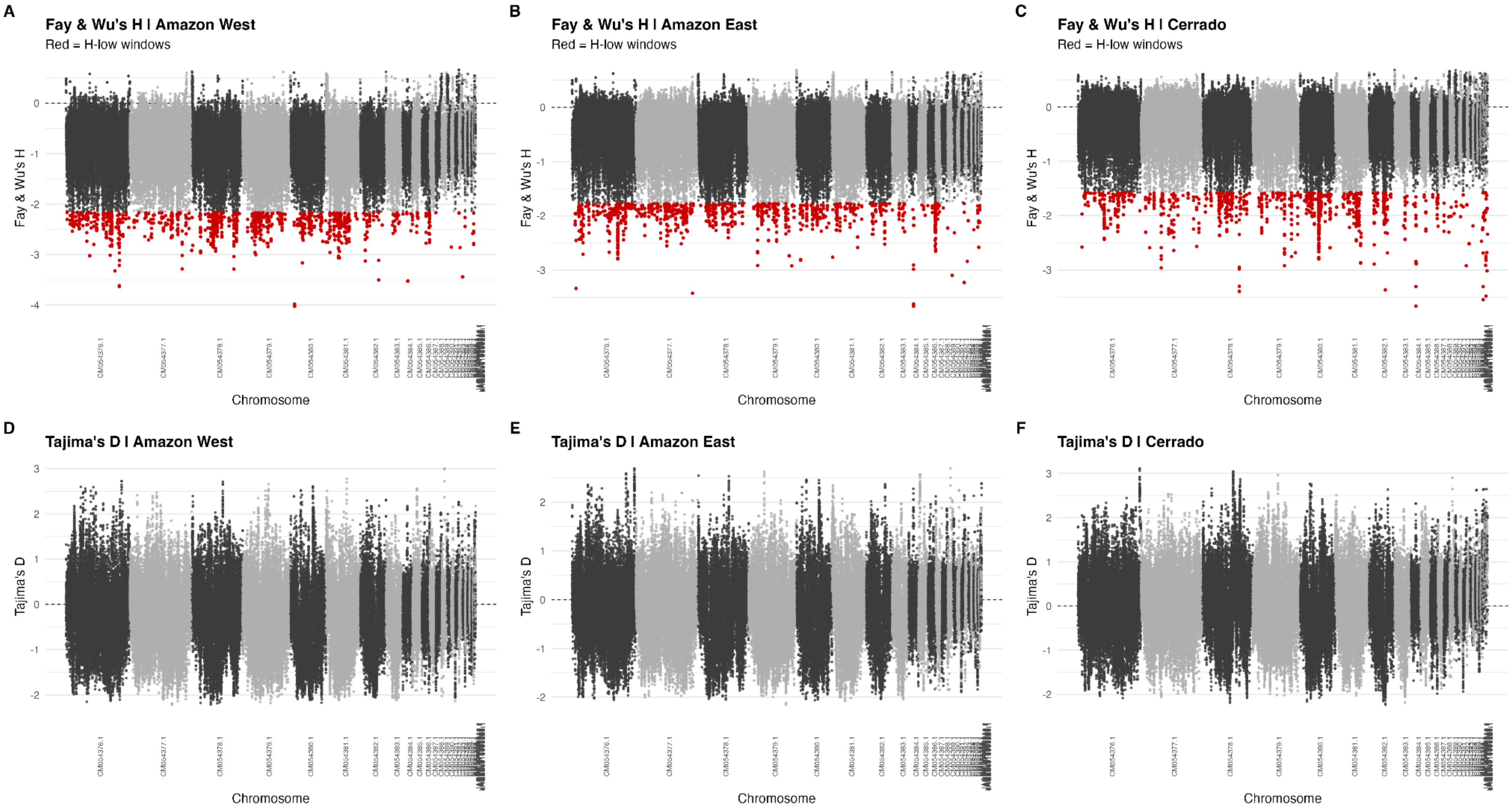

**Figure S41**. Genome-wide Manhattan plots for Fay & Wu's H and Tajima's D in *Ara chloropterus*. Data was visualized independently for Amazon West (A and D), Amazon East (B and E), and Cerrado (C and F) populations. Low-H windows, defined separately for each species and biome as windows in the lower 1% of the empirical Fay and Wu's H distribution, are shown in red.

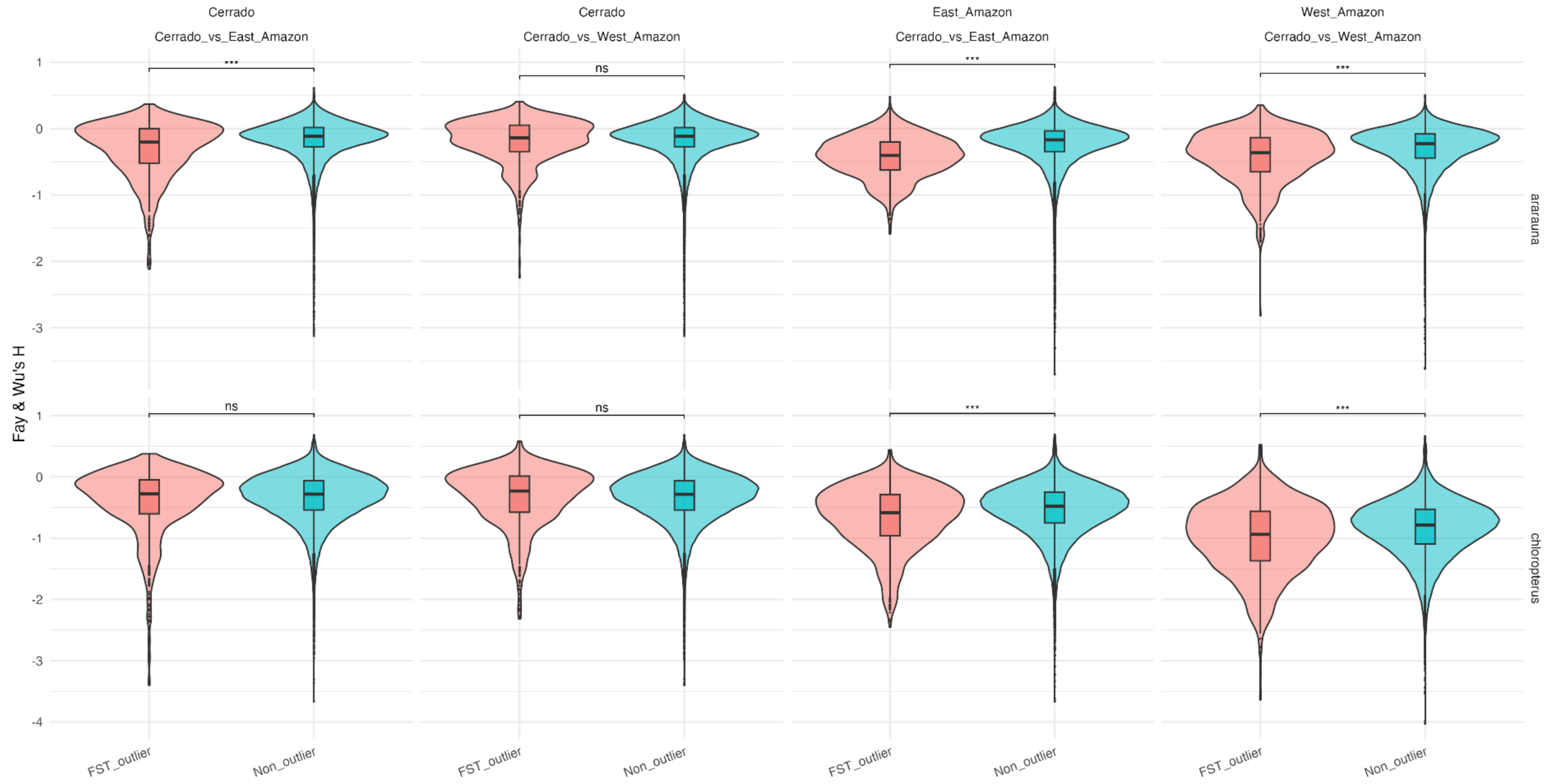

**Figure S42**. Fay & Wu's H in $F_{ST}$ outlier and non-outlier windows. $F_{ST}$ outliers were defined as the top 1% of the empirical $F_{ST}$ distribution within each species and population comparison. For each comparison, Fay & Wu's H was compared between $F_{ST}$ outlier windows and non-outlier windows using a one-sided Wilcoxon rank-sum test. Asterisks indicate Benjamini–Hochberg-adjusted significance levels; ns = not significant.

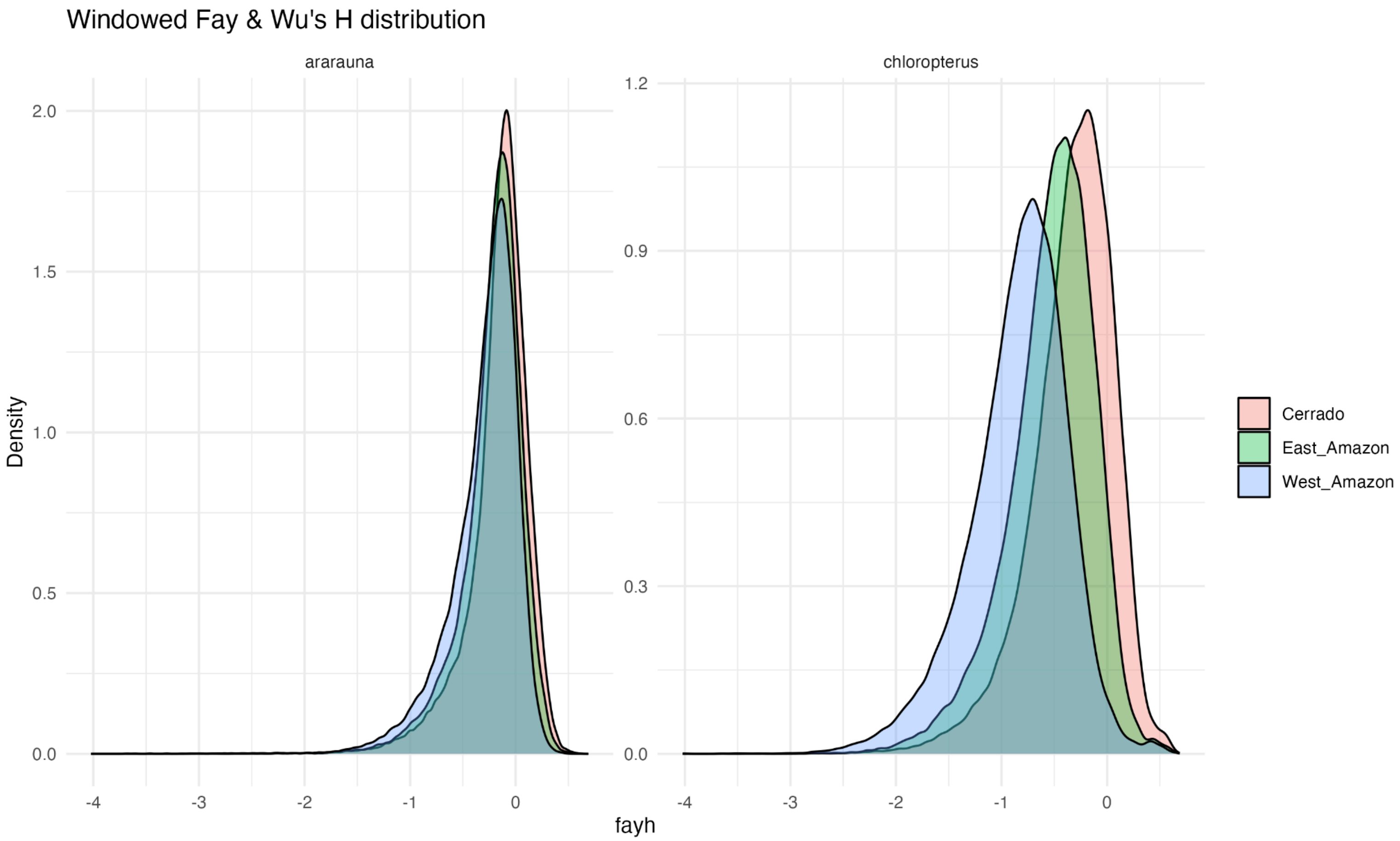

**Figure S43**. Distribution of Fay & Wu's *H* across genomic windows in *Ara ararauna* and *Ara chloropterus* per population.

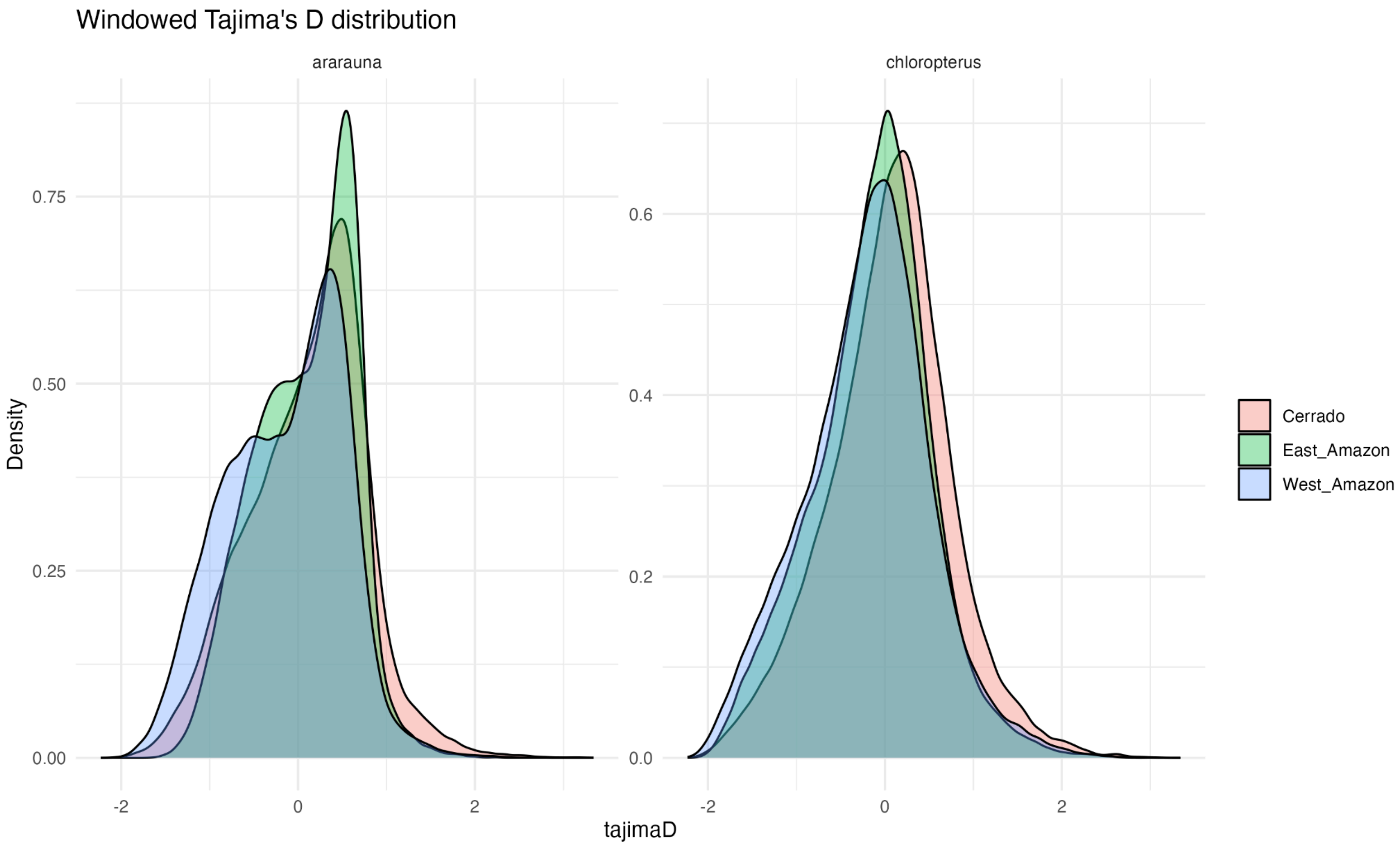

**Figure S44**. Distribution of Tajima's D across genomic windows in *Ara ararauna* and *Ara chloropterus* per population.

**Table S1.** Sample metadata and sequencing statistics for *Ara ararauna* individuals included in this study. Specimen information (voucher number, locality, and sex) is based on original tag descriptions available from VertNet (https://vertnet.org/). In addition we report the sex identity determined using SATC (Sex Assignment Through Coverage) (Nursyifa et al. 2022). Genome-wide sequencing statistics were calculated after filtering out low-quality reads, mapped to the *Ara ararauna* bAraAra1.hap1 reference genome (GCA_028858755.1), as described in the Materials and Methods. Mitochondrial DNA coverage is based on mapping to species-specific mitogenomes available on NCBI (accessions KF946546.1, NC_029319.1, and NC_047199.1), and similarly reflect the mapped reads after filtering. Abbreviation of source material (in "Voucher Number"): FMNH: Field Museum of Natural History, USA; LSUMZ: Louisiana Museum of Natural History, USA; MNHN: Muséum national d'Histoire naturelle, France; RMNH: Rijksmuseum van Natuurlijke Historie, Netherlands; YPM: Yale Peabody Museum, USA; MVZ: Museum of Vertebrate Zoology, USA; MUSM: Museo de Historia Natural de San Marcos, Peru; LGEMA: Laboratório de Genética e Evolução Molecular de Aves, Brazil.

| **Voucher Number** | **Species** | **Country** | **Province** | **Locality** | **Sex indicated on specimen voucher** | **Sex determined via SATC** | **Sample type** | **Genome-wide coverage** | **Endogenous portion** | **Biome grouping for genome-wide analyses** | **Mitogenome coverage** | **Mitogenome Size** | **Mitochondrial haplogroup** | **NCBI accession number** |
|---|---|---|---|---|---|---|---|---|---|---|---|---|---|---|
| LSUMZ35768 | *Ara ararauna* | Bolivia | | Santa cruz | | | Toe pad | 0.00 | 0.02 | | 84.55 | 16,981 | A | PV918067 |
| FMNH179049 | *Ara ararauna* | Bolivia | Santa Cruz | Rio Ichilo | Male | | Toe pad | 0.01 | 0.16 | | 43.77 | 16,981 | A | PV918069 |
| MNHN1960_247 | *Ara ararauna* | Bolivia | Ichilo | Buena Vista, Ichilo, Santa Cruz | | Female (ZW) | Toe pad | 0.77 | 0.71 | Amazon-West | 171.35 | 16,980 | A | PV918072 |

| | | | | | | | | | | | | | | |
|---|---|---|---|---|---|---|---|---|---|---|---|---|---|---|
| FMNH179046 | *Ara ararauna* | Bolivia | Santa Cruz | Ichilo, Buenavista | Male | Female (ZW) | Toe pad | 1.03 | 0.80 | Amazon-West | 376.76 | 16,981 | A | PV918070 |
| MNHN1960_249 | *Ara ararauna* | Bolivia | Ichilo | Buena Vista, Ichilo, Santa Cruz | | Female (ZW) | Toe pad | 1.16 | 0.51 | Amazon-West | 513.28 | 16,981 | A | PV918068 |
| LSUMZ35769 | *Ara ararauna* | Bolivia | Cochabamba | Chipiriri, Chapare Province | Male | Male (ZZ) | Toe pad | 1.18 | 0.68 | Amazon-West | 254.01 | 16,981 | A | PV918066 |
| LSUMZ37245 | *Ara ararauna* | Bolivia | Santa cruz | Buena vista, Ichilo Province | Male | Male (ZZ) | Toe pad | 1.47 | 0.68 | Amazon-West | 294.65 | 16,981 | A | PV918062 |
| FMNH62484 | *Ara ararauna* | Bolivia | Santa Cruz | Ichilo, Buenavista | Male | Male (ZZ) | Toe pad | 3.78 | 0.73 | Amazon-West | 1,076.01 | 16,981 | A | PV918065 |
| LGEMA 9050 | *Ara ararauna* | Brazil | PA | Redenção. Projeto Indígena Pinkaiti | | Male (ZZ) | Blood | 1.77 | 0.83 | Cerrado | 19.65 | 16,980 | A | PV918087 |
| LGEMA 15314 | *Ara ararauna* | Brazil | MS | Campo Grande | | Male (ZZ) | Blood | 12.69 | 0.92 | Cerrado | 146.34 | 16,980 | A | PV918084 |
| LGEMA 15991 | *Ara ararauna* | Brazil | MS | Campo Grande | | Male (ZZ) | Blood | 2.01 | 0.92 | Cerrado | 44.08 | 16,980 | A | PV918083 |
| LGEMA 6495 | *Ara ararauna* | Brazil | GO | Região Nordeste Do | | Female (ZW) | Blood | 2.54 | 0.89 | Cerrado | 18.32 | 16,980 | A | PV918093 |

| | | | | | | | | | | | | | | |
|---|---|---|---|---|---|---|---|---|---|---|---|---|---|---|
| | | | | Estado de Goiás | | | | | | | | | | |
| LGEMA 4810 | *Ara ararauna* | Brazil | GO | Mineiros. Parque Nacional das Emas | | Male (ZZ) | Blood | 2.83 | 0.90 | Cerrado | 36.90 | 16,980 | A | PV918082 |
| LGEMA 5254 | *Ara ararauna* | Brazil | PA | Ilha Mexiana | | Female (ZW) | Blood | 2.88 | 0.91 | Amazon-East | 40.38 | 16,980 | B2 | PV918118 |
| LGEMA 7540 | *Ara ararauna* | Brazil | MS | Corginho. Fazenda Altamira | | Female (ZW) | Blood | 2.99 | 0.92 | | 110.67 | 16,980 | B3 | PV918127 |
| LGEMA 4804 | *Ara ararauna* | Brazil | GO | Mineiros. Parque Nacional das Emas | | Female (ZW) | Blood | 3.00 | 0.89 | Cerrado | 17.59 | 16,980 | B3 | PV918124 |
| LGEMA 4812 | *Ara ararauna* | Brazil | GO | Mineiros. Parque Nacional das Emas | | Female (ZW) | Blood | 3.06 | 0.88 | Cerrado | 21.08 | 16,980 | B3 | PV918122 |
| LGEMA 18958 | *Ara ararauna* | Brazil | MT | Cidade de Rondonópolis MT | | Male (ZZ) | Blood | 3.06 | 0.91 | Cerrado | 9.36 | 16,980 | B3 | PV918130 |
| LGEMA 7470 | *Ara ararauna* | Brazil | TO | Peixe | | Female (ZW) | Blood | 3.09 | 0.93 | Cerrado | 45.29 | 16,980 | A | PV918075 |

| | | | | | | | | | | | | | | |
|---|---|---|---|---|---|---|---|---|---|---|---|---|---|---|
| LGEMA 5081 | *Ara ararauna* | Brazil | MG | Serra do Cipó | | Female (ZW) | Blood | 3.10 | 0.88 | Cerrado | 14.93 | 16,980 | B3 | PV918121 |
| LGEMA 7536 | *Ara ararauna* | Brazil | MS | Corginho. Fazenda Araras | | Male (ZZ) | Blood | 3.11 | 0.93 | Cerrado | 148.41 | 16,980 | B3 | PV918128 |
| LGEMA 5083 | *Ara ararauna* | Brazil | MG | Serra do Cipó | | Female (ZW) | Blood | 3.12 | 0.88 | Cerrado | 17.86 | 16,980 | B3 | PV918123 |
| LGEMA 7474 | *Ara ararauna* | Brazil | TO | São Valério | | Female (ZW) | Blood | 3.14 | 0.91 | Cerrado | 57.10 | 16,980 | A | PV918092 |
| LGEMA 7533 | *Ara ararauna* | Brazil | MS | Corginho. Fazenda Araras | | Female (ZW) | Blood | 3.22 | 0.92 | Cerrado | 106.92 | 16,980 | A | PV918080 |
| LGEMA 5253 | *Ara ararauna* | Brazil | PA | Ilha Caviana. Rio Busutubinha | | Male (ZZ) | Blood | 3.23 | 0.94 | Amazon-East | 59.08 | 16,981 | B2 | PV918116 |
| LGEMA 12339 | *Ara ararauna* | Brazil | MT | Paranaíta | | Female (ZW) | Blood | 3.26 | 0.74 | Central | 6,583.80 | 16,980 | B3 | PV918126 |
| LGEMA 13488 | *Ara ararauna* | Brazil | AC | Rio Branco | | Male (ZZ) | Blood | 3.26 | 0.93 | Amazon-West | 85.93 | 16,980 | A | PV918085 |
| LGEMA 7472 | *Ara ararauna* | Brazil | TO | São Valério | | Male (ZZ) | Blood | 3.33 | 0.94 | Cerrado | 59.07 | 16,980 | A | PV918079 |
| LGEMA 15989 | *Ara ararauna* | Brazil | MS | Campo Grande | | Female (ZW) | Blood | 3.40 | 0.92 | Cerrado | 59.48 | 16,980 | B3 | PV918132 |

| | | | | | | | | | | | | | | |
|---|---|---|---|---|---|---|---|---|---|---|---|---|---|---|
| LGEMA 8934 | *Ara ararauna* | Brazil | TO | Palmas. Perímetro urbano | | Female (ZW) | Blood | 3.52 | 0.92 | Cerrado | 200.44 | 16,980 | A | PV918071 |
| LGEMA 5087 | *Ara ararauna* | Brazil | MG | Serra do Cipó | | Male (ZZ) | Blood | 3.63 | 0.89 | Cerrado | 16.01 | 16,980 | B3 | PV918131 |
| LGEMA 8935 | *Ara ararauna* | Brazil | TO | Palmas. Resgate-Lajea do | | Male (ZZ) | Blood | 3.63 | 0.93 | Cerrado | 41.78 | 16,980 | A | PV918086 |
| LGEMA 6494 | *Ara ararauna* | Brazil | GO | Região Nordeste Do Estado de Goiás | | Male (ZZ) | Blood | 3.66 | 0.93 | Cerrado | 71.95 | 16,980 | B3 | PV918135 |
| LGEMA 15986 | *Ara ararauna* | Brazil | MS | Campo Grande | | Female (ZW) | Blood | 3.70 | 0.91 | Cerrado | 137.27 | 16,980 | B3 | PV918133 |
| LGEMA 7532 | *Ara ararauna* | Brazil | MS | Corginho. Fazenda Pedra Bonita | | Female (ZW) | Blood | 3.76 | 0.91 | Cerrado | 195.78 | 16,980 | B3 | PV918125 |
| LGEMA 6499 | *Ara ararauna* | Brazil | GO | Região Nordeste Do Estado de Goiás | | Male (ZZ) | Blood | 4.01 | 0.90 | Cerrado | 63.15 | 16,980 | A | PV918095 |
| LGEMA 7542 | *Ara ararauna* | Brazil | MS | Corginho. Fazenda Altamira | | Male (ZZ) | Blood | 4.05 | 0.93 | Cerrado | 232.20 | 16,980 | A | PV918074 |

| | | | | | | | | | | | | | | |
|---|---|---|---|---|---|---|---|---|---|---|---|---|---|---|
| LGEMA 6505 | *Ara ararauna* | Brazil | GO | Região Nordeste Do Estado de Goiás | | Male (ZZ) | Blood | 4.08 | 0.92 | Cerrado | 56.38 | 16,981 | A | PV918094 |
| LGEMA 4807 | *Ara ararauna* | Brazil | GO | Mineiros. Parque Nacional das Emas | | Male (ZZ) | Blood | 4.43 | 0.91 | Cerrado | 38.93 | 16,981 | B3 | PV918129 |
| LGEMA 4808 | *Ara ararauna* | Brazil | GO | Mineiros. Parque Nacional das Emas | | Male (ZZ) | Blood | 5.23 | 0.91 | Cerrado | 35.95 | 16,980 | A | PV918088 |
| LGEMA 6502 | *Ara ararauna* | Brazil | GO | Região Nordeste Do Estado de Goiás | | Male (ZZ) | Blood | 5.26 | 0.91 | Cerrado | 29.65 | 16,979 | A | PV918078 |
| LGEMA 15984 | *Ara ararauna* | Brazil | MS | Campo Grande | | Female (ZW) | Blood | 5.53 | 0.92 | Cerrado | 200.75 | 16,980 | A | PV918081 |
| MVZ138151 | *Ara ararauna* | Colombia | Putumayo | Puerto Asis, Putumayo | Male | Male (ZZ) | Toe pad | 1.46 | 0.69 | Amazon-West | 608.49 | NA | NA | NA |
| FMNH190744 | *Ara ararauna* | Colombia | Antioquia | Nechi | Female | Female (ZW) | Toe pad | 2.61 | 0.54 | Isolated | 4,419.73 | 16,980 | B3 | PV918120 |
| MNHN1937_919 | *Ara ararauna* | Ecuador | Pastaza | El Singuin, Canelos, | Male | Male (ZZ) | Toe pad | 1.52 | 0.75 | Amazon-West | 146.82 | 16,981 | A | PV918056 |

| | | | | | | | | | | | | | | |
|---|---|---|---|---|---|---|---|---|---|---|---|---|---|---|
| | | | | Pastaza, Pastaza | | | | | | | | | | |
| captive | *Ara ararauna* | France | Captive | Beauval Zoo | | | Feather | 0.10 | 0.06 | | 65.06 | 16,980 | C | PV918107 |
| MNHN1992_2006 | *Ara ararauna* | French Guiana | Camopi | Trois sauts, Camopi, Cayenne | | | Toe pad | 0.11 | 0.45 | | 9.84 | NA | NA | NA |
| MNHN1937_920 | *Ara ararauna* | French Guiana | Camopi | Trois sauts, Camopi, Cayenne | | Female (ZW) | Toe pad | 1.01 | 0.76 | Amazon-East | 286.12 | 16,982 | B1 | PV918112 |
| MNHN1963_767 | *Ara ararauna* | French Guiana | | Fleuve Maroni, Saint Laurent du Maroni | | Female (ZW) | Toe pad | 1.55 | 0.64 | Amazon-East | 3,510.61 | 16,980 | A | PV918091 |
| FMNH43868 | *Ara ararauna* | Guyana | | Courentyne River | | | Toe pad | 0.00 | 0.01 | | 36.63 | NA | NA | NA |
| FMNH32214 | *Ara ararauna* | Guyana | | Courentyne River | | | Toe pad | 0.01 | 0.04 | | 235.11 | 16,980 | B2 | PV918117 |
| FMNH40261 | *Ara ararauna* | Guyana | | | NA | Male (ZZ) | Toe pad | 4.08 | 0.77 | Amazon-East | 1,085.72 | 16,980 | B2 | PV918115 |
| MUSMf11053 | *Ara ararauna* | Peru | Madre de Dios | Tambopata: TRC | | | Feather | 0.00 | 0.01 | | 100.29 | 16,980 | A2 | PV918106 |
| MUSMf11125 | *Ara ararauna* | Peru | Loreto | Centro Turistico | | | Feather | 0.00 | 0.00 | | 2.56 | NA | NA | NA |

| | | | | | | | | | | | | | | |
|---|---|---|---|---|---|---|---|---|---|---|---|---|---|---|
| | | | | Quistococha, Iquitos | | | | | | | | | | |
| MUSMf11126 | *Ara ararauna* | Peru | Loreto | Centro Turistico Quistococha, Iquitos | | | Feather | 0.00 | 0.00 | | 0.41 | NA | NA | NA |
| LSUMZ31260 | *Ara ararauna* | Peru | Ucayali | Ucayali | | | Toe pad | 0.01 | 0.09 | | 312.81 | 16,980 | A | PV918073 |
| LSUMZ34008 | *Ara ararauna* | Peru | Ucayali | Ucayali | | | Toe pad | 0.01 | 0.08 | | 336.83 | 16,981 | A | PV918077 |
| LSUMZ84327 | *Ara ararauna* | Peru | Madre de Dios | | Male | | Toe pad | 0.01 | 0.19 | | 8.35 | NA | NA | NA |
| MUSMf11130 | *Ara ararauna* | Peru | Loreto | Artesanias ANACONDA, Iquitos | | | Feather | 0.01 | 0.01 | | 4.95 | NA | NA | NA |
| MUSMf11131 | *Ara ararauna* | Peru | Loreto | Artesanias ANACONDA, Iquitos | | | Feather | 0.01 | 0.01 | | 18.93 | NA | NA | NA |
| MUSMf11146 | *Ara ararauna* | Peru | Ucayali | Bolsa de plumas, Mercado Nº2, Pucallpa | | | Feather | 0.01 | 0.01 | | 1.13 | NA | NA | NA |
| MUSMf11110A | *Ara ararauna* | Peru | Loreto | Artesanias ANACONDA, Iquitos | | | Feather | 0.02 | 0.02 | | 0.41 | NA | NA | NA |

| | | | | | | | | | | | | | | |
|---|---|---|---|---|---|---|---|---|---|---|---|---|---|---|
| MUSMf11134 | *Ara ararauna* | Peru | Ucayali | Comunidad "Once de Agosto", Yarinacocha, Pucallpa | | | Feather | 0.02 | 0.04 | | 112.16 | 16,981 | A | PV918063 |
| MUSMf11137 | *Ara ararauna* | Peru | Ucayali | Parque Natural de Pucallpa, Pucallpa | | | Feather | 0.02 | 0.02 | | 27.69 | 16,981 | B1 | PV918111 |
| MUSMf11139 | *Ara ararauna* | Peru | Ucayali | "La Jungla", Lago Yarinacocha, Pucallpa | | | Feather | 0.02 | 0.02 | | 67.79 | 16,982 | A | PV918064 |
| LSUMZ34009 | *Ara ararauna* | Peru | Ucayali | Rio Curanja, Balta | Male | | Toe pad | 0.03 | 0.27 | | 157.15 | 16,981 | A2 | PV918096 |
| MUSMf11105 | *Ara ararauna* | Peru | Loreto | Centro de Artesanías en plaza de Iquitos | | | Feather | 0.04 | 0.06 | | 10.98 | NA | NA | NA |
| MUSMf11118 | *Ara ararauna* | Peru | Loreto | CREA, Iquitos | | | Feather | 0.07 | 0.07 | | 105.87 | 16,982 | A2 | PV918105 |
| MUSMf11111C | *Ara ararauna* | Peru | Loreto | “El Serpentario” Rio momón, Iquitos | | | Feather | 0.12 | 0.13 | | 156.14 | 16,981 | A3 | PV918059 |
| MUSMf11121 | *Ara ararauna* | Peru | Loreto | CREA, Iquitos | | | Feather | 0.14 | 0.09 | | 350.43 | 16,980 | A2 | PV918097 |

| | | | | | | | | | | | | | | |
|---|---|---|---|---|---|---|---|---|---|---|---|---|---|---|
| MUSMf11136 | *Ara ararauna* | Peru | Ucayali | "Costa del Ucayali", Yarinacocha, Pucallpa | | | Feather | 0.21 | 0.09 | | 465.94 | 16,981 | B1 | PV918113 |
| MUSMf11096B | *Ara ararauna* | Peru | Loreto | Centro de Artesanías en plaza de Iquitos | | Male (ZZ) | Feather | 0.85 | 0.80 | Amazon-West | 200.05 | 16,980 | B3 | PV918119 |
| YPM081423 | *Ara ararauna* | Peru | Cuzco | Huajyumbe | Male | Female (ZW) | Toe pad | 0.88 | 0.72 | Amazon-West | 291.53 | 16,981 | A2 | PV918061 |
| YPM081422 | *Ara ararauna* | Peru | Cuzco | Huajyumbe | Male | Male (ZZ) | Toe pad | 1.08 | 0.73 | Amazon-West | 413.08 | 16,981 | A2 | PV918102 |
| YPM081424 | *Ara ararauna* | Peru | Cuzco | Huajyumbe | Male | Female (ZW) | Toe pad | 1.21 | 0.77 | Amazon-West | 387.74 | 16,981 | A | PV918076 |
| LSUMZ63922 | *Ara ararauna* | Peru | Ucayali | Rio Curanja, Balta | Female | Male (ZZ) | Toe pad | 1.23 | 0.57 | Amazon-West | 251.18 | 16,981 | A2 | PV918100 |
| MUSM11059 | *Ara ararauna* | Peru | Madre de Dios | Tambopata: TRC | | Male (ZZ) | Toe pad | 1.24 | 0.72 | Amazon-West | 246.75 | 16,980 | A3 | PV918057 |
| MUSMf11096A | *Ara ararauna* | Peru | Loreto | Centro de Artesanías en plaza de Iquitos | | Male (ZZ) | Feather | 1.30 | 0.65 | Amazon-West | 297.60 | 16,981 | A2 | PV918104 |
| MUSMf11116 | *Ara ararauna* | Peru | Loreto | CREA, Iquitos | | Male (ZZ) | Feather | 1.40 | 0.25 | Amazon-West | 3,174.35 | 16,980 | A2 | PV918103 |

| | | | | | | | | | | | | | | |
|---|---|---|---|---|---|---|---|---|---|---|---|---|---|---|
| MUSMf11117 | *Ara ararauna* | Peru | Loreto | CREA, Iquitos | | Female (ZW) | Feather | 1.43 | 0.34 | Amazon-West | 1,817.03 | 16,980 | A2 | PV918098 |
| MUSMf11128 | *Ara ararauna* | Peru | Loreto | Centro de Rescate Pilpintawasi, Iquitos | | Male (ZZ) | Feather | 1.44 | 0.64 | Amazon-West | 616.75 | 16,980 | A3 | PV918058 |
| MUSM865 | *Ara ararauna* | Peru | Loreto | Rio Pacaya | Male | Male (ZZ) | Toe pad | 1.51 | 0.65 | Amazon-West | 611.53 | 16,981 | A2 | PV918099 |
| MUSM1980 | *Ara ararauna* | Peru | Ucayali | Yarinacocha, rio Ucayali | Female | Male (ZZ) | Toe pad | 1.83 | 0.72 | Amazon-West | 478.67 | NA | NA | NA |
| MUSM1981 | *Ara ararauna* | Peru | Ucayali | Yarinacocha (Rio Ucayali) | Female | Female (ZW) | Toe pad | 1.94 | 0.59 | Amazon-West | 549.25 | 16,981 | A2 | PV918060 |
| MUSMf11115A | *Ara ararauna* | Peru | Loreto | Fundo Pedrito, Barrio Florido, Iquitos | | Female (ZW) | Feather | 1.97 | 0.53 | Amazon-West | 29.64 | 16,980 | B2 | PV918114 |
| MUSMf11132 | *Ara ararauna* | Peru | Captive | San Isidro ("Blady") | | Female (ZW) | Feather | 2.37 | 0.62 | Amazon-West | 698.88 | 16,981 | A2 | PV918101 |
| RMNH208365 | *Ara ararauna* | Suriname | | | | | Toe pad | 0.01 | 0.01 | | 2,323.62 | 16,980 | B3 | PV918134 |
| FMNH260145 | *Ara ararauna* | Suriname | Nickerie | Kaiserberg Airstrip, Zuid River | Female | Female (ZW) | Toe pad | 1.37 | 0.66 | Amazon-East | 4,276.61 | 16,980 | A | PV918089 |
| RMNH65980 | *Ara ararauna* | Suriname | | Awarra savanne | Female | Female (ZW) | Toe pad | 1.47 | 0.40 | Amazon-East | 4,286.38 | 16,981 | C | PV918109 |

| | | | | | | | | | | | | | | |
|---|---|---|---|---|---|---|---|---|---|---|---|---|---|---|
| RMNH36195 | *Ara ararauna* | Suriname | | Suriname rivier | Female | Female (ZW) | Toe pad | 1.48 | 0.69 | Amazon-East | 1,114.27 | 16,979 | C | PV918110 |
| RMNH36194 | *Ara ararauna* | Suriname | | Suriname rivier | Male | Male (ZZ) | Toe pad | 1.56 | 0.72 | Amazon-East | 2,778.88 | 16,979 | C | PV918108 |
| captive | *Ara ararauna* | Uk | Captive | | | Female (ZW) | Feather | 0.27 | 0.48 | | 1,980.13 | 16,980 | A | PV918090 |

**Table S2.** Sample metadata and sequencing statistics for *Ara chloropterus* individuals included in this study. Specimen information (voucher number, locality, and sex) is based on original tag descriptions available from VertNet (https://vertnet.org/). In addition we report the sex identity determined using Sex Assignment Through Coverage (Nursyifa et al. 2022). Genome-wide sequencing statistics reflect filtered reads (Phred ≥ 25) mapped to the *Ara ararauna* bAraAra1.hap1 reference genome (GCA_028858755.1). Mitochondrial DNA coverage is based on mapping to species-specific mitogenomes available on NCBI (accessions KF946546.1, NC_029319.1, and NC_047199.1). Abbreviation of source material (in "Voucher Number"): FMNH: Field Museum of Natural History, USA; LSUMZ: Louisiana Museum of Natural History, USA; MNHN: Muséum national d'Histoire naturelle, France; RMNH: Rijksmuseum van Natuurlijke Historie, Netherlands; YPM: Yale Peabody Museum, USA; MVZ: Museum of Vertebrate Zoology, USA; MUSM: Museo de Historia Natural de la Universidad Nacional de San Marcos, Peru; LGEMA: Laboratório de Genética e Evolução Molecular de Aves, Brazil.

| **Voucher Number** | **Species** | **Country** | **Province** | **Locality** | **Sex (Voucher)** | **Sex (SATC)** | **Type of Sample** | **Genome-wide coverage (MinQ ≥ 25)** | **Endogenous content (MinQ ≥ 25)** | **Biome-population (Genome-wide Analyses)** | **Mitogenome Coverage (MinQ ≥ 25)** | **Mitogenome Size** | **Mitochondrial Haplogroup** | **NCBI Accession Number** |
|---|---|---|---|---|---|---|---|---|---|---|---|---|---|---|
| FMNH179050 | *Ara chloropterus* | Bolivia | Santa Cruz | Rio Ichilo | Female | | Toe pad | 0.00 | 0.03 | | 24.53 | 16,994 | A | PV918175 |
| FMNH179051 | *Ara chloropterus* | Bolivia | Santa Cruz | Rio Ichilo | Female | | Toe pad | 0.01 | 0.49 | | 16.56 | 16,994 | A | PV918149 |
| FMNH179052 | *Ara chloropterus* | Bolivia | Santa Cruz | Rio Surutú | Male | Male (ZZ) | Toe pad | 2.16 | 0.58 | Amazon-West | 913.67 | 16,994 | A | PV918178 |
| FMNH179053 | *Ara chloropterus* | Bolivia | Santa Cruz | Rio Surutú | Male | | Toe pad | 0.01 | 0.44 | | 6.15 | NA | NA | NA |

| | | | | | | | | | | | | | | |
|---|---|---|---|---|---|---|---|---|---|---|---|---|---|---|
| FMNH179054 | *Ara chloropterus* | Bolivia | Santa Cruz | Rio Yapacani | Male | Male (ZZ) | Toe pad | 1.12 | 0.58 | Amazon-West | 283.69 | 16,994 | A | PV918176 |
| FMNH179055 | *Ara chloropterus* | Bolivia | Santa Cruz | Rio Yapacani | Female | | Toe pad | 0.02 | 0.35 | | 3.72 | NA | NA | NA |
| FMNH179056 | *Ara chloropterus* | Bolivia | Cochabamba | El Palmar, Yungas | Female | Female (ZW) | Toe pad | 1.14 | 0.53 | Amazon-West | 263.35 | 16,994 | A | PV918174 |
| LSUMZ123498 | *Ara chloropterus* | Bolivia | Beni | 50 km by road N Yacumo on road to Rurrenabaque | Male | Male (ZZ) | Toe pad | 1.53 | 0.78 | Amazon-West | 112.45 | 16,994 | D | PV918141 |
| MNHN1960-250 | *Ara chloropterus* | Bolivia | Santa Cruz | Rio Surutú, Ichilo | | Male (ZZ) | Toe pad | 1.41 | 0.65 | Amazon-West | 277.20 | 16,994 | A | PV918151 |
| MNHN1996-531 | *Ara chloropterus* | Bolivia | | | Male | Male (ZZ) | Toe pad | 0.30 | 0.48 | | 87.36 | NA | NA | NA |
| YPM038365 | *Ara chloropterus* | Bolivia | Cochabamba | Chapare | Female | Female (ZW) | Toe pad | 1.42 | 0.68 | Amazon-West | 416.54 | 16,994 | A | PV918177 |
| YPM038366 | *Ara chloropterus* | Bolivia | Beni | Chutihuara (Municipality of Reyes) | | Male (ZZ) | Toe pad | 3.00 | 0.59 | Amazon-West | 622.62 | 16,994 | A | PV918163 |
| YPM038367 | *Ara chloropterus* | Bolivia | Beni | Chutihuara (Municipality of Reyes) | | | Toe pad | 0.04 | 0.46 | | 29.92 | 16,994 | A | PV918179 |
| FMNH257844 | *Ara chloropterus* | Brazil | PA | Fordlandia, Rio Tapajos | Female | | Toe pad | 0.03 | 0.55 | | 24.75 | 16,995 | E | PV918136 |
| FMNH257845 | *Ara chloropterus* | Brazil | PA | Fordlandia, Rio Tapajos | | | Toe pad | 0.01 | 0.31 | | 9.12 | NA | NA | NA |

| | | | | | | | | | | | | | | |
|---|---|---|---|---|---|---|---|---|---|---|---|---|---|---|
| LGEMA10315 | *Ara chloropterus* | Brazil | MT | Barão de Melgaço, SESC Pantanal | | Female (ZW) | Blood | 6.34 | 0.87 | Cerrado-South | 78.60 | 16,994 | A | PV918153 |
| LGEMA11272 | *Ara chloropterus* | Brazil | MT | Barão de Melgaço, SESC Pantanal | | Female (ZW) | Blood | 5.48 | 0.87 | Cerrado-South | 82.70 | 16,992 | D | PV918142 |
| LGEMA11457 | *Ara chloropterus* | Brazil | PA | Canaã dos Carajás | | Female (ZW) | Blood | 3.70 | 0.88 | Amazon-East | 36.84 | 16,994 | B1 | PV918196 |
| LGEMA11458 | *Ara chloropterus* | Brazil | PA | Canaã dos Carajás | | Male (ZZ) | Blood | 4.01 | 0.88 | Amazon-East | 38.32 | 16,994 | B1 | PV918194 |
| LGEMA11459 | *Ara chloropterus* | Brazil | PA | Canaã dos Carajás | | Female (ZW) | Blood | 4.50 | 0.88 | Amazon-East | 62.53 | 16,993 | A | PV918155 |
| LGEMA11460 | *Ara chloropterus* | Brazil | PA | Canaã dos Carajás | | Female (ZW) | Blood | 4.20 | 0.90 | Amazon-East | 85.94 | 16,993 | A | PV918156 |
| LGEMA11461 | *Ara chloropterus* | Brazil | PA | Canaã dos Carajás | | Female (ZW) | Blood | 20.68 | 0.88 | Amazon-East | 1216.59 | 16,994 | B1 | PV918193 |
| LGEMA11487 | *Ara chloropterus* | Brazil | PI | Boa Vista, Frente da casa | | Male (ZZ) | Blood | 3.75 | 0.89 | Cerrado-North | 39.41 | 16,995 | B1 | PV918188 |
| LGEMA11488 | *Ara chloropterus* | Brazil | PI | Boa Vista, Subida da Serra do Suvaco | | Female (ZW) | Blood | 3.40 | 0.88 | Cerrado-North | 22.46 | 16,994 | B2 | PV918205 |
| LGEMA11489 | *Ara chloropterus* | Brazil | PI | Boa Vista, Serra do Capucho | | Female (ZW) | Blood | 4.08 | 0.88 | Cerrado-North | 54.44 | 16,995 | B2 | PV918206 |
| LGEMA11499 | *Ara chloropterus* | Brazil | PI | São Gonçalo de Gurguéia, Serra do Curralinho | | Male (ZZ) | Blood | 5.10 | 0.89 | Cerrado-North | 86.71 | 16,995 | B2 | PV918207 |

| | | | | | | | | | | | | | | |
|---|---|---|---|---|---|---|---|---|---|---|---|---|---|---|
| LGEMA11500 | *Ara chloropterus* | Brazil | PI | São Gonçalo de Gurguéia, Serra do Curralinho | | Male (ZZ) | Blood | 4.94 | 0.89 | Cerrado-North | 57.06 | 16,995 | B2 | PV918208 |
| LGEMA11504 | *Ara chloropterus* | Brazil | PI | São Gonçalo de Gurguéia, Serra do Guará | | Female (ZW) | Blood | 3.87 | 0.89 | Cerrado-North | 127.78 | 16,995 | B2 | PV918204 |
| LGEMA11924 | *Ara chloropterus* | Brazil | RO | Parque Nacional de Pacaás Novos | | Female (ZW) | Blood | 2.89 | 0.85 | Amazon-West | 21.02 | 16,993 | E | PV918138 |
| LGEMA4437 | *Ara chloropterus* | Brazil | MS | Nhecolândia | | Female (ZW) | Blood | 3.20 | 0.83 | Cerrado-South | 45.10 | 16,992 | D | PV918143 |
| LGEMA4439 | *Ara chloropterus* | Brazil | MS | Nhecolândia | | Male (ZZ) | Blood | 2.27 | 0.83 | Cerrado-South | 9.77 | NA | NA | |
| LGEMA4441 | *Ara chloropterus* | Brazil | MS | Nhecolândia | | Male (ZZ) | Blood | 3.32 | 0.86 | Cerrado-South | 31.53 | 16,995 | B2 | PV918201 |
| LGEMA4976 | *Ara chloropterus* | Brazil | MS | Nhecolândia | | Male (ZZ) | Blood | 2.95 | 0.84 | Cerrado-South | 31.99 | 16,993 | E | PV918137 |
| LGEMA4986 | *Ara chloropterus* | Brazil | MS | Abobral | | Male (ZZ) | Blood | 2.87 | 0.86 | Cerrado-South | 27.70 | 16,992 | D | PV918139 |
| LGEMA4988 | *Ara chloropterus* | Brazil | MS | Abobral | | Male (ZZ) | Blood | 2.80 | 0.86 | Cerrado-South | 20.76 | 16,992 | D | PV918147 |
| LGEMA9149 | *Ara chloropterus* | Brazil | MT | Barão de Melgaço, SESC Pantanal | | Female (ZW) | Blood | 6.00 | 0.87 | Cerrado-South | 95.65 | 16,992 | D | PV918144 |
| LGEMA9151 | *Ara chloropterus* | Brazil | MT | Barão de Melgaço, SESC Pantanal | | Female (ZW) | Blood | 7.63 | 0.87 | Cerrado-South | 68.40 | 16,992 | D | PV918145 |

| | | | | | | | | | | | | | | |
|---|---|---|---|---|---|---|---|---|---|---|---|---|---|---|
| LGEMA9153 | *Ara chloropterus* | Brazil | MT | Barão de Melgaço, SESC Pantanal | | Male (ZZ) | Blood | 4.24 | 0.90 | Cerrado-South | 38.39 | 16,995 | B2 | PV918203 |
| LGEMA9154 | *Ara chloropterus* | Brazil | MT | Barão de Melgaço, SESC Pantanal | | Female (ZW) | Blood | 3.31 | 0.87 | Cerrado-South | 41.90 | 16,992 | D | PV918146 |
| LSUMZ210126 | *Ara chloropterus* | Brazil | RO | Reserva Biologica Rebid Duro Preto | Male | Male (ZZ) | Tissue | 1.91 | 0.74 | Amazon-West | 13298.07 | 16,993 | A | PV918164 |
| FMNH190745 | *Ara chloropterus* | Colombia | Antioquia | Cuturu | Female | Female (ZW) | Toe pad | 0.60 | 0.46 | Isolated | 482.97 | 16,994 | A | PV918150 |
| FMNH190746 | *Ara chloropterus* | Colombia | Antioquia | Cuturu | Male | | Toe pad | 0.05 | 0.42 | | 49.53 | 16,994 | D | PV918140 |
| FMNH261079 | *Ara chloropterus* | Colombia | Boyaca | Bojaba | Male | | Toe pad | 0.02 | 0.44 | | 69.61 | 16,994 | A | PV918166 |
| FMNH261080 | *Ara chloropterus* | Colombia | Arauca | Rio Arauca | Female | | Toe pad | 0.02 | 0.41 | | 27.39 | 16,994 | C | PV918183 |
| FMNH261081 | *Ara chloropterus* | Colombia | Boyaca | Bojaba | Male | Male (ZZ) | Toe pad | 1.50 | 0.74 | Amazon-West | 216.88 | 16,994 | C | PV918184 |
| FMNH261082 | *Ara chloropterus* | Colombia | Arauca | Rio Arauca | Male | Female (ZW) | Toe pad | 1.55 | 0.73 | Amazon-West | 185.68 | 16,994 | A | PV918167 |
| FMNH371813 | *Ara chloropterus* | Colombia | | | Female | | Toe pad | 0.03 | 0.39 | | 4.62 | NA | NA | NA |
| MNHN1996-530 | *Ara chloropterus* | Ecuador | Pastaza | Sarayacu, Rio Bobonnaza / Oriente | Male | | Toe pad | 0.02 | 0.07 | | 94.75 | 16,994 | B2 | PV918200 |

| | | | | | | | | | | | | | | |
|---|---|---|---|---|---|---|---|---|---|---|---|---|---|---|
| MNHN1992-2003 | *Ara chloropterus* | French Guiana | Cayenne | Trois sauts, Camopi, Cayenne | | | Toe pad | 0.03 | 0.42 | | 1.28 | NA | NA | NA |
| MNHN1992-2007 | *Ara chloropterus* | French Guiana | Cayenne | Trois sauts, Camopi, Cayenne | | Male (ZZ) | Toe pad | 1.52 | 0.55 | Amazon-East | 506.71 | 16,995 | B1 | PV918187 |
| FMNH108112 | *Ara chloropterus* | Guyana | E Demerara-W Coast Berbice | Rockstone, Essequibo River | Female | Female (ZW) | Toe pad | 2.38 | 0.65 | Amazon-East | 441.78 | 16,994 | B1 | PV918199 |
| FMNH108113 | *Ara chloropterus* | Guyana | Mazaruni-Potaro | Oko Mountains | Male | Male (ZZ) | Toe pad | 1.43 | 0.68 | Amazon-East | 116.90 | 16,994 | B1 | PV918197 |
| FMNH62485 | *Ara chloropterus* | Guyana | E Berbice-Corentyne | New Amsterdam | Female | Male (ZZ) | Toe pad | 0.22 | 0.22 | | 83.04 | 16,994 | B1 | PV918186 |
| NOR1 | *Ara chloropterus* | Norway | | Trondheim (Captive) | Female | Female (ZW) | Feather | 0.70 | 0.60 | Captive | 187.96 | 16,995 | B2 | PV918202 |
| FMNH251522 | *Ara chloropterus* | Peru | Madre de Dios | Collpa, Rio Tambopata | Male | Female (ZW) | Toe pad | 1.76 | 0.64 | Amazon-West | 283.25 | 16,994 | A | PV918161 |
| FMNH251523 | *Ara chloropterus* | Peru | Madre de Dios | Collpa, Rio Tambopata | | Male (ZZ) | Toe pad | 1.51 | 0.66 | Amazon-West | 356.68 | 16,994 | A | PV918157 |
| FMNH286657 | *Ara chloropterus* | Peru | Pasco | Cacazu | Male | Female (ZW) | Toe pad | 1.54 | 0.56 | Amazon-West | 475.91 | 16,995 | A | PV918170 |
| FMNH333804 | *Ara chloropterus* | Peru | Loreto | La Union, Contamana Mts | Male | Male (ZZ) | Toe pad | 0.69 | 0.80 | Amazon-West | 49.49 | 16,994 | A | PV918180 |
| LSUMZ31264 | *Ara chloropterus* | Peru | Ucayali | Rio Curanja, Balta | Female | Female (ZW) | Toe pad | 0.23 | 0.55 | | 33.18 | NA | NA | NA |

| | | | | | | | | | | | | | | |
|---|---|---|---|---|---|---|---|---|---|---|---|---|---|---|
| LSUMZ31265 | *Ara chloropterus* | Peru | Ucayali | Rio Curanja, Balta | Female | Female (ZW) | Toe pad | 4.36 | 0.39 | Amazon-West | 2986.95 | 16,994 | A | PV918158 |
| LSUMZ31266 | *Ara chloropterus* | Peru | Ucayali | Rio Curanja, Balta | Male | Male (ZZ) | Toe pad | 0.37 | 0.68 | | 9.26 | NA | NA | NA |
| LSUMZ34006 | Ara chloropterus | Peru | Ucayali | Rio Curanja, Balta | | | Toe pad | 0.00 | 0.28 | | 0.19 | NA | NA | NA |
| LSUMZ34007 | *Ara chloropterus* | Peru | Ucayali | Rio Curanja, Balta | Female | | Toe pad | 0.02 | 0.48 | | 39.70 | 16,994 | A | PV918162 |
| LSUMZ51577 | *Ara chloropterus* | Peru | Ucayali | Rio Curanja, Balta | Female | Female (ZW) | Toe pad | 1.56 | 0.71 | Amazon-West | 82.36 | 16,994 | A | PV918181 |
| LSUMZ84329 | *Ara chloropterus* | Peru | Madre de Dios | Quebrada Juliaca on Rio Heath | Female | | Toe pad | 0.00 | 0.04 | | 1.79 | NA | NA | NA |
| MUSM11066 | *Ara chloropterus* | Peru | Madre de Dios | TRC, Tambopata | | | Toe pad | 0.01 | 0.17 | | 3.81 | NA | NA | NA |
| MUSM11070 | *Ara chloropterus* | Peru | Madre de Dios | TRC, Tambopata | | | Toe pad | 0.03 | 0.17 | | 36.28 | 16,994 | A | PV918159 |
| MUSM11460 | *Ara chloropterus* | Peru | Ucayali | Lado Oeste del rio shesha, ca 65km ENE Pucallpa | Female | | Toe pad | 0.06 | 0.09 | | 545.77 | 16,994 | A | PV918173 |
| MUSM1976 | *Ara chloropterus* | Peru | San Martin | Tocache- R, Huallaga | Male | Male (ZZ) | Toe pad | 1.75 | 0.57 | Amazon-West | 1198.39 | 16,994 | A | PV918171 |
| MUSM1977 | *Ara chloropterus* | Peru | Ucayali | Yarinacocha (Rio Ucayali) | | Male (ZZ) | Toe pad | 1.64 | 0.60 | Amazon-West | 545.97 | 16,994 | A | PV918165 |

| | | | | | | | | | | | | | | |
|---|---|---|---|---|---|---|---|---|---|---|---|---|---|---|
| MUSM5721 | *Ara chloropterus* | Peru | Lima | Parque de las leyendas (Lima Zoo) | | Male (ZZ) | Toe pad | 0.61 | 0.72 | Amazon-West | 71.50 | 16,995 | A | PV918168 |
| MUSMf11054 | *Ara chloropterus* | Peru | Madre de Dios | TRC, Tambopata | | | Feather | 0.01 | 0.00 | | 58.06 | 16,994 | A | PV918160 |
| MUSMf11109A | *Ara chloropterus* | Peru | Loreto | Artesanias ANACONDA, Iquitos | | | Feather | 0.18 | 0.21 | | 4.78 | NA | NA | NA |
| MUSMf11120 | *Ara chloropterus* | Peru | Loreto | Iquitos, CREA | | Male (ZZ) | Feather | 1.41 | 0.57 | Amazon-West | 386.41 | 16,994 | A | PV918172 |
| MUSMf11129 | *Ara chloropterus* | Peru | Loreto | Iquitos, Centro de Rescate Pilpintawasi | | Male (ZZ) | Feather | 0.34 | 0.14 | | 140.47 | 16,991 | A | PV918169 |
| MUSMf11133 | *Ara chloropterus* | Peru | Lima | San Isidro (Rojo) | | Female (ZW) | Feather | 1.66 | 0.67 | Amazon-West | 368.18 | 16,994 | A | PV918154 |
| FMNH260147 | *Ara chloropterus* | Suriname | Nickerie | Kaiserberg Airstrip, Zuid River | | Male (ZZ) | Toe pad | 1.05 | 0.55 | Amazon-East | 355.85 | 16,994 | B1 | PV918191 |
| RMNH65981 | *Ara chloropterus* | Suriname | Sipaliwini | Lombok Val | Female | | Toe pad | 0.05 | 0.08 | | 1425.24 | 16,994 | B1 | PV918198 |
| FMNH355149 | *Ara chloropterus* | USA | Florida | Captive | Female | | Toe pad | 0.01 | 0.18 | | 2.07 | NA | NA | NA |
| FMNH371812 | *Ara chloropterus* | USA | Florida | Captive | Male | | Toe pad | 0.03 | 0.55 | | 3.30 | NA | NA | NA |
| FMNH110180 | *Ara chloropterus* | Venezuela | Zulia | Machiques, Rio Cogollo | Male | Male (ZZ) | Toe pad | 2.85 | 0.63 | Amazon-West | 296.85 | 16,994 | C | PV918182 |

| | | | | | | | | | | | | | |
|---|---|---|---|---|---|---|---|---|---|---|---|---|---|
| FMNH318668 | *Ara chloropterus* | Venezuela | Amazonas | Cerro de la Neblina, Base Camp | Male | Male (ZZ) | Toe pad | 1.76 | 0.64 | Amazon-East | 216.56 | 16,994 | B1 | PV918189 |
| FMNH34366 | *Ara chloropterus* | Venezuela | Tachira | Orope | Female | | Toe pad | 0.01 | 0.04 | | 65.06 | 16,994 | B1 | PV918185 |
| FMNH34367 | *Ara chloropterus* | Venezuela | Tachira | Orope | Male | | Toe pad | 0.05 | 0.25 | | 32.79 | 16,993 | A | PV918148 |
| FMNH34368 | *Ara chloropterus* | Venezuela | Tachira | Orope | Male | Male (ZZ) | Toe pad | 2.44 | 0.42 | Amazon-West | 1345.23 | 16,993 | A | PV918152 |
| FMNH81440 | Ara chloropterus | Venezuela | Delta Amacuro | Piacoa | Male | Male (ZZ) | Toe pad | 0.91 | 0.54 | Amazon-East | 197.63 | 16,994 | B1 | PV918195 |
| LSUMZ226526 | *Ara chloropterus* | Venezuela | Amazonas | Cerro de La Neblina, Base Camp | Male | Male (ZZ) | Tissue | 2.56 | 0.75 | Amazon-East | 20092.60 | 16,994 | B1 | PV918190 |
| LSUMZ843 | *Ara chloropterus* | Venezuela | Aragua | Urama | Male | | Toe pad | 0.00 | 0.10 | | 2.63 | NA | NA | NA |
| MNHN1930-272 | *Ara chloropterus* | Venezuela | Caura | | | | Toe pad | 0.13 | 0.29 | | 113.68 | NA | NA | NA |
| MNHN1930-273 | *Ara chloropterus* | Venezuela | Caura | | | Female (ZW) | Toe pad | 1.16 | 0.55 | Amazon-East | 442.81 | 16,994 | B1 | PV918192 |

**Table S3.** Sample metadata and sequencing statistics for *Ara severus* individuals included in this study. Specimen information (voucher number, locality, and sex) is based on original tag descriptions available from VertNet (https://vertnet.org/). In addition we report the sex identity determined using Sex Assignment Through Coverage (Nursyifa et al. 2022). Genome-wide sequencing statistics reflect filtered reads (Phred ≥ 25) mapped to the *Ara ararauna* bAraAra1.hap1 reference genome (GCA_028858755.1). Mitochondrial DNA coverage is based on mapping to species-specific mitogenomes available on NCBI (accessions KF946546.1, NC_029319.1, and NC_047199.1). Abbreviation of source material (in "Voucher Number"): FMNH: Field Museum of Natural History, USA; MVZ: Museum of Vertebrate Zoology, USA; MUSM: Museo de Historia Natural de la Universidad Nacional de San Marcos.

| Voucher Number | Species | Country | Province | Locality | Sex (Voucher) | Sex (SATC) | Type of Sample | Genome-wide coverage (MinQ ≥ 25) | Endogenous content (MinQ ≥ 25) | Mitogenome Coverage (MinQ ≥ 25) | Mitogenome Size | Mitochondrial Haplogroup | NCBI Accession Number |
|---|---|---|---|---|---|---|---|---|---|---|---|---|---|
| FMNH179066 | *Ara severus* | Bolivia | Santa Cruz | Santa Cruz, Ichilo, Buenavista | Male | Female | Toe pad | 4.39 | 0.61 | 964.72 | 16,996 | A1 | PV918229 |
| FMNH179067 | *Ara severus* | Bolivia | Santa Cruz | Santa Cruz, Ichilo, Buenavista | Male | Male | Toe pad | 2.45 | 0.63 | 900.95 | 16,996 | Unassigned | PV918242 |
| FMNH179068 | *Ara severus* | Bolivia | Santa Cruz | Santa Cruz, Ichilo, Buenavista | Male | Male | Toe pad | 4.13 | 0.38 | 2422.48 | 16,996 | A1 | PV918239 |
| FMNH179069 | *Ara severus* | Bolivia | Santa Cruz | Santa Cruz, Ichilo, Buenavista | Male | Male | Toe pad | 0.02 | 0.34 | 12.52 | 13,966 (partial) | A1 | PV918238 |

| | | | | | | | | | | | | | |
|---|---|---|---|---|---|---|---|---|---|---|---|---|---|
| FMNH179070 | *Ara severus* | Bolivia | Santa Cruz | Santa Cruz, Ichilo, Buenavista | Female | Female | Toe pad | 0.03 | 0.45 | 18.75 | NA | NA | NA |
| FMNH179071 | *Ara severus* | Bolivia | Santa Cruz | Nueva Moka | Female | Male | Toe pad | 0.05 | 0.37 | 36.55 | 16,996 | A1 | PV918240 |
| FMNH179072 | *Ara severus* | Bolivia | Santa Cruz | Cercado | Male | Male | Toe pad | 0.03 | 0.42 | 16.87 | 16,997 | A1 | PV918241 |
| FMNH257846 | *Ara severus* | Brazil | Para | Tapaiuna, Rio Tapajos | Female | Female | Toe pad | 0.51 | 0.68 | 159.50 | 16,995 | C | PV918211 |
| FMNH257847 | *Ara severus* | Brazil | Para | Tapaiuna, Rio Tapajos | Male | Male | Toe pad | 1.02 | 0.57 | 445.57 | 16,995 | C | PV918212 |
| FMNH257848 | *Ara severus* | Brazil | Para | Tapaiuna, Rio Tapajos | Male | Male | Toe pad | 0.01 | 0.52 | 4.24 | NA | NA | NA |
| FMNH257849 | *Ara severus* | Brazil | Para | Fordlandia, Rio Tapajos | Female | Female | Toe pad | 0.01 | 0.44 | 6.12 | NA | NA | NA |
| MVZ123802 | *Ara severus* | Colombia | Meta | Ocoa, 5 km S Villaviciencio | Female | Female | Toe pad | 0.96 | 0.61 | 351.82 | 16,995 | B1 | PV918259 |
| MVZ123803 | *Ara severus* | Colombia | Meta | Ocoa, 5 km S Villaviciencio | Male | Male | Toe pad | 0.02 | 0.49 | 13.72 | 14,261 (partial) | B2 | PV918247 |
| MVZ140645 | *Ara severus* | Colombia | Putumayo | Puerto Asis, Putumayo | Male | Male | Toe pad | 0.05 | 0.56 | 18.84 | 12,953 (partial) | B1 | PV918256 |
| FMNH261083 | *Ara severus* | Colombia | Arauca | Rio Arauca | Male | Male | Toe pad | 0.94 | 0.50 | 3688.07 | 16,996 | A2 | PV918218 |
| FMNH261084 | *Ara severus* | Colombia | Arauca | Rio Arauca | Female | Female | Toe pad | 0.05 | 0.52 | 66.02 | 16,996 | A2 | PV918220 |

| | | | | | | | | | | | | | |
|---|---|---|---|---|---|---|---|---|---|---|---|---|---|
| FMNH261085 | *Ara severus* | Colombia | Boyaca | Hacienda Bojabá | Male | Male | Toe pad | 0.95 | 0.47 | 971.99 | 16,996 | A2 | PV918219 |
| FMNH261086 | *Ara severus* | Colombia | Boyaca | Hacienda Bojabá | Male | Male | Toe pad | 0.02 | 0.44 | 16.69 | 15,558 (partial) | A2 | PV918216 |
| FMNH286761 | *Ara severus* | Colombia | Putumayo | San Antonio, Guamuez | Male | Male | Toe pad | 0.02 | 0.53 | 11.62 | 16,995 | B1 | PV918253 |
| FMNH286762 | *Ara severus* | Colombia | Putumayo | San Antonio Guamuez | Female | Male | Toe pad | 0.83 | 0.63 | 302.27 | 16,997 | A1 | PV918226 |
| FMNH286763 | *Ara severus* | Colombia | Putumayo | San Antonio Guamuez | Male | Male | Toe pad | 0.03 | 0.44 | 20.15 | 16,997 | B1 | PV918257 |
| FMNH286764 | *Ara severus* | Colombia | Putumayo | San Antonio Guamuez | Male | Female | Toe pad | 0.00 | 0.07 | 10.81 | 15,593 (partial) | B1 | PV918258 |
| FMNH292740 | *Ara severus* | Colombia | Putumayo | Estacion de Bombeo Guamuez | Male | Female | Toe pad | 0.01 | 0.16 | 39.32 | 16,995 | B1 | PV918249 |
| FMNH40259 | *Ara severus* | Colombia | | Rio Cauca | Female | Female | Toe pad | 8.84 | 0.62 | 3266.97 | 16,996 | B2 | PV918245 |
| FMNH50926 | *Ara severus* | Colombia | Cauca | Valle del Cauca, florida | Female | Male | Toe pad | 6.35 | 0.61 | 1814.67 | 16,996 | B2 | PV918243 |
| FMNH50927 | *Ara severus* | Colombia | Cundimarca | Cundimarca, El Roble, above Fusagasuga | Male | Male | Toe pad | 1.37 | 0.60 | 1419.78 | 16,995 | B2 | PV918244 |
| FMNH248517 | *Ara severus* | Colombia | Meta | Caño Yerly, Serrania de la Macarena | Female | Female | Toe pad | 0.96 | 0.54 | 285.18 | 16,996 | B2 | PV918248 |

| | | | | | | | | | | | | | |
|---|---|---|---|---|---|---|---|---|---|---|---|---|---|
| FMNH248518 | *Ara severus* | Colombia | Meta | Rio Guapaya, Serrania de la Macarena | Female | Female | Toe pad | 0.05 | 0.55 | 8.89 | NA | NA | NA |
| FMNH248519 | *Ara severus* | Colombia | Meta | Los Micos, San Juan de Arama | Male | Male | Toe pad | 0.02 | 0.46 | 18.28 | 16,997 | B1 | PV918252 |
| FMNH371815 | *Ara severus* | Ecuador | Los Rios | Centro Científico Río Palenque | Male | Male | Toe pad | 1.05 | 0.43 | 536.60 | 16,994 | B1 | PV918255 |
| FMNH77376 | *Ara severus* | Ecuador | Napo | Rio Suno | Male | Male | Toe pad | 0.89 | 0.27 | 1483.81 | 16,998 | A1 | PV918228 |
| FMNH119062 | *Ara severus* | Ecuador | Manabi | San Mateo | Male | Male | Toe pad | 0.62 | 0.55 | 170.28 | 16,995 | B1 | PV918254 |
| FMNH119063 | *Ara severus* | Ecuador | Manabi | San Mateo | Male | Male | Toe pad | 0.03 | 0.51 | 8.05 | NA | NA | NA |
| MVZ165114 | *Ara severus* | Peru | Amazonas | Rio Cenepa on bank, Huampami 0.5 mi W | Female | Female | Toe pad | 0.92 | 0.52 | 463.08 | 16,995 | A1 | PV918232 |
| MVZ169556 | *Ara severus* | Peru | Madre de Dios | Albergue, Cuzco Amazónico, ca. 12 km E (by air) Puerto Maldonado, Rio Madre de Dios, Peru -12.6 / -69.07289 | Male | Male | Toe pad | 1.57 | 0.67 | 589.57 | 16,998 | A1 | PV918235 |
| MUSM1985 | *Ara severus* | Peru | Huanuco | Rio Pachitea (altura de Tournavista y Puerto Inca), | Male | Male | Toe pad | 0.16 | 0.36 | 224.75 | 16,998 | A1 | PV918213 |

| | | | | | | | | | | | | | |
|---|---|---|---|---|---|---|---|---|---|---|---|---|---|
| | | | | Boca del Rio Macui | | | | | | | | | |
| MUSM1988 | *Ara severus* | Peru | Ucayali | Yarinacocha (Rio Ucayali) | Male | Male | Toe pad | 3.07 | 0.57 | 465.60 | 16,997 | A1 | PV918214 |
| MUSM1989 | *Ara severus* | Peru | Junin | La Merced Huallaga | | Female | Toe pad | 4.82 | 0.70 | 1026.42 | 16,996 | A1 | PV918233 |
| MUSM30585 | *Ara severus* | Peru | Amazonas | Puerto Galilea (Rio Santiago) | | Female | Toe pad | 3.17 | 0.64 | 377.78 | 16,996 | B2 | PV918246 |
| MUSM3991 | *Ara severus* | Peru | Amazonas | Puerto Galilea (Rio Santiago) | | Male | Toe pad | 2.30 | 0.72 | 420.36 | 16,996 | B1 | PV918251 |
| MUSM3993 | *Ara severus* | Peru | Amazonas | Puerto Galilea (Rio Santiago) | | Male | Toe pad | 2.99 | 0.53 | 1878.70 | 16,996 | A1 | PV918236 |
| FMNH251527 | *Ara severus* | Peru | Cuzco | Hda Villacarmen | Male | Male | Toe pad | 1.21 | 0.57 | 313.23 | 16,996 | A1 | PV918227 |
| FMNH251528 | *Ara severus* | Peru | Cuzco | Hda Villacarmen | Female | Female | Toe pad | 1.39 | 0.66 | 187.08 | 16,996 | A1 | PV918231 |
| FMNH251529 | *Ara severus* | Peru | Madre de Dios | Rio Inambari, boca | Female | Female | Toe pad | 0.89 | 0.60 | 922.13 | 16,997 | A1 | PV918237 |
| FMNH282553 | *Ara severus* | Peru | Amazonas | Puerto Galilea, Rio Santiago | | Male | Toe pad | 0.87 | 0.54 | 1004.37 | 16,997 | B1 | PV918250 |
| FMNH222872 | *Ara severus* | Peru | Madre de Dios | Boca Amigo | Male | Male | Toe pad | 0.90 | 0.55 | 196.83 | 16,997 | A1 | PV918234 |

| | | | | | | | | | | | | | |
|---|---|---|---|---|---|---|---|---|---|---|---|---|---|
| FMNH251526 | *Ara severus* | Peru | Cuzco | Hda Villacarmen | Male | Male | Toe pad | 1.16 | 0.69 | 240.65 | 16,996 | A1 | PV918230 |
| FMNH47325 | *Ara severus* | Suriname | Paramaribo | Paramaribo, vicinity | Male | Male | Toe pad | 2.84 | 0.55 | 807.75 | 16,995 | C | PV918209 |
| FMNH47326 | *Ara severus* | Suriname | Paramaribo | Paramaribo, vicinity | Female | Female | Toe pad | 2.71 | 0.56 | 540.56 | 16,995 | C | PV918210 |
| FMNH34497 | *Ara severus* | Venezuela | Zulia | Encontrados | Male | Male | Toe pad | 4.83 | 0.40 | 786.52 | 16,996 | A2 | PV918221 |
| FMNH34498 | *Ara severus* | Venezuela | Zulia | Encontrados | Female | Female | Toe pad | 5.88 | 0.45 | 3193.50 | 16,996 | A2 | PV918217 |
| FMNH34499 | *Ara severus* | Venezuela | Zulia | Encontrados | Female | Female | Toe pad | 3.07 | 0.56 | 617.73 | 16,995 | A2 | PV918225 |
| FMNH34500 | *Ara severus* | Venezuela | Zulia | Encontrados | Female | Female | Toe pad | 5.23 | 0.45 | 2112.02 | 16,993 | A2 | PV918222 |
| FMNH34501 | *Ara severus* | Venezuela | Zulia | Encontrados | Male | Male | Toe pad | 4.75 | 0.58 | 1967.47 | 16,993 | A2 | PV918224 |
| FMNH34502 | *Ara severus* | Venezuela | Zulia | Encontrados | Male | Male | Toe pad | 6.31 | 0.58 | 2410.54 | 16,996 | A2 | PV918215 |
| FMNH43295 | *Ara severus* | Venezuela | Zulia | Zulia, Catatumbo | Female | Female | Toe pad | 6.53 | 0.51 | 2970.72 | 16,996 | A2 | PV918223 |
| FMNH46978 | *Ara severus* | Venezuela | Zulia | Encontrados | Male | Male | Toe pad | 0.03 | 0.54 | 5.68 | NA | NA | NA |

**Table S4.** Intra- and inter-clade genetic distances among mitogenome-defined clades in *Ara ararauna*. Distances were estimated using the Species Delimitation plugin in Geneious (Masters, Fan, and Ross 2011) based on the inferred maximum-likelihood phylogeny. Values represent mean pairwise genetic distances within (intra) and between (inter) clades, based on tree topology.

| ***Ara ararauna*** | | | |
|---|---|---|---|
| Clade | Nearest Neighbor | Intra Dist | Inter Dist |
| A | C | 0.002 | 0.007 |
| A1 | A2 | 0.001 | 0.002 |
| A2 | A1 | 0.001 | 0.002 |
| A3 | A2 | 0.001 | 0.003 |
| B | C | 0.003 | 0.005 |
| B1 | B3 | 5.96E-4 | 0.005 |
| B2 | B3 | 0.001 | 0.003 |
| B3 | B2 | 0.001 | 0.003 |
| C | B | 8.78E-4 | 0.005 |

**Table S5.** Intra- and inter-clade genetic distances among mitogenome-defined clades in *Ara chloropterus*. Distances were estimated using the Species Delimitation plugin in Geneious (Masters, Fan, and Ross 2011) based on the inferred maximum-likelihood phylogeny. Values represent mean pairwise genetic distances within (intra) and between (inter) clades, based on tree topology.

| ***Ara chloropterus*** | | | |
|---|---|---|---|
| Clade | Nearest Neighbor | Intra Dist | Inter Dist |
| A | C | 0.001 | 0.003 |
| B | C | 0.003 | 0.003 |
| B1 | B2 | 0.002 | 0.003 |
| B2 | C | 0.002 | 0.003 |
| C | A | 0.002 | 0.003 |
| D | C | 7.94E-4 | 0.004 |
| E | D | 6.39E-4 | 0.006 |

**Table S6.** Intra- and inter-clade genetic distances among mitogenome-defined clades in *Ara severus*. Distances were estimated using the Species Delimitation plugin in Geneious (Masters, Fan, and Ross 2011) based on the inferred maximum-likelihood phylogeny. Values represent mean pairwise genetic distances within (intra) and between (inter) clades, based on tree topology.

| ***A. severus*** | | | |
|---|---|---|---|
| Clade | Nearest Neighbor | Intra Dist | Inter Dist |
| A | B | 0.002 | 0.004 |
| A1 | A2 | 0.001 | 0.002 |
| A2 | A1 | 8.77E-4 | 0.002 |
| B | A | 0.002 | 0.004 |
| B1 | B2 | 0.001 | 0.002 |
| B2 | B1 | 0.002 | 0.002 |
| C | B | 9.31E-4 | 0.008 |

**Table S7.** Summary statistics of autosomal heterozygosity (H) for three *Ara* species and their respective populations. For each species and population, the number of individuals (N), mean heterozygosity, standard deviation (sd), and range (min–max) are reported. Species-level values of *A. ararauna* and *A. chloropterus* include only the three main populations. “Isolated” and “Captive” are referring to individuals outside major biomes and populations.

| Species | Kruskal-Wallis test | Biome | N | mean heterozygosity | median | SD | min | max |
|---|---|---|---|---|---|---|---|---|
| *A. ararauna* | 2.2e-16 | All | 62 | 0.00102220968466054 | 0.000847282620094763 | 0.000371253047546309 | 0.000559045722566256 | 0.0022709980924101 |
| *A. chloroptera* | | All | 55 | 0.000723628159197142 | 0.000499690220959666 | 0.000403805171309007 | 0.000227714017149364 | 0.00200406092688419 |
| *A. severus* | | All | 38 | 0.00170767253433377 | 0.00167514831731022 | 0.000350497611198296 | 0.00112044449060496 | 0.00277697917671987 |
| *A. ararauna* | 0.00029 | West Amazon | 24 | 0.00121741212312722 | 0.00124593511305818 | 0.000378935495366211 | 0.000595243454200956 | 0.00182546968609571 |
| *A. ararauna* | | East Amazon | 9 | 0.00124438589697938 | 0.0012091608414555 | 0.000481099702361587 | 0.000652735056961437 | 0.0022709980924101 |
| *A. ararauna* | | Cerrado | 30 | 0.000785673188773603 | 0.000778636981041553 | 8.87145554274078e-05 | 0.000559045722566256 | 0.00103082680266404 |
| *A. ararauna* | | Isolated | 1 | 0.00162588118083761 | 0.00162588118083761 | NA | NA | NA |
| *A. ararauna* | | Central | 1 | 0.00083018866214883 | 0.00083018866214883 | NA | NA | NA |
| *A. chloroptera* | 0.00013 | West Amazon | 24 | 0.000912712566718414 | 0.000917479043451757 | 0.000391985162157897 | 0.000336292662134106 | 0.00200406092688419 |
| *A. chloroptera* | | East Amazon | 13 | 0.00079655368103035 | 0.00066038280 7087321 | 0.000430058864588031 | 0.000347479558951232 | 0.0016513039133029 |
| *A. chloroptera* | | Cerrado | 18 | 0.000411236874248008 | 0.00041108427 7658427 | 5.29052269712447e-05 | 0.000334361520837479 | 0.000499690220959666 |
| *A.* | | Cerra | 6 | 0.000411 | 0.000415 | 0.0000529 | 0.000334 | 0.000472 |

| | | | | | | | | |
|---|---|---|---|---|---|---|---|---|
| *chloroptera* | | do North | | | | | | |
| *A. chloroptera* | | Cerrado South | 12 | 0.000411 | 0.000406 | 0.0000552 | 0.000341 | 0.000500 |
| *A. chloroptera* | | Isolated | 1 | 0.00135652786598712 | 0.00135652786598712 | NA | NA | NA |
| *A. chloroptera* | | Captive | 1 | 0.000227714017149364 | 0.000227714017149364 | NA | NA | NA |

**Table S8.** Global $F_{ST}$ values (weighted) estimated among populations in *A. ararauna*

| **Population 1** | **Population 2** | **Fst (Unweight)** | **Fst (Weight)** | **nObs** |
|---|---|---|---|---|
| Amazon | Cerrado | 0.019927 | 0.047230 | 499612007 |
| Amazon_West | Amazon_East | 0.035992 | 0.047007 | 438415081 |
| Amazon_West | Cerrado | 0.031323 | 0.050594 | 472297970 |
| Amazon_East | Cerrado | 0.090302 | 0.069760 | 863994275 |

**Table S9.** Global $F_{ST}$ values (weighted) estimated among populations in *A. chloropterus*

| **Population 1** | **Population 2** | **Fst (Unweight)** | **Fst (Weight)** | **nObs** |
|---|---|---|---|---|
| Amazon | Cerrado | 0.009540 | 0.048250 | 760698655 |
| Amazon_West | Amazon_East | 0.027780 | 0.044086 | 604378758 |
| Cerrado_N | Cerrado_S | 0.039462 | 0.102744 | 908849307 |
| Cerrado_N | Amazon_West | 0.014663 | 0.121077 | 589442985 |
| Cerrado_N | Amazon_East | 0.019226 | 0.059689 | 880627914 |
| Cerrado_S | Amazon_West | 0.018270 | 0.076637 | 608023766 |
| Cerrado_S | Amazon_East | 0.030061 | 0.067384 | 912063601 |

**Table S10.** Model selection (under BIC) and best partition scheme in *A. ararauna* inferred by IQ-TREE v2.0.3 (Nguyen et al. 2015). The phylogenetic-tree construction was performed using the following command: iqtree -s alignment.phy -p partition.nex -m MFP+MERGE -B 1000 -nt AUTO. Initially the mitogenome was partitioned by gene and codon, resulting in 67 schemes (partition.nex). This scheme was tested, partitions were merged into four groups, and the best model for those groups were inferred.

| **Partition** | **Genes** | **Model** |
|---|---|---|
| 1 | tRNA-Phe, 12S, tRNA-Val, 16S, tRNA-leu, ND1-1, tRNA-Ile, tRNA-Met, ND2-1, tRNA-Trp, tRNA-Ala, tRNA-Asn, tRNA-Cys, tRNA-Tyr, COX1-1, tRNA-Ser, tRNA-Asp, COX2-1, tRNA-Lys, COX3-1, tRNA-Gly, ND3a-1, ND3b-1, tRNA-Arg, ND4L-1, ND4-1, tRNA-Ser2, ND5-1, Cytb-1, tRNA-Pro, ND6-2, tRNA-Glu | HKY+F+R2 |
| 2 | ND1-2, ND2-2, COX1-2, COX2-2, ATP8-2, ATP6-2, ND3a-2, ND3b-2, ND4L-2, ND4-2, ND5-2, Cytb-2 | HKY+F+I |
| 3 | ND1-3, ND2-3, COX1-3, COX2-3, ATP8-1, ATP8-3, ATP6-3, COX3-3, ND3a-3, ND3b-3, ND4L-3, ND4-3, ND5-3, Cytb-3, ND6-3 | TN+F+G4 |
| 4 | tRNA-Gln, ATP6-1, COX3-2, tRNA-His, tRNA-Leu, tRNA-Thr, ND6-1, D-loop | HKY+F+R2 |

**Table S11.** Model selection (under BIC) and best partition scheme in *A. chloropterus* inferred by IQ-TREE v2.0.3 (Nguyen et al. 2015). The phylogenetic-tree construction was performed using the following command: iqtree -s alignment.phy -p partition.nex -m MFP+MERGE -B 1000 -nt AUTO. Initially the mitogenome was partitioned by gene and codon, resulting in 67 schemes (partition.nex). This scheme was tested, partitions were merged into four groups, and the best model for those groups were inferred.

| **Partition** | **Genes** | **Model** |
|---|---|---|
| 1 | tRNA-Phe, 12S, tRNA-Val, 16S, tRNA-leu, ND1-1, tRNA-Met, ND2-1, tRNA-Asn, tRNA-Cys, tRNA-Tyr, COX1-1, tRNA-Asp, COX2-1, tRNA-Lys, ATP8-2, ATP6-1, tRNA-Gly, ND3b-1, tRNA-Arg, ND4L-1, ND4-1, tRNA-His, tRNA-Leu, ND5-1, Cytb-1, tRNA-Pro, tRNA-Glu | HKY+F+I |
| 2 | ND1-2, ND2-2, COX1-2, COX2-2, COX3-2, ND3a-1, ND3a-2, ND3b-2, ND4L-2, ND4-2, ND5-2, Cytb-2 | HKY+F+I |
| 3 | ND1-3, ND2-3, COX1-3, COX2-3, ATP8-3, ATP6-3, COX3-3, ND3a-3, ND3b-3, ND4L-3, ND4-3, ND5-3, Cytb-3, ND6-3, ND6-2, ND6-1 | TN+F |
| 4 | tRNA-Ile, tRNA-Gln, tRNA-Trp, tRNA-Ala, tRNA-Ser, ATP8-1, ATP6-2, COX3-1, tRNA-Ser2, tRNA-Thr, D-loop | HKY+F+R2 |

**Table S12.** Model selection (under BIC) and best partition scheme in *A. severus* inferred by IQ-TREE v2.0.3 (Nguyen et al. 2015). The phylogenetic-tree construction was performed using the following command: iqtree -s alignment.phy -p partition.nex -m MFP+MERGE -B 1000 -nt AUTO. Initially the mitogenome was partitioned by gene and codon, resulting in 67 schemes (partition.nex). This scheme was tested, partitions were merged into four groups, and the best model for those groups were inferred.

| Partition | Genes | Model |
|---|---|---|
| 1 | tRNA-Phe, 12S, 16S, tRNA-leu, ND1-1, tRNA-Met, ND2-1, tRNA-Trp, tRNA-Ala, tRNA-Asn, tRNA-Cys, tRNA-Tyr, COX1-1, COX2-1, ATP8-1, ATP6-1, COX3-1, tRNA-Gly, ND3a-1, ND3b-1, tRNA-Arg, ND4L-1, ND4-1, tRNA-His, tRNA-Ser2, tRNA-Leu, ND5-2, Cytb-1, tRNA-Thr, tRNA-Pro, ND6-2, tRNA-Glu | HKY+F+I |
| 2 | tRNA-Val, tRNA-Ile, ND2-2, tRNA-Ser, tRNA-Asp, tRNA-Lys, ATP8-2, D-loop | HKY+F+R2 |
| 3 | ND1-2, tRNA-Gln, COX1-2, COX2-2, ATP6-2, COX3-2, ND3a-2, ND3b-2, ND4L-2, ND4-2, ND5-3, Cytb-2 | HKY+F+I |
| 4 | ND1-3, ND2-3, COX1-3, COX2-3, ATP8-3, ATP6-3, COX3-3, ND3a-3, ND3b-3, ND4L-3, ND4-3, ND5-1, Cytb-3, ND6-3, ND6-1 | TIM2+F+G4 |

**Table S13.** Summary statistics of individual inbreeding coefficients (F) estimated with ngsF for three *Ara* species across populations. For each species–population group, the number of individuals (N), mean, standard deviation (sd), and range (min–max) of F are reported. "ALL" refers to the combined set of individuals per species. Species-level values of *A. ararauna* and *A. chloropterus* include only the three main populations.

| species | Population | N | mean inbreeding (F) | sd | min | max |
|---|---|---|---|---|---|---|
| *Ara ararauna* | All | 62 | 0.011326064516129 | 0.0173209321407734 | 0 | 0.107779 |
| *Ara chloropterus* | All | 55 | 0.019947 | 0.0435414026730966 | 0 | 0.275289 |
| *Ara severus* | All | 38 | 0.00624860526315789 | 0.0107657697771235 | 0.00029 | 0.05719 |
| *Ara ararauna* | WestAmazon | 23 | 0.00655 | 0.0221977083317584 | 0.000273 | 0.107779 |
| *Ara ararauna* | EastAmazon | 9 | 0.0105164444444444 | 0.0195230400367816 | 0 | 0.046407 |
| *Ara ararauna* | Cerrado | 30 | 0.0152306 | 0.0108652439349293 | 0.002291 | 0.042812 |
| *Ara chloropterus* | WestAmazon | 24 | 0.00636070833333333 | 0.0177145701207623 | 3.8e-05 | 0.084248 |
| *Ara chloropterus* | EastAmazon | 13 | 0.0275493846153846 | 0.075162379458896 | 0 | 0.275289 |
| *Ara chloropterus* | Cerrado | 18 | 0.0325714444444444 | 0.0337078619546769 | 0.003623 | 0.120303 |
| *Ara chloropterus* | Cerrado North | 6 | 0.05012150 | 0.05254713 | 0.003623 | 0.120303 |
| *Ara chloropterus* | Cerrado South | 12 | 0.02379642 | 0.01577597 | 0.005343 | 0.058115 |

**Table S14.** Comparison of Fay & Wu's H in $F_{ST}$ outliers vs non-outlier windows. $F_{ST}$ outlier windows were defined as the top 1% of the empirical $F_{ST}$ distribution within each species and population comparison, after filtering windows with fewer than 100 callable sites and excluding the Z chromosome. For each species, comparison, and biome, Fay & Wu's H was compared between $F_{ST}$ outlier and non-outlier windows using a one-sided Wilcoxon rank-sum test, testing whether $F_{ST}$ outlier windows had lower H values than non-outlier windows. P-values were adjusted across tests using the Benjamini–Hochberg correction, asterisks show significant results.

| **Species** | **Comparison** | **Number of FST outlier windows** | **Number of non FST outlier windows** | **Number of genes in FST outlier windows** | **Biome** | **median H in FST outlier windows** | **median H in FST non-outlier windows** | **mean H in FST outlier windows** | **mean H in FST non-outlier windows** | **Wilcoxon rank-sum W** | **p_value** | **p_adj_BH** | **direction_median** | **direction_mean** |
|---|---|---|---|---|---|---|---|---|---|---|---|---|---|---|
| *Ara ararauna* | Cerrado vs East_Amazon | 1039 | 102842 | 168 | Cerrado | -0.205275 | -0.115623 | -0.3126608662175168 | -0.16355157197448514 | 43490260 | 2.549020669711413e-25 | 6.797388452563768e-25 * | FST_outlier_lower_H | FST_outlier_lower_H |
| *Ara ararauna* | Cerrado vs East_Amazon | 1039 | 102842 | 168 | East_Amazon | -0.404684 | -0.1704625000000002 | -0.42889412030798846 | -0.22342237537192974 | 31319996 | 3.286995395001122e-117 | 2.6295963160008974e-116 * | FST_outlier_lower_H | FST_outlier_lower_H |
| *Ara ararauna* | Cerrado vs West_Amazon | 1032 | 102103 | 259 | Cerrado | -0.141328 | -0.11574 | -0.19055919089147286 | -0.16336706423905273 | 51361255.5 | 0.08208755519847456 | 0.10945007359796607 | FST_outlier_lower_H | FST_outlier_lower_H |
| *Ara ararauna* | Cerrado vs West_Amazon | 1032 | 102092 | 259 | West_Amazon | -0.3643645 | -0.2278065000000002 | -0.42655820833333336 | -0.2959811430278572 | 41815691 | 1.7185073358819506e-30 | 6.874029343527802e-30 * | FST_outlier_lower_H | FST_outlier_lower_H |
| *Ara chloropterus* | Cerrado vs East_Amazon | 1046 | 103531 | 211 | Cerrado | -0.2749195 | -0.281899 | -0.40907754684512426 | -0.33288028521891994 | 53025585.5 | 0.1242400548919938 | 0.1419886341622786 | FST_outlier_higher_H | FST_outlier_lower_H |
| *Ara* | Cerrad | 104 | 103 | 211 | East | -0.5 | -0.479 | -0.6630 | -0.5317 | 466 | 6.22658 | 9.96253 | FST_o | FST_o |

| | | | | | | | | | | | | | | |
|---|---|---|---|---|---|---|---|---|---|---|---|---|---|---|
| *chloropterus* | o vs East_Amazon | 6 | 531 | | _Amazon | 839945 | 948 | 885305927342 | 748391592856 | 55384.5 | 5240823462e-15 | 638531754e-15 * | utlier_lower_H | utlier_lower_H |
| *Ara chloropterus* | Cerrado vs West_Amazon | 1042 | 103068 | 269 | Cerrado | -0.230842 | -0.2834105 | -0.3317992226487524 | -0.3352155681491831 | 56669014.5 | 0.998956046113 1106 | 0.998956046113 1106 | FST_outlier_higher_H | FST_outlier_higher_H |
| *Ara chloropterus* | Cerrado vs West_Amazon | 1042 | 103066 | 269 | West_Amazon | -0.938701 | -0.787503 | -0.9978167994241842 | -0.8395303357654319 | 45502995.5 | 1.039485979349 1519e-17 | 2.078971958698 3038e-17 * | FST_outlier_lower_H | FST_outlier_lower_H |

**Table S15** Total candidate gene list. These genes overlap with $F_{ST}$ outlier windows per comparison per species, which also exhibit low Fay & Wu's H. Because $F_{ST}$ was estimated in genomic windows rather than directly per gene, $F_{ST}$ values are summarized from all $F_{ST}$ windows overlapping each gene. The table reports the number of overlapping $F_{ST}$ windows, the number of overlapping $F_{ST}$ outlier windows, the maximum and mean $F_{ST}$ across overlapping windows, the maximum $F_{ST}$ among outlier windows, and the pairwise comparisons in which each gene overlapped $F_{ST}$ signal.

| Species | Biome | Gene Label | ID | gene_chr | gene_start | gene_end | GeneLabel_gff | n_FST_windows_overlapping_gene | n_FST_outlier_windows_overlapping_gene | max_Fst_overlapping_gene | mean_Fst_overlapping_gene | max_Fst_outlier_gene | FST_comparisons_with_overlap | FST_comparisons_with_outlier |
|---|---|---|---|---|---|---|---|---|---|---|---|---|---|---|
| ararauna | Cerrado | SEC31A | FUNCG00000000608 | CM054376.1 | 63279966 | 63321373 | SEC31A | 20 | 3 | 0.223744 | 0.10233335 | 0.223744 | Cerrado_vs_East_Amazon;Cerrado_vs_West_Amazon | Cerrado_vs_East_Amazon |
| ararauna | Cerrado | ST18 | FUNCG00000002285 | CM054377.1 | 42025789 | 42214896 | ST18 | 48 | 5 | 0.2766 | 0.08080285416666666 | 0.2766 | Cerrado_vs_East_Amazon;Cerrado_vs_West_Amazon | Cerrado_vs_East_Amazon |
| ararauna | Cerrado | FUNCG00000002304 | FUNCG00000002304 | CM054377.1 | 45240241 | 45305569 | FUNCG000000002304 | 22 | 2 | 0.213565 | 0.09046709090909091 | 0.213565 | Cerrado_vs_East_Amazon;Cerrado_vs_West_Amazon | Cerrado_vs_East_Amazon |
| ararauna | Cerrado | DNAH5 | FUNCG0000 | CM054 | 645313 | 647010 | DNAH5 | 44 | 2 | 0.2040 | 0.07127097 | 0.2040 | Cerrado_vs_East_Amazon;C | Cerrado_vs_East_Amazon |

| | | | | | | | | | | | | | | |
|---|---|---|---|---|---|---|---|---|---|---|---|---|---|---|
| a | o | | 0002462 | 377.1 | 06 | 87 | | | | 37 | 727272727 | 37 | errado_vs_West_Amazon | |
| arauna | Cerrado | LTN1 | FUNCG000000003637 | CM054378.1 | 31889289 | 31918944 | LTN1 | 16 | 2 | 0.235275 | 0.0992710625 | 0.235275 | Cerrado_vs_East_Amazon;Cerrado_vs_West_Amazon | Cerrado_vs_East_Amazon |
| arauna | Cerrado | RWDD2B | FUNCG000000003638 | CM054378.1 | 31920537 | 31924055 | RWDD2B | 10 | 3 | 0.235275 | 0.1212899 | 0.235275 | Cerrado_vs_East_Amazon;Cerrado_vs_West_Amazon | Cerrado_vs_East_Amazon |
| arauna | Cerrado | USP16 | FUNCG000000003639 | CM054378.1 | 31926729 | 31946899 | USP16 | 14 | 5 | 0.259631 | 0.13330857142857144 | 0.259631 | Cerrado_vs_East_Amazon;Cerrado_vs_West_Amazon | Cerrado_vs_East_Amazon |
| arauna | Cerrado | CCT8 | FUNCG000000003640 | CM054378.1 | 31949326 | 31961285 | CCT8 | 14 | 6 | 0.259631 | 0.1436072142857143 | 0.259631 | Cerrado_vs_East_Amazon;Cerrado_vs_West_Amazon | Cerrado_vs_East_Amazon |
| arauna | Cerrado | GYG2 | FUNCG000000003872 | CM054378.1 | 58667087 | 58707591 | GYG2 | 18 | 2 | 0.208787 | 0.07101166666666667 | 0.208787 | Cerrado_vs_East_Amazon;Cerrado_vs_West_Amazon | Cerrado_vs_East_Amazon |
| arauna | Cerrado | FUNCG000000003873 | FUNCG000000003873 | CM054378.1 | 58713730 | 58743300 | FUNCG000000003873 | 16 | 3 | 0.209233 | 0.0951805625 | 0.209233 | Cerrado_vs_East_Amazon;Cerrado_vs_West_Amazon | Cerrado_vs_East_Amazon |
| arauna | Cerrado | BIVM | FUNCG000000004044 | CM054378.1 | 76643996 | 76656572 | BIVM | 12 | 1 | 0.179106 | 0.099582 | 0.131415 | Cerrado_vs_East_Amazon;Cerrado_vs_West_Amazon | Cerrado_vs_West_Amazon |
| arauna | Cerrado | POGLUT2 | FUNCG000000004045 | CM054378.1 | 76661382 | 76675815 | POGLUT2 | 12 | 4 | 0.198948 | 0.1368024166666668 | 0.198948 | Cerrado_vs_East_Amazon;Cerrado_vs_West_Amazon | Cerrado_vs_East_Amazon;Cerrado_vs_West_Amazon |
| araraun | Cerrad | TEX30 | FUNCG0000 | CM054 | 766778 | 766840 | TEX30 | 12 | 5 | 0.1989 | 0.14503008 | 0.1989 | Cerrado_vs_East_Amazon;C | Cerrado_vs_East_Amazon;Cerra |

| | | | | | | | | | | | | | | |
|---|---|---|---|---|---|---|---|---|---|---|---|---|---|---|
| a | o | | 0004046 | 378.1 | 71 | 80 | | | | 48 | 333333334 | 48 | errado_vs_West_Amazon | do_vs_West_Amazon |
| ararauna | Cerrado | METTL21C | FUNCG00000004047 | CM054378.1 | 76710023 | 76713756 | METTL21C | 10 | 4 | 0.170569 | 0.131961 | 0.128055 | Cerrado_vs_East_Amazon;Cerrado_vs_West_Amazon | Cerrado_vs_West_Amazon |
| ararauna | Cerrado | Tpp2 | FUNCG00000004048 | CM054378.1 | 76714962 | 76766928 | Tpp2 | 20 | 4 | 0.170569 | 0.1109193 | 0.128055 | Cerrado_vs_East_Amazon;Cerrado_vs_West_Amazon | Cerrado_vs_West_Amazon |
| ararauna | Cerrado | NALCN | FUNCG00000004051 | CM054378.1 | 77411582 | 77644676 | NALCN | 56 | 5 | 0.272505 | 0.118105375 | 0.272505 | Cerrado_vs_East_Amazon;Cerrado_vs_West_Amazon | Cerrado_vs_East_Amazon |
| ararauna | Cerrado | CLYBL | FUNCG00000004060 | CM054378.1 | 78280851 | 78415799 | CLYBL | 36 | 12 | 0.380639 | 0.1502885555555556 | 0.380639 | Cerrado_vs_East_Amazon;Cerrado_vs_West_Amazon | Cerrado_vs_East_Amazon;Cerrado_vs_West_Amazon |
| ararauna | Cerrado | TM9SF2 | FUNCG00000004061 | CM054378.1 | 78468726 | 78516772 | TM9SF2 | 20 | 5 | 0.270757 | 0.12197865 | 0.270757 | Cerrado_vs_East_Amazon;Cerrado_vs_West_Amazon | Cerrado_vs_East_Amazon |
| ararauna | Cerrado | GPC6 | FUNCG00000004085 | CM054378.1 | 80685760 | 80881370 | GPC6 | 50 | 10 | 0.209895 | 0.07812266 | 0.209895 | Cerrado_vs_East_Amazon;Cerrado_vs_West_Amazon | Cerrado_vs_West_Amazon |
| ararauna | Cerrado | Eif5 | FUNCG00000006955 | CM054380.1 | 69951101 | 69955623 | Eif5 | 10 | 3 | 0.156295 | 0.1382778 | 0.152384 | Cerrado_vs_East_Amazon;Cerrado_vs_West_Amazon | Cerrado_vs_West_Amazon |
| ararauna | Cerrado | AHNAK2 | FUNCG00000006982 | CM054380.1 | 71834603 | 71905756 | AHNAK2 | 24 | 5 | 0.259212 | 0.124626083333333333 | 0.259212 | Cerrado_vs_East_Amazon;Cerrado_vs_West_Amazon | Cerrado_vs_East_Amazon |
| ararauna | Cerrad | QRICH2 | FUNCG0000 | CM054 | 719112 | 719175 | QRICH2 | 10 | 5 | 0.2592 | 0.1462121 | 0.2592 | Cerrado_vs_East_Amazon;C | Cerrado_vs_East_Amazon |

| | | | | | | | | | | | | | | |
|---|---|---|---|---|---|---|---|---|---|---|---|---|---|---|
| a | o | | 0006983 | 380.1 | 36 | 31 | | | | 12 | | 12 | errado_vs_West_Amazon | |
| ararauna | Cerrado | FUNCG000000007069 | FUNCG000000007069 | CM054380.1 | 79858815 | 79891933 | FUNCG000000007069 | 18 | 3 | 0.235844 | 0.0686358888888889 | 0.235844 | Cerrado_vs_East_Amazon;Cerrado_vs_West_Amazon | Cerrado_vs_West_Amazon |
| ararauna | Cerrado | PLAU | FUNCG000000007809 | CM054381.1 | 31122598 | 31129070 | PLAU | 10 | 1 | 0.135973 | 0.0681728 | 0.135973 | Cerrado_vs_East_Amazon;Cerrado_vs_West_Amazon | Cerrado_vs_West_Amazon |
| ararauna | Cerrado | Sorbs1 | FUNCG000000008203 | CM054381.1 | 67786738 | 67861694 | Sorbs1 | 26 | 2 | 0.202286 | 0.09095203846153846 | 0.202286 | Cerrado_vs_East_Amazon;Cerrado_vs_West_Amazon | Cerrado_vs_East_Amazon |
| ararauna | Cerrado | Antxr1 | FUNCG000000008248 | CM054381.1 | 69709274 | 69763607 | Antxr1 | 22 | 4 | 0.162677 | 0.08749663636363636 | 0.162677 | Cerrado_vs_East_Amazon;Cerrado_vs_West_Amazon | Cerrado_vs_West_Amazon |
| ararauna | Cerrado | FAS | FUNCG000000008271 | CM054381.1 | 71234923 | 71246693 | FAS | 12 | 2 | 0.213818 | 0.1023113333333334 | 0.213818 | Cerrado_vs_East_Amazon;Cerrado_vs_West_Amazon | Cerrado_vs_East_Amazon |
| ararauna | Cerrado | RYK | FUNCG000000008719 | CM054382.1 | 13841079 | 13893777 | RYK | 20 | 6 | 0.336577 | 0.13743905 | 0.336577 | Cerrado_vs_East_Amazon;Cerrado_vs_West_Amazon | Cerrado_vs_East_Amazon |
| ararauna | Cerrado | EPHA4 | FUNCG000000008774 | CM054382.1 | 17942568 | 18044780 | EPHA4 | 30 | 2 | 0.147464 | 0.08507706666666667 | 0.147464 | Cerrado_vs_East_Amazon;Cerrado_vs_West_Amazon | Cerrado_vs_West_Amazon |
| ararauna | Cerrado | EIF4A2 | FUNCG000000008985 | CM054382.1 | 35466670 | 35473357 | EIF4A2 | 12 | 5 | 0.236048 | 0.1285885 | 0.236048 | Cerrado_vs_East_Amazon;Cerrado_vs_West_Amazon | Cerrado_vs_East_Amazon |
| ararauna | Cerrad | KNG1 | FUNCG0000 | CM054 | 354902 | 355093 | KNG1 | 12 | 3 | 0.2113 | 0.11354058 | 0.2113 | Cerrado_vs_East_Amazon;C | Cerrado_vs_East_Amazon |

| | | | | | | | | | | | | | | |
|---|---|---|---|---|---|---|---|---|---|---|---|---|---|---|
| a | o | | 0008986 | 382.1 | 59 | 41 | | | | 98 | 333333333 | 98 | errado_vs_West_Amazon | |
| ararauna | Cerrado | FETUB | FUNCG000000008987 | CM054382.1 | 35517816 | 35549945 | FETUB | 16 | 2 | 0.211398 | 0.0877150625 | 0.211398 | Cerrado_vs_East_Amazon;Cerrado_vs_West_Amazon | Cerrado_vs_East_Amazon |
| ararauna | Cerrado | CKAP5 | FUNCG000000009052 | CM054382.1 | 37353579 | 37357484 | CKAP5 | 10 | 2 | 0.221557 | 0.1030465 | 0.221557 | Cerrado_vs_East_Amazon;Cerrado_vs_West_Amazon | Cerrado_vs_East_Amazon |
| ararauna | Cerrado | SST | FUNCG000000009053 | CM054382.1 | 37363591 | 37364953 | SST | 10 | 3 | 0.221557 | 0.11793100000000001 | 0.221557 | Cerrado_vs_East_Amazon;Cerrado_vs_West_Amazon | Cerrado_vs_East_Amazon |
| ararauna | Cerrado | TP63 | FUNCG000000009057 | CM054382.1 | 38124980 | 38227946 | TP63 | 30 | 4 | 0.155826 | 0.1051478 | 0.140606 | Cerrado_vs_East_Amazon;Cerrado_vs_West_Amazon | Cerrado_vs_West_Amazon |
| ararauna | Cerrado | BRINP2 | FUNCG000000009109 | CM054382.1 | 44390717 | 44396341 | BRINP2 | 10 | 1 | 0.201259 | 0.1425291 | 0.201259 | Cerrado_vs_East_Amazon;Cerrado_vs_West_Amazon | Cerrado_vs_East_Amazon |
| ararauna | Cerrado | RBBP6 | FUNCG000000014155 | CM054395.1 | 91243 | 105918 | RBBP6 | 4 | 2 | 0.217551 | 0.19002249999999998 | 0.217551 | Cerrado_vs_East_Amazon | Cerrado_vs_East_Amazon |
| ararauna | East_Amazon | POGLUT2 | FUNCG000000004045 | CM054378.1 | 76661382 | 76675815 | POGLUT2 | 6 | 1 | 0.198948 | 0.1569215 | 0.198948 | Cerrado_vs_East_Amazon | Cerrado_vs_East_Amazon |
| ararauna | East_Amazon | TEX30 | FUNCG000000004046 | CM054378.1 | 76677871 | 76684080 | TEX30 | 6 | 1 | 0.198948 | 0.166544 | 0.198948 | Cerrado_vs_East_Amazon | Cerrado_vs_East_Amazon |
| araraun | East_A | Tagap | FUNCG0000 | CM054 | 838617 | 838639 | Tagap | 5 | 5 | 0.2896 | 0.281976 | 0.2896 | Cerrado_vs_East_Amazon | Cerrado_vs_East_Amazon |

| | | | | | | | | | | | | | | |
|---|---|---|---|---|---|---|---|---|---|---|---|---|---|---|
| a | mazon | | 0004099 | 378.1 | 74 | 26 | | | | 54 | | 54 | | |
| ararauna | East_Amazon | IPMK | FUNCG000000007968 | CM054381.1 | 45328518 | 45374605 | IPMK | 10 | 1 | 0.217145 | 0.131844 | 0.217145 | Cerrado_vs_East_Amazon | Cerrado_vs_East_Amazon |
| ararauna | East_Amazon | PPP2R3A | FUNCG000000008749 | CM054382.1 | 15250851 | 15294942 | PPP2R3A | 9 | 3 | 0.233547 | 0.13292088888889 | 0.233547 | Cerrado_vs_East_Amazon | Cerrado_vs_East_Amazon |
| ararauna | East_Amazon | MSL2 | FUNCG000000008750 | CM054382.1 | 15303425 | 15319538 | MSL2 | 6 | 4 | 0.233547 | 0.1996133333333334 | 0.233547 | Cerrado_vs_East_Amazon | Cerrado_vs_East_Amazon |
| ararauna | East_Amazon | LPP | FUNCG000000009054 | CM054382.1 | 37689749 | 37899840 | LPP | 26 | 26 | 0.584017 | 0.4508361153846154 | 0.584017 | Cerrado_vs_East_Amazon | Cerrado_vs_East_Amazon |
| ararauna | East_Amazon | OR14J1 | FUNCG000000015127 | CM054400.1 | 901277 | 911814 | OR14J1 | 6 | 2 | 0.286093 | 0.140838 | 0.286093 | Cerrado_vs_East_Amazon | Cerrado_vs_East_Amazon |
| ararauna | East_Amazon | Pim1 | FUNCG000000015128 | CM054400.1 | 917647 | 919835 | Pim1 | 5 | 2 | 0.286093 | 0.1580038 | 0.286093 | Cerrado_vs_East_Amazon | Cerrado_vs_East_Amazon |
| ararauna | West_Amazon | crhr2 | FUNCG000000002793 | CM054377.1 | 105039900 | 105201503 | crhr2 | 22 | 1 | 0.125369 | 0.07766377272727273 | 0.125369 | Cerrado_vs_West_Amazon | Cerrado_vs_West_Amazon |
| ararauna | West_Amazon | IL1RAPL1 | FUNCG000000003787 | CM054378.1 | 45119475 | 45334894 | IL1RAPL1 | 27 | 11 | 0.153865 | 0.103064 | 0.153865 | Cerrado_vs_West_Amazon | Cerrado_vs_West_Amazon |
| ararauna | West_A | FUNCG0000 | FUNCG0000 | CM054 | 736581 | 737326 | FUNCG00000 | 9 | 5 | 0.1336 | 0.11253511 | 0.1336 | Cerrado_vs_West_Amazon | Cerrado_vs_West_Amazon |

| | | | | | | | | | | | | | | |
|---|---|---|---|---|---|---|---|---|---|---|---|---|---|---|
| a | mazon | 0004031 | 0004031 | 378.1 | 37 | 72 | 004031 | | | 69 | 111111111 | 69 | | |
| ararauna | West_Amazon | BIVM | FUNCG000000004044 | CM054378.1 | 76643996 | 76656572 | BIVM | 6 | 1 | 0.131415 | 0.08861533333333334 | 0.131415 | Cerrado_vs_West_Amazon | Cerrado_vs_West_Amazon |
| ararauna | West_Amazon | POGLUT2 | FUNCG000000004045 | CM054378.1 | 76661382 | 76675815 | POGLUT2 | 6 | 3 | 0.149822 | 0.11668333333333333 | 0.149822 | Cerrado_vs_West_Amazon | Cerrado_vs_West_Amazon |
| ararauna | West_Amazon | TEX30 | FUNCG000000004046 | CM054378.1 | 76677871 | 76684080 | TEX30 | 6 | 4 | 0.149822 | 0.12351616666666668 | 0.149822 | Cerrado_vs_West_Amazon | Cerrado_vs_West_Amazon |
| ararauna | West_Amazon | FUNCG000000004103 | FUNCG000000004103 | CM054378.1 | 84687616 | 84697200 | FUNCG000000004103 | 6 | 6 | 0.211006 | 0.17598466666666668 | 0.211006 | Cerrado_vs_West_Amazon | Cerrado_vs_West_Amazon |
| ararauna | West_Amazon | FUNCG000000006016 | FUNCG000000006016 | CM054379.1 | 126215695 | 126216005 | FUNCG000000006016 | 3 | 3 | 0.16252 | 0.16252 | 0.16252 | Cerrado_vs_West_Amazon | Cerrado_vs_West_Amazon |
| ararauna | West_Amazon | Rcn1 | FUNCG000000006495 | CM054380.1 | 29560755 | 29571134 | Rcn1 | 6 | 2 | 0.128263 | 0.108912 | 0.128263 | Cerrado_vs_West_Amazon | Cerrado_vs_West_Amazon |
| ararauna | West_Amazon | NUBPL | FUNCG000000006735 | CM054380.1 | 51438290 | 51526051 | NUBPL | 14 | 2 | 0.133546 | 0.06108121428571429 | 0.133546 | Cerrado_vs_West_Amazon | Cerrado_vs_West_Amazon |
| ararauna | West_Amazon | C14orf132 | FUNCG000000006902 | CM054380.1 | 65425854 | 65511164 | C14orf132 | 14 | 13 | 0.191208 | 0.15601342857142858 | 0.191208 | Cerrado_vs_West_Amazon | Cerrado_vs_West_Amazon |
| ararauun | West_A | FUNCG0000 | FUNCG0000 | CM054 | 798588 | 798919 | FUNCG00000 | 9 | 3 | 0.2358 | 0.09770655 | 0.2358 | Cerrado_vs_West_Amazon | Cerrado_vs_West_Amazon |

| | | | | | | | | | | | | | | |
|---|---|---|---|---|---|---|---|---|---|---|---|---|---|---|
| a | mazon | 0007069 | 0007069 | 380.1 | 15 | 33 | 007069 | | | 44 | 555555556 | 44 | | |
| ararauna | West_Amazon | GLUD1 | FUNCG000000007869 | CM054381.1 | 37913219 | 37943091 | GLUD1 | 8 | 5 | 0.144715 | 0.1291155 | 0.144715 | Cerrado_vs_West_Amazon | Cerrado_vs_West_Amazon |
| ararauna | West_Amazon | FUNCG000000007870 | FUNCG000000007870 | CM054381.1 | 37947009 | 37950075 | FUNCG000000007870 | 6 | 2 | 0.143127 | 0.1178188333333333 | 0.143127 | Cerrado_vs_West_Amazon | Cerrado_vs_West_Amazon |
| ararauna | West_Amazon | G2e3 | FUNCG000000011150 | CM054385.1 | 21812417 | 21822734 | G2e3 | 6 | 1 | 0.126779 | 0.0822426666666667 | 0.126779 | Cerrado_vs_West_Amazon | Cerrado_vs_West_Amazon |
| ararauna | West_Amazon | G2e3 | FUNCG000000011151 | CM054385.1 | 21827690 | 21858680 | G2e3 | 8 | 1 | 0.126779 | 0.0805305 | 0.126779 | Cerrado_vs_West_Amazon | Cerrado_vs_West_Amazon |
| ararauna | West_Amazon | FUNCG000000011603 | FUNCG000000011603 | CM054387.1 | 534279 | 534496 | FUNCG000000011603 | 5 | 1 | 0.141804 | 0.0765214 | 0.141804 | Cerrado_vs_West_Amazon | Cerrado_vs_West_Amazon |
| chloropterus | Cerrado | SLIT2 | FUNCG000000000217 | CM054376.1 | 19073205 | 19356746 | SLIT2 | 66 | 5 | 0.218525 | 0.08447628787878787 | 0.218525 | Cerrado_vs_East_Amazon;Cerrado_vs_West_Amazon | Cerrado_vs_West_Amazon |
| chloropterus | Cerrado | FUNCG000000001140 | FUNCG000000001140 | CM054376.1 | 93134660 | 93152908 | FUNCG000000001140 | 14 | 2 | 0.240456 | 0.08568542857142857 | 0.240456 | Cerrado_vs_East_Amazon;Cerrado_vs_West_Amazon | Cerrado_vs_West_Amazon |
| chloropterus | Cerrado | APPL2 | FUNCG000000001141 | CM054376.1 | 93170576 | 93206538 | APPL2 | 16 | 4 | 0.240456 | 0.110626875 | 0.240456 | Cerrado_vs_East_Amazon;Cerrado_vs_West_Amazon | Cerrado_vs_West_Amazon |
| chloropt | Cerrad | STK3 | FUNCG0000 | CM054 | 224443 | 226649 | STK3 | 48 | 4 | 0.2265 | 0.07418066 | 0.2265 | Cerrado_vs_East_Amazon;C | Cerrado_vs_West_Amazon |

| | | | | | | | | | | | | | | |
|---|---|---|---|---|---|---|---|---|---|---|---|---|---|---|
| erus | o | | 0002087 | 377.1 | 89 | 48 | | | | 14 | 666666667 | 14 | errado_vs_West_Amazon | |
| chloropterus | Cerrado | IKZF1 | FUNCG000000002513 | CM054377.1 | 73479286 | 73571204 | IKZF1 | 30 | 6 | 0.215468 | 0.1270724 | 0.215468 | Cerrado_vs_East_Amazon;Cerrado_vs_West_Amazon | Cerrado_vs_East_Amazon |
| chloropterus | Cerrado | SPATA48 | FUNCG000000002514 | CM054377.1 | 73599976 | 73624945 | SPATA48 | 16 | 3 | 0.215444 | 0.1329163125 | 0.215444 | Cerrado_vs_East_Amazon;Cerrado_vs_West_Amazon | Cerrado_vs_East_Amazon |
| chloropterus | Cerrado | CDH12 | FUNCG000000002555 | CM054377.1 | 81373299 | 81540066 | CDH12 | 44 | 6 | 0.200792 | 0.09327781818181818 | 0.200792 | Cerrado_vs_East_Amazon;Cerrado_vs_West_Amazon | Cerrado_vs_West_Amazon |
| chloropterus | Cerrado | Mc4r | FUNCG000000002562 | CM054377.1 | 86197539 | 86198534 | Mc4r | 10 | 2 | 0.204478 | 0.110499 | 0.204478 | Cerrado_vs_East_Amazon;Cerrado_vs_West_Amazon | Cerrado_vs_East_Amazon |
| chloropterus | Cerrado | PHACTR1 | FUNCG000000002636 | CM054377.1 | 93313623 | 93466437 | PHACTR1 | 40 | 2 | 0.170738 | 0.09136815 | 0.170738 | Cerrado_vs_East_Amazon;Cerrado_vs_West_Amazon | Cerrado_vs_East_Amazon |
| chloropterus | Cerrado | LSM5 | FUNCG000000002684 | CM054377.1 | 98115528 | 98119358 | LSM5 | 10 | 2 | 0.17873 | 0.1056602 | 0.17873 | Cerrado_vs_East_Amazon;Cerrado_vs_West_Amazon | Cerrado_vs_East_Amazon |
| chloropterus | Cerrado | Cnksr2 | FUNCG000000003807 | CM054378.1 | 48481409 | 48703243 | Cnksr2 | 54 | 4 | 0.224311 | 0.091613 | 0.224311 | Cerrado_vs_East_Amazon;Cerrado_vs_West_Amazon | Cerrado_vs_West_Amazon |
| chloropterus | Cerrado | Slc5a7 | FUNCG000000003966 | CM054378.1 | 66776888 | 66796983 | Slc5a7 | 14 | 1 | 0.197874 | 0.09218428571428572 | 0.197874 | Cerrado_vs_East_Amazon;Cerrado_vs_West_Amazon | Cerrado_vs_West_Amazon |
| chloropt | Cerrad | CCDC138 | FUNCG0000 | CM054 | 670562 | 670930 | CCDC138 | 18 | 8 | 0.2999 | 0.12910411 | 0.2999 | Cerrado_vs_East_Amazon;C | Cerrado_vs_West_Amazon |

| | | | | | | | | | | | | | | |
|---|---|---|---|---|---|---|---|---|---|---|---|---|---|---|
| erus | o | | 0003970 | 378.1 | 85 | 48 | | | | 36 | 111111111 | 36 | errado_vs_West_Amazon | |
| chloropterus | Cerrado | EDAR | FUNCG000000003971 | CM054378.1 | 67107567 | 67170354 | EDAR | 24 | 7 | 0.297731 | 0.1226321666666667 | 0.297731 | Cerrado_vs_East_Amazon;Cerrado_vs_West_Amazon | Cerrado_vs_West_Amazon |
| chloropterus | Cerrado | SH3RF3 | FUNCG000000003973 | CM054378.1 | 67213364 | 67466001 | SH3RF3 | 60 | 7 | 0.297731 | 0.09758745 | 0.297731 | Cerrado_vs_East_Amazon;Cerrado_vs_West_Amazon | Cerrado_vs_West_Amazon |
| chloropterus | Cerrado | MYO16 | FUNCG000000004030 | CM054378.1 | 73218749 | 73521412 | MYO16 | 72 | 4 | 0.195518 | 0.07220729166666667 | 0.195518 | Cerrado_vs_East_Amazon;Cerrado_vs_West_Amazon | Cerrado_vs_West_Amazon |
| chloropterus | Cerrado | NALCN | FUNCG000000004051 | CM054378.1 | 77411582 | 77644676 | NALCN | 56 | 10 | 0.214139 | 0.12142669642857143 | 0.214139 | Cerrado_vs_East_Amazon;Cerrado_vs_West_Amazon | Cerrado_vs_East_Amazon;Cerrado_vs_West_Amazon |
| chloropterus | Cerrado | DIAPH3 | FUNCG000000004151 | CM054378.1 | 98060936 | 98274157 | DIAPH3 | 52 | 24 | 0.276369 | 0.12187407692307692 | 0.276369 | Cerrado_vs_East_Amazon;Cerrado_vs_West_Amazon | Cerrado_vs_East_Amazon |
| chloropterus | Cerrado | ADGB | FUNCG000000005366 | CM054379.1 | 84604832 | 84717571 | ADGB | 32 | 5 | 0.239924 | 0.1005865 | 0.239924 | Cerrado_vs_East_Amazon;Cerrado_vs_West_Amazon | Cerrado_vs_East_Amazon |
| chloropterus | Cerrado | RAB32 | FUNCG000000005367 | CM054379.1 | 84727764 | 84747110 | RAB32 | 14 | 7 | 0.239924 | 0.13777007142857142 | 0.239924 | Cerrado_vs_East_Amazon;Cerrado_vs_West_Amazon | Cerrado_vs_East_Amazon |
| chloropterus | Cerrado | GRM1 | FUNCG000000005368 | CM054379.1 | 84789142 | 84970581 | GRM1 | 48 | 1 | 0.17438 | 0.0500459375 | 0.17438 | Cerrado_vs_East_Amazon;Cerrado_vs_West_Amazon | Cerrado_vs_East_Amazon |
| chloropt | Cerrad | SMYD3 | FUNCG0000 | CM054 | 986258 | 990347 | SMYD3 | 92 | 1 | 0.1981 | 0.06530042 | 0.1981 | Cerrado_vs_East_Amazon;C | Cerrado_vs_West_Amazon |

| | | | | | | | | | | | | | | |
|---|---|---|---|---|---|---|---|---|---|---|---|---|---|---|
| erus | o | | 0005500 | 379.1 | 21 | 07 | | | | 44 | 391304347 | 44 | errado_vs_West_Amazon | |
| chloropterus | Cerrado | ARL14EP | FUNCG000000006489 | CM054380.1 | 28702804 | 28706015 | ARL14EP | 10 | 1 | 0.171576 | 0.0941306 | 0.171576 | Cerrado_vs_East_Amazon;Cerrado_vs_West_Amazon | Cerrado_vs_East_Amazon |
| chloropterus | Cerrado | MPPED2 | FUNCG000000006490 | CM054380.1 | 28743486 | 28844481 | MPPED2 | 30 | 13 | 0.229826 | 0.1347433333333333 | 0.229826 | Cerrado_vs_East_Amazon;Cerrado_vs_West_Amazon | Cerrado_vs_East_Amazon |
| chloropterus | Cerrado | SNX4 | FUNCG000000008086 | CM054381.1 | 59655138 | 59689502 | SNX4 | 16 | 5 | 0.232856 | 0.1084586875 | 0.232856 | Cerrado_vs_East_Amazon;Cerrado_vs_West_Amazon | Cerrado_vs_East_Amazon |
| chloropterus | Cerrado | CNTNAP5 | FUNCG000000008122 | CM054381.1 | 63924228 | 64216952 | CNTNAP5 | 68 | 17 | 0.234612 | 0.11747010294117646 | 0.234612 | Cerrado_vs_East_Amazon;Cerrado_vs_West_Amazon | Cerrado_vs_East_Amazon;Cerrado_vs_West_Amazon |
| chloropterus | Cerrado | HIF1AN | FUNCG000000008229 | CM054381.1 | 69043935 | 69051106 | HIF1AN | 12 | 6 | 0.310903 | 0.1612596666666666 | 0.310903 | Cerrado_vs_East_Amazon;Cerrado_vs_West_Amazon | Cerrado_vs_East_Amazon |
| chloropterus | Cerrado | FUNCG000000008230 | FUNCG000000008230 | CM054381.1 | 69051370 | 69052802 | FUNCG000000008230 | 10 | 5 | 0.310903 | 0.1668274 | 0.310903 | Cerrado_vs_East_Amazon;Cerrado_vs_West_Amazon | Cerrado_vs_East_Amazon |
| chloropterus | Cerrado | NDUFB8 | FUNCG000000008231 | CM054381.1 | 69053124 | 69056688 | NDUFB8 | 10 | 5 | 0.310903 | 0.1668274 | 0.310903 | Cerrado_vs_East_Amazon;Cerrado_vs_West_Amazon | Cerrado_vs_East_Amazon |
| chloropterus | Cerrado | SEC31B | FUNCG000000008232 | CM054381.1 | 69062351 | 69092198 | SEC31B | 16 | 7 | 0.310903 | 0.1455733125 | 0.310903 | Cerrado_vs_East_Amazon;Cerrado_vs_West_Amazon | Cerrado_vs_East_Amazon |
| chloropt | Cerrad | RABGAP1L | FUNCG0000 | CM054 | 436646 | 439090 | RABGAP1L | 58 | 5 | 0.2164 | 0.06975674 | 0.2164 | Cerrado_vs_East_Amazon;C | Cerrado_vs_West_Amazon |

| | | | | | | | | | | | | | | |
|---|---|---|---|---|---|---|---|---|---|---|---|---|---|---|
| erus | o | | 0009099 | 382.1 | 61 | 67 | | | | 98 | 137931034 | 98 | errado_vs_West_Amazon | |
| chloropterus | Cerrado | RBBP6 | FUNCG000000014155 | CM054395.1 | 91243 | 105918 | RBBP6 | 6 | 1 | 0.216404 | 0.0682801666666667 | 0.216404 | Cerrado_vs_East_Amazon;Cerrado_vs_West_Amazon | Cerrado_vs_West_Amazon |
| chloropterus | Cerrado | RBBP6 | FUNCG000000014156 | CM054395.1 | 133050 | 144044 | RBBP6 | 9 | 3 | 0.216404 | 0.1007100000000001 | 0.216404 | Cerrado_vs_East_Amazon;Cerrado_vs_West_Amazon | Cerrado_vs_West_Amazon |
| chloropterus | East_Amazon | FUNCG000000001478 | FUNCG000000001478 | CM054376.1 | 121437852 | 121440271 | FUNCG000000001478 | 6 | 3 | 0.167517 | 0.1614131666666666 | 0.167517 | Cerrado_vs_East_Amazon | Cerrado_vs_East_Amazon |
| chloropterus | East_Amazon | GABBR2 | FUNCG000000002469 | CM054377.1 | 65111796 | 65572497 | GABBR2 | 51 | 1 | 0.166038 | 0.08292945098039216 | 0.166038 | Cerrado_vs_East_Amazon | Cerrado_vs_East_Amazon |
| chloropterus | East_Amazon | DIAPH3 | FUNCG000000004151 | CM054378.1 | 98060936 | 98274157 | DIAPH3 | 26 | 24 | 0.276369 | 0.20495530769230077 | 0.276369 | Cerrado_vs_East_Amazon | Cerrado_vs_East_Amazon |
| chloropterus | East_Amazon | FBXO25 | FUNCG000000004951 | CM054379.1 | 35362335 | 35388944 | FBXO25 | 7 | 3 | 0.175231 | 0.11651257142857142 | 0.175231 | Cerrado_vs_East_Amazon | Cerrado_vs_East_Amazon |
| chloropterus | East_Amazon | TMEM181 | FUNCG000000005312 | CM054379.1 | 78900466 | 78933032 | TMEM181 | 8 | 4 | 0.185512 | 0.135746625 | 0.185512 | Cerrado_vs_East_Amazon | Cerrado_vs_East_Amazon |
| chloropterus | East_Amazon | Tulp4 | FUNCG000000005313 | CM054379.1 | 78954334 | 79085894 | Tulp4 | 18 | 3 | 0.185512 | 0.1346976111111112 | 0.185512 | Cerrado_vs_East_Amazon | Cerrado_vs_East_Amazon |
| chloropt | East_A | Kif26b | FUNCG0000 | CM054 | 983916 | 986060 | Kif26b | 26 | 16 | 0.2232 | 0.17005861 | 0.2232 | Cerrado_vs_East_Amazon | Cerrado_vs_East_Amazon |

| | | | | | | | | | | | | | | |
|---|---|---|---|---|---|---|---|---|---|---|---|---|---|---|
| erus | mazon | | 0005499 | 379.1 | 75 | 74 | | | | 47 | 53846154 | 47 | | |
| chloropterus | East_Amazon | SHISA9 | FUNCG000000006361 | CM054380.1 | 17618990 | 17802437 | SHISA9 | 24 | 2 | 0.188313 | 0.08300441666666666 | 0.188313 | Cerrado_vs_East_Amazon | Cerrado_vs_East_Amazon |
| chloropterus | East_Amazon | LUZP2 | FUNCG000000006474 | CM054380.1 | 26903364 | 27093115 | LUZP2 | 24 | 6 | 0.184153 | 0.116367 | 0.184153 | Cerrado_vs_East_Amazon | Cerrado_vs_East_Amazon |
| chloropterus | East_Amazon | AKAP6 | FUNCG000000006737 | CM054380.1 | 51781308 | 52019062 | AKAP6 | 28 | 1 | 0.162095 | 0.10360685714285714 | 0.162095 | Cerrado_vs_East_Amazon | Cerrado_vs_East_Amazon |
| chloropterus | East_Amazon | RNLS | FUNCG000000007980 | CM054381.1 | 48035761 | 48107298 | RNLS | 12 | 5 | 0.230846 | 0.15713258333333333 | 0.230846 | Cerrado_vs_East_Amazon | Cerrado_vs_East_Amazon |
| chloropterus | West_Amazon | PDS5A | FUNCG000000000265 | CM054376.1 | 26579529 | 26663355 | PDS5A | 14 | 5 | 0.22501 | 0.15398671428571428 | 0.22501 | Cerrado_vs_West_Amazon | Cerrado_vs_West_Amazon |
| chloropterus | West_Amazon | ATXN7L1 | FUNCG000000001628 | CM054376.1 | 138798240 | 138912157 | ATXN7L1 | 17 | 5 | 0.228951 | 0.1329904705882353 | 0.228951 | Cerrado_vs_West_Amazon | Cerrado_vs_West_Amazon |
| chloropterus | West_Amazon | TBC1D22A | FUNCG000000001662 | CM054376.1 | 141185955 | 141366888 | TBC1D22A | 23 | 6 | 0.252302 | 0.1252377391304348 | 0.252302 | Cerrado_vs_West_Amazon | Cerrado_vs_West_Amazon |
| chloropterus | West_Amazon | Slc5a7 | FUNCG000000003966 | CM054378.1 | 66776888 | 66796983 | Slc5a7 | 7 | 1 | 0.197874 | 0.14646528571428571 | 0.197874 | Cerrado_vs_West_Amazon | Cerrado_vs_West_Amazon |
| chloropt | West_A | LIMS1 | FUNCG0000 | CM054 | 669981 | 670171 | LIMS1 | 7 | 4 | 0.2999 | 0.19692014 | 0.2999 | Cerrado_vs_West_Amazon | Cerrado_vs_West_Amazon |

| | | | | | | | | | | | | | | |
|---|---|---|---|---|---|---|---|---|---|---|---|---|---|---|
| erus | mazon | | 0003968 | 378.1 | 29 | 53 | | | | 36 | 285714285 | 36 | | |
| chloropterus | West_Amazon | Ranbp2 | FUNCG000000003969 | CM054378.1 | 67022268 | 67055134 | Ranbp2 | 8 | 8 | 0.299936 | 0.248300375 | 0.299936 | Cerrado_vs_West_Amazon | Cerrado_vs_West_Amazon |
| chloropterus | West_Amazon | CCDC138 | FUNCG000000003970 | CM054378.1 | 67056285 | 67093048 | CCDC138 | 9 | 8 | 0.299936 | 0.229426444444443 | 0.299936 | Cerrado_vs_West_Amazon | Cerrado_vs_West_Amazon |
| chloropterus | West_Amazon | EDAR | FUNCG000000003971 | CM054378.1 | 67107567 | 67170354 | EDAR | 12 | 7 | 0.297731 | 0.204492416666666668 | 0.297731 | Cerrado_vs_West_Amazon | Cerrado_vs_West_Amazon |
| chloropterus | West_Amazon | SH3RF3 | FUNCG000000003973 | CM054378.1 | 67213364 | 67466001 | SH3RF3 | 30 | 7 | 0.297731 | 0.149051433333333334 | 0.297731 | Cerrado_vs_West_Amazon | Cerrado_vs_West_Amazon |
| chloropterus | West_Amazon | Itgbl1 | FUNCG000000004050 | CM054378.1 | 77260248 | 77392864 | Itgbl1 | 18 | 6 | 0.285587 | 0.1714815 | 0.285587 | Cerrado_vs_West_Amazon | Cerrado_vs_West_Amazon |
| chloropterus | West_Amazon | NALCN | FUNCG000000004051 | CM054378.1 | 77411582 | 77644676 | NALCN | 28 | 4 | 0.214139 | 0.120889357142857 15 | 0.214139 | Cerrado_vs_West_Amazon | Cerrado_vs_West_Amazon |
| chloropterus | West_Amazon | SMYD3 | FUNCG000000005500 | CM054379.1 | 98625821 | 99034707 | SMYD3 | 46 | 1 | 0.198144 | 0.047450826086956 52 | 0.198144 | Cerrado_vs_West_Amazon | Cerrado_vs_West_Amazon |
| chloropterus | West_Amazon | AKAP6 | FUNCG000000006737 | CM054380.1 | 51781308 | 52019062 | AKAP6 | 28 | 11 | 0.337587 | 0.196275285714285 7 | 0.337587 | Cerrado_vs_West_Amazon | Cerrado_vs_West_Amazon |
| chloropt | West_A | RABGAP1L | FUNCG0000 | CM054 | 436646 | 439090 | RABGAP1L | 29 | 5 | 0.2164 | 0.10637158 | 0.2164 | Cerrado_vs_West_Amazon | Cerrado_vs_West_Amazon |

| | | | | | | | | | | | | |
|---|---|---|---|---|---|---|---|---|---|---|---|---|
| erus | mazon | | 0009099 | 382.1 | 61 | 67 | | | | 98 | 620689655 | 98 |

**Table S16.** Complete list of genes overlapping $F_{ST}$ outlier windows per comparison per species.

| *Ara ararauna* - Cerrado vs East Amazon | *Ara ararauna* - Cerrado vs West Amazon | *Ara chloropterus* - Cerrado vs East Amazon | *Ara chloropterus* - Cerrado vs West Amazon |
|---|---|---|---|
| ACTA2 | ABCC4 | ADGB | ABCA13 |
| ADAMTS1 | ABCC5 | ADGRF5 | AHSA1 |
| ADAMTS5 | ACTL6B | ADHFE1 | AKAP6 |
| AHNAK2 | ADK | AKAP12 | AMER2 |
| AKAP7 | ALPI | AKAP17A | ANK3 |
| ANAPC10 | ALX1 | AKAP6 | ANKRD35 |
| ANKRD50 | ANO4 | ALDH1A2 | APPL2 |
| APP | ARF4 | ARHGAP11A | ARGLU1 |
| ARF4 | Alpi | ARHGAP5 | ARHGAP5 |
| ARHGAP10 | Antxr1 | ARL14EP | ARHGEF2 |
| ASTN1 | Atp6v1c2 | ASMT | ARHGEF5 |
| Amotl2 | BCL11B | ASMTL | ASB11 |
| Atp5pf | BDKRB1 | ASNS | ASB9 |
| BOD1L1 | BDKRB2 | BOC | ATAD1 |

| | | | |
|---|---|---|---|
| BRINP2 | BDMD1 | Baalc | ATXN7L1 |
| BTAF1 | BEGAIN | C1GALT1 | Acaa1a |
| CAMK2G | BHLHE22 | CCDC170 | Ace2 |
| CCDC178 | BICC1 | CDKAL1 | CAMK1D |
| CCDC28A | BIVM | CEFIP | CASP7 |
| CCDC50 | BOD1L1 | CFAP206 | CCDC138 |
| CCNL1 | Bicdl1 | CFAP44 | CCN2 |
| CCSER2 | C14orf132 | CGA | CDC5L |
| CCT8 | C4orf33 | CHST9 | CDH12 |
| CDH18 | C8orf88 | CIBAR1 | CDH22 |
| CDK6 | CAMK2G | CLPTM1L | CEP290 |
| CFAP47 | CAPN10 | CNTNAP5 | CHRM2 |
| CGNL1 | CARTPT | COL1A2 | CIBAR1 |
| CHODL | CCDC50 | CPB1 | CIPC |
| CKAP5 | CCDC93 | CRH | CLTRN |
| CLASP1 | CCNK | CSF2RA | CNTNAP5 |
| CLYBL | CCSER2 | Cd40 | CPNE8 |
| CSGALNACT1 | CELF5 | DCBLD1 | CRYZL1 |
| CSTF3 | CHODL | DCLRE1C | CTPS2 |
| Cacna2d3 | CHRND | DCUN1D5 | CWC22 |
| DENND6A | CHRNG | DIAPH3 | CXCL12 |

| | | | |
|---|---|---|---|
| DNAH5 | CHUK | DIO2 | CYB5R3 |
| DNAJC24 | CIBAR1 | DLEU7 | Cd40 |
| DNM3 | CIT | DMTF1 | Cnksr2 |
| DPYS | CLASP1 | DNAJC1 | Comt |
| DYNLT2B | CLGN | DTNA | DAAM2 |
| Dnah8 | CLYBL | Dnajc15 | DCAF5 |
| Dusp11 | CPED1 | Drgx | DCBLD1 |
| EDNRA | CREBBP | Dync2h1 | DCUN1D5 |
| EGF | CSGALNACT1 | EAPP | DECR2 |
| EIF4A2 | CYP2J2 | EDN2 | DERL2 |
| ELOVL6 | CYP2J4 | ELP4 | DIO2 |
| ELP4 | CYP7B1 | ENOX1 | DIO3 |
| EPB41L3 | Cop1 | ESR1 | DLEC1 |
| EURL | Cops7b | FAM98B | DONSON |
| FAM180A | Cwf19l1 | FBXO25 | DPP6 |
| FAS | Cysltr2 | FERD3L | Ddx27 |
| FASLG | DCT | FUNCG00000000677 | Ddx51 |
| FETUB | DENND6A | FUNCG00000000678 | Dync1h1 |
| FUNCG00000001123 | DGKB | FUNCG00000001478 | Dync2h1 |
| FUNCG00000002286 | DIO3 | FUNCG00000002674 | Dyrk2 |
| FUNCG00000002304 | DIS3L2 | FUNCG00000003029 | EDAR |

| | | | |
|---|---|---|---|
| FUNCG00000003620 | DMTF1 | FUNCG00000003051 | EFCC1 |
| FUNCG00000003621 | DNAAF2 | FUNCG00000003620 | EFNB2 |
| FUNCG00000003622 | DNAH1 | FUNCG00000003621 | EIF3I |
| FUNCG00000003624 | DNAJB4 | FUNCG00000003622 | ENOX1 |
| FUNCG00000003625 | DSL1 | FUNCG00000003875 | ENPEP |
| FUNCG00000003626 | DTD2 | FUNCG00000004040 | ENPP1 |
| FUNCG00000003635 | DVL3 | FUNCG00000004834 | ENPP3 |
| FUNCG00000003873 | Ddx18 | FUNCG00000005330 | EPB42 |
| FUNCG00000004209 | Dnah1 | FUNCG00000006831 | ERCC6 |
| FUNCG00000004256 | EFL1 | FUNCG00000006832 | ESR1 |
| FUNCG00000004824 | EIF2B5 | FUNCG00000006834 | EXD2 |
| FUNCG00000004825 | EIF3M | FUNCG00000007828 | Egln3 |
| FUNCG00000005288 | ELP4 | FUNCG00000008230 | Elmo2 |
| FUNCG00000008135 | ENKUR | FUNCG00000011588 | FAM13C |
| FUNCG00000008897 | EPB41L3 | FUNCG00000013793 | FAM167B |
| FUNCG00000009050 | EPHA4 | FUNCG00000013794 | FBXO8 |
| GABPA | EPHB4 | FUNCG00000013795 | FUNCG00000000571 |
| GYG2 | ERLIN1 | FUNCG00000013796 | FUNCG00000000572 |
| Gabrr3 | ESYT2 | FUNCG00000013797 | FUNCG00000001123 |
| Glp1r | EXOSC3 | FUNCG00000013798 | FUNCG00000001140 |
| Grm7 | Ecel1 | FUNCG00000013799 | FUNCG00000001381 |

| | | | |
|---|---|---|---|
| HGSNAT | Eif4e2 | FUNCG00000013800 | FUNCG00000001521 |
| IFNGR1 | Eif5 | FUNCG00000013801 | FUNCG00000002666 |
| IL20RA | FBLN1 | FUNCG00000013802 | FUNCG00000002830 |
| IL22RA2 | FCHO1 | FUNCG00000013803 | FUNCG00000003010 |
| IMMP1L | FMO1 | FUNCG00000013804 | FUNCG00000003029 |
| INTS10 | FNDC3A | FUNCG00000013805 | FUNCG00000003033 |
| IPMK | FUNCG00000001123 | FUNCG00000013806 | FUNCG00000003673 |
| IPO5 | FUNCG00000001478 | FUNCG00000013807 | FUNCG00000003972 |
| Itgbl1 | FUNCG00000002238 | FUNCG00000013808 | FUNCG00000004040 |
| JAM2 | FUNCG00000002239 | FUNCG00000013809 | FUNCG00000004830 |
| KDM5A | FUNCG00000002240 | FUNCG00000013810 | FUNCG00000005941 |
| KL | FUNCG00000002364 | FUNDC1 | FUNCG00000006813 |
| KLHL14 | FUNCG00000002763 | FZD1 | FUNCG00000006829 |
| KNG1 | FUNCG00000002998 | FZD6 | FUNCG00000006830 |
| LIN54 | FUNCG00000003620 | Fam107b | FUNCG00000006831 |
| LPP | FUNCG00000003621 | Fmn2 | FUNCG00000006832 |
| LRP12 | FUNCG00000003649 | GABBR2 | FUNCG00000006834 |
| LTN1 | FUNCG00000004031 | GALNT18 | FUNCG00000006938 |
| Lipm | FUNCG00000004057 | GJB7 | FUNCG00000007602 |
| MAP3K5 | FUNCG00000004087 | GNAL | FUNCG00000007692 |
| MAP7 | FUNCG00000004088 | GOPC | FUNCG00000008031 |

| | | | |
|---|---|---|---|
| MASP1 | FUNCG00000004101 | GPR158 | FUNCG00000009653 |
| MRPL39 | FUNCG00000004102 | GRM1 | FUNCG00000010253 |
| MRTFB | FUNCG00000004103 | GTSF1 | FUNCG00000011753 |
| MSL2 | FUNCG00000006016 | Gpr158 | FUNCG00000013793 |
| MYZAP | FUNCG00000006913 | HDAC9 | FUNCG00000013794 |
| Map3k7cl | FUNCG00000006914 | HIF1AN | FUNCG00000013795 |
| Marchf4 | FUNCG00000006915 | HIPK2 | FUNCG00000013796 |
| Mcf2l | FUNCG00000006954 | HIVEP3 | FUNCG00000013797 |
| NALCN | FUNCG00000007069 | HPCAL1 | FUNCG00000013798 |
| NCAM2 | FUNCG00000007774 | HSPA14 | FUNCG00000013799 |
| NDFIP2 | FUNCG00000007843 | ICOSLG | FUNCG00000013800 |
| NPY | FUNCG00000007845 | IKZF1 | FUNCG00000013801 |
| OR14J1 | FUNCG00000007870 | ILDR1 | FUNCG00000013802 |
| PANK1 | FUNCG00000007871 | ITPRID2 | FUNCG00000013803 |
| PCCB | FUNCG00000007881 | IYD | FUNCG00000013804 |
| PCMTD1 | FUNCG00000008061 | Inpp4b | FUNCG00000013805 |
| PCYT1A | FUNCG00000008198 | JMJD1C | FUNCG00000013806 |
| PDGFD | FUNCG00000009024 | KDM6A | FUNCG00000013807 |
| PIGC | FUNCG00000010559 | KHDRBS2 | FUNCG00000013808 |
| POGLUT2 | FUNCG00000011603 | KIF6 | FUNCG00000013809 |
| POMK | FUNCG00000011604 | Kif26b | FUNCG00000013810 |

| | | | |
|---|---|---|---|
| PPP2R3A | FUNCG00000012598 | LSM5 | FUNCG00000014800 |
| PRMT9 | FUNCG00000012599 | LUZP2 | FUNCG00000014801 |
| Pex7 | FUNCG00000015159 | Lpcat1 | FUNCG00000015129 |
| Pim1 | Fubp1 | MAP1LC3C | FUNCG00000015236 |
| QRICH2 | G2e3 | MARCHF7 | Fgf14 |
| RASAL2 | GALNT13 | MEIG1 | GABBR2 |
| RBBP6 | GCN1 | MEP1A | GINM1 |
| RBM26 | GIPC2 | MINDY4 | GOPC |
| REPS1 | GLUD1 | MLLT10 | GRPR |
| RFC4 | GNA11 | MMP9 | HDAC1 |
| RHAG | GNG11 | MPPE1 | HSP90AA1 |
| RPL37A | GPC6 | MPPED2 | ISM2 |
| RWDD2B | GRID1 | MTMR6 | ITPR2 |
| RYK | GRM8 | MYBL1 | ITSN1 |
| Rcn1 | Gdf10 | Mc4r | Irf2bpl |
| SEC16B | Grm7 | Mrps9 | Itgbl1 |
| SEC31A | Gzmk | NAA15 | KATNA1 |
| SFTPA1 | HEATR5A | NALCN | KCNF1 |
| SIM2 | HGSNAT | NDUFA12 | KCNK16 |
| SLC13A4 | HPCAL1 | NDUFB8 | KDM5A |
| SLC35D3 | HSD11B1L | NFKBIZ | KIF6 |

| | | | |
|---|---|---|---|
| SLC51A | IL1RAPL1 | NPSR1 | KYNU |
| SLC6A13 | INTS10 | NRBF2 | Kcnc1 |
| SLITRK5 | Itgbl1 | NUP107 | LATS1 |
| SST | KCNF1 | Ncoa5 | LCK |
| ST18 | KCNH5 | ODC1 | LIMS1 |
| STAMBPL1 | KCNMA1 | OPRM1 | LRP11 |
| STARD13 | KDM5A | PACRG | MEI4 |
| STMN2 | KIF1C | PAX6 | METTL21C |
| SUCO | KIFAP3 | PCDH8 | MIF |
| Sh2d4a | KLHDC1 | PDE1A | MINPP1 |
| Slmap | LPP | PDIA5 | MLLT10 |
| Sorbs1 | LRR1 | PHACTR1 | MROH7 |
| TAGAP | Lss | PRSS12 | MRPS6 |
| TEX30 | MAP1S | Piezo2 | MTHFD1L |
| TM9SF2 | MAT1A | Ppp1r14c | MTMR6 |
| TMEM200C | MCPH1 | Ppp1r1c | MTMR9L |
| TMPRSS15 | MEPCE | Prim2 | MYO16 |
| Tagap | METTL21C | QtsA-12155 | Map3k3 |
| Tcf12 | MEX3D | RAB32 | Mrps9 |
| USP16 | MGAT2 | RAP1B | NALCN |
| Xrcc5 | MGAT4C | RASGRP1 | NALF1 |

| | | | |
|---|---|---|---|
| ZIC1 | MPPED2 | RGS17 | NBEA |
| ccdc77 | MYOZ1 | RGS7 | NME4 |
| cyyr1 | Mmp16 | RMND1 | NOBOX |
| smarcal1 | NCAM2 | RNLS | NPAS3 |
| zic4 | NCLN | ROS1 | NRAP |
| | NDST2 | RRS1 | NRBP2 |
| | NECAB1 | RUNX2 | NUP43 |
| | NEK10 | Rbm12b1 | Ncoa5 |
| | NEXN | Rcn1 | Nup58 |
| | NOL10 | SCG5 | OLA1 |
| | NTS | SEC31B | OPN5 |
| | NUBPL | SHISA9 | OR14J1 |
| | NUDT3 | SLC12A5 | OSGEP |
| | ODC1 | SLC25A6 | PAPSS2 |
| | OR14J1 | SLC35A1 | PCMT1 |
| | PACSIN1 | SLC35E3 | PDS5A |
| | PAX6 | SMIM8 | PEX5L |
| | PCDH20 | SNX4 | PIGA |
| | PDIA6 | SNX6 | PITX2 |
| | PHYHIPL | SPATA48 | PLEKHG1 |

| | | | |
|---|---|---|---|
| | PIP4P2 | SPRED1 | PPFIBP1 |
| | PLAU | SPTSSA | PPP2R2C |
| | PLS1 | SYNE1 | PPP2R5C |
| | POGLUT2 | Sox4 | PREPL |
| | POLE2 | TAC1 | PRKG1 |
| | POMK | TDRD3 | PRKG2 |
| | POT1 | TGFBR2 | PTEN |
| | PREX2 | TMEM181 | PUF60 |
| | PRKG1 | TMEM243 | Phf14 |
| | PRRX1 | TMEM67 | Pim1 |
| | PRSS12 | TRIM55 | Plekhs1 |
| | PRSS56 | TUB | Pou3f4 |
| | PRTFDC1 | Tert | Ppil4 |
| | PRXL2A | Tmem106b | Ppp1r14c |
| | Pcca | Tulp4 | Ptprk |
| | Pim1 | Twist1 | RAB11FIP3 |
| | RAB35 | VIPR2 | RAB12 |
| | RASSF9 | VWA3B | RABGAP1L |
| | RBP3 | VWDE | RASGEF1B |
| | RIBC2 | VXN | RBBP6 |

| | | | |
|---|---|---|---|
| | RNPEPL1 | WNT8B | RNF25 |
| | RPS10 | XYLT1 | RNF31 |
| | RPS29 | ZBTB2 | ROS1 |
| | RUNX1T1 | ZMIZ1 | RSBN1L |
| | Rcn1 | ZNF292 | RSPO2 |
| | S1pr4 | Zswim8 | Ranbp2 |
| | SAFB | fam124a | Rbm12b1 |
| | SCAF4 | myct1 | SAMD3 |
| | SCLT1 | nr2c1-a | SERGEF |
| | SETD3 | rnaseh2b | SGK1 |
| | SHLD2 | | SH3RF3 |
| | SLC6A13 | | SLC12A5 |
| | SLITRK6 | | SLC25A13 |
| | SOCS1 | | SLC2A11 |
| | SOD1 | | SLC6A13 |
| | SPAG7 | | SLCO5A1 |
| | SPDEF | | SLIT2 |
| | STMN2 | | SMARCB1 |
| | SUB1 | | SMPX |
| | SULT1B1 | | SMYD3 |

| | | | |
|---|---|---|---|
| | SYT15 | | SNRPB |
| | Scoc | | SPIDR |
| | Sh2d4a | | STK3 |
| | Slmap | | STK36 |
| | Smc1b | | SULF1 |
| | TAGAP | | SUPT3H |
| | TEX30 | | SYAP1 |
| | TFPI2 | | SYNE1 |
| | TFR2 | | Slc5a7 |
| | THNSL1 | | Son |
| | TLE5 | | Sptlc2 |
| | TP63 | | TBC1D22A |
| | TRPC1 | | TGM6 |
| | Tmprss11e | | TIAM1 |
| | Tpp2 | | TMEM67 |
| | Tyms | | TNRC6B |
| | UGT2A2 | | TTLL4 |
| | WASF3 | | TUBGCP3 |
| | Wdr25 | | TXLNG |
| | YES1 | | Tmem106b |

| | | | |
|---|---|---|---|
| | Ythdc1 | | Tpp2 |
| | ZBTB14 | | Ttc5 |
| | ZNF407 | | Txnrd2 |
| | ZNF488 | | VIPAS39 |
| | Zswim8 | | VIT |
| | ccdc77 | | VPREB3 |
| | ccdc85c | | VTG1 |
| | crhr2 | | VWDE |
| | lrrfip2 | | Vegfd |
| | micos13 | | Vgll2 |
| | ncapg2 | | WASHC4 |
| | slc12a9 | | WDR17 |
| | smim19 | | WDR20 |
| | tcf3 | | XPO6 |
| | | | XYLT1 |
| | | | Xylb |
| | | | Zdhhc22 |
| | | | bcs1l |
| | | | ccdc77 |
| | | | ccndbp1 |

| | | | |
|---|---|---|---|
| | | | cep44 |
| | | | mmp11 |
| | | | myct1 |

**Table S17.** Software tools: versions and sources used for all genome annotations

| **Software tool** | **Version** | **Source** |
|---|---|---|
| AGAT | v1.4 | https://github.com/NBISweden/AGAT |
| BUSCO | v5.8.3 | https://gitlab.com/ezlab/busco |
| BWA-MEM | v0.7.18 | https://github.com/lh3/bwa |
| DIAMOND | v2.1.8 | https://github.com/bbuchfink/diamond |
| EMBLmyGFF3 | v2.2 | https://github.com/NBISweden/EMBLmyGFF3 |
| EvidenceModeler | v2.1.0 | https://github.com/EVidenceModeler/EVidenceModeler |
| Funannotate | v1.8.17 | https://github.com/nextgenusfs/funannotate |
| GALBA | 1.0.9 | https://github.com/Gaius-Augustus/GALBA |
| gfastats | 1.3.6 | https://github.com/vgl-hub/gfastats |
| InterProScan | v5.47-82 | https://www.ebi.ac.uk/interpro/search/sequence/ |
| minimap2 | 2.28 | https://github.com/lh3/minimap2 |
| miniprot | 0.11-r234 | https://github.com/lh3/miniprot |
| RED | v2018.09.10 | https://github.com/BioinformaticsToolsmith/Red |
| samtools | 1.17 | https://github.com/samtools/samtools |

**Table S18.** Genome and annotation statistics. * BUSCO scores based on the aves BUSCO set using v5.7.1. C = complete [S = single copy, D = duplicated], F = fragmented, M = missing, n = number of orthologues in comparison (8338) **Number of genes annotated with a functional domain as found by InterProScan.

| | *Ara ararauna* | *Ara chloropterus* | *Ara severus* |
|---|---|---|---|
| # contigs | 353 | 459691 | 182905 |
| contig N50 (kb) | 38811 | 4.2 | 16.9 |
| BUSCO (genome) | C:98.7%[S:98.4%,D:0.3%],F:0.3%,M:1.0% | C:45.3%[S:45.2%,D:0.1%],F:28.6%,M:26.1% | C:73.8%[S:73.6%,D:0.2%],F:18.1%,M:8.1% |
| Number genes | 17033 | 26554 | 25175 |
| Number of genes with names | 14182 | 19156 | 19992 |
| Number of genes with functional domain** | 16222 | 24241 | 23649 |
| Mean mRNA length (bp) | 27708 | 6264 | 8670 |
| Mean CDS length (bp) | 1705 | 776 | 991 |
| BUSCO (proteins) | C:95.7%[S:95.3%,D:0.4%],F:0.6%,M:3.7% | C:39.3%[S:39.2%,D:0.1%],F:26.7%,M:34.0% | C:71.1%[S:70.8%,D:0.3%],F:17.9%,M:11.0% |

**Table S19**. Number of candidate genes evaluated in GO enrichment analysis. We show the total number of candidate genes ($F_{ST}$ outliers + LowH), and $F_{ST}$ outlier genes, as well as their respective number of background genes. In addition we show the number of those genes that exhibit GO terms.

| **Species** | **Comparison** | **Analysis** | **n_candidate_total** | **n_candidate_with_BP_GO** | **candidate_GO_fraction** | **n_background_total** | **n_background_with_BP_GO** | **background_GO_fraction** |
|---|---|---|---|---|---|---|---|---|
| *Ara ararauna* | Cerrado_vs_East_Amazon | FST outliers | 168 | 77 | 0.45833333333333333 | 14865 | 6431 | 0.4326269761183989 |
| *Ara ararauna* | Cerrado_vs_West_Amazon | FST outliers | 259 | 114 | 0.440154440154440016 | 14793 | 6411 | 0.43338065301155954 |
| *Ara chloropterus* | Cerrado_vs_East_Amazon | FST outliers | 211 | 75 | 0.355450236966682465 | 14913 | 6451 | 0.43257560517669147 |
| *Ara chloropterus* | Cerrado_vs_West_Amazon | FST outliers | 269 | 115 | 0.42750929368029740 | 14869 | 6435 | 0.43277960858161274 |
| **Species** | **Biome** | **Analysis** | **n_candidate_total** | **n_candidate_with_BP_GO** | **candidate_GO_fraction** | **n_background_total** | **n_background_with_BP_GO** | **background_GO_fraction** |
| ararauna | Cerrado | FST outliers + LowH | 37 | 18 | 0.4864864864864865 | 14865 | 6431 | 0.4326269761183989 |
| ararauna | East_Amazon | FST outliers + LowH | 9 | 4 | 0.4444444444444444 | 14865 | 6431 | 0.4326269761183989 |
| ararauna | West_Amazon | FST outliers + LowH | 17 | 3 | 0.17647058823529413 | 14792 | 6411 | 0.43340995132504057 |
| chloropterus | Cerrado | FST outliers + LowH | 33 | 16 | 0.48484848484848486 | 14914 | 6452 | 0.43261365160252113 |
| chloropterus | East_Amazon | FST outliers + LowH | 11 | 3 | 0.2727272727272727 | 14913 | 6451 | 0.43257560517669147 |
| chloropterus | West_Amazon | FST outliers + LowH | 14 | 5 | 0.35714285714285715 | 14869 | 6435 | 0.43277960858161274 |
